# RMeDPower2 for Biology: guiding the design, experimental structure and analyses of experiments generating repeated measures datasets

**DOI:** 10.1101/2022.07.18.500490

**Authors:** Min-Gyoung Shin, Naufa Amirani, Stephanie Lam, Noura Al Bistami, Edward Vertudes, Krishna Raja, Julia A. Kaye, Reuben Thomas, Steven Finkbeiner

## Abstract

Lack of experimental reproducibility has plagued efforts to understand biology at both basic biomedical and preclinical levels. The cause is often improperly powered experiments and the use of inadequate statistical tools. To overcome these problems, we developed RMeDPower2, a complete, user-friendly package of tools in R that will allow scientists that are not deeply familiar with statistical analyses to predict the scope and size of biological data they need when conducting experiments with a repeated measures design. RMeDPower2 is based on Generalized Linear Mixed Effects Models (GLMM), which are better suited to the statistical analysis of these experiments than ANOVA or t-tests. We illustrate the use of RMeDPower2 and compare it to t-test for power calculations, using our own pilot studies of iPSC-derived motor neurons (iMNs) from sporadic ALS (sALS) patients versus healthy controls. We report that sALS iMNs display reduced numbers of soma- emanating processes compared to control iMNs using RMeDPower2. We expect RMeDPower2 to find applications far beyond cell assays, from single-cell RNAseq experiments to brain slice electrophysiology or animal behavior.

**Motivation:** The lack of rigor and reproducibility in biomedical research has caused a crisis that has been highlighted in the popular literature and has become a focus for the National Institutes of Health^1–3^. It has been estimated that the majority of published empirical observations cannot be reproduced^4–9^, rendering nearly futile any effort to build on these observations to further our understanding of basic biological mechanisms or design effective therapeutic approaches. Further, the resources and time spent attempting to reproduce findings from low-quality or incorrectly acquired data are estimated to cost the global scientific community about 200 billion dollars per year^10^. The root cause lies in experimental designs that are not structured or powered adequately for conclusive statistical analyses. Since all biomedical researchers cannot be expected to have a deep knowledge of statistics or easy access to trained statisticians, tools are desperately needed to help them check the design of their experiments and apply adequate statistical power estimation. Not only could this improve our confidence in scientific outcomes, it could help make biological experiments more time-efficient and cost-effective. For example, if a researcher could estimate how many experiments should be performed and how many cell lines, animals or tissue samples should be collected to achieve sufficient statistical power to test their hypothesis, they may adjust their experimental design to fit their time or budgetary constraints without jeopardizing the quality of their findings. Another source of scientific errors comes from technologies such as scRNA-seq, whose advances are leading to a rapid increase in studies involving so-called “pseudo-replication”, which treats non-independent measures as if they were independent. For example, carrying out multiple measurements on a single sample instead of using separate, independent samples would represent non-independent replication. The risk of pseudo-replication^11^ (illustrated further below), can be remedied by the implementation of rigorous statistical methods that apply to all aspects of the data arising from such designs.

## Introduction

Experiments with a repeated measures design are actually common in biomedical research. These experiments are characterized by multiple measurements made on the same biological unit, as opposed to several independent units. For example, measuring the expression of a given gene or protein in multiple cells from the same diseased tissue sample using technologies such as single cell RNA sequencing (scRNA-Seq)^12, 13^ or flow cytometry^14^ is a repeated measures experiment. Other examples include longitudinal or time-lapse imaging, where phenotypic measurements such as the size, shape or live-dead status^15–25^ of individual cells are acquired multiple times within the same culture; behavioral measurements of live animals, such as a mouse modelling Alzheimer’s disease (AD) that is assessed for its learning capacity across multiple trials^26^; or electrophysiological studies that evaluate multiple cells derived from multiple brain tissue slices of the same AD mouse model^27^. In all these cases, the units being measured repeatedly (e.g., cells or animals) are not independent, since they are either the exact same cell or animal, or derived from the same tissue sample. This dependence is why such designs are referred to as “repeated measures”.

Commonly used statistical tests like t-tests, one-way and two-ANOVA treat all observations as independent of each other, and thus fail to account for the repeated measures nature of the data. Failure to account for the repeated measures nature of the data attributes overstates the statistical power of the measurements by treating all observations as independent and not taking into account their inherent dependency or correlation. However, these tests continue to be used^28, 29^, resulting in elevated rates of false discovery. A false positive occurs when the null hypothesis (“there is no difference between condition 1 and condition 2”) is rejected even though the evidence against it is insufficient. Conversely, a false negative happens when a statistical test fails to detect an association between two events, preventing the rejection of the null hypothesis. Both occur in underpowered studies—studies based on a dataset too small to reveal the expected effect, or where the effect size may be overestimated —and will lead to false conclusions within a given dataset^30, 31^.

A better way to analyze repeated measures datasets is to use Linear Mixed effects Models (LMM) and Generalized Linear Mixed effects Models (GLMM). Both are classes of statistical models designed to work with data generated under repeated measure designs. LMM is based on the assumption of normal probability distribution while a GLMM handles data that display a non-normal distribution. Both LMM and GLMM models deal with repeated measurements from the same source by adding random effect variables. Random effect variables account for the correlation of the observations arising from factors representing the different repeated sampling sources. This correlation of observations results in what is commonly referred to as batch-effects associated with the repeated sampling sources. For example, all observations in a given batch may have their values increased or decreased by a certain amount irrespective of their underlying biological condition. If it isn’t properly accounted for, it can lead to biased results and inflated p-values or coefficients. The main parameters of interest that researchers evaluate (such as the difference in mean measurements between control and treatment or disease) are modeled as fixed effects. Ignoring the random effects will result in too many false positives and bias the estimates of the parameters of interest. This problem is especially common in unbalanced experiments. For example, in a cell culture experiment, results can vary from batch to batch because of uncontrollable factors like different operators, reagents, or microscope settings. Since researchers care about the main effects (like treatment differences) rather than this random variation, these batch effects should be accounted for. LMM and GLMM models are well-suited for handling this.

Like all statistical models, LMM and GLMM come with their own sets of assumptions. Real data collected in experimental settings are often messy: they may include outliers; they may not perfectly meet probability distribution assumptions required by the specific mixed model chosen by users; and the true association between the response of interest and the chosen predictor may not be linear. Therefore, assessing the degree of violation of the assumptions of all models including LMM and GLMM is critical and often overlooked by scientists when evaluating their data.

In addition to ensuring that the assumptions with any given model are accurately vetted, it’s also crucial to establish that the sample size will result in an adequately powered set of data to maximize the reproducibility of the findings. For example, induced pluripotent stem cells (iPSCs) have transformed the ability of researchers to model human disease in a dish^32–34^, but the system poses numerous challenges. Genetic variability results in variation across cell lines^35–37^. Further, a typical neuronal differentiation can take months and use dozens of morphogens and reagents, and hence is subject to fluctuations due to time, batch and cell culturist. The variability that arises from the instruments used, morphogens or reagents applied, genetic variability, and other unknown factors makes it challenging to determine how much data is required to reliably interpret a set of results^38, 39^.

Statistical tools to perform power analyses in the context of repeated measures experiments have been lacking in situations where the error distributions are not normal, and for arbitrary experimental designs involving multiple factors. Simulation-based solutions are typically the only recourse in situations where one does not have access to simple formulae^40^ for sample-size estimations. The R package SIMR^41^, which is built for GLMM^41^, provides one such solution. Users can calculate power or sample size for a given regression model that includes single, multiple fixed or random effects in a general setting. For example, the SIMR package simulates values of responses with user-provided input values or pilot data to estimate the statistical power needed to capture given effects. Commonly used statistical software like SPSS^42^, STATA^43^ and SAS^44^ (that are behind paywalls) do not provide convenient implementation of simulation-based sample size estimations in the general repeated measures experimental scenario. Moreover, these packages remain cumbersome and difficult for non-specialists to use. To address this challenge and make the tools more accessible, we developed RMeDPower2 (**<u>R</u>**epeated **<u>Me</u>**asures **<u>D</u>**esign and **<u>P</u>**ower **2**\*). *= the number 2 is used here because it is the second version of RMedPower.

In this paper, we describe the development and features of RMeDPower2, which is based on R^45^. RMeDPower2 provides complete functionality to analyze data coming from repeated measures experiments such as those detailed above. RMeDPower2 tests the modeling assumptions needed to perform accurate analysis of any given dataset, identifies outlier observations as well as outlier units (i.e. batch, animal, tissue slice etc.), estimates statistical power to perform sample size calculations, estimates parameters of interest and also visualizes the association being tested. The functionality is currently limited to testing associations of one predictor (continuous or categorical, e.g., disease status or brain pathology) along with one covariate (continuous or categorical, e.g., time or sex status) and a potential interaction between the predictor and the covariate in the context of nested/hierarchical or crossed experimental designs (see Methods for an explanation of nested and crossed designs). It also allows for random or subject-level slopes when either the predictor or covariate are continuous. In sum, RMeDPower2 constitutes a comprehensive suite of statistical tools designed to rigorously and effectively model repeated measures data, and is accessible to all biologists with only a minimal experience of using R and a basic understanding of their experimental design and statistics. RMeDPower2 is not meant to replace the expertise of a trained statistician, but rather to empower biologists to independently carry out robust experimental design, model diagnostics, and parameter estimation to arrive at a well-informed, evidence-based understanding of their data before consulting one.

We illustrate the usefulness of RMeDPower2 by applying it to two datasets consisting of images of iPSC-derived motor neurons (iMNs) from patients with sporadic ALS (sALS) and unaffected controls. We report that sALS iMNs display sigincanlty fewer branches compared to controls. We also showcase RMeDPower2’s utility with other datasets (**Note S1**) such as the analysis of single-cell gene expression data, mouse behavioral measurements and electrophysiological studies, demonstrating its utility to analyze a wide range of data types.

## Results

### The RMeDPower2 workflow

RMeDPower2 helps scientists design experiments involving repeated measures, including not only the modeling stage and the planning for the collection of data, but also the estimation and visualization of the parameters of interest. The flowchart in **Figure 1** illustrates the use of RMeDPower2.

**Figure 1.**
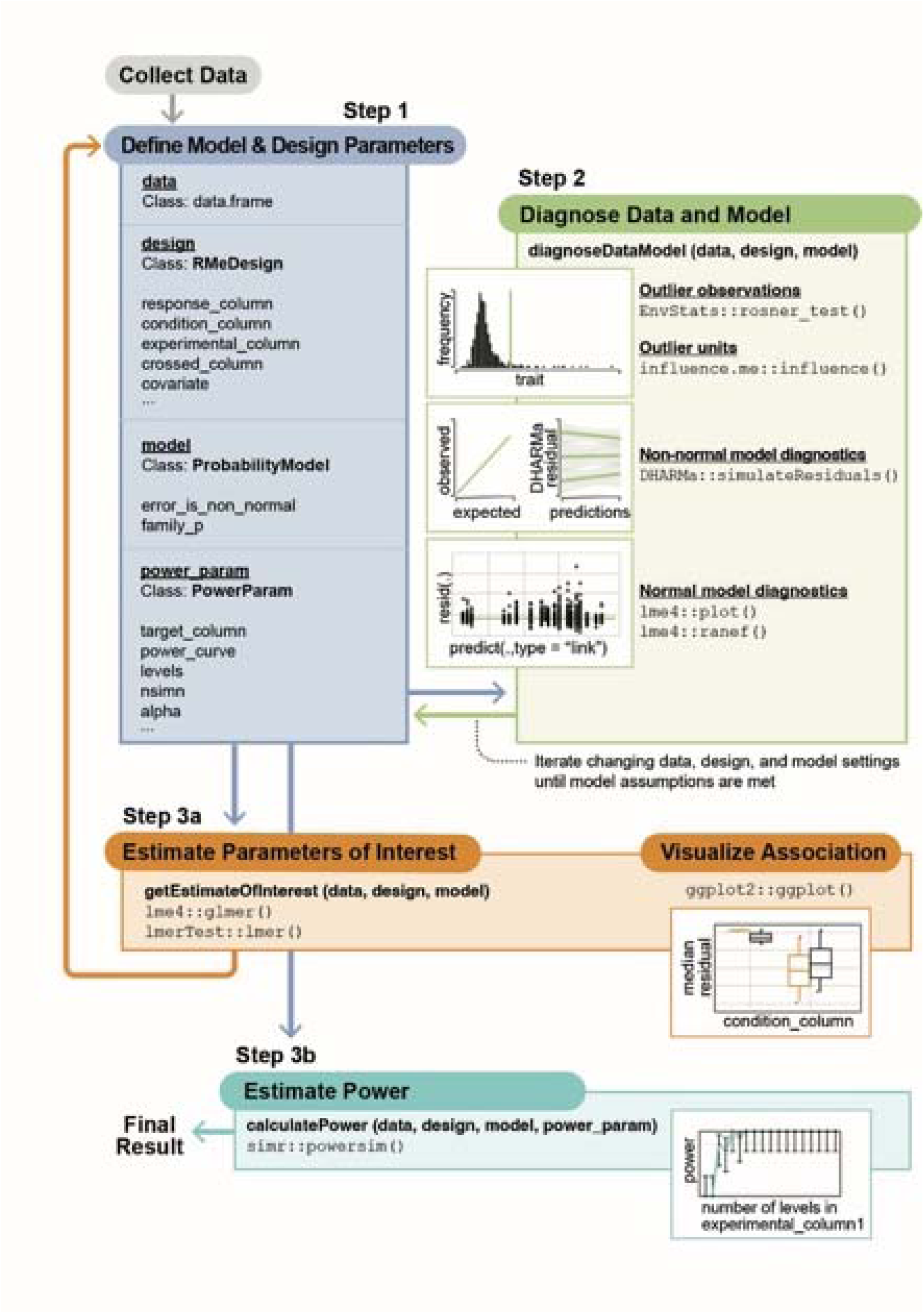
RMeDPower2 analysis flow chart. Flow chart illustrating the steps in involved in the planning/design for data to answer biological questions of interest (“Define Model & Design Parameters”), verification of modeling assumptions, identification of outliers observations and units (“Diagnose Data and Model”), planning of additional experiments to have adequate statistical power for the observed association (“Estimate Power”) and finally for the derivation of the estimates parameters of interest from experiments involving repeated measures designs (“Estimate Parameters of Interest”). The first step involves the definition of 3 S4 class objects capturing the design *(RMeDesign*), model for the data (*ProbabilityModel*) and the parameter choices for sample size estimations (*PowerParam*). These three objects along with the input data are used as input to the three primary functions – *diagnoseDataModel*, *calculatePower* and *getEstimatesOfInterest*. RMeDPower2 brings together the relevant functionality implemented in multiple R Packages. Inside the panels related to the steps in the flowchart, the R package names are placed before the “::” symbol, while the associated function names are placed after this symbol. The flowchart represents a typical scenario where a researcher collects pilot data under a repeated measures scenario to answer a question of interest. They then need to come up with an appropriate model for the data that they can use to generate their parameter of interest. This will involve making and testing assumptions regarding the probability distribution underlying the data generating distribution, identifying outliers that may otherwise result in violation of the modeling assumptions or strongly influence the estimate of the interest (*diagnoseDataModel*). Once this is done, they will need to ensure that they have a design for the data that is adequately powered to estimate the observed association with the pilot data (*calculatePower*). They may then need to generate new experimental data, again diagnose the data and verify the modeling assumptions (*diagnoseDataModel*) and finally generate and visualize the parameters of interest (*getEstimatesOfInterest*).

The input data is assumed to be organized into a table with column names or headers identifying the key variables in the experiment (e.g. disease status, drug treatment, batch replicate, plate or other variables capturing the different repeated measures or experimental design, etc.). These column names are then used to define the 3 primary S4 objects (a data type or organization of data in R) capturing the experimental design (*RMeDesign*), model assumptions (*ProbabilityModel*) and sample size estimations (*PowerParam*) (**Figure 1**: Step 1). The user needs to specify the assumed probability distribution of the response being modeled (e.g. Normal or Binomial or Poisson). For example, a normal probability distribution where data is continuous and follows a Gaussian distribution such as the size or shape of cells in a population. Alternatively, data that that has binomial probability distribution occurs when there are only two possible outcomes such as cell is determined to be alive or dead or a cell’s gene expression is activated or inactivated. A Poisson probability distribution is typically used to model counts such as the number of sequencing reads or cells mapping to a given gene or cell-type for a given biological RMeDPower2 is therefore capable of modeling both Gaussian and non-Gaussian data. In addition, this step allows the user to specify parameters required for sample size calculation or power analysis to determine the number of cells, lines, experimental batches, etc., which aids in experimental planning. For example, the user may want to know how many experimental batches would be needed to have at least 80% statistical power to estimate a given disease effect on cell-size at a Type I error rate of 0.05. The 3 S4 class objects^46^ are used as inputs in one framework that brings together the functionality of multiple existing R packages: lme4^47^ (implementation of LMM and GLMM), lmerTest^48^ (approximation of p-values for LMM), influence.ME^49^ (identification of outlier units), EnvStats^50^ (identification of outlier observations), DHARMa^51^ (testing of modeling assumptions for non-normal distributions), simr^41, 52^ (sample-size calculations) and tidyverse^53^ (data manipulation and visualization).

The function *diagnoseDataModel* tests the modeling assumptions and identifies potential outliers (**Figure 1**: Step 2). Next the *calculatePower* function performs sample-size estimations (**Figure 1**: Step 3a) while the *getEstimatesOfInterest* estimates the parameter of interest and visualizes the resulting association (**Figure 1**: Step 3a). With each step in this workflow, the user can go back and change some settings if the model assumptions seem violated, or re-test the assumptions after having removed outlier observations or outlier units, generate more data based on the estimates from *calculatePower* and repeat the workflow with the newly gathered data. There is additional information along with a convenient Rshiny app guiding a user on how to appropriately specify the relevant objects in RMeDPower2 [https://gladstone-institutes.github.io/RMeDPower2/articles/RMeDPower2_Class_Configuration_Guide.html].

### Simulation scheme for sample-size calculations

We now describe the simulation scheme using an example from a cellular assay where a user can evaluate the effect size comparing the effect of a drug (e.g., control vs. treated) that contain multiple lines representing each class from a pilot set of data. Consider a situation where a user wants to compare the response of 6 different cell lines to a given treatment. The pilot dataset consists of measurements of the 6 cell lines split into 2 plates, conducted over 3 experimental batches (**Figure 2**). To determine the impact of experimental design on statistical power, users can explore multiple alternative scenarios. They can increase the number of independent experiments to 6 instead of 3, for a total of 12 plates (**Figure 2A**), or they can increase the number of plates per experiment (**Figure 2B**). They might also examine the effect of expanding the number of cell lines within each experiment (**Figure 2C**). These 3 types of variable expansion can be accomplished by setting ‘*level=1*’ in the PowerParams object. **Tables S1-S3,** in addition to **Figure 2**, show the change in the number of cells required depending on the expansion type (experiment, plate or cell line), especially when the cell numbers distribution is asymmetric (e.g. the populations or cell counts differ across experiments). Asymmetry in cell numbers may occur because of different reasons, for example if a diseased cell line dies more rapidly than the control line, or if transduction of a biosensor occurs to differing extents between cell lines.

**Figure 2.**
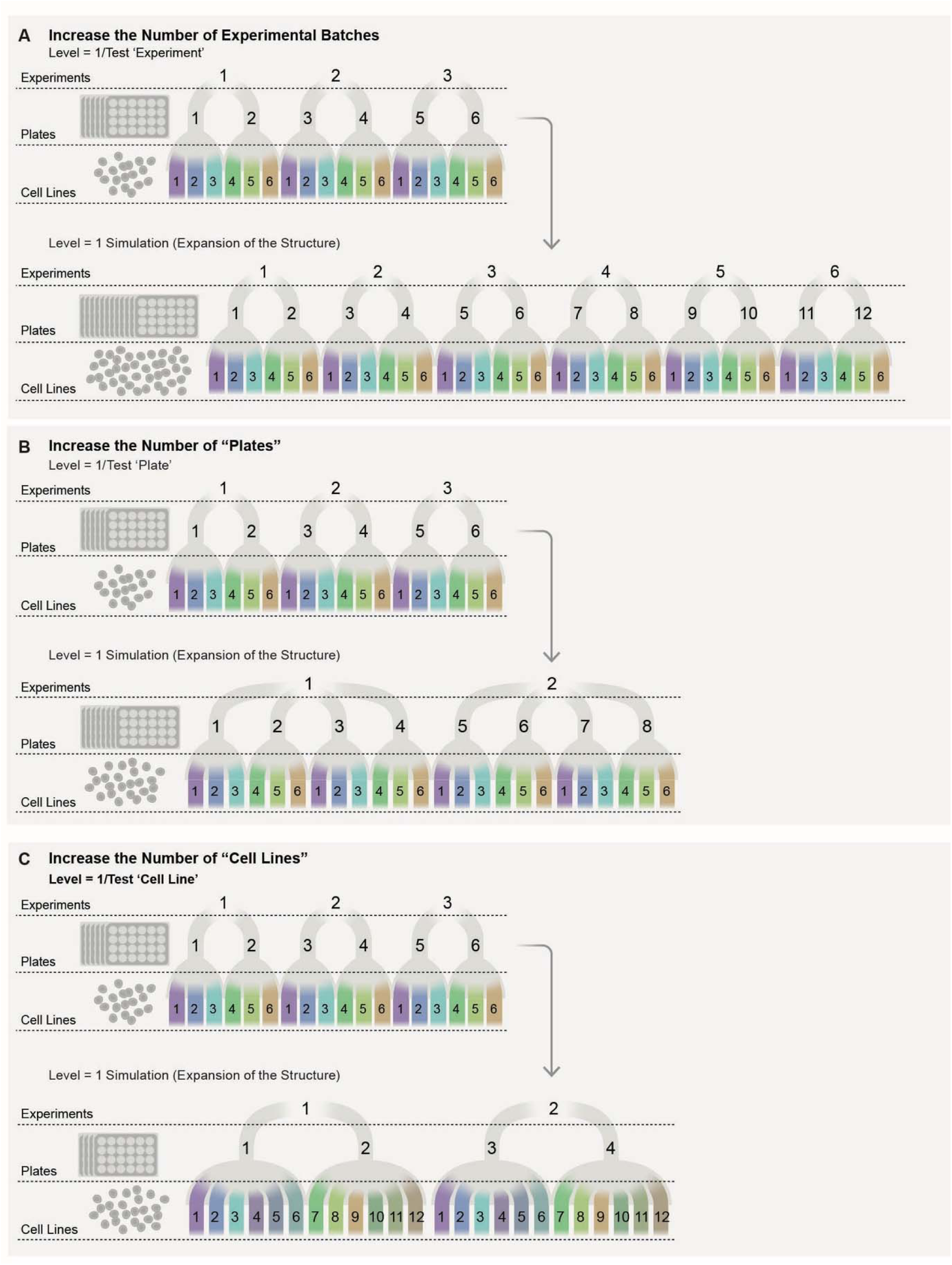
Example scenario of cellular assay involving multiple experimental batches, plates and cell-lines. The cell-lines are assumed to be drawn from either a diseased or a healthy population. The biological question centers around identifying mean differences in an individual cell characteristic (e.g., size) between the diseased and healthy populations. **A.** Illustration of the simulation of experimental variability to model increase in the number of experimental batches or ‘experiments’ for sample-size estimations. If *level*=1 (level is an attribute of the *PowerParam* class) is set, RMeDPower2 simulates the effect of adding in additional experiments by inheriting the experimental design structure from the existing data. This describes the situation where the pilot study involves data from 3 experiments, with 2 plates used per experiment, 3 cell lines within each plate, resulting in a total of 6 cell lines. In this case, we simulate the effect of doubling the number of experimental replicates, which results in 6 experiments containing 12 plates. **B.** Example of simulation of plate variability to model an increase in the number of ‘plates’ for sample-size estimations. If *level*=1 is set, RMeDPower2 simulates new plates by inheriting the experimental design structure from the existing data. Here we start with 2 experiments containing 4 plates and we simulate the effect of doubling the number of plates per experiment to 8 plates. **C.** Example of simulation of cell-line variability to model increase in the number of cell lines for sample-size estimations. If *level*=1 is set, RMeDPower2 simulates new cell lines by inheriting the experimental design structure from the existing data. In this case, we start with 2 experiments containing 4 plates with 6 cell lines, and we simulate an increase in the number of cell lines such that we have 12 cell lines.

Alternatively, users may want to examine the power of increasing the total number of cells they measure per experiment. For example, by doubling the number of cells for each cell line by increasing the number of wells or plates, they can assess the effect in twice the number of cells per cell line without changing the number of experiments and cell lines (**Figure 3A**). Alternatively, one might want to assess the effect of increasing the total number of cells by increasing the number of cells per cell line per well rather than by adding more plates (**Figure 3B**). Expansion results for plates and cell lines may differ if cells are asymmetrically distributed in different settings. **Tables S4** and **S5** show the different results users can get in an asymmetric distributed cell scenario. This type of variable expansion can be accomplished by setting ‘*level=0’*. An approach may be chosen based on reagent costs or the time required to complete each experiment.

**Figure 3.**
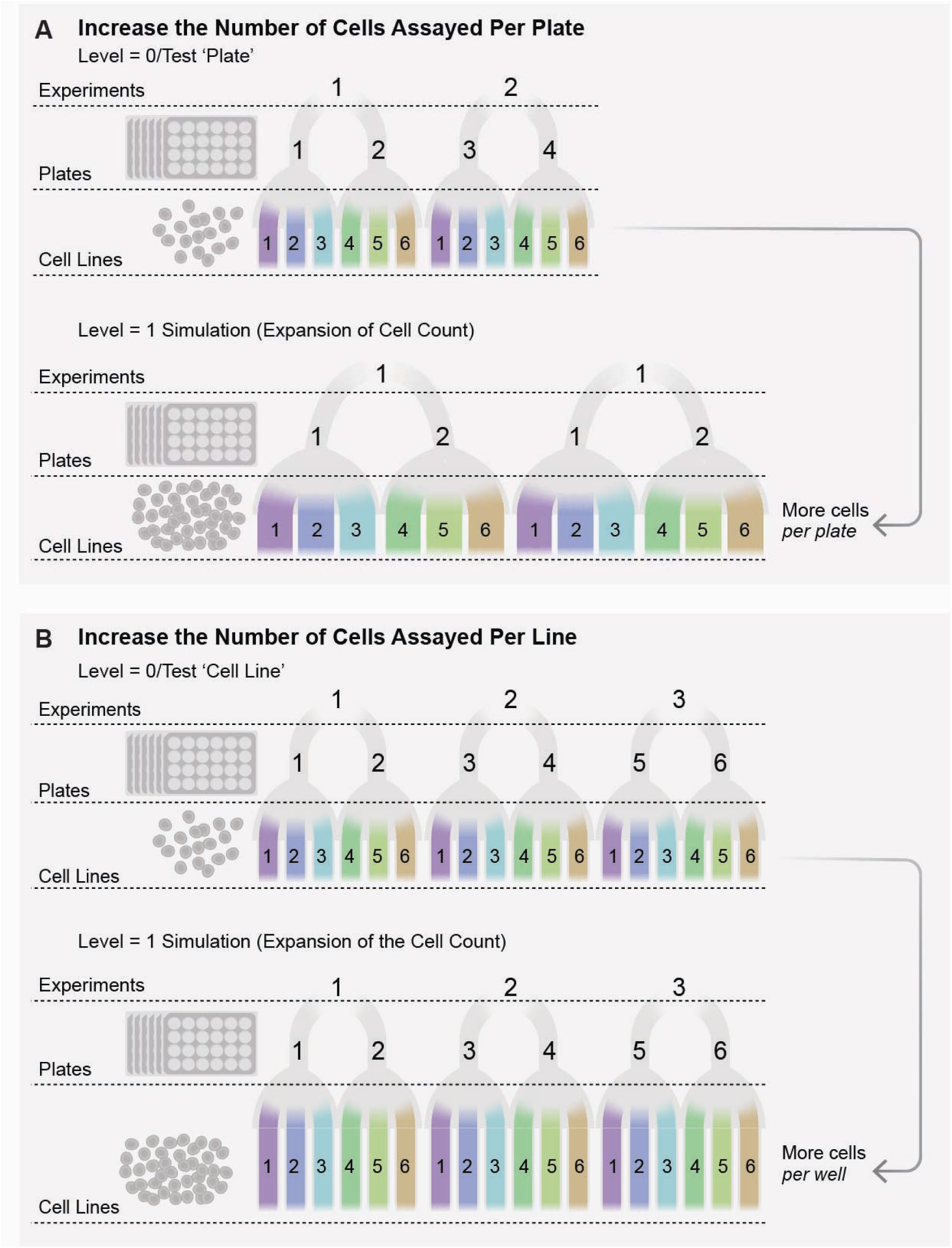
Example of cellular assay involving multiple experimental batches, plates and cell-lines. The cell lines are assumed to be drawn from either a diseased or a healthy population. The biological question centers around identifying mean differences in an individual cell characteristic (e.g., size) between the diseased and healthy populations. **A.** Example of simulation to model the power estimates as a function of increasing the number of cells assayed per plate. If *level*=0 (*level* is an attribute of the *PowerParam* class) is set, RMeDPower2 multiplies the number of cells per plate by *M/N*, where *N* is the maximum number of cells per plate and M is a value assigned to the parameter *max_size* (another attribute of the *PowerParam* class). **B.** Example of simulation to model the power estimates as a function of increasing the number of cells assayed per cell-line per well. If *level*=0 is set, RMeDPower2 multiplies the number of cells per cell line by *M/N*, where *N* is the maximum number of cells per cell line and *M* is a value assigned to the parameter *max_size*.

These simulation analyses can inform a user on how to best carry out a set of studies in the most cost- and time-effective way. One might initially assume that increasing power simply requires conducting more experiments; however, this approach can be both time-consuming and costly. Simulated scenarios, by contrast, may reveal that increasing the number of plates per experiment while performing fewer experiments (scenario 2B) can achieve the same power as scenario 2A, but at a lower cost.

### Applying RMeDPower2 to the study of morphological differences in iPSC-derived neurons from ALS patient lines and controls

We present two case studies illustrating the potential of RMeDPower2 for rigorous statistical estimation of the morphological differences in iPSC-derived neurons. The first case study is based on simulated neuronal perimeter data while the second one is based on real neuronal sholl feature (that quantifies the complexity of dendritic arbors) data. The two case studies illustrate differing complexities of the data and underlying statistical models. The first case study models change in simulated neuronal perimeter as a function of disease status (ALS vs control) based on a normal probability distribution assumption while the second one models changes in the sholl feature levels (modeled using a negative binomial probability distribution) as a function of disease status, time elapsed and the interaction between these two variables.

#### Case study 1

Previously, cortical and motor neurons were found to be smaller in the postmortem brain of ALS patients compared to controls^54^. However, animal ALS models display larger growth cones and neurites than control animals^55^. We wanted to examine if we could detect differences in the size and shape of motor neurons differentiated from induced pluripotent cells (iMNs) from control and sporadic ALS patients. To this end, we measured the soma perimeter from images of iMNs that were transduced with Synapsin:EGFP, a fluorescent marker expressed in neurons (**Figure 4A**), as a pilot set of data we fed to RMeDPower2.

**Figure 4.**
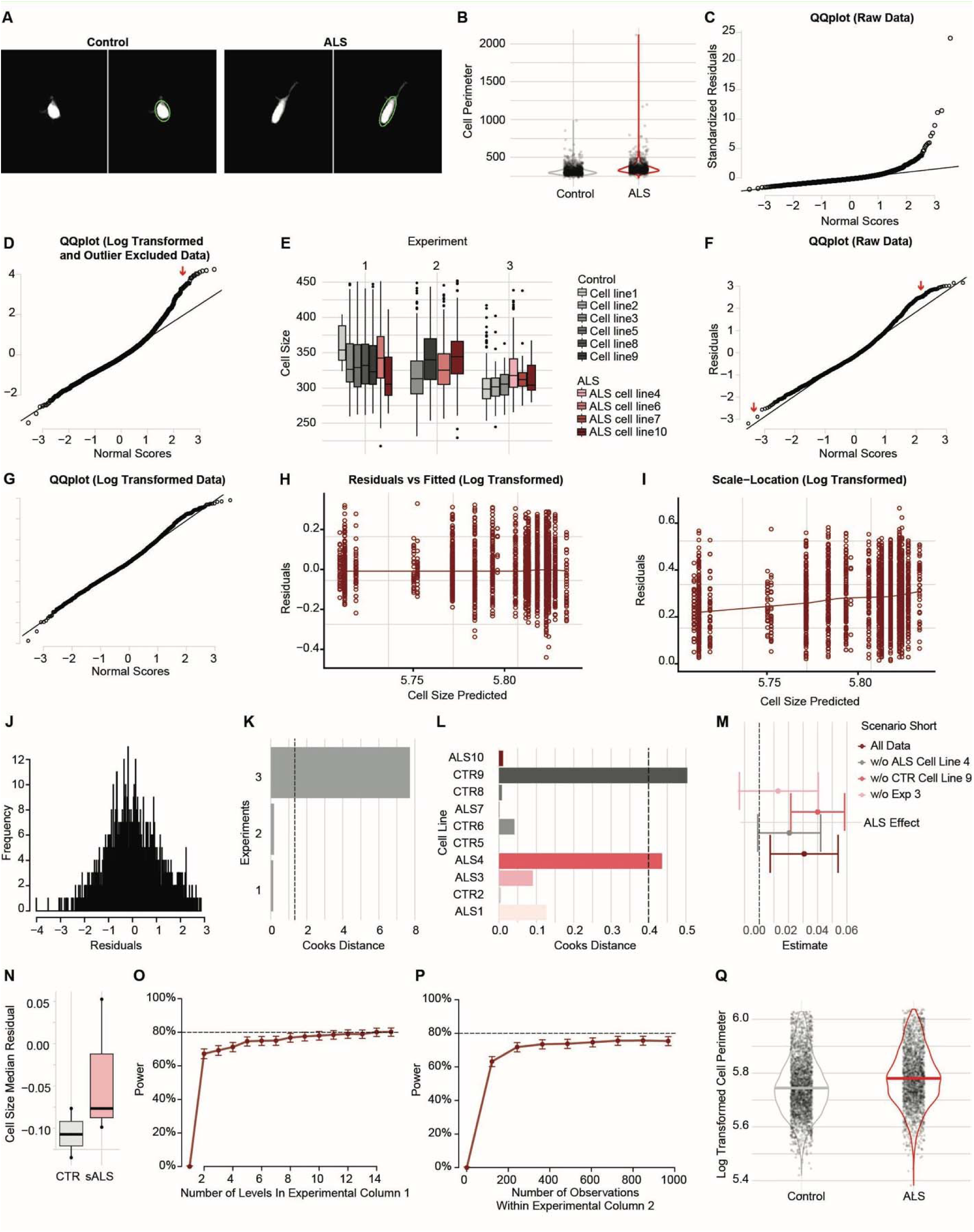
Workflow summarizing the data transformations and analysis to model neuronal soma perimeter of ALS and control /iMNs. **A.** Example images of control and sALS patient iMNs expressing a GFP. Images were acquired by RM and subjected to soma perimeter extraction using the eclipse function in Python. **B.** Violin plots depicting the distributions of the 1271 soma perimeter observations across the sALS lines and the 1317 observations across the control lines. Note, the data shown in this panel and in all the remaining panels and figures are based on simulated data with variance characteristics that match the real data. **C.** Q-Q plot of the distribution of residuals from the LMM fit with soma perimeter in its natural scale demonstrating the deviation of this distribution from normality (indicated by red arrow). The x-axis (Normal Scores) represents the quantiles from a standard normal distribution while the y-axis the quantiles of the observed residuals that were centered by their mean and scaled by their standard deviation (Standardized Residuals). The experimental batch and cell lines were modeled as random effects while the ALS status was modeled as a fixed effect in this LMM fit. **D.** Q-Q plot of the distribution of residuals from the LMM fit (with the same random and fixed effects) with soma perimeter in its logarithmic scale. The interpretation of the x and y-axis labels are the same as in **C**. Deviations of the distribution from normality are indicated by the red arrow. **E.** The top 5% of observations in **B.** are filtered out and the resulting distributions across the 3 experimental batches and 10 patient cell lines are plotted. The 6 control lines are colored in different shades of grey while the 4 ALS lines are colored in different shades of red. **F.** Q-Q plot of the distribution of residuals from the LMM fit with soma perimeter in its natural scale using the filtered data demonstrating the deviation of this distribution from normality. The interpretation of the x and y-axis labels are the same as in **C**. Red arrows indicate deviations from normality. **G.** Q-Q plot of the distribution of residuals from the LMM fit (with the same random and fixed effects) with the filtered data soma perimeter in logarithmic scale demonstrating a better fit to the normality assumption than in **C**, **D** and **F**. The interpretation of the x and y-axis labels are the same as in **C**. **H.** The horizontal best fit solid line to residuals versus the fitted values from the LMM fit in **G** does not suggest violation of linearity. **I.** The almost horizontal best fit solid line to square-root of the absolute value of the residuals versus the fitted values from the LMM fit in **G** suggests only a small violation of homoscedasticity assumption. **J.** Distribution of the residuals of the LMM fit in **G** did not result in any identified outlier observations using the Rosner’s test. **K.** Cook’s distance estimates for each of the three experimental batches using the LMM fit in **G** suggested that experimental batch 3 has a strong influence on the parameter estimates based on a threshold of 4/3 (the vertical dashed line). **L.** Cook’s distance estimates for each of the 10 cell-lines using the LMM fit in **G** suggested that cell-lines 4 and 9 have a strong influence on the final parameter estimates based on a threshold of 4/10 (the vertical dashed line). The 6 control lines are colored in different shades of grey while the 4 ALS lines are colored in different shades of red. **M.** Estimates (including corresponding 95% confidence intervals) of change in mean soma perimeter between the ALS lines and control lines using all the data and also after removing all observations from the experiment 3 and cell lines 4 and 9, individually. **N.** Distribution of the median of the residuals across all observations within each of the 3 experimental batches by ALS status of the observations. The residuals are computed using a LMM fit to the log-transformed soma perimeter observation and modeling the experimental batch and cell line as random effect and excluding the ALS status of these observations. The estimate of difference in log-transformed neuronal soma perimeters between ALS and control cell-line is 0.031 and p=0.03. **O.** Sample-size estimates as a function of the number of experimental batches a. Estimates of sample size as a function of the number of experimental batches, based on the data and model fit shown in Figure **4G**. The power curve shows that at least 8 experimental batches or more are needed to achieve a statistical power close to 80%. **P.** Sample-size estimates as a function of the number of cells per cell-line using the data and the model fit in Figure **4G**. The power curve indicates that 250 cells or more per cell-line to achieve a statistical power close to 80%. **Q.** Distribution of the median of the residuals across all observations within each of the 8 experimental batches by ALS status of the observations. The residuals are computed using a LMM fit to the log-transformed perimeter observation and modeling the experimental batch and cell line as random effect and excluding the ALS status of these observations. The estimate of difference in log-transformed neuronal soma perimeters between ALS and control cell-line is 0.027 and p=0.006.

Note that for the purpose of illustrating the functionality of the RMeDPower2 package, the ***data used here and shown in* *Figure 4* *are simulated*** from real cell soma perimeter measurements obtained from live cell imaging of control and ALS iMNs. The difference in cell soma perimeters we actually measured between the ALS and control lines corresponded to an effect size of around 1, while for the simulated data used here, the effect size was made closer to 2 (**Figure S1**). ***However, the number of experimental batches and the numbers of ALS and control cell lines used, as well as the variances of measurements due to these two sources are exactly the same in the simulated data as in the real data***.

### Define Design and Model Parameters (Figure 1: Step 1)

Our pilot data consist of neuronal soma perimeter measurements from ten different lines of iMNs (some derived from ALS patients and other from control lines) obtained across three independent experiments (**Table S6)**. Hereafter in this section, selected attributes for the 3 S4 objects in **Figure 1: Step 1** are highlighted as italicized terms in parenthesis. We are interested in testing the effect of the ALS status of each cell (*condition_column*) on its perimeter (*response_column*). We have repeated measures across the observations in each experiment batch and in each cell-line (*experimental_columns*). The design is similar to the one illustrated in Figure 2A, where only one plate is used per experiment. Further, cells from the same cell lines are measured across multiple experimental batches. Therefore, the cell line status (*crossed_columns*) of each observation is considered *crossed* with the experimental batch where the measurement is made. We assume that cell soma perimeter measurements across observations from the same cell line within the same experimental batch are normally distributed (*family_p*). We will explore below the effect of increasing the number of experimental batches and/or the number of cells per cell line (*target_column*) on the statistical power to detect our observed association.

### Diagnosis of Data and Model for the Cell Perimeter Observations (Figure 1: Step 2)

The cell soma perimeters of control and ALS cell lines did not appear obviously different when represented as a violin plot (**Figure 4B**). However, this finding is not necessarily accurate, since these plots consider all datapoints as independent, ignoring the “repeated measures” nature of our experimental design. This is a frequent yet often overlooked mistake that biologists make when handling their data.

We used an LMM to estimate the association between cell soma perimeter and ALS disease status. The validity of the inferences derived from this model fit depends on whether or not certain assumptions are met (specifically these assumptions center around the normality of residuals and that of the estimated random effects, no observable trend in each of the residuals versus fitted and the scale-location plots). To verify the normality of residuals assumption, we generated a Quantile-Quantile (QQ) plot of the residuals of the LMM model fit using the natural scale cell-perimeters. Residuals represent the difference between observed values and those predicted by a model. They indicate how far each data point deviates from the model’s expected value. When subsets of extreme values exist, these points can deviate substantially from the expected normal distribution. Alternatively, the data may need to be transformed, as discussed below.

The QQ plot reveals that there is a clear deviation from the diagonal at the higher end of the distribution (**Figure 4C**, red arrow), suggesting that the data are not normally distributed. Extreme outliers or values that span several orders of magnitude can distort normality. To address this issue, a user has two options: remove the outliers or apply a log transformation. Excluding outliers can eliminate extreme residuals, while a log transformation reduces skewness and helps the distribution approximate normality. Using a Rosner’test^56^, which is a statistical method for detecting multiple outliers in a dataset, we removed extreme values. In addition, we log- transformed the raw data and visualized using a QQ plot. This resulted in a visible skew from the expected normal distribution (**Figure 4D**, red arrow). Alternate parameterizations based on assigning the *family_p* parameter for the data generating distribution of cell size to the Gamma probability distribution did not improve this situation (data not shown).

Further examination of the distribution of the observed data in **Figure 4B** revealed a small fraction of extreme measurements at the higher end. We therefore filtered out the top 5% of the observations in part to ensure more robust estimates, like the trimmed mean parameter for location estimate of a parameter^57^, and in part to meet the normal distribution assumption required by LMM. Plotting the resulting distribution of the cell perimeters across the three experimental batches and 10 cell lines (**Figure 4E)** revealed that measurements from experiment 3 were lower than those from experiments 1 and 2 across all cell lines, evidence of a batch effect. QQ plots of the filtered data (after the top 5% of observations are removed) showed that deviation from normality was still evident for the QQ plot using the natural scale (**Figure 4F**, red arrows), but much reduced in the plot using the log-transformed perimeter values (**Figure 4G**). Therefore, we modeled the filtered, log-transformed perimeter values.

The plot of residuals versus the fitted values is used to evaluate the adequacy of the linear model fit to the data^47^. If the best fit line for this plot is horizontal or close to being so then the linear model is assumed to be reasonable. In addition, the plot of the square-root of the absolute values of the residuals versus the fitted values is used to assess the homoscedasticity or equal-variance assumption for the model used^47^. Similarly, if the best fit line for this plot is horizontal then the equal variance assumption is reasonable for the fitted model. To understand the best fit line, there are examples where these quality control plots would suggest violation of the assumptions or where these assumptions are met for linear model^58^. The best fit lines to the residuals versus the fitted values was reasonably horizontal (**Figure 4H)** and the square-root of the absolute values of the residuals versus the fitted values (**Figure 4I)** suggest both the LMM model assumptions of linearity and homoscedasticity are reasonable. In other words, the check for model fits in **Figures 4H** and **4I** show that the model looks like a good fit.

The inferences from the LMM could be strongly influenced by some individual observations or by all observations within a batch or a cell line. Therefore, we reapplied the Rosner’test^56^ to the filtered data, but we did not identify any individual observations as outliers (**Figure 4J**). While Rosner’s test is effective at detecting outliers within multiple datasets, it does not account for situations where an entire experiment or cell line behaves as an outlier. To capture these broader deviations, a different method is required. To identify experimental batch or cell line outliers, we estimated how influential all observations within each experimental batch and cell line were on the estimate of our parameter of interest, i.e., the mean difference in the perimeter of cells derived from ALS lines versus control lines, using Cook’s distance^59^. Cook’s Distance is a way to measure how much influence each entire set of related data has on a regression model. We can visualize the output of the Cook’s distance using a bar plot. It was clear that the observations from experimental batch 3 displayed a high Cook’s distance that crossed the suggested threshold^49^ of 4/n (where n is the number of experimental batches being tested for their influence, i.e. n = 3) for identifying influential units (**Figure 4K**). Similarly, the Cook’s distance estimates for cell lines 4# and 9# passed the suggested threshold of 4/n (with n = 10 for 10 cell lines) and qualified as influential cell lines (**Figure 4L**).

This warranted further investigation on whether or not there was something unusual with the observations made in experimental batch 3 (example: the images from the microscope displayed low signal to noise and may have skewed the measurements) and also for cell line 4# and 9# (example: differentiation runs, genetic background was different from the other lines) that would suggest observations linked to this experiment and cell lines may need to be removed from the analysis. Our investigation did not indicate anything **untoward** and therefore we ran a sensitivity analysis where we dropped all observations linked to either experiment 3, cell line 4# or cell line 9# to examine the resulting estimate (**Figure 4M**). After refitting the model, we visualized the resulting change in ALS status associated mean soma perimeter value (**Figure 4M**). Despite there being slight changes in the estimated changes in the mean, the conclusions we would draw of an increase in the soma cell perimeter in ALS lines remains irrespective of whether or not we drop observations from the experimental batch or the two cell lines. This analysis confirms that the interpretation of results is not meaningfully driven by any single experimental batch or cell line.

### Derivation of the Estimate of Interest – the Mean Difference in Cell Perimeters (Figure 1: Step 3a)

We next derived our estimate of interest based on the model of the log-transformed data using the *getEstimatesOfInterest* function. The difference in the mean log-transformed cell size between ALS lines and control lines was 0.031, and the significance (p-value) of this difference 0.03 (as indicated above, this conclusion is based on simulated data.

The function also generates a visualization of this association using boxplots (**Figure 4N)**, where the individual points represent the median of the observations’ residuals in each of the experimental batches across either all control lines or all ALS lines. The residuals calculated in **Figure 4N** are based on a model fit that excludes the ALS status of the observations. The goal here is to provide a clear visualization of the estimated relationship between perimeter and disease class, free from confounding effects related to specific experiments or cell lines. We see a small increase in the median residual of cell soma sizes for the ALS iMNs, in line with the estimates we obtained from the model fit. Note that the residuals from the model fit, summarized by taking the median of all residuals within a disease class at the experiment level, are visualized here instead of the actual perimeter values. The reason is that observed changes in the actual perimeter values could potentially be confounded due to variation in these values between experimental batches and cell-lines, whereas the residual values will not be confounded by these variables. The summary at the (more-or-less independent) experiment level rather than using individual residual values per disease class should better illustrate the association, particularly in situations where there is a relatively large number of observations per experiment.

### Calculate Power – influence of increasing the number of batches or the number of cells (Figure 1: Step 3b)

For the same statistical power, users might intuitively expect that a larger number of experimental batches would be needed to estimate differences across groups that display a small effect size. Effect size is defined as the difference of mean cell sizes between the disease conditions scaled by the standard deviation of this difference. To examine if this is true, we ran simulations using data shaped by real observations. For each scenario, we looked at different effect sizes and calculated how many experimental batches would be needed to reach 80% power for detecting those effects. Note that this represents the only analysis in this manuscript where the results are based on the observed cell size data. For purposes of illustrating the functionality of RMeDPower2, simulated data with an effect size of 2 was used in all other analyses. Using RMedPower2 for this calculation, we found that we would need close to 15 experimental batches to reach 80% power when the effect size was around 1. The number of required batches decreased as the effect size increased, as expected, eventually reaching a plateau of value 3 with high effect sizes (**Figure S1**). As a reference for this simulation result, we used the sample-size calculations based on the two-sample t-test implemented in the pwr.t.test function from the R library pwr^40^. Specifically, we asked the number of samples per group that would be required to reach 80% power at 0.05 Type I error level for a two-group comparison across the range of the observed effect sizes. Both analyses showed that fewer experiments would be required as effect size increased, but the t-test predicted a smaller number of experiments across all effect sizes than did RMedPower2. This difference reflects the variability between ALS or control lines that was modeled in the RMedPower2’s, but not the t-test’s power estimates.

Given that the observed difference of 0.03 units in the simulated cell soma size between ALS and control lines had a 72.1% statistical power with a 95% confidence interval (69.2, 74.9) at type I error rate of 0.05, our next step is to see how we could increase the statistical power of our experiments. Using the pilot data and the model parameter settings, we arrived at the end of the **Diagnose Data and Model** section (**Figure 1: Step 3**). Using the *calculatePower* function, we generated a Power Curve (PC) (**Figure 4O**) to estimate the statistical power to detect the observed change in mean cell size between the ALS and the control iMNs as a function of increasing number of experimental batches. A PC is a graph that shows how likely a statistical test is to detect a true effect (if it exists) under different conditions. In this way, one can determine the probability that a test correctly rejects the null hypothesis (i.e., it finds a real effect when there is one). A PC plots the power probability (on the y-axis) against variables that change in the study, such as the number of experimental batches or number of observations as shown in **Figure 4 O, P** (on the x-axis). Using this curve, we can predict that with at least 8 independent experimental batches, we will achieve close to 80% power (or to be exact 78.20% power with a 95% confidence interval (75.51, 80.72)) to detect robust associations between the cell soma perimeter and ALS status of the cells (**Figure 4O**).

Another way to increase power might be to increase the number of cells assayed for each line (as illustrated in **Figure 3B).** In this case, the PC showed that the power to detect the observed association between ALS status and cell size did not improve substantially beyond around 250 cells per cell line, at which point power estimate plateaued near 70% (**Figure 4P)**. In other words, we will need at least around 250 cells per cell line to maximize the power to detect the observed association of cell soma perimeter. In the pilot data, the median number of cells per cell line is 246. We have elaborated on simulations underlying power estimation to Section 3 in Note S1.

### Analysis and Visualizations of Final Dataset (Figure 1: Step 3a)

Power calculations based on our pilot dataset indicated that 8 experimental replicates would provide a set of associations between ALS status and the cell perimeter that achieves close to 80% statistical power with a type I error rate of 0.05 (**Figure 4O)**. We therefore replicated the experiment 5 more times to achieve 8 experimental replicates, and also increased the median number of cells per cell lines using observations across all batches to 407. Given the results from **Figure 4 O P**, we could have picked a larger (than 8) number of batches, but there were only marginal improvements in the estimated statistical power beyond eight experimental batches. Using the log-transformed values of the observations across all 8 batches (**Table S7**) we derived an estimate of 0.028 as the mean difference in cell soma perimeter with an associated significance of p = 0.004 (**Figure 4 Q**), thus validating the predictions based on the pilot data. The estimated power to detect the observed association with data from the 8 experimental batches is 93.70% with 95% confidence interval of (92.01, 95.13).

The power simulations depend on the variance components for the different technical factors estimated from the input dataset. However, in some cases there may not be enough data to estimate the variance components, or the variance of the response due to differences in each of the experimental factors. For example, the pilot data might have only a single category for ‘experimental batch’, ‘plate’ or ‘line’ (**Table S8**). When this occurs, the user needs to provide Intra-Class correlation Coefficients (ICC) values that reflects the proportion of variance of the responses due to each of the different experimental factors. These values can be estimated from another dataset for which the variance components are assumed to be similar to those inherent for the new response being considered in the input dataset. **Figure S2** shows the PC as a function of the number of experimental batches when ICC values for experimental batch, cell line and plate are set to be equal to 0.8, 0.05 and 0.05 respectively (based on estimates from other data sets not shown here). In this example, we observe that we need at least 6 experimental batches to have at least 80% power to detect the observed association.

Here we have illustrated the functionality of RMeDPower2 using neuronal cell morphology data from differentiated ALS patient and control subject cell lines. Note again that these analyses were based on a response (*cell_size2*) reflecting a larger effect size than present in the observed data. This choice was made to illustrate RMeDPower2’s functionality more clearly with the data we had in hand. The focus here is on the package itself, not on the actual cell morphology differences between the disease and control lines. That said, the estimate of the actual difference in the observed mean log-transformed neuronal cell soma perimeter between the ALS and patient lines, using observations from all 8 experimental batches and following the same process illustrated in **Figure 1**, is 0.015 and the associated significance is 0.07. This is suggestive of a real increase in perimeter of neurons derived from ALS patient lines compared to controls, though our power simulations (**Figure S1**) demonstrate that data across at least 15 experimental batches should be acquired before making this claim. These findings are consistent with previous reports examining neuronal size in mouse models of ALS that reported larger somas than in control mice^60, 61^.

#### Case study 2

In this next case study, we describe a significant finding of ALS-associated changes in the sholl feature in iMNs in a real data set. Several studies have reported that motor neurons derived from sALS iPSC lines have consistently displayed progressive neurite shortening and regression, reduced neurite outgrowth and branching^62–64^. We hypothesized that neurite branching may be altered in sALS iMNs compered to controls. Therefore, we analyzed iMNs to determine if there were dynamic and static changes associated with sALS iMNs. Here we will illustrate the steps we followed using RMedPower to investigate this.

iMNs were transduced with the RGEDI biosensor (RGEDI-P2a-EGFP)^65^, which combines an EGFP (green) marker of cell morphology with a Ca^++^-sensitive red fluorescent marker that only activates when cells are dying^65^. iMNs were transduced with RGEDI on differentiation day DIV15–17 and subjected ∼ 1 week later (∼DIV27) to daily time-lapse imaging using RM^15, 17–22, 66–68^ in both the red and green channels^65, 69^ for 9 days^70^. iMNs displayed typical neuronal morphology with dendrites and branching processes. Images were cropped as previously described to contain only the soma and emanating processes (**Figure 5A top**). We subjected the crops to a sholl feature extraction algorithm^71^ that captures how many structures extend from the object in the middle of each crop and can be interpreted as a way to characterize the number of processes emanating from each neuron (**Figure 5A bottom**). We measured this on an individual neuron basis in 5 control and 5 ALS cell lines at two time-points (time-points 1 and 6) across 8 experiments. The distribution of sholl values grouped by ALS status and over the two time- points is shown in **Figure 5B**. No obvious differences in this feature value by either ALS status or time were apparent. However, we see an obvious effect over time when the distribution of the sholl values is separated by experiment and cell line (**Figure 5C**). We also see an experiment effect where all observations in certain experiments (e.g. Experiment 1) have relatively large sholl values while other experiments (e.g. Experiment 6) have relatively small sholl values, irrespective of ALS status and time-point of the observations. We define a RMeDesign object with the sholl values as the *response_column*, ALS status as the *condition_column*, time-point as the *covariate* and include an interaction between these two variables. Like how we modeled the cell perimeter data, the *experimental_columns* attribute is chosen to be experimental batch and cell line while cell line is considered to be part of *crossed_columns*. The Poisson probability distribution is an obvious choice to model the count nature of the sholl feature. However, the Q-Q plot of the residuals generated by the *diagnoseDataModel* function does not suggest a good fit (**Figure 5D)**. This leads us to the more flexible Negative Binomial distribution as an alternate probability distribution. The Q-Q plot of the resulting residuals suggests a better fit (**Figure 5E**), while the fit looks even better after individual outlier observations are removed (**Figure 5F**). Hereafter for the rest of the analysis, we use the data with the individual outliers removed. The distribution of the residuals within each combination of ALS status and time-point of observation mostly exhibits the desired uniform distribution (**Figure 5G**). The Q-Q plots for the random effects associated with each experiment and cell-line approximately follows the expected normal distribution (**Figures 5 H, I**). One experiment and two cell-lines are identified as influential in the parameter estimates based on the Cook’s distance metric (**Figures 5 J, K**). We plot the parameter estimates after removing the groups of observations linked to the identified influential experiment and two cell-lines (**Figure 5L**). Our model leads to three main findings. In control cell lines, mean Sholl values decline by 0.15 units between time points 1 and 6 (p = 2.2e-16), while sALS lines decline an additional 0.03 units over the same period (p = 0.002). sALS lines decline 0.05 units less than controls over this interval (p = 0.015). When we refit the model excluding three groups of identified outliers, the parameter estimates change slightly but the overall conclusions remain consistent. However, the conclusions one draws remains the same irrespective of which group of observations are not included in the final model fit. Therefore, the final estimates (using the function *getEstimatesOfInterest*) that we visualize (in **Figure 5M**) are based all groups of observations. We are well powered (100.0% statistical power and a 95% confidence interval of (96.38, 100.0) at a type I error rate of 0.05 estimated using the *calculate_power* function) to make conclusions based on the associations of sholl feature values at the observed effect sizes. This type of change over time we call the <u>m</u>orphology <u>c</u>hange <u>o</u>ver time or MCOT^72^ to reflect a time-by-disease-status interaction term.

**Figure 5.**
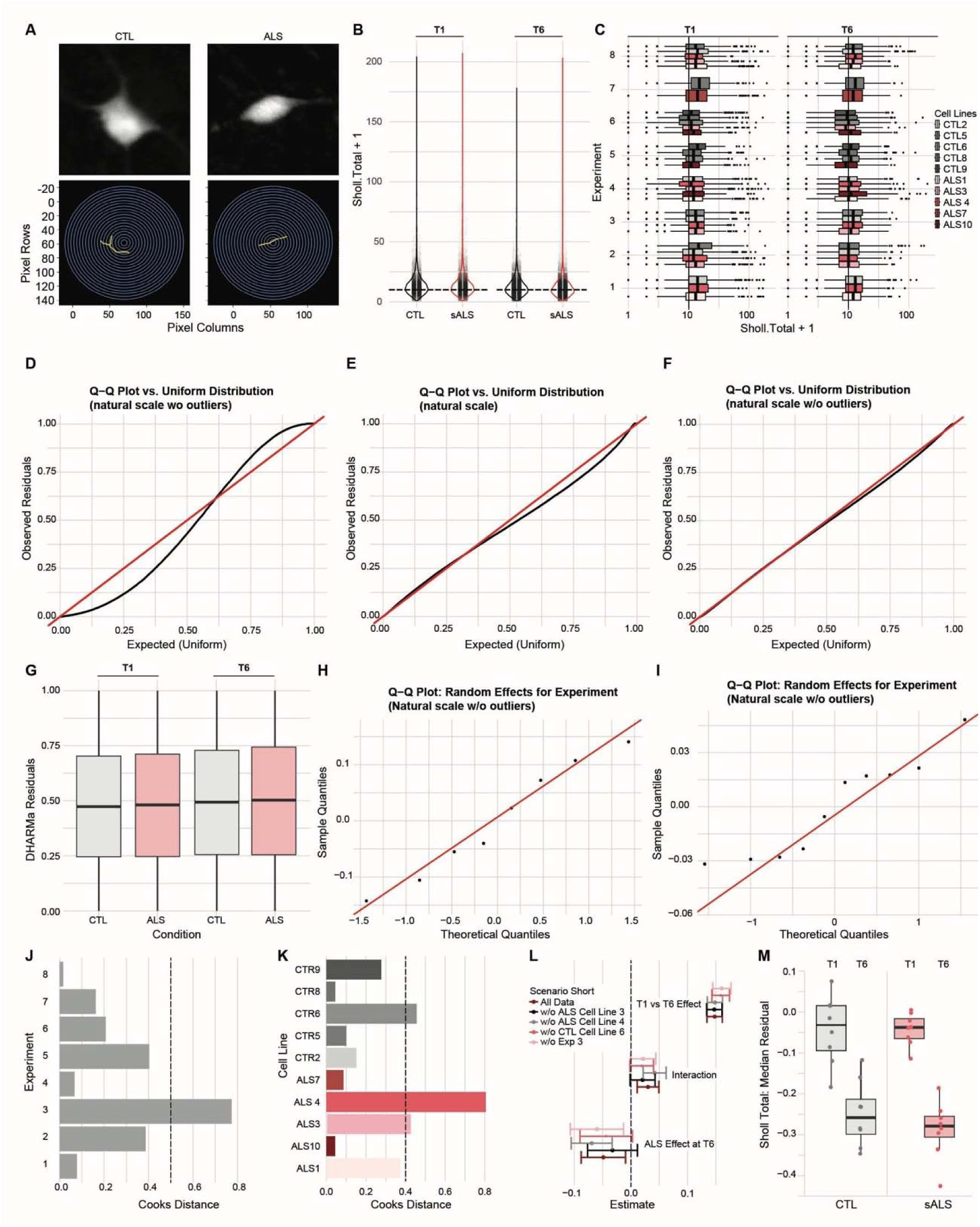
Modeling changes in the mean sholl feature between ALS and control lines. **B.** Example crop images of control (CTL) and sALS iMNs as acquired by RM. **C.** Violin plots depicting the distributions of sholl observations across sALS and control lines over two time-points labeled T1 and at T6. These correspond to days *in vitro* (DIV), T1 (∼ DIV27) and at T6 (∼ DIV33). The dashed horizontal line is drawn at a sholl count of 10. **D.** The distribution of the sholl observations across the 3 experimental batches and 10 patient cell lines are plotted at timepoints 6 and 1. The 6 control lines are colored in different shades of grey while the 4 sALS lines are colored in different shades of red. Vertical solid lines are drawn at a sholl count of 10. **E.** Q-Q plot of the distribution of scaled residuals from the GLMM fit of sholl count assuming a Poisson probability distribution demonstrating the deviation of this distribution from uniformity. The x-axis represents the quantiles from a uniform distribution while the y-axis the quantiles of the observed scaled residuals. The experimental batch and cell lines were modeled as random effects while the sALS status, time-point and the interaction between these two variables were modeled as fixed effects in this GLMM fit. **F.** Q-Q plot of the distribution of scaled residuals from the GLMM fit of sholl count assuming a Negative Binomial probability distribution results in a better fit compared to C. The interpretation of the x and y-axis labels are the same as in **C**. **G.** Q-Q plot of the distribution of scaled residuals from the GLMM fit of sholl count without individual outlier observations and assuming a Negative Binomial probability distribution results in an even better fit compared to D. The interpretation of the x and y-axis labels are the same as in **C**. **H.** Distributions represented as boxplots of the scaled residuals from the GLMM fit of sholl count corresponding to the model used in E. among the four groups of observations corresponding case and control status at each of the two time-points. **I.** Q-Q plot of the distribution of random effects corresponding to experimental batch from the GLMM fit of sholl count corresponding to the model used in E. The x-axis (Normal Scores) represents the quantiles from a standard normal distribution while the y-axis the quantiles of the observed residuals that were centered by their mean and scaled by their standard deviation (Standardized Residuals). **J.** Q-Q plot of the distribution of random effects corresponding to cell line from the GLMM fit of sholl count corresponding to the model used in E. The interpretation of the x and y-axis labels are the same as in **G**. **K.** Cook’s distance estimates for each of the eight experimental batches using the GLMM fit of sholl count corresponding to the model used in E suggested that experimental batch 3 has a strong influence on the parameter estimates based on a threshold of 4/8 (the vertical dashed line). **L.** Cook’s distance estimates for each of the nine cell lines using the GLMM fit of sholl count corresponding to the model used in E suggested that sALS cell lines 3 and 4 and control cell line 6 strongly influence the parameter estimates based on a threshold of 4/10 (the vertical dashed line). **M.** Estimates (including corresponding 95% confidence intervals) of changes in three parameters of interest using all the data and also after removing all observations from the experiment 3, ALS cell lines 3 and 9 and control cell line 6 individually. The three parameters of interest are *ALSeffectAtT6* that represents the difference in rates of sholl counts between the sALS and control lines at time T6, *T1vsT6effect* that represents the difference in rates of sholl counts between time-points T1 and T6 for the control cell lines and *Interaction* that represents the additional change in the rates of sholl counts from time points T6 to T1 for the ALS lines over the corresponding change in the control lines. **N.** Distribution of the median of the residuals across all observations within each of the 8 experimental batches by ALS status and time-point of the observations. The scaled residuals are computed using a GLMM fit to the sholl counts and modeling the experimental batch and cell line as random effect and excluding the ALS status and the time-point of these observations. In summary, the mean Sholl values decline by 0.15 units between time points 1 and 6 in control lines (p = 2.2e-16), with sALS lines showing an additional 0.03 unit decline (p = 0.002) but 0.05 units less overall than controls (p = 0.015).

Consistent with prior reports of impaired neuronal morphology in sALS^62–64^, here we show new results that recapitulate these phenotypes in our sALS cohort but in different patient lines that previously reported. However, by leveraging an interaction term in our model, we extend beyond static single-timepoint measurements to capture the temporal dynamics of neurite degeneration and present novel findings in the context of sALS. This approach affords a substantially richer characterization of disease progression in neurodegeneration.

#### Other Applications

Aside from cellular assays, RMeDPower2 can be useful across a broad range of applications (**Figure 6**) involving repeated measures with non-normal, discrete or binary distribution (where GLMM are needed), and also in situations where there is a need to model the effect of a confounder to the association between the chosen predictor and response variable (e.g., sex of subjects from which the cell lines are derived). In sections 2.2-2.4 in **Note S1**, we illustrate the utility and implementation of RMeDPower2 in the analyses of single cell RNA-seq, of mouse behavior and also electrophysiology data^73–75^. To avoid the pseudo replication bias associated with the estimation for single-cell differential gene expression studies^76^, Poisson and Negative Binomial-based mixed effects models have been proposed in the literature^28, 77^. These same papers have found their performance wasn’t as good as the simpler pseudo-bulk-based models that were also evaluated. One of the key insights we derive while diagnosing these models using RMeDPower2 is that they are not good models for the data we tested them on. This could explain their low performance.

**Figure 6.**
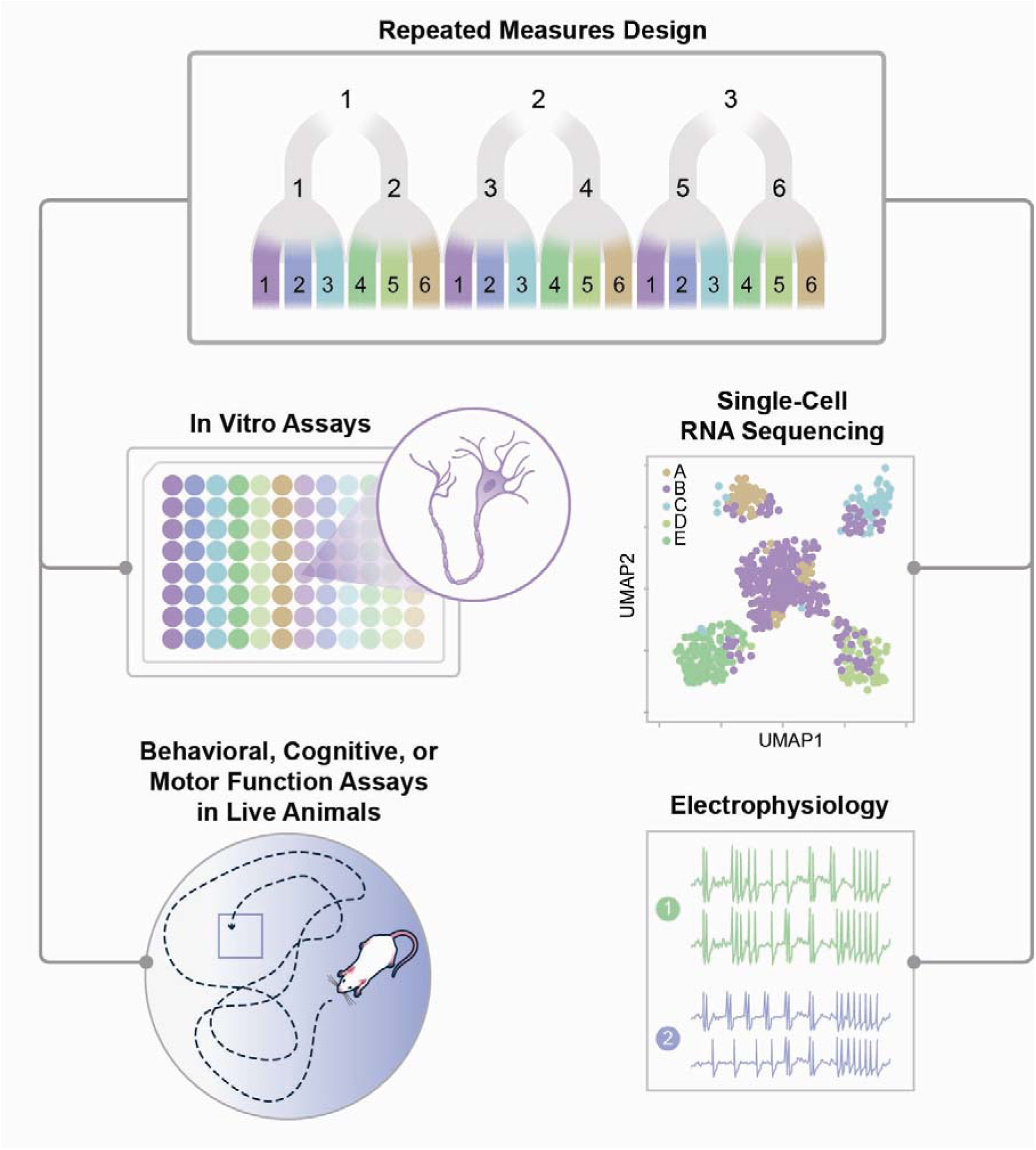
Application areas for RMeDPower2 involving repeated measures designs. The application areas include cellular, single cell RNA-seq, mouse behavior and mouse brain electrophysiology assays. Implementation of models using RMeDPower2 using data from each of these areas are provided in the **S1**.

## Discussion

Here we have showcased the RMeDPower2 R package which supports both categorical and continuous covariates, covariate-by-condition interaction terms, and random slopes — capabilities that extend the range of experimental designs that can be accommodated. We demonstrated the utility and diverse functionality of this tool in detecting differences in iPSC– derived sALS iMNs compared to unaffected controls. Using a simple statistical tool such as a two-sample t-test in GraphPad Prism (a tool commonly used in biology) to compare the mean of neuronal cell soma perimeter observations across all ALS lines versus all control lines, one would have overestimated (by a lot) the significance of the difference between control and ALS perimeter sizes: the simple t- test gives a p-value of 1.1×10^-7^ whereas the LMM gives a p-value of 0.03. This is because the t-test incorrectly treats observations made within each experimental batch and for a given cell line as being independent of each other, whereas the LMM accounts for the fact that they are not.

We model a Sholl-derived neurite feature in sporadic ALS versus control motor neurons over time, using time as a categorical covariate and interaction term. We also show that we were adequately powered to make this claim and that the statistical model we based our findings on met the required assumptions. Further, one can easily model the interaction of time which one cannot easily do with typical statical software packages that are currently available.

RMeDPower2 is not only applicable to simple two-group comparisons but can also capture the temporal dynamics of neuronal phenotypes — a particularly important capability for longitudinal iPSC-based studies. To further support users with limited statistical background, we have also added a step-by-step tutorial (Note S1) that guides biologists through multiple data types and experimental designs, and we have included a recommendation in the Methods section that users unfamiliar with linear mixed models consult with a statistician during the analysis phase. Taken together, the core aim of RMeDPower2 is to make rigorous, mixed-effects-based experimental design and analysis accessible to the broader biological research community, without sacrificing statistical integrity.

Experimental reproducibility has been a major challenge to advancing our understanding of biological phenomena and disease mechanisms, and to our ability to devise powerful therapeutic approaches. Indeed, the majority of published empirical observations cannot be reproduced^4–9^, and misleading findings may steer future directions or studies in the wrong direction. The wasted money and effort to replicate faulty, underpowered studies creates a major hurdle in scientific discovery and breakthrough. There are two major underlying causes of this lack of reproducibility: 1) experiments that are designed poorly and are vastly underpowered for the phenotype of interest; and 2) the lack of statistical models that adequately account for all of the observed variation that occurs within biological replicates. Variability is known to come from several sources—including cell line variability, batch-to-batch variation in reagents, user variation in handling methods, and instrument or technical variability—and can also be unexplained. Typical statistical models such t-tests and ANOVA that are often used in user-friendly software such as GraphPad Prism fail to account for the effects that come from this variation. Therefore, effective statistical models are needed to guide experimental design as well as to help researchers apply adequate statistical power estimation. To examine the utility of RMeDPower2 and how it compares with other statistical tools commonly used, we contrasted the different attributes and utilities available with each package in **Table 1**. We examined 13 aspects of functionality or implementation that are commonly used in situations where one is working with data from repeated measures experimental design scenarios. The availability of each of the 13 features across 18 tools are indicated. RMeDPower2 clearly brings together the most functionality across these defined features **Table 1**.

**Table 1.**
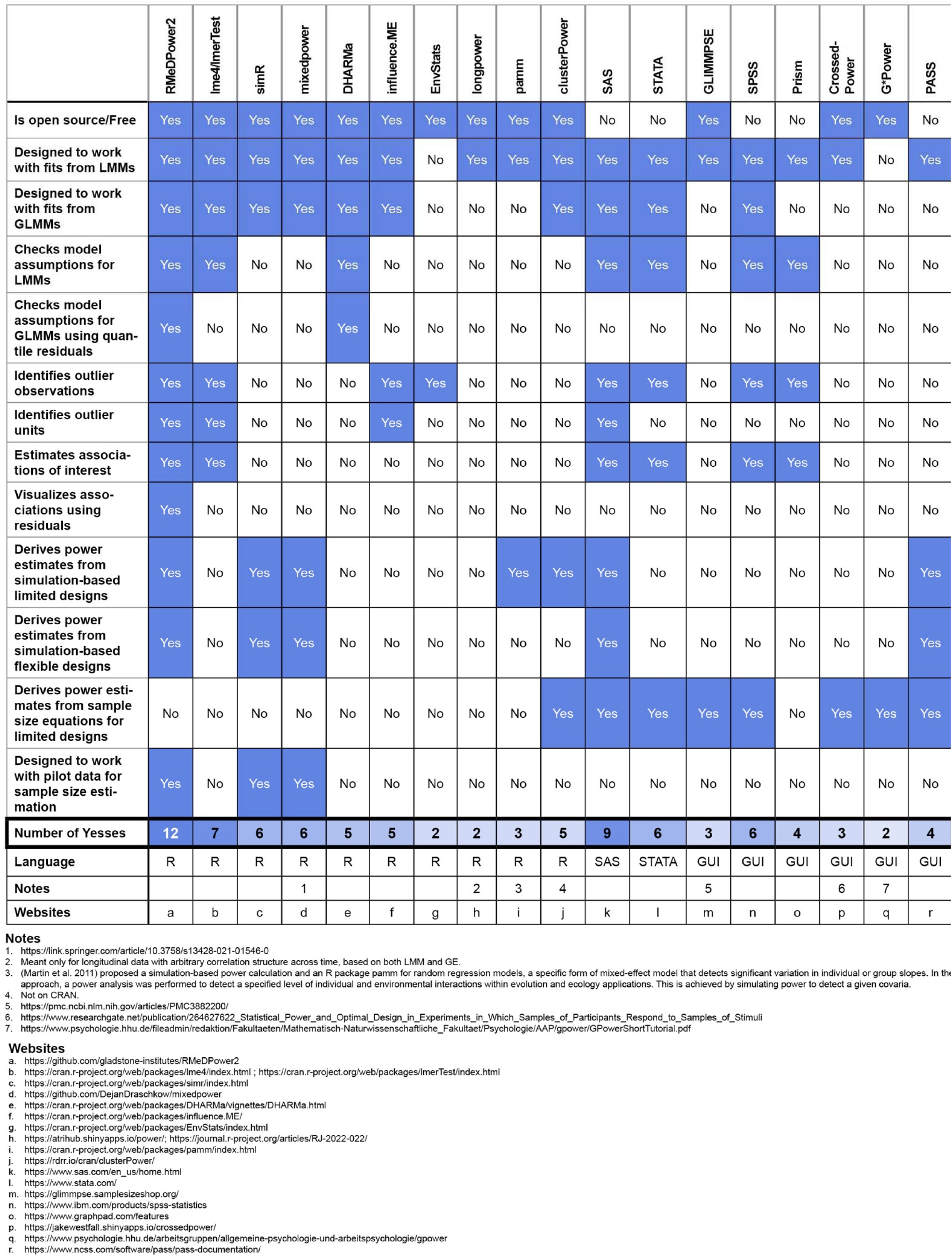
Comprehensive comparison of RMeDPower2 to other statistical tools. RMeDPower2 has the greatest functionality compared to other statistical software.

RMeDPower2, in the setting of repeated measures studies, provides tools to researchers that will help them navigate the efforts needed to move from an idea for a scientific hypothesis, to the experimental planning for generation of the data needed to test the hypothesis, to testing assumptions for an appropriate statistical model and also to the identification of outliers for the data collected, and finally to test, estimate and visualize the parameter of interest associated with the scientific hypothesis. Throughout this process, there are objective measures that one can use to make appropriate modeling choices and/or for the removal of outliers.

### Limitations

Although RMedPower2 can handle a variety of biological experimental designs at a level that other power packages have not attempted, it still has some limitations. For example, the current version of RMedPower2 is made to accept one response variable and one condition/confounder/predictor variable (modeled as fixed effects) with a possible interaction between the two variables and any number of experimental/technical variables (modeled as random effects) and also the possibility of including a random slope variable. Therefore, if the experimental design does not meet these specifications, the package cannot be applied. Box-Cox and other transformations^78, 79^ for continuous data represent more general methods than the log-transform to meet the normality assumptions. Future work to develop these features of RMedPower2 will provide even wider utility for users.

## Methods

**Note**: Users who are not familiar with mixed effects models are strongly encouraged to consult a statistician to verify their parameter choices and the interpretation of the conclusions that they draw from the results provide by RMeDPower2’s functionality.

### Specification of the design for the data using the *RMeDesign* S4 object

Users will first have to define an S4 object of class *RMeDesign* that will force them to answer the relevant design and outlier identification parameter questions such as: what is the response of interest? What is the predictor for this association? Is the predictor a categorical or continuous variable? Is there a covariate or confounder? Is the covariate a categorical or continuous variable? Is there an interaction between the predictor and the covariate? What is their chosen Type I error for outlier identification? What are the experimental factors capturing the repeated measures? If there are multiple factors, then are they linked in a *nested* or *crossed* manner? An experimental factor B is considered *nested* within another experimental factor A if all levels of factor B occur under only one level of factor A. For example, if an experimenter is measuring a set of variables from cells that are growing in multiple assay plates, this type of experiment would be considered to be *nested* within a given experimental batch of observations. On the other hand, an experimental factor B is considered *crossed* with another factor A if all levels of factor B can occur with any level of factor A. For example, cells from the same cell lines could be assayed across multiple experimental batches, and in this case the different cell lines would be considered to be *crossed* with the experimental batches. Is there a variable (predictor or covariate) that would represent a random slope? In case the response of interest is a proportion, users can use the ‘total_column’ parameter to provide the total number of samples over which the proportion is estimated. For example, this option is useful when testing cell-cluster membership associations in data sets such as single RNA seq data.

### Specification of parameters for sample size calculation using the *PowerParams* S4 object

To determine the optimal sample size of a set of experiments, we used simulation-based sample size estimations with user-specifications that relay experimental variables of interest (e.g., animals, plates, experimental batches). Users can also select the upper bound of number of samples to test, type I error, the effect size, and Intra Class Correlation (ICC) coefficients for variance components for the experimental variables.

### Specification of the error distribution using the *ProbabilityModel* S4 object

Users can specify a probability distribution for the data generating distribution, which could either be normal or chosen from a set of other distributions in the ‘*ProbabilityModel*’ object using the *family_p* attribute. Poisson, Binomial, Negative Binomial and Gamma probability distributions are the possible non-normal distributions one could choose from.

### Derivation of parameter estimates and visualization of the association using *getEstimatesOfInterest* function

The *lmer* function in the R package lmerTest^48^ is used to fit the mixed models and get estimates for the parameter of interest for models based on normal distributions, while the *glmer* function in lme4^47^ generates corresponding estimates for models based on non-normal distributions. The association of the response and predictor is visualized by examining the distribution of the residuals of the model fit excluding the predictor variable, but including the remaining variables used in the model. If the predictor is a categorical variable, then the distributions of the mean/median of the estimated residuals across all observations within each level of the first random effect (or the experimental column that is specified first in the design object) are visualized as a function of the levels of the predictor variable as a boxplot. Otherwise, the association is visualized as a scatter plot with best-fit lines between the residuals and the predictor.

### Checks for model assumptions using *diagnoseDataModel* function

The assumptions underlying mixed effects models are described in detail elsewhere^47^, and they require a linear relationship between the response of interest and the chosen predictor. Mixed effects models also require homoscedasticity such that the variance of the residuals across all fitted values displays a normal distribution for the estimated random effects. For situations where the error distribution is normal, the *plot* function that is part of lme4 package is used to visualize the residual versus fitted values. Further, the scale-location plot allows the user to assess the degree of violation of the linearity and the homoscedasticity. In addition, the models specified with the normality assumption are also evaluated based on fits to the log-transformed response values, while the Quantile-Quantile (QQ) plots are used to verify the normality assumptions for the residuals and the random effects. However, for the case of non-normal distributions, the interpretation of the (Pearson or Deviance) residuals versus fitted plots are not straightforward and can be difficult to diagnosis^80^. For this reason, we incorporated the DHARMa package, which is used to generate randomized quantile residuals and generates plots that are easier to assess for distributional, linearity and homoscedasticity assumptions. The Rosner’s test^56^ implemented in the EnvStats package is used to identify individual outlier observations using the residuals from models assuming a normal error distribution, while the same test is used on the quantile residuals transformed to a standard normal distribution. The *influence* function that is part of the influence.ME package is used to find outlier units or sets of individual observations grouped by a given random effect (or experimental variable) that has an effect (as quantified by the Cook’s distance^59^) on the estimated parameters of the chosen model. It is up to the user to examine the various outputs from this function, using the experimental and biological background information for the data to decide on either changing parameters (e.g., changing the assumed probability generating distribution) or working with the model based on log-transformed response values and/or the removal of individual outlier observations and experimental units.

### Sample size calculations using the *calculatePower* function

The functionality here broadly builds upon the power simulation structure of the simr package^41^ to enable power calculations based on a generalized LMM. As in simr, *calculatePower* simulates response variables, which correspond to direct measurements, based on given variance components of the random effects estimated from a user-provided pilot dataset. The power estimation is based on the assumption that the relationship between the condition variable and the response variable in the pilot dataset, as captured by the effect size, is a true association. The simulated response variables are refitted into a LMM or GLMM and the total number of significant association results are counted to estimate power to detect the association at a chosen Type I error level. In cases where the pilot dataset does not have enough data for each experimental variable, users can provide ICC values which can be used to estimate variance components. These will come from prior knowledge based on empirical data. Here, a high ICC value indicates that the variable contributes to a relatively high variance relative to the overall variance estimate of the response variable.

### iPSC differentiation and imaging

iPSC lines from control and patient derived iPSCs from sALS patients were obtained through the Answer ALS (AALS) consortium^68^. AALS is the largest repository of ALS patient samples, and contains publicly available patient-specific iPSCs and multi-OMIC data from motor neurons derived from those iPSCs—including RNA Seq, proteomics and whole-genome sequence data— all of which are matched with longitudinal clinical data^68, 81^.

iPSCs were thawed in mTESR Plus medium supplemented with ROCK inhibitor and cultured on growth factor–reduced Matrigel. Cells were maintained in mTESR Plus and passaged approximately 1:5 using ReLeSR or Versene. All lines were expanded, cryopreserved in mFreSR, and stored in liquid nitrogen at ∼ 8 million cells per vial.

For the cell soma perimeter experiments, 7 control lines [9BP3iCTR (female), 2AE8iCTR (female), 5NWTiCTR (female), 6ZBiCTR (male), 0YX7iCTR (male), 2518iCTR (male), 2PFYiCTR (male)] and 6 sALS lines [9DT1iALS (female), 5ZP7iALS (female), 5GBQiALS (male), 5FFPiALS (female), 6WKViALS (female), 3DE3iALS (female)] were used. We differentiated 8 separate experimental batches of each line into iMNs, transduced with the Synapsin::GFP morphology marker from SignaGen and imaged on day ∼25 using RM^15, 17–20, 22, 66, 67, 82^ as previously described^68^.

For the sholl experiments, 10 iPSC lines [five controls, 2AE8iCTR, 5NWTiCTR, 6GNGiCTR, 6ZKZiCTR, 7PFJiCTR, and five sALS, 0XLRiALS, 3DE3iALS, 5GBQiALS, 6WKViALS, 8RBNiALS] were subjected to a slightly different motor neuron differentiation protocol that was also based on previous reports^68^ but contained a freezing step for neuronal progenitors on Day 16 as previously described^83^. Briefly, iPSCs were grown to ∼90% confluency, dissociated with Accutase, and replated at 4 million cells per T25 flask in mTESR Plus with ROCK inhibitor. When cultures reached ∼90% confluency (1–2 days later), differentiation was initiated (Day 0) using Stage 1 neural induction media containing IMDM/F12, N2, B27, NEAA, and the small molecules LDN193189, CHIR99021, and SB431542. Cells were fed daily with Stage 1 media until Day 6. On Day 6, cells were transitioned to Stage 2 patterning media, which consisted of Stage 1 media supplemented with all-trans retinoic acid (ATRA) and SAG to induce motor neuron progenitor cell (MNPC) fate. Cells were split 1:2 and cultured in Stage 2 media until Day 13, with media changes every other day. At Day 13, MNPCs were cryopreserved.

For neuronal differentiation, MNPCs were thawed and plated onto poly-L-ornithine and laminin coated plates in Stage 3 maturation media, consisting of Stage 2 media supplemented with db- cAMP, Compound E, DAPT, ascorbic acid, BDNF, and GDNF. Cells were maintained in Stage 3 media with media changes every other day until Day 20. Beginning at Day 20, cells matured into neurons (MoNOs). Cells were transduced with GEDI^65^ lentiviral biosensor around Day 17 (MOI 5–10). On Day 20, neurons were dissociated and replated into coated 384-well imaging plates at 30,000 cells per well. After Day 25 the cultures were considered induced motor neurons (iMNs). Imaging experiments were performed from approximately Day 27–36.

### Image Analysis

High-content robotic microscopy (RM) was used to acquire large image montages (6144 × 6144 pixels) of iMNs expressing the Synapisn: EGFP or the GEDI biosensor from control and sALS.

The raw images are run through our custom-built imaging pipeline assembled in Galaxy software^84^ in order to obtain object crops that contain a single cell from raw image tiles. Crops containing single cells are then contrast-enhanced with 1.5% saturation, normalized, denoised. Pixels that deviate from their neighborhood median by threshold are removed using FIJI^85^. A smoothing algorithm is applied to remove extraneous debris/cells around the central cell. The crops were then subjected to our morphological feature-based pipeline, and the perimeter of each neuronal soma was captured using the contour ellipse^86^ function adapted for use in python^70, 87^.

For each cropped cell, a series of quantitative features describing cell shape and perimeter, or complexity by the sholl intercept as captured by Sholl Intersections Analysis^88^ with Skeletonization transformation^89^ were computed using Python image-analysis libraries (OpenCV, Scikit-Image, Pillow).

### Datasets

#### Cell perimeter feature

Three datasets derived from the above cell perimeter-based experiments were used for the analysis described in this manuscript. The cell lines are de-identified in these datasets since the main focus here is on the illustrating and explaining of the functionality that we incorporated into RMeDPower2. The first dataset (**Tables S6)**, representative of a pilot data set, is composed of data derived from three (of the original eight) experimental batches with 10 cell lines (4 ALS and 6 control lines), and 6 cell perimeter-based responses. The first response variable, ‘cell_size1’, corresponds to the measured perimeter values in both the control and the ALS lines, while observed values only for the ALS lines are increased by 4, 8, 18, 26, and 36 units respectively for the remaining 5 responses (’cell_size2’, ‘cell_size3’, ‘cell_size4’, ‘cell_size5’ and ‘cell_size6’). These six responses in order imply ALS-specific effect sizes of 1.1, 1.56, 2.02, 3.16, 4.08 and 5.2 based on an estimate of the difference between mean perimeter between the ALS and the control lines derived from fits of each of the six raw perimeter responses to LMMs with the experimental batch and cell-lines being modeled as random effects. The ‘cell_size2’ response is used in the models implemented for this manuscript. The largest 5% of the ‘cell_size2’ perimeter values across all the experiments and cell-lines are filtered out from these data, resulting in 2458 observations. The second dataset represents another pilot experiment that consists of one plate, two cell lines, and one response variable representing cell area, corresponding to situations where there are not enough replicate experimental units. (**Table S8**). Since data in **Table S8** cannot be used for estimating variance components due to an insufficient number of replicates of experimental batches, plates and cell-lines, these data are used to demonstrate the situation where users are required to provide Intra Class Correlation coefficients (ICC) values for each of the experimental batch, plate and cell-line factors. Based on given ICC values, the package will calculate variable components and simulate the response variable to assess power. The third dataset (**Table S7**) consists of cell-perimeter measurements with observations from all 8 experimental batches and the 13 cell-lines. This includes the cell perimeter values corresponding to the 6 effect sizes described above for **Table S6**. This data set is used to validate the sample-size predictions for the required number of experimental batches as a function of the effect-size for the ALS-specific differences in mean cell perimeter. These predictions are based on 56 pilot data sets, each with 3 experimental batches (sampled from the original 8 batches) that are required to have adequate statistical power (80% or a Type II error rate of 0.2, i.e., if one repeats experiments 100 times, each time by sampling observations from three new experimental batches, then in 80 of these experiments one would detect a significant difference between the ALS and the control lines with the assumption that there are true differences) to detect changes at a Type I error rate of 0.05 (i.e., if one repeats 100 times experiments by sampling observations from three new experimental batches, then in at least 5 of these experiments one would detect a significant difference between the ALS and the control lines when there aren’t any differences) in the mean cell perimeter between the ALS and control cell lines. For cases where 80% power was never achieved in the range of 1 to 15 experimental batches, the value 15 was assigned as the required number of experiments.

#### Sholl feature

The data is available in **Table S9**. This is based on a random 10% down sampling of the original experimental data in order to speed up the computations for the resulting model fits, diagnostics and power estimation. The original data had 601,030 observations and the down sampling was performed within each experiment and cell line combination. The main conclusions are not altered we draw from the data are not altered based on alternate random sampling of the data (data not shown).

## Operation and Code Availability

The source code of the RMeDPower2 is available on https://github.com/gladstone-institutes/RMeDPower2 and on CRAN https://cran.r-project.org/web/packages/RMeDPower2/index.html. To reproduce the results of examples in this paper, R >= 4.0.4, simr >= 1.0.5, and lme4 >= 1.1-27.1, lmertest >=3.1-3 are required based on RMeDPower2.

## Acknowledgments and Funding

We would like to acknowledge these sources of funding that supported this work: NSF 1761941, NIH R37NS101996, P01 AG54407, R01 LM013617, NIH U01MH115747, ALS FindingACure, Target ALS IL-2023-C4-L2, U.S. Army Medical Research Acquisition Activity (USAMRAA): W81XWH-22-1-0721, W81XWH-20-1-0710 W81XWH-18-1-0696, and CIRM (DISC0-13914), CIRM (DISC0-16039), NIH P01AG073082, the ALS Association, the Robert Packard Center for ALS Research, the JSRM Foundation, Tad Taube Foundation and Answer ALS. We also would like to thank all Tami Tolpa, Alex Pico, Kathryn Claiborn, Kelley Nelson and Gayane Abramova that helped on various aspects of this work.

## Contributions

Conceived and designed the study. SF, JAK, RT

Designed the overall work and oversaw all development MS, JAK, RT

Wrote the manuscript with input and edits from all the authors MS, JAK, RT

Provided project leadership JAK, RT, MS

Differentiated iPSCs from control and ALS lines into iMNs and subjected to RM imaging EV, KR, NA

Performed integrative analysis and computational modeling MS, NA, SL, RT

Developed the statistical models MS, RT

Optimized the statistical models MS, NA, SL, RT, JAK

Ran and optimized the R code MS, NA, SL, RT

---

**Figure S1.**
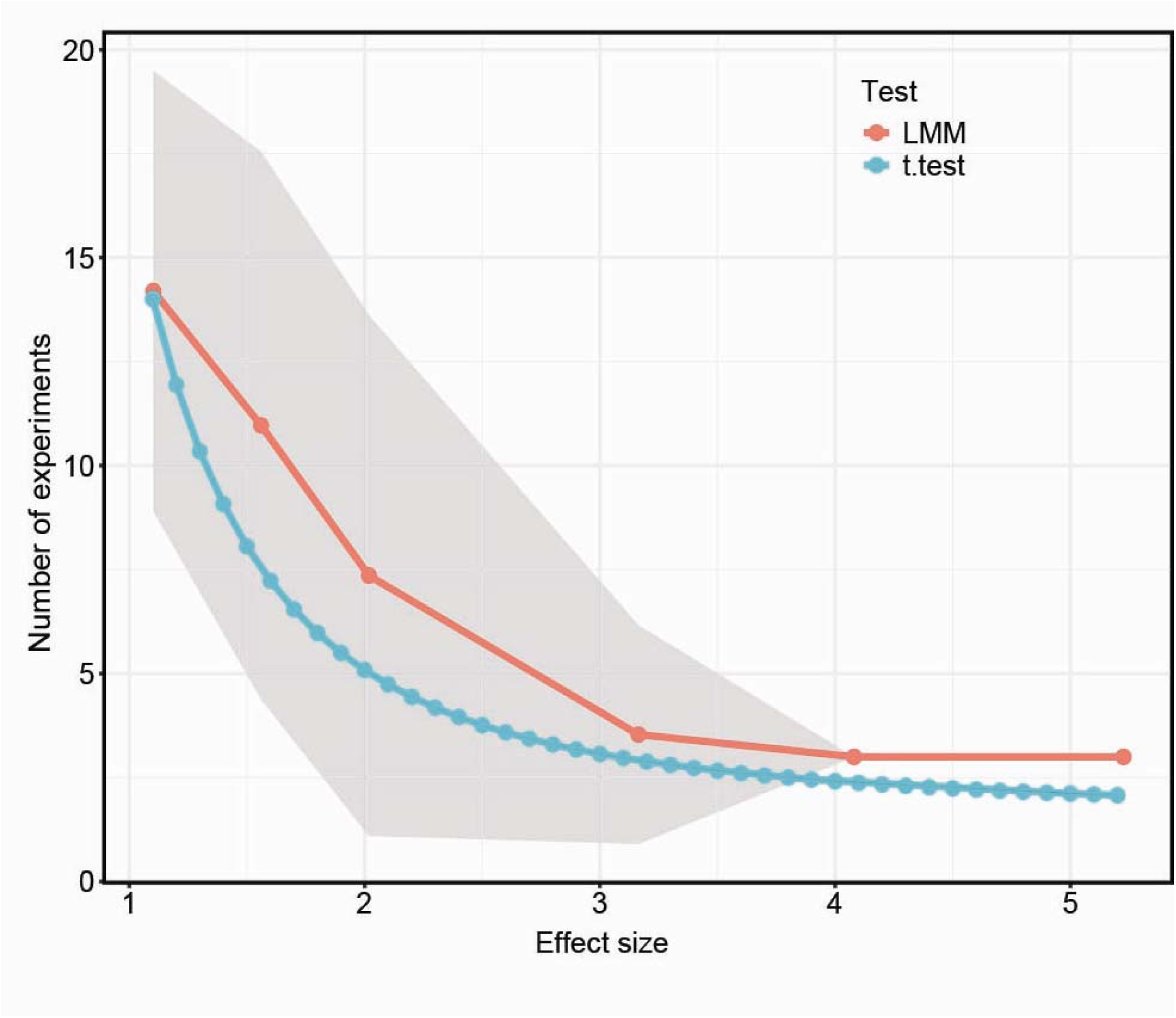
The number of experiments required to achieve 80% power at various effect sizes using data provided in Table S3. Results are calculated from the mean number of experiments estimated by simulation (red) using RMeDPower2 and the two-sample t-test (blue), where gray intervals represent standard deviations across sample size estimates derived from 56 pilot data sets, each with 3 experimental batches sampled from the 8 experimental batches in **Table S3**. The models are based on cell perimeter observations with experiments and cell-lines models as random effects with the ALS status of the cell-lines being modeled as a fixed effect.

**Figure S2.**
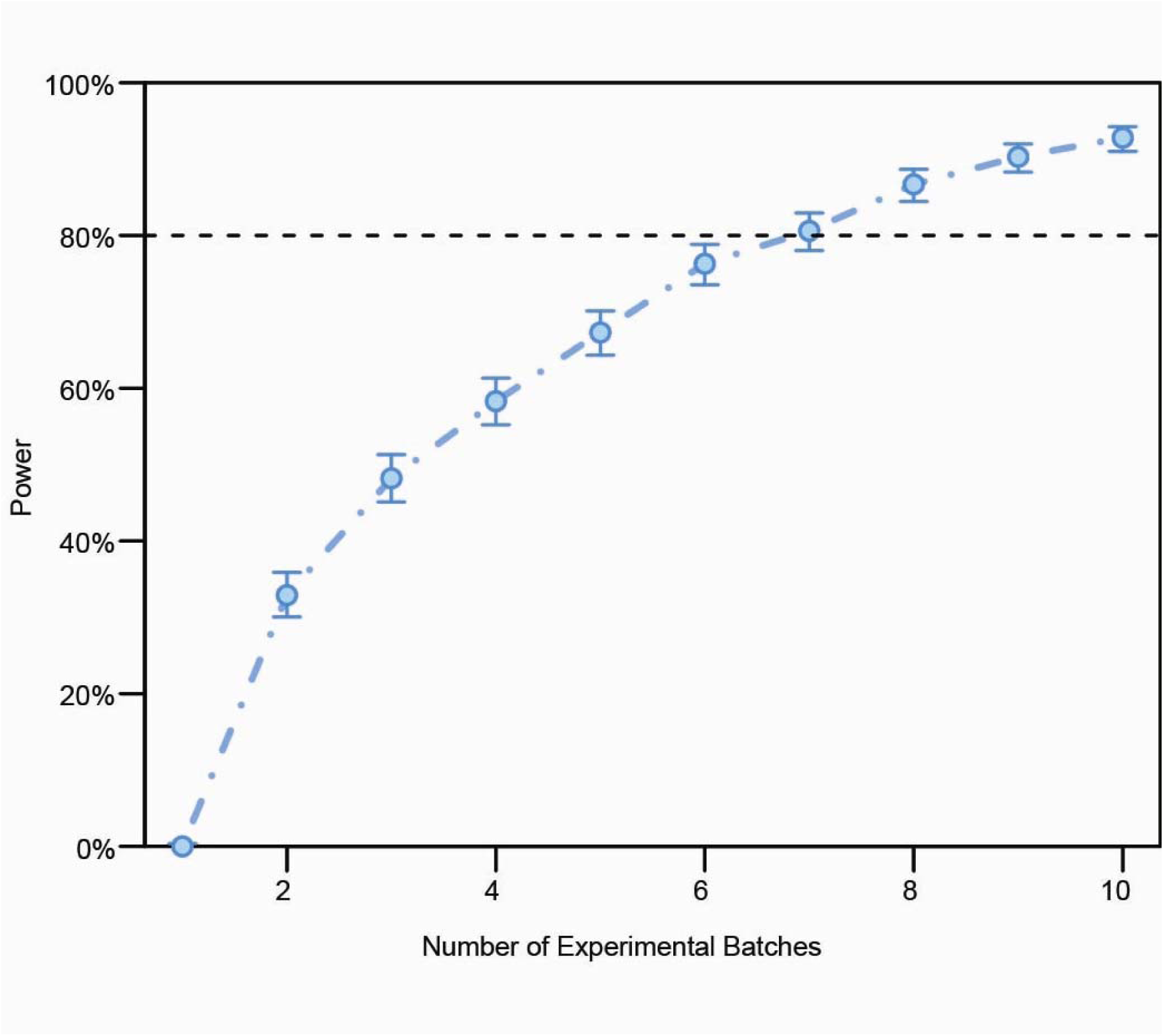
Sample-size estimates as a function of the number of experimental batches using pilot data. Here we that estimate the sample size required to allow for estimates of variability due from the cell perimeter dataset (**Table S8**) that occur from three technical sources – experimental batch, plate and cell-line. In a scenario like this a user can provide estimates of variability from these three sources using Intra-Class Correlation (ICC) coefficient values. ICC values of each of the above technical sources were set to 0.8, 0.05, and 0.05, respectively.

**Table S1.**
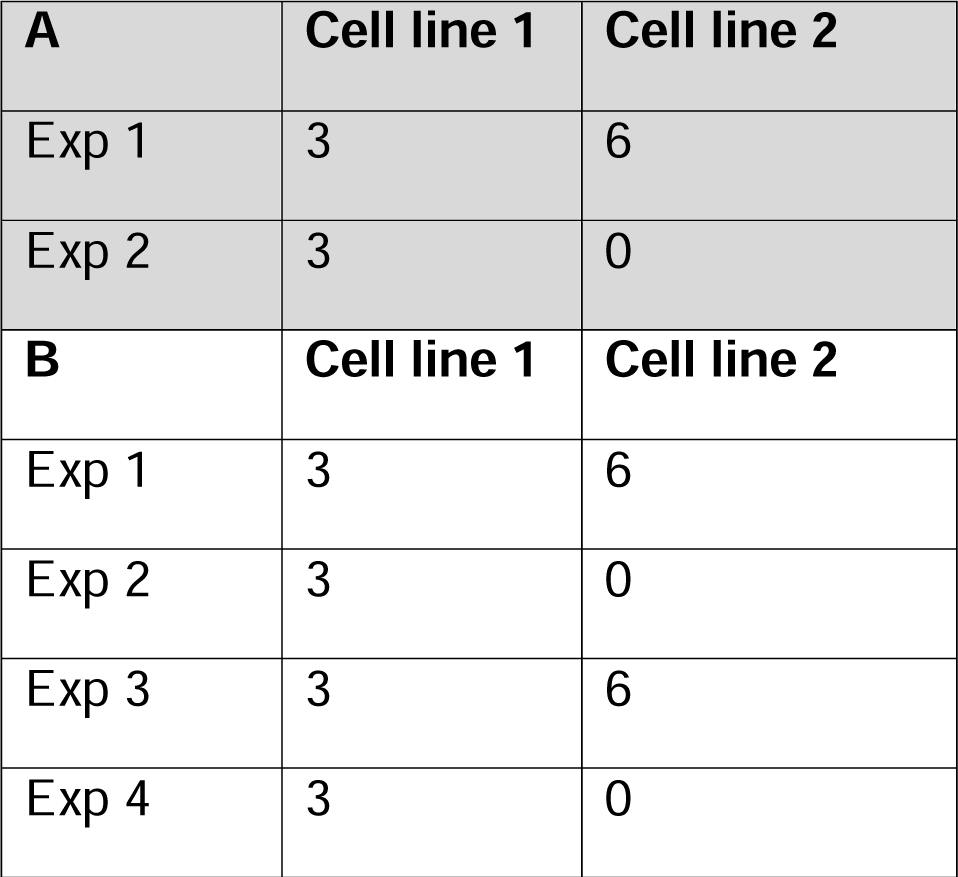
Example of changes in cell counts in a simulation of experimental batch variability. (A) Original data structure and (B) changed data structure after simulation. Simulation based on level=1 keeps the same structure and increases the number of experiments. This describes the situation where the pilot study involved 2 experiments, in the first experiment two cell lines are used with 3 and 6 cells assayed, respectively, and in the second experiment only 3 cells from the first experiment are assayed.

**Table S2.**
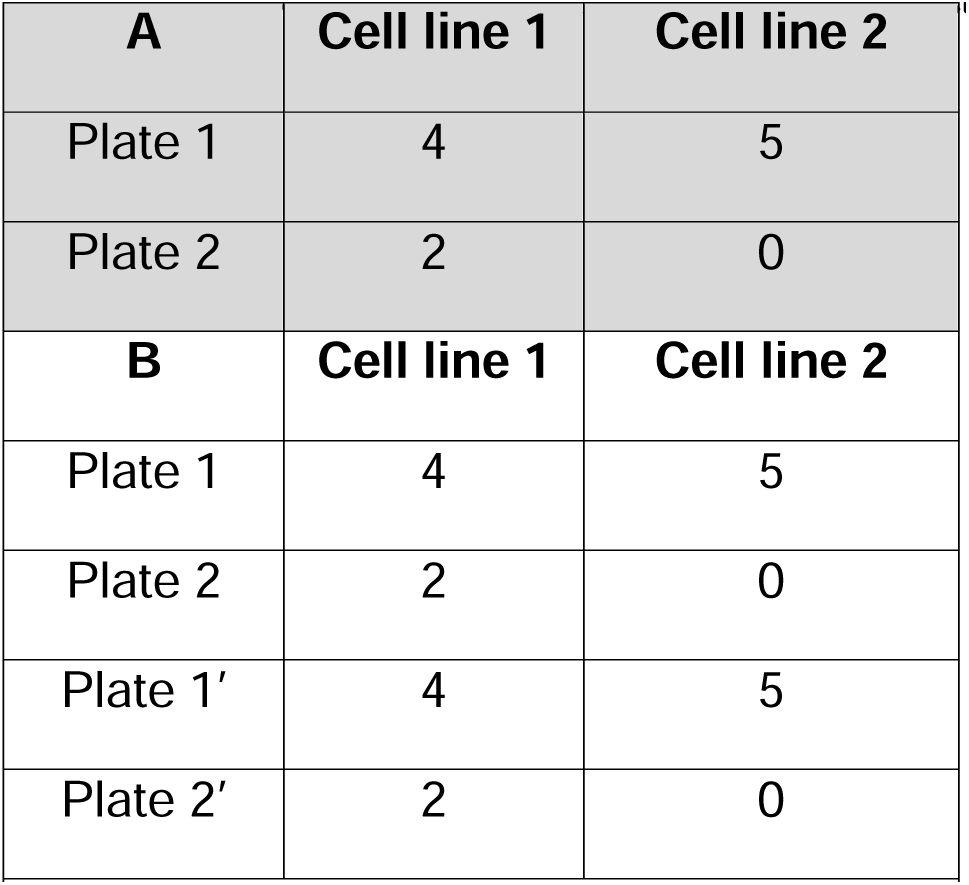
Example of changes in cell counts in a simulation of plate variability. (A) Original data structure and (B) changed data structure after simulation. Simulation based on level=1 keeps the same structure and increases the number of plates.

**Table S3.**
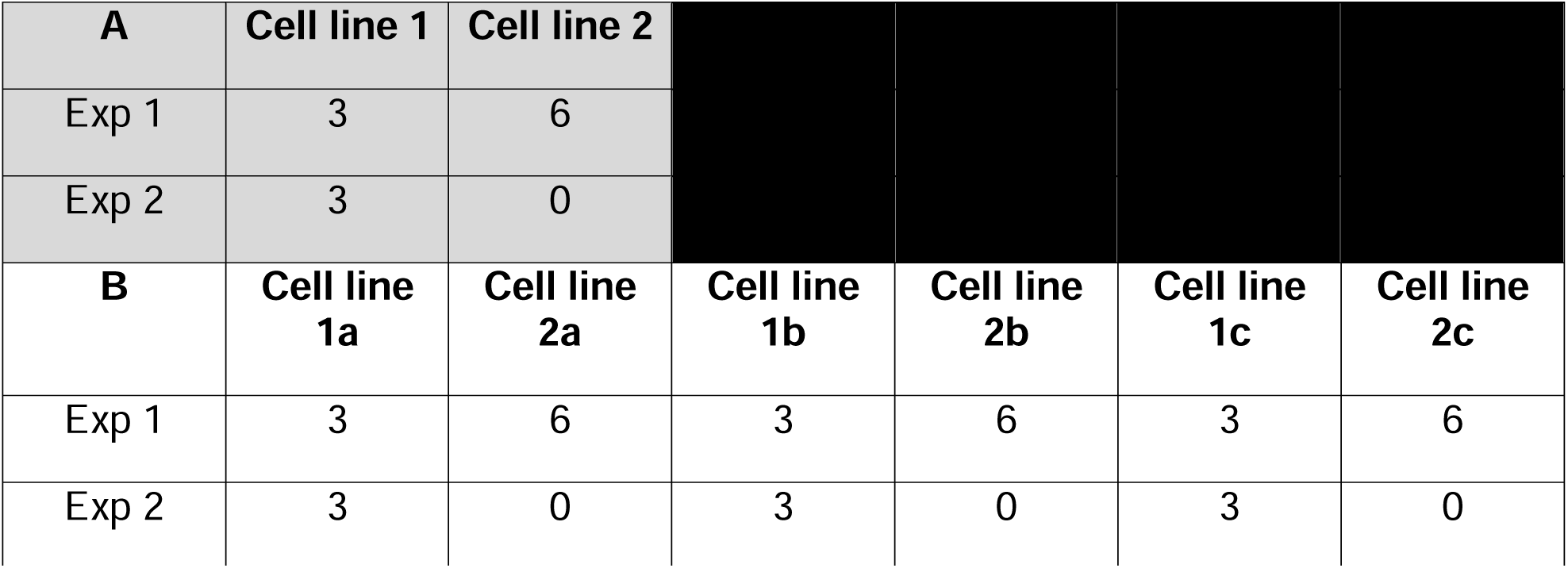
Example of changes in cell counts in a simulation of cell line variability. (A) Original data structure and (B) changed data structure after simulation. If level=1 is set, RMeDPower simulates new cell lines by inheriting the experimental design structure from the existing data. An example of a simulation in tabular format shows the change in the number of cells per cell line and experiment after simulation. Simulation based on level=1 keeps the same structure and increases the number of cell lines.

**Table S4.**
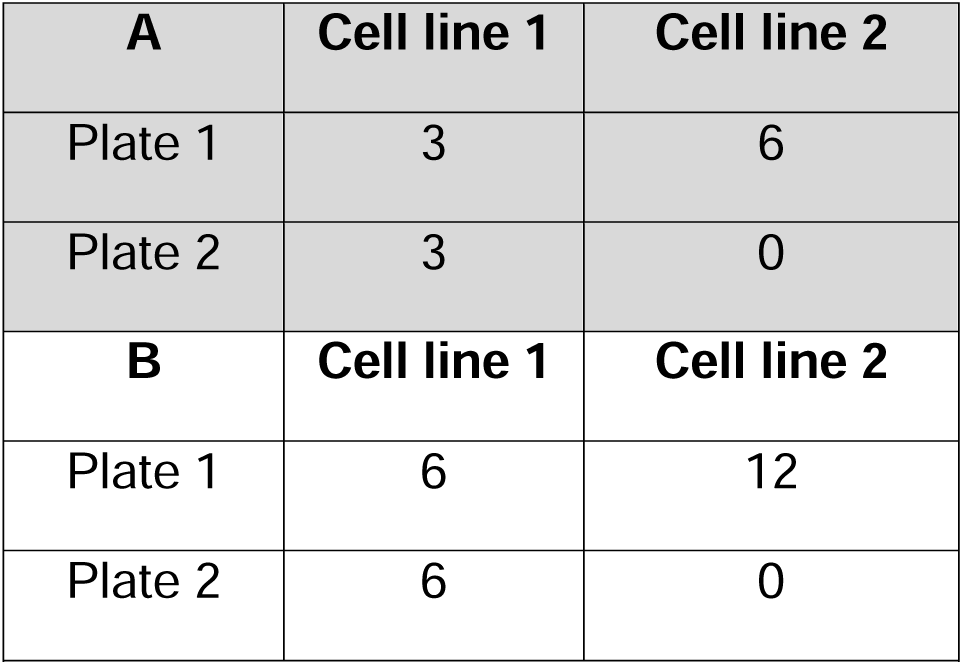
Example of plate-based cell count expansion simulation. (A) Original data structure and (B) changed data structure after simulation. The simulation is based on level=0 which results in increased number of cells per plate.

**Table S5.**
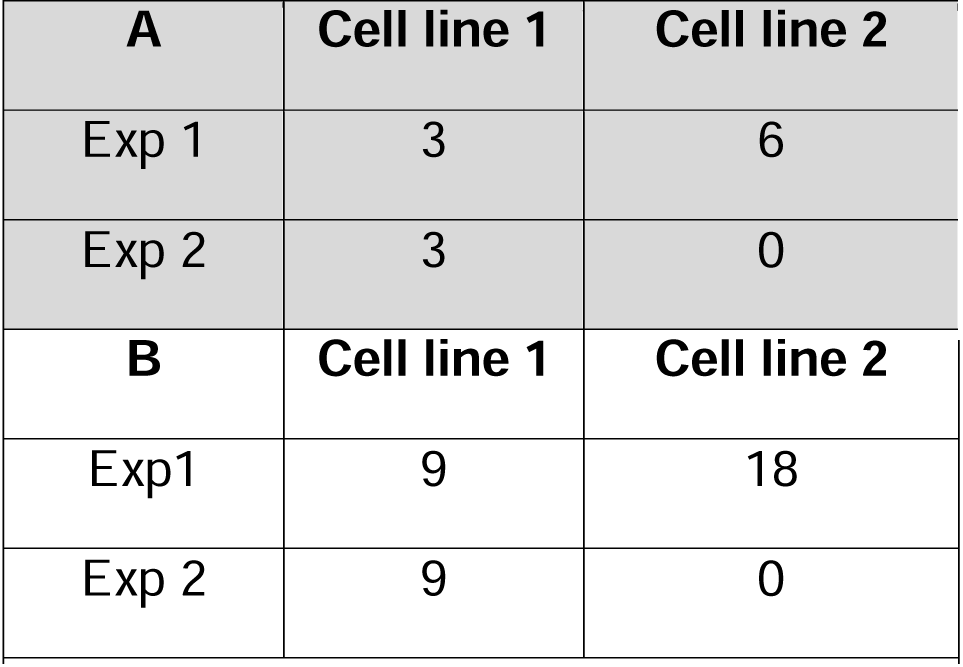
Example of cell line-based cell count expansion simulation. (A) Original data structure and (B) changed data structure after simulation. An example of a simulation in tabular format shows the change in the number of cells per cell line and experiment after simulation. The simulation is based on level=0 which results in an increased number of cells per cell line.

## References Cited

1. NIH. (2024) Enhancing reproducibility through rigor and transparency. Available from: https://grants.nih.gov/policy-and-compliance/policy-topics/reproducibility

2. NIH. (2024) Guidance: Rigor and reproducibility in grant applications. Available from: https://grants.nih.gov/policy-and-compliance/policy-topics/reproducibility/guidance

3. NIH. (2024) Training and other resources for rigor and reproducibility. Available from: https://grants.nih.gov/policy-and-compliance/policy-topics/reproducibility/training

4. Chan AW, Song F, Vickers A, Jefferson T, Dickersin K, Gøtzsche PC, Krumholz HM, Ghersi D, van der Worp HB. (2014) Increasing value and reducing waste: Addressing inaccessible research. Lancet 383:257–266. PMCID: PMC4533904

5. Young SS, Bang H, Oktay K. (2009) Cereal-induced gender selection? Most likely a multiple testing false positive. Proc. Biol. Sci. 276:1211–1222. PMCID: PMC2660953

6. Begley CG, Ioannidis JP. (2015) Reproducibility in science: Improving the standard for basic and preclinical research. Circ. Res. 116:116–126.

7. Dirnagl U, Duda GN, Grainger DW, Reinke P, Roubenoff R. (2022) Reproducibility, relevance and reliability as barriers to efficient and credible biomedical technology translation. Adv. Drug Deliv. Rev. 182:114118.

8. Peers IS, Ceuppens PR, Harbron C. (2012) In search of preclinical robustness. Nat. Rev. Drug. Discov. 11:733–734.

9. Macleod MR, Michie S, Roberts I, Dirnagl U, Chalmers I, Ioannidis JP, Al-Shahi Salman R, Chan AW, Glasziou P. (2014) Biomedical research: Increasing value, reducing waste. Lancet 383:101–104.

10. Belluz J. (2015) Most research spending is wasted on bad studies. These billionaires want to change that. Available from: https://www.vox.com/2015/10/4/9440931/arnold-foundation-meta-research.

11. Hurlbert SH. (1984) Pseudoreplication and the design of ecological field xxperiments. Ecol. Monogr. 54:187–211.

12. Haque A, Engel J, Teichmann SA, Lönnberg T. (2017) A practical guide to single-cell RNA-sequencing for biomedical research and clinical applications. Genome Med. 9:75. PMCID: PMC5561556

13. Bressan E, Reed X, Bansal V, Hutchins E, et al. (2023) The foundational data initiative for Parkinson disease: Enabling efficient translation from genetic maps to mechanism. Cell. Genom. 3:100261. PMCID: PMC10025424

14. McKinnon KM. (2018) Flow cytometry: An overview. Curr. Protoc. Immunol. 120:5.1.1–5.1.11. PMCID: PMC5939936

15. HD iPSC Consortium. (2012) Induced pluripotent stem cells from patients with Huntington’s disease show CAG-repeat-expansion-associated phenotypes. Cell Stem Cell 11:264–278. PMCID: PMC3804072

16. HD iPSC Consortium. (2017) Developmental alterations in Huntington’s disease neural cells and pharmacological rescue in cells and mice. Nat. Neurosci. 20:648–660. PMCID: PMC5610046

17. Arrasate M, Finkbeiner S. (2005) Automated microscope system for determining factors that predict neuronal fate. Proc. Natl. Acad. Sci. U. S. A. 102:3840–3845. PMCID: PMC553329

18. Arrasate M, Finkbeiner S. (2012) Protein aggregates in Huntington’s disease. Exp. Neurol. 238:1–11. PMCID: PMC3909772

19. Arrasate M, Mitra S, Schweitzer ES, Segal MR, Finkbeiner S. (2004) Inclusion body formation reduces levels of mutant huntingtin and the risk of neuronal death. Nature 431:805–810. PMCID: PMC15483602

20. Miller J, Arrasate M, Brooks E, Libeu CP, et al. (2011) Identifying polyglutamine protein species *in situ* that best predict neurodegeneration. Nat. Chem. Biol. 7:925–934. PMCID: PMC3271120

21. Mitra S, Tsvetkov AS, Finkbeiner S. (2009b) Single neuron ubiquitin-proteasome dynamics accompanying inclusion body formation in Huntington disease. J. Biol. Chem. 284:4398–4403. PMCID: PMCID: PMC2640959

22. Tsvetkov AS, Arrasate M, Barmada S, Ando DM, Sharma P, Shaby BA, Finkbeiner S. (2013) Proteostasis of polyglutamine varies among neurons and predicts neurodegeneration. Nat. Chem. Biol. 9:586–592. PMCID: PMC3900497

23. Barmada SJ, Skibinski G, Korb E, Rao EJ, Wu JY, Finkbeiner S. (2010b) Cytoplasmic mislocalization of TDP-43 is toxic to neurons and enhanced by a mutation associated with familial amyotrophic lateral sclerosis. J. Neurosci. 30:639–649. PMCID: PMC2821110

24. Bilican B, Serio A, Barmada SJ, Nishimura AL, et al. (2012) Mutant induced pluripotent stem cell lines recapitulate aspects of TDP-43 proteinopathies and reveal cell-specific vulnerability. Proc. Natl. Acad. Sci. U. S. A. 109:5803–5808. PMCID: PMC3326463

25. Serio A, Bilican B, Barmada SJ, Ando DM, et al. (2013) Astrocyte pathology and the absence of non-cell autonomy in an induced pluripotent stem cell model of TDP-43 proteinopathy. Proc. Natl. Acad. Sci. U. S. A. 110:4697–4702. PMCID: PMC3607024

26. Jones EA, Gillespie AK, Yoon SY, Frank LM, Huang Y. (2019) Early hippocampal sharp-wave ripple deficits predict later learning and memory impairments in an Alzheimer’s disease mouse model. Cell Rep. 29:212–2133.e4. PMCID: PMC7437815

27. Martinez-Losa M, Tracy TE, Ma K, Verret L, et al. (2018) Nav1.1-Overexpressing interneuron transplants restore brain rhythms and cognition in a mouse model of Alzheimer’s disease. Neuron 98:75–89.e5. PMCID: PMC5886814

28. Squair JW, Gautier M, Kathe C, Anderson MA, et al. (2021) Confronting false discoveries in single-cell differential expression. Nat. Commun. 12:5692. PMCID: PMC8479118

29. Lazic SE. (2010) The problem of pseudoreplication in neuroscientific studies: Is it affecting your analysis? BMC Neurosci. 11:5. PMCID: PMC2817684

30. Vadillo MA, Konstantinidis E, Shanks DR. (2016) Underpowered samples, false negatives, and unconscious learning. Psychon. Bull. Rev. 23:87–102. PMCID: PMC4742512

31. Button KS, Ioannidis JP, Mokrysz C, Nosek BA, Flint J, Robinson ES, Munafò MR. (2013) Power failure: Why small sample size undermines the reliability of neuroscience. Nat. Rev. Neurol. 14:365–376.

32. Sayed N, Liu C, Wu JC. (2016) Translation of human-induced pluripotent stem cells: From clinical trial in a dish to precision medicine. J. Am. Coll. Cardiol. 67:2161–2176. PMCID: PMC5086255

33. Tiscornia G, Vivas EL, Izpisúa Belmonte JC. (2011) Diseases in a dish: Modeling human genetic disorders using induced pluripotent cells. Nat. Med. 17:1570–7156.

34. Xie YZ, Zhang RX. (2015) Neurodegenerative diseases in a dish: The promise of iPSC technology in disease modeling and therapeutic discovery. Neurol. Sci. 36:21–27. PMCID: PMC4282683

35. Burrows CK, Banovich NE, Pavlovic BJ, Patterson K, Gallego Romero I, Pritchard JK, Gilad Y. (2016) Genetic variation, not cell type of origin, underlies the majority of identifiable regulatory differences in iPSCs. PLoS Genet. 12:e1005793. PMCID: PMC4727884

36. DeBoever C, Li H, Jakubosky D, Benaglio P, et al. (2017) Large-scale profiling reveals the influence of genetic variation on gene expression in human induced pluripotent stem cells. Cell Stem Cell 20:533–546.e7. PMCID: PMC5444918

37. Kilpinen H, Goncalves A, Leha A, Afzal V, et al. (2017) Common genetic variation drives molecular heterogeneity in human iPSCs. Nature 546:370–375. PMCID: PMC5524171

38. Volpato V, Webber C. (2020) Addressing variability in iPSC-derived models of human disease: Guidelines to promote reproducibility. Dis. Model Mech. 13:PMCID: PMC6994963

39. Carcamo-Orive I, Hoffman GE, Cundiff P, Beckmann ND, et al. (2017) Analysis of transcriptional variability in a large human iPSC library reveals genetic and non-genetic determinants of heterogeneity. Cell Stem Cell 20:518–532.e9. PMCID: PMC5384872

40. Champely S, Ekstrom C, Dalgaard P, Gill J, Weibelzahl S, Anandkumar A, Ford C, Volcic R, De Rosario H. (2020) pwr: Basic functions for power analysis. https://cran.r-project.org/web/packages/pwr/.

41. Green P, MacLeod CJ. (2016) SIMR: An R package for power analysis of generalized linear mixed models by simulation. Methods Ecol. Evol. 7:493–498.

42. IBM. (2009) IBM SPSS software.

43. STATA. (2021) Stata 17 [software].

44. SAS. (2020) SAS/STAT version 15.2 [software].

45. R Core Team. (2024) R: A language and environment for statistical computing. R Foundation for Statistical Computing. Available from: https://www.R-project.org/

46. Bates D. (2003) Coverting package to S4. R News 3:6–8.

47. Bates D, Mächler M, Bolker B, Walker S. (2015) Fitting linear mixed-effects models using lme4. J. Stat. Softw. 67:1–48.

48. Kuznetsova A, Brockhoff PB, Christensen RHB. (2017) lmerTest package: Tests in linear mixed effects models. J. Stat. Softw. 82:1–26.

49. Nieuwenhuis R, te Grotenhuis M, Pelzer B. (2012) influence.ME: Tools for detecting influential data in mixed Eefects models. The R Journal 4:38–47.

50. Millard SP. (2013) EnvStats: An R Package for environmental statistics. Springer, New York

51. Hartig F, Lohse L, Melina de Souza M. (2022) DHARMa: Residual diagnostics for hierarchical (multi-level / mixed) regression models. Available from: https://CRAN.R-project.org/package=DHARMa

52. Green P, MacLeod C, Alday P. (2023) simr: Power analysis for generalised linear mixed models by simulation. Available from: https://cran.r-project.org/web/packages/simr/index.html

53. Wickham H, Averick M, Bryan J, Chang W, et al. (2019) Welcome to the tidyverse. J. Open Source Softw. 4:1686.

54. Kiernan JA, Hudson AJ. (1991) Changes in sizes of cortical and lower motor neurons in amyotrophic lateral sclerosis. Brain 114 (Pt 2):843–853.

55. Osking Z, Ayers JI, Hildebrandt R, Skruber K, Brown H, Ryu D, Eukovich AR, Golde TE, Borchelt DR, Read TA, Vitriol EA. (2019) ALS-linked SOD1 mutants enhance neurite outgrowth and branching in adult motor neurons. iScience 11:294–304. PMCID: PMC6327879

56. Rosner B. (1983) Percentage points for a generalized ESD many-outlier procedure. Technometrics 25:165–172.

57. Bickel PJ. (1965) On some robust estimates of location. Ann. Math. Stat. 36:847–858.

58. Kim B. (2015) Understanding diagnostic plots for linear regression analysis. UVA Library StatLab Available from: https://library.virginia.edu/data/articles/diagnostic-plots.

59. Cook RD. (1977) Detection of influential observation in linear regression. Technometrics 19:15–18.

60. Dukkipati SS, Garrett TL, Elbasiouny SM. (2018) The vulnerability of spinal motoneurons and soma size plasticity in a mouse model of amyotrophic lateral sclerosis. J. Physiol. 596:1723–1745. PMCID: PMC5924829

61. Shoenfeld L, Westenbroek RE, Fisher E, Quinlan KA, Tysseling VM, Powers RK, Heckman CJ, Binder MD. (2014) Soma size and Cav1.3 channel expression in vulnerable and resistant motoneuron populations of the SOD1G93A mouse model of ALS. Physiol. Rep. 2:e12113. PMCID: PMC4246589

62. Ye L, Dittlau KS, Sicart A, Janky R, Van Damme P, Van Den Bosch L. (2025) Sporadic ALS hiPSC-derived motor neurons show axonal defects linked to altered axon guidance pathways. Neurobiol. Dis. 206:106815.

63. Bye CR, Qian E, Lim K, Daniszewski M, et al. (2026) Large-scale drug screening in iPSC-derived motor neurons from sporadic ALS patients identifies a potential combinatorial therapy. Nat. Neurosci. 29:40–52. PMCID: PMC12779551

64. Fujimori K, Ishikawa M, Otomo A, Atsuta N, Nakamura R, Akiyama T, Hadano S, Aoki M, Saya H, Sobue G, Okano H. (2018) Modeling sporadic ALS in iPSC-derived motor neurons identifies a potential therapeutic agent. Nat. Med. 24:1579–1589.

65. Linsley JW, Shah K, Castello N, Chan M, et al. (2021a) Genetically encoded cell-death indicators (GEDI) to detect an early irreversible commitment to neurodegeneration. Nat. Commun. 12:5284. PMCID: PMC8421388

66. Mitra S, Tsvetkov AS, Finkbeiner S. (2009a) Protein turnover and inclusion body formation. Autophagy 5:1037–1038. PMCID: PMC2892253

67. Shaby BA, Skibinsk iG, Ando M, LaDow ES, Finkbeiner S. (2016) A three-groups model for high-throughput survival screens. Biometrics 72:936–944. PMCID: PMC4965338

68. Baxi EG, Thompson T, Li J, Kaye JA, et al. (2022) Answer ALS, a large-scale resource for sporadic and familial ALS combining clinical and multi-omics data from induced pluripotent cell lines. Nat. Neurosci. 25:226–237. PMCID: PMC8825283

69. Wang B, Vartak R, Zaltsman Y, Naing ZZC, et al. (2024) A foundational atlas of autism protein interactions reveals molecular convergence. bioRxiv 2023.12.03.569805 [Preprint]. February 4, 2024. Available from: 10.1101/2023.12.03.569805. PMCID: PMC10705567

70. Kaye J, Amirani N, Chan Ú, Al Bistami N, et al. (2026) Predictive cellular signatures from live human motor neurons distinguish TDP-43 ALS and enable ALS subtype stratification. bioRxiv 2026.04.22.719920 [Preprint]. April 24, 2026. Available from: 10.64898/2026.04.22.719920.

71. Kutzing MK, Langhammer CG, Luo V, Lakdawala H, Firestein BL. (2010) Automated Sholl analysis of digitized neuronal morphology at multiple scales. J. Vis. Exp. 2354. PMCID: PMC3159598

72. Stocksdale JT, Leventhal MJ, Lam S, Xu YX, et al. (2025) Intersecting impact of CAG repeat and huntingtin knockout in stem cell-derived cortical neurons. Neurobiol. Dis. 210:106914.

73. Koutsodendris N, Blumenfeld J, Agrawal A, Traglia M, et al. (2023) Neuronal APOE4 removal protects against tau-mediated gliosis, neurodegeneration and myelin deficits. Nat. Aging 3:275–296. PMCID: PMC10154214

74. Possin KL, Sanchez PE, Anderson-Bergman C, Fernandez R, et al. (2016) Cross-species translation of the Morris maze for Alzheimer’s disease. J. Clin. Invest. 126:779–783. PMCID: PMC4731157

75. Chang CW, Evans MD, Yu X, Yu GQ, Mucke L. (2021) Tau reduction affects excitatory and inhibitory neurons differently, reduces excitation/inhibition ratios, and counteracts network hypersynchrony. Cell Rep. 37:109855. PMCID: PMC8648275

76. Zimmerman KD, Espeland MA, Langefeld CD. (2021) A practical solution to pseudoreplication bias in single-cell studies. Nat. Commun. 12:738. PMCID: PMC7854630

77. Crowell HL, Soneson C, Germain PL, Calini D, Collin L, Raposo C, Malhotra D, Robinson MD. (2020) Muscat detects subpopulation-specific state transitions from multi-sample multi-condition single-cell transcriptomics data. Nat. Commun. 11:6077. PMCID: PMC7705760

78. Box GEP, Cox DR. (2018) An analysis of transformations. J. R. Stat. Soc. Ser. B Stat. Method. 26:211–243.

79. Yeo I-K, Johnson RA. (2000) A new family of power transformations to improve normality or symmetry. Biometrika 87:954–959.

80. Dunn PK, Smyth GK. (1996) Randomized quantile residuals. J. Comput. Graph. Stat. 5:236–244.

81. Answer ALS. (2024) Neuromine. Available from: https://dataportal.answerals.org/home

82. Mitra S, Tsvetkov AS, Finkbeiner S. (2009) Single neuron ubiquitin-proteasome dynamics accompanying inclusion body formation in Huntington disease. J. Biol. Chem. 284:4398– 4403. PMCID: PMC2640959

83. Aikio M, Odeh HM, Wobst HJ, Lee BL, et al. (2025) Opposing roles of p38α-mediated phosphorylation and PRMT1-mediated arginine methylation in driving TDP-43 proteinopathy. Cell Rep. 44:115205. PMCID: PMC11831926

84. Jalili V, Afgan E, Gu Q, Clements D, Blankenberg D, Goecks J, Taylor J, Nekrutenko A. (2020) The Galaxy platform for accessible, reproducible and collaborative biomedical analyses: 2020 update. Nucleic Acids Res. 48:W395–W402. PMCID: PMC7319590

85. Schindelin J, Arganda-Carreras I, Frise E, Kaynig V, et al. (2012) Fiji: An open-source platform for biological-image analysis. Nat. Methods 9:676–682. PMCID: PMC3855844

86. Zhou C, Wang G, Huang H, Song L, Xue K. (2019) Edge detection based on joint iteration ghost imaging. Opt. Express 27:27295–27307.

87. Python. (2024) Python [software]. Available from: https://www.python.org/

88. Bird AD, Cuntz H. (2019) Dissecting sholl analysis into its functional components. Cell Rep. 27:3081–3096.e5.

89. Koch E, Rosolowsky EW. (2025) FilFinder. Available from: https://github.com/e-koch/FilFinder?tab=readme-ov-file

