## Supplementary Note for "RMeDPower2 for Biology: guiding the design, experimental structure and analyses of experiments generating repeated measures datasets"

### *RMeDPower2*: helping design and analyses of data from repeated measures experiments

true true

**Abstract**

Biomedical research very often involves data generated from repeated measures experiments. RMeDPower2 is an R package that provides complete functionality to analyse data coming from repeated measures experiments, i.e., where one has repeated measures from the same biological/independent units or samples. RMeDPower2 helps test the modeling assumptions one makes, identify outlier observations, outlier units at different levels of the design, estimates statistical power or perform sample size calculations, estimate parameters of interest and also to visualize the association being tested. The functionality is limited to testing associations of one predictor (continuous or categorical, e.g., disease status or brain pathology) along with one another covariate (e.g., gender status) in the context of hierarchical or crossed experimental designs. This vignette illustrates the use of this package in multiple contexts relevant to biomedical research.

RMeDPower2 defines the experimental design for the data, probability model of the data generating distribution and necessary parameters required for sample size calculation using convenient S4 class objects. It uses these objects in one framework that brings together the functionality implemented in multiple R packages - lme4 (implementation of linear mixed effects models), influence.ME (identification of outlier units), EnvStats (identification of outlier observations), DHARMa (testing of modeling assumptions for non-normal distributions), simr (sample-size calculations) and tidyverse (data manipulation and visualization).

### Contents

**1 Getting started** **2**

1.1 Installing `RMeDPower2` 2
1.2 Quick tutorial 2
1.3 Input data 3
1.4 Three key classes of `RMeDPower2` 3
1.5 Three key functions of `RMeDPower2` 7

**2 Application examples** **8**

2.1 Plate assays 8
2.2 Single-cell RNA-seq assays 32
2.3 Behavior assays 68
2.4 Brain electrical signal assays 86

**3 Understanding simulations to estimate the power** **126**

**References** **139**

**4 Session info** **139**

### 1 Getting started

#### 1.1 Installing RMeDPower2

RMeDPower2 can be installed using the following commands

```
install.packages("devtools")
library(devtools)
install_github('gladstone-institutes/RMeDPower2', build_vignettes=TRUE, force = TRUE)
```

#### 1.2 Quick tutorial

The workflow can be accomplished in 5 main steps as described in the sections below (1.2.1-.2.5).

##### 1.2.1 Load libraries and read input data

The first step is to load RMeDPower2 and read-in the input data as a data frame. This is accomplished by the following commands:

```
suppressPackageStartupMessages(library(RMeDPower2))
options(warn = -1)
data <- plate_assay_pilot_data
```

Example data has experiment, line as experience columns, classification as the condition column, and cell size as the response column.

```
knitr::kable(data[1:5,] )
```

| experiment | line | classification | cell_size1 | cell_size2 | cell_size3 | cell_size4 | cell_size5 | cell_size6 |
|---|---|---|---|---|---|---|---|---|
| experiment1 | cellline1 | 0 | 353.8401 | 353.8401 | 353.8401 | 353.8401 | 353.8401 | 353.8401 |
| experiment1 | cellline1 | 0 | 456.3522 | 456.3522 | 456.3522 | 456.3522 | 456.3522 | 456.3522 |
| experiment1 | cellline1 | 0 | 350.7909 | 350.7909 | 350.7909 | 350.7909 | 350.7909 | 350.7909 |
| experiment1 | cellline1 | 0 | 387.4861 | 387.4861 | 387.4861 | 387.4861 | 387.4861 | 387.4861 |
| experiment1 | cellline1 | 0 | 403.9912 | 403.9912 | 403.9912 | 403.9912 | 403.9912 | 403.9912 |

##### 1.2.2 Specify design, model and power parameters

The following set of commands allows users to specify parameters for experimental design, assumed probability models and distributions, and power simulations.

1. Use a text editor to modify the template jsonfile capturing the design for the data and read in this file.

```
design <- readDesign(file.path(template_dir,"design_cell_assay.json"))
```

2. Use a text editor to modify the template jsonfile capturing the error probability model and read in this file.

```
model <- readProbabilityModel(file.path(template_dir,"prob_model.json"))
```

3. Use a text editor to modify the template jsonfile capturing the power parameters and read in this file.

```
power_param <- readPowerParams(file.path(template_dir,"power_param.json"))
```

##### 1.2.3 Diagnose data and model

The next set of commands will allow the user to evaluate the datatype, examine if the model is appropriate and identity outliers.

1. To diagnose the data and model, test model assumptions, identify outlier observations, outlier biological/independent units.

```
diagnose_res <- diagnoseDataModel(data, design, model)
data_update = diagnose_res$Data_updated
```

2. Modify design objects (e.g., remove outliers), model objects (e.g., change probability distribution assumption) if needed based on output of step 1 above.

```
diagnose_res_update <- diagnoseDataModel(data_update, design_update, model_update)
```

3. Repeat steps 1 and 2 until the models assumptions are met. For example, users can choose to remove outlier-experiments so that the distribution is close to normal.

###### 1.2.4 Statistical power estimation

This section of code will allow the user to run sample-size calculations from the pilot input data.

```
power_res <- calculatePower(data, design, model, power_param)
```

###### 1.2.5 Estimate parameters of interest

Once we have run the models, the user can estimate and visualize parameters of interest in order to investigate the association between response variable and condition variables.

```
estimate_res <- getEstimatesOfInterest(data, design, model, print_plots = F)
```

##### 1.3 Input data

Users will have to ensure that the input data is organized in a format that will allow RMedPower2 to read in the data. The data should be structured in a table with rows representing the individual observations or responses and columns representing not only the corresponding explanatory variables but also other variables that capture the hierarchical or crossed design of how the data was generated. It is important that the columns have names or headers - these column names will be used to define the relevant S4 class objects.

For example, the next below illustrates the input data from a cellular assay where each of the 2,588 observations represents a measurement of size (`cell_size2`) of a single cell, from a given cell line (`line`) that is either from a control or disease condition (`classification`) and is measured in a given experimental batch (`experiment`).

```
## 'data.frame':    2588 obs. of  4 variables:
##  $ experiment    : Factor w/ 3 levels "experiment1",..: 1 1 1 1 1 1 1 1 1 3 ...
##  $ line          : Factor w/ 10 levels "cellline1","cellline10",..: 1 1 1 1 1 1 1 1 1 1 ...
##  $ classification: Factor w/ 2 levels "0","1": 1 1 1 1 1 1 1 1 1 1 ...
##  $ cell_size2    : num  354 456 351 387 404 ...
```

##### 1.4 Three key classes of `RMeDPower2`

Here, users will learn how to set up parameters for experimental designs, probability and distributional assumptions, and power simulations.

###### 1.4.1 `RMeDesign` S4 class

The underlying design for the data will be stored in an object (R data structure with variables) of this class. The slots of this class are:

1. `response_column`: character that is the column name in the input data table that captures the responses or individual observations. This has to be set by the user.

2. `covariate`: character that is the column name in the input data table that captures the covariate information. This will be used as a fixed effect in the mixed effects model of the data. This slot is set to `NULL` if there are no covariates.
3. `condition_column`: character that is the column name in the input data table that captures the predictor/independent variable information. This will be used as a fixed effect in the mixed effects model of the data.This has to be set by the user.
4. `experimental_columns`: character vector that represents the column names in the input data table that captures the experimental variables of the design. These will be used as random effects in the mixed effects model of the data. The order of the names in this character vector is important. The first element captures the top hierarchical level of the design, the second element the next level of the hierarchy and so on. The hierarchy is explicitly modeled in the specification of the mixed effects models. The hierarchy will not be modeled for variables specified in the `crossed_columns` slot (see below). This has to be set by the user.
5. `condition_is_categorical`: logical value equal to TRUE if the variable in the `condition_column` is to be considered as a categorical variable and is equal to FALSE otherwise. Defaults to TRUE.
6. `crossed_columns`: character vector that represents the column names in the input data table that are a subset of the values in `experimental_columns` representing crossed factors. Defaults to `NULL`.
7. `total_column`: character that is the column name in the input table that will be used to offset values in the mixed effects models. Useful for modeling count data. Defaults to `NULL`.
8. `outlier_alpha`: numeric value that is the p-value threshold used to identify outlier observations. Defaults to 0.05.
9. `na_action`: character equal to *complete* or *unique*. To be used in the context where the input data has multiple responses that will each be sequentially modeled. For example, from a single set of data, a user is gathering numerous observations. Here we refer to *complete* in the scenario where the observed outliers that identified for one response will also be identified as outliers for all the other responses while *unique* refers to the situation where the outlier observations identified for one response will only be used for the model of that particular response. Defaults to *complete*.
10. `include_interaction`: Whether to include condition * covariate interaction
11. `random_slope_variable`: Variable for random slopes (typically "condition_column")
12. `covariate_is_categorical`: Specify whether the covariate variable is categorical. TRUE: Categorical, FALSE: Continuous.

To create an object of class RMeDesign, the following code is used:

```
design <- new("RMeDesign")
print(design)
```

```
## An object of class "RMeDesign"
## Slot "response_column":
## [1] "response_column"
##
## Slot "covariate":
## NULL
##
## Slot "condition_column":
## [1] "condition_column"
##
## Slot "condition_is_categorical":
## [1] TRUE
##
```

```
## Slot "experimental_columns":
## [1] "experimental_column"
##
## Slot "crossed_columns":
## NULL
##
## Slot "total_column":
## NULL
##
## Slot "outlier_alpha":
## [1] 0.05
##
## Slot "na_action":
## [1] "complete"
##
## Slot "include_interaction":
## [1] NA
##
## Slot "random_slope_variable":
## NULL
##
## Slot "covariate_is_categorical":
## [1] NA
```

In practice, for a given data set the design can be input from a jsonfile. The user can use a text editor to modify the `design_template.json` file available with the package to input the relevant information based on the column names of the input data. For example, the design for the cell size data referred to above can be read using the function `readDesign`.

```r
design <- readDesign(file.path(template_dir, "design_cell_assay.json"))
print(design)
```

```
## An object of class "RMeDesign"
## Slot "response_column":
## [1] "cell_size2"
##
## Slot "covariate":
## NULL
##
## Slot "condition_column":
## [1] "classification"
##
## Slot "condition_is_categorical":
## [1] TRUE
##
## Slot "experimental_columns":
## [1] "experiment" "line"
##
## Slot "crossed_columns":
## [1] "line"
##
## Slot "total_column":
## NULL
##
## Slot "outlier_alpha":
```

```
## [1] 0.05
##
## Slot "na_action":
## [1] "complete"
##
## Slot "include_interaction":
## [1] NA
##
## Slot "random_slope_variable":
## NULL
##
## Slot "covariate_is_categorical":
## [1] NA
```

##### 1.4.2 `ProbabilityModel` S4 class

The error probability distribution is specified by two slots in objects of the `ProbabilityModel` class.

1. `error_is_non_normal`: Is the logical value that is TRUE if the underlying probability distribution is not assumed to be Normal and is FALSE otherwise.

2. `family_p`: This character value specifies the probability distribution. Accepted values are *poisson*(`=poisson(link="log")`), *binomial*(`=binomial(link="logit")`), *bionomial_log*(`=binomial(link="log")`), *Gamma_log*(`=Gamma(link="log")`), *Gamma*(`=Gamma(link="inverse")`), *negative_binomial*. Defaults to `NULL`.

```r
model <- readProbabilityModel(file.path(template_dir, "prob_model.json"))
print(model)
```

```
## An object of class "ProbabilityModel"
## Slot "error_is_non_normal":
## [1] FALSE
##
## Slot "family_p":
## NULL
```

##### 1.4.3 `PowerParams` S4 class

The parameters described below are necessary for sample-size estimation and are stored in the following slots of the `PowerParams` class.

1. `target_columns`: This describes the character vector with column names of experimental variables for which the sample-size estimation is to be performed
2. `power_curve`: This is numeric 1 or 0. 1: Power simulation over a range of sample sizes or levels. 0: Power calculation over a single sample size or a level. Defaults to 1.
3. `nsimn`: The number of simulations to estimate power. Defaults to 10.
4. `levels`: numeric 1 or 0. 1: Amplify the number of corresponding target parameter. 0: Amplify the number of samples from the corresponding target parameter, ex) If target_columns = c("experiment","cell_line") and if you want to expand the number of experiment and sample more cells from each cell line, set levels = c(1,0).
5. max_size: Maximum levels or sample sizes to test. Default if set to `NULL` equals the current level or the current sample size x 5. ex) If max_levels = c(10,5), it will test upto 10 experiments and 5 cell lines.
6. `alpha`: Type I error for sample-size estimation. Defaults to 0.05.
7. `breaks`: Levels /sample sizes of the variable to be specified along the power curve. Default if set to NULL equals max (1, round (the number of current levels / 5)).

8. `effect_size`: A 3 dimensional numeric vector given the proposed effect sizes/parameter estimates for the condition_column, covariate and interaction term between these two variables, respectively in that order. Depending on the model not all elements of the effect_size would make sense. For example, in situations, where there is only the condition column, the second and third elements of effect_size should be equal to NA. If you know the effect size of your condition variable, the effect size can be provided as a parameter. Default set to `NULL`,i.e., if the effect size is not provided then it will be estimated from your data.
9. `icc`: Intra-class coefficients for each of the experimental variables. Used only in the situation when `error_is_non_normal=FALSE` and when the data does not allow variance component estimation.

```
power_param <- readPowerParams(file.path(template_dir, "power_param.json"))
print(power_param)
```

```
## An object of class "PowerParams"
## Slot "target_columns":
## [1] "experiment"
##
## Slot "power_curve":
## [1] 1
##
## Slot "nsimn":
## [1] 10
##
## Slot "levels":
## [1] 1
##
## Slot "max_size":
## [1] 1
##
## Slot "alpha":
## [1] 0.05
##
## Slot "breaks":
## NULL
##
## Slot "effect_size":
## NULL
##
## Slot "icc":
## NULL
```

#### 1.5 Three key functions of `RMeDPower2`

The following are functions for users to manipulate data, estimate power, and perform association analysis.

##### 1.5.1 `diagnoseDataModel(data, design, model)` function

This set of code, tests the model assumptions by using quantile-quantile, residuals vs fitted and residuals versus predicted plots. If needed, the code also transforms the responses if feasible and identifies outlier observations and also outlier experimental units.

##### 1.5.2 `calculatePower(data, design, model, power_param)` function

This set of code performs sample-size calculations for given experimental design/target variable.

###### 1.5.3 getEstimatesOfInterest(data, design, model) function

This string estimates parameter of interest and also plots the resulting association with predictor of interest using residuals from the model that removes the effects of the modeled experimental variables.

An overview of the anticipated usage of the functionality of this package is shown in the figure below.

### 2 Application examples

Here we illustrate the application of RMeDPower2 using data derived from a range of domains in biomedical research.

#### 2.1 Plate assays

iPSC lines from control and patient derived iPSCs from sALS patients were obtained through the Answer ALS (AALS) consortium36 . AALS is the largest repository of ALS patient samples, and contains publicly available patient-specific iPSCs and multi-OMIC data from motor neurons derived from those iPSCs—including RNA Seq, proteomics and whole genome sequence data— all of which is matched with longitudinal clinical data (https://dataportal.answerals.org/home)36.

We used 6 control lines and 4 sALS lines and differentiated each line into motor neurons (iMNs). iMNs were transduced with a morphology marker and imaged on day ~25 using RM37-45 as previously described36. Images were processed and the soma size of neurons were captured using contour ellipse46 function adapted for use in python (https://www.python.org/).

The next step is to read the data and perform data distribution diagnostics, association analysis, and power simulation.

First, we will read the data from the package and json files with specified parameters:

```
template_dir <- system.file("input_templates/cell_assay_data", package = "RMeDPower2")
data <- plate_assay_pilot_data
design <- readDesign(file.path(template_dir,"design_cell_assay.json"))
model <- readProbabilityModel(file.path(template_dir,"prob_model.json"))
power_param <- readPowerParams(file.path(template_dir,"power_param.json"))
```

A visualization of the distribution of cell_size2 across experiments and cell-lines as box-plots and also in terms of empirical cumulative distribution plots suggests the observations have an extremely long tail. We will therefore filter out the top 5 percent of the observations to ensure the results are not skewed by these extreme observations. We also see that the observations in experiment 3 are in general lower than those in the other two ex-

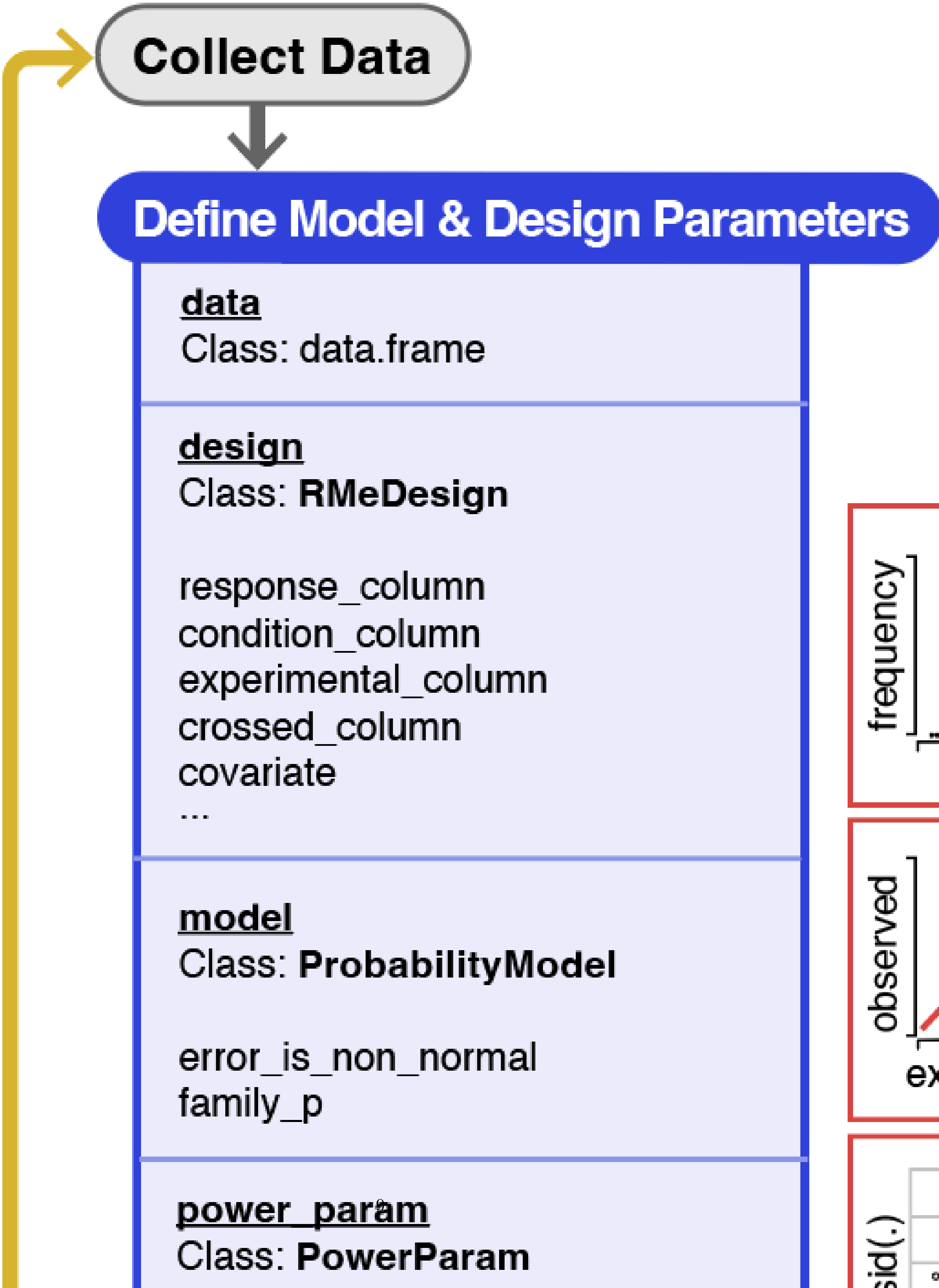

Collect Data
Define Model & Design Parameters
data
Class: data.frame
design
Class: RMeDesign
response_column
condition_column
experimental_column
crossed_column
covariate
...
model
Class: ProbabilityModel
error_is_non_normal
family_p
power_param
Class: PowerParam
frequency
observed

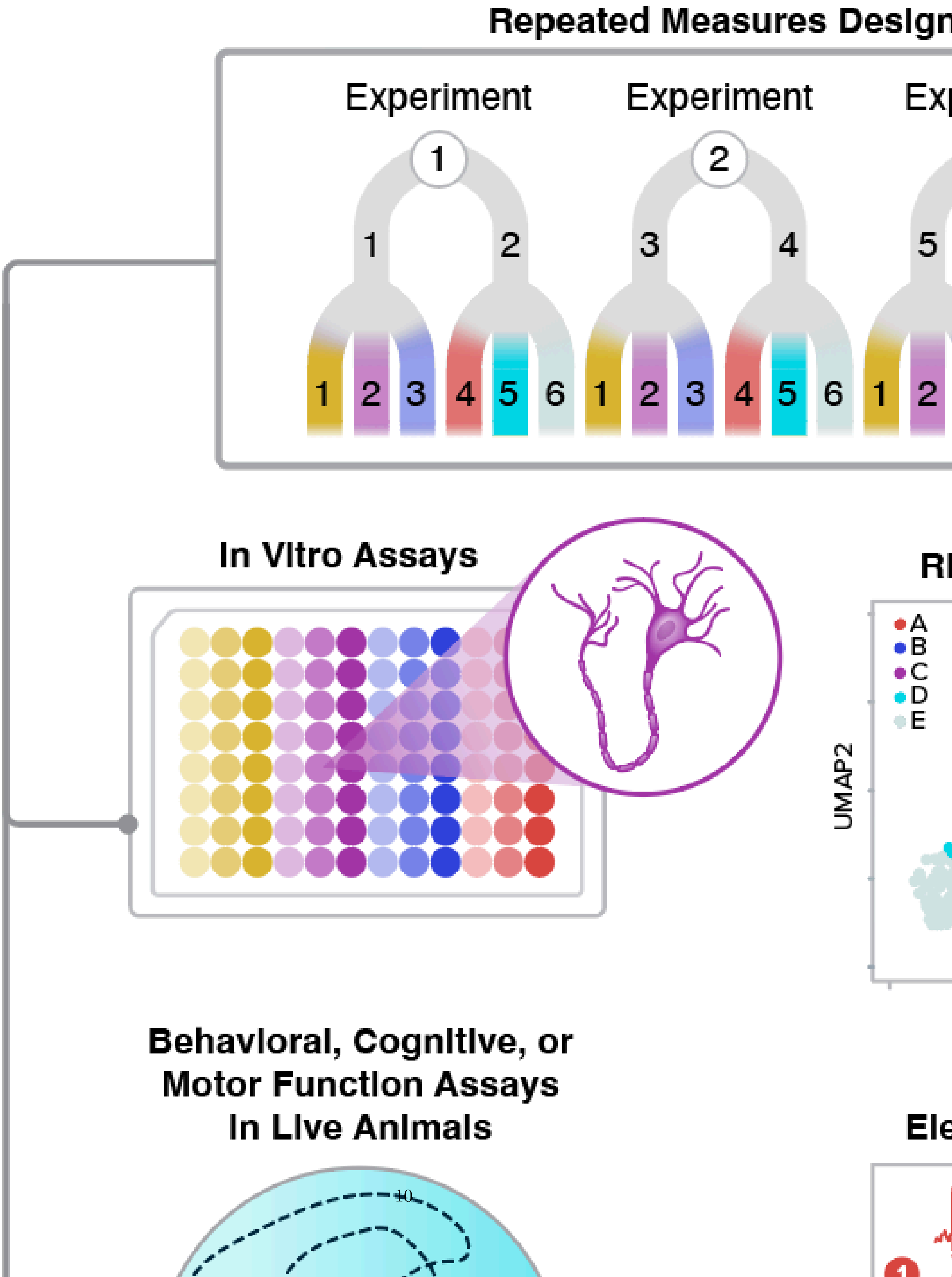
Repeated Measures Design
Experiment
Experiment
1
2
1
2
3
4
5
1
2
3
4
5
6
1
2
3
4
5
6
1
2
In Vitro Assays
A
B
C
D
E
UMAP2
Behavioral, Cognitive, or
Motor Function Assays
In Live Animals

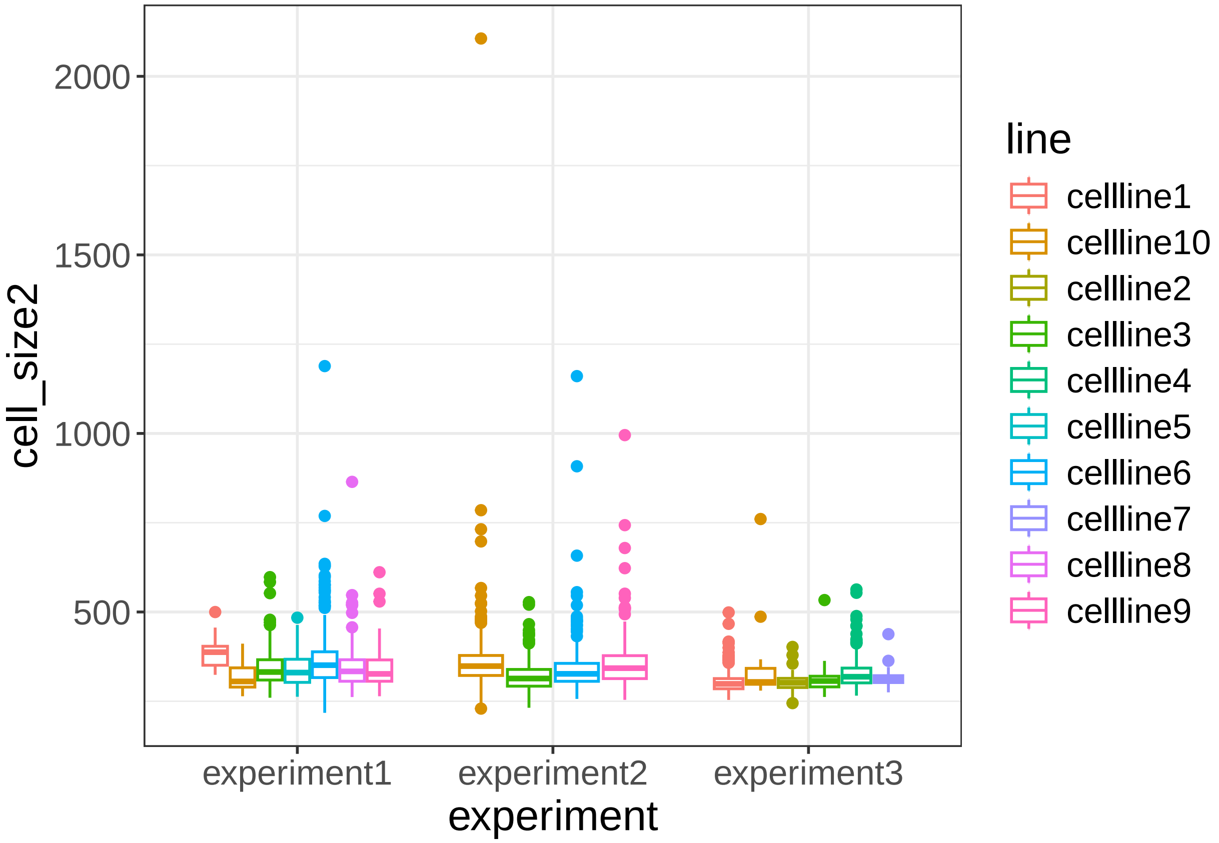

cell_size2
2000
1500
1000
500
experiment1
experiment2
experiment3
experiment
line
cellline1
cellline10
cellline2
cellline3
cellline4
cellline5
cellline6
cellline7
cellline8
cellline9


periments.

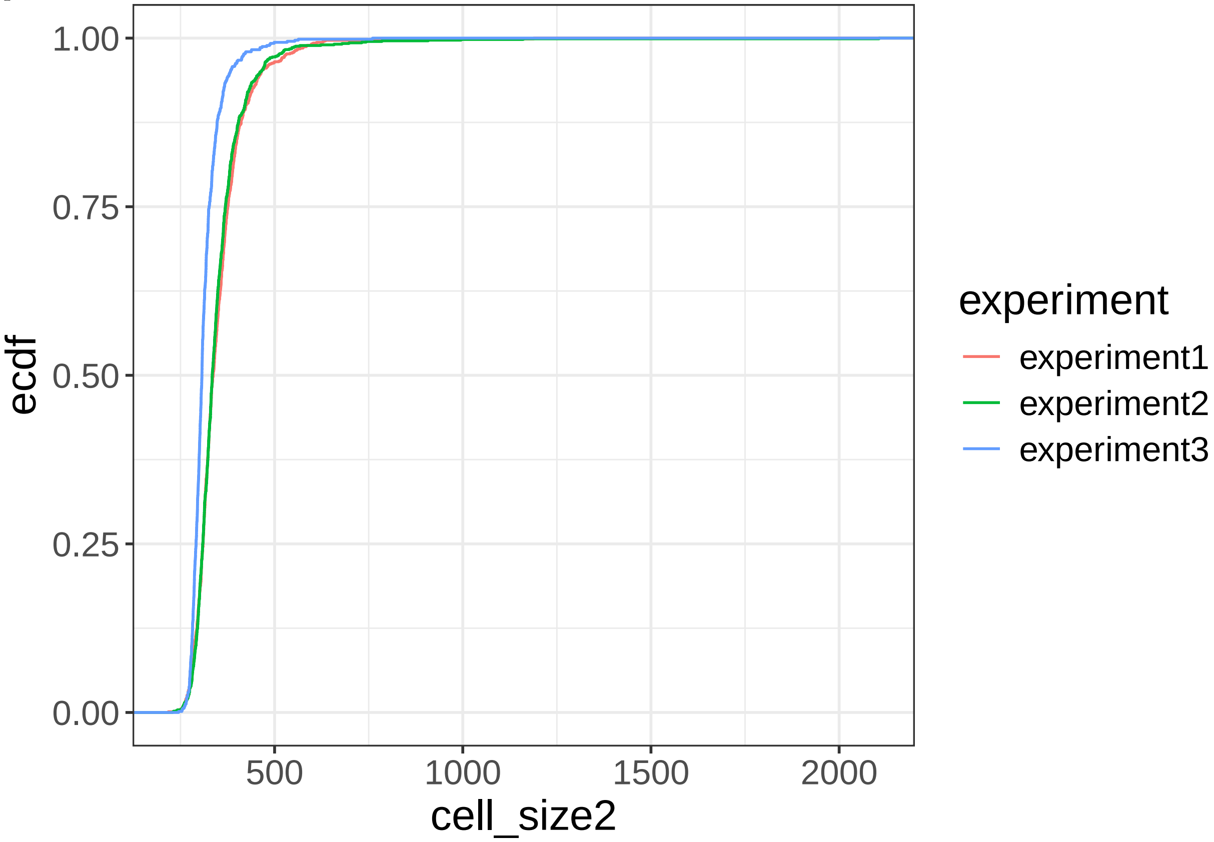

ecdf
1.00
0.75
0.50
0.25
0.00
500
1000
1500
2000
cell_size2
experiment
experiment1
experiment2
experiment3

```r
data %<>% filter(cell_size2 < quantile(cell_size2, 0.95))
```

We would like to estimate the difference in the mean cell-sizes across all the observations derived from all experimental batches between the ALS disease lines and the control cell lines (modeled by the *classification* variable). The first step is run diagnosis on the data and model using the 'diagnoseDataModel' function. The Q-Q plot of the residuals of the LMM model fit to the log-transformed cell-sizes suggests a better fit than the one fit to the cell sizes in the natural scale , assuming that the data generating distribution is based on the Normal distribution. Therefore, we will hereafter model the log-transformed cell sizes. The reasonably horizontal best fit lines for the residuals from this versus the fitted values, and the square-root of the absolute values of the residuals versus the fitted values, Figure 7e suggest both the LMM model assumptions of linearity and homoscedasticity are reasonable. There were no identified individual outlier observations based on the application of the Rosner's test to the distribution of the residuals in.

```r
diagnose_res <- diagnoseDataModel(data, design, model)
```

The *diagnoseDataModel* function fits at least one model representing one in the natural scale of the response of interest. Additionally, if individual observations are inferred as outliers using the rosner's test applied to the residuals of the fitted model then another model is fit without these outlier observations. In addition, if the assumed probability distribution of the residuals of the model is normal then a further additional model are fit using logarithm (log) transformed responses and another one if there are outliers in the log-transformed model fit.

To identify all the models fit to the data

```r
print(diagnose_res$models)
```

```
## [1] "Natural scale" "Log scale"
```

We see that two models were fit to this data - one in the natural scale of cell size and the other in the logarithmic scale. No individual observations were identified as outliers in each of these models.

Let us look the basic QC plots for the models fit in the natural scale of the responses.

```r
plot_no <- 1
plots <- diagnose_res$diagnostic_plots$natural_scale$plots
captions <- diagnose_res$diagnostic_plots$natural_scale$captions
for (i in seq_along(plots)) {
  print(plots[[i]])
  cat(sprintf("\n\n\n**Figure %d:** %s. %s\n\n", plot_no,names(plots)[i], captions[i]))
  plot_no <- plot_no + 1
}
```

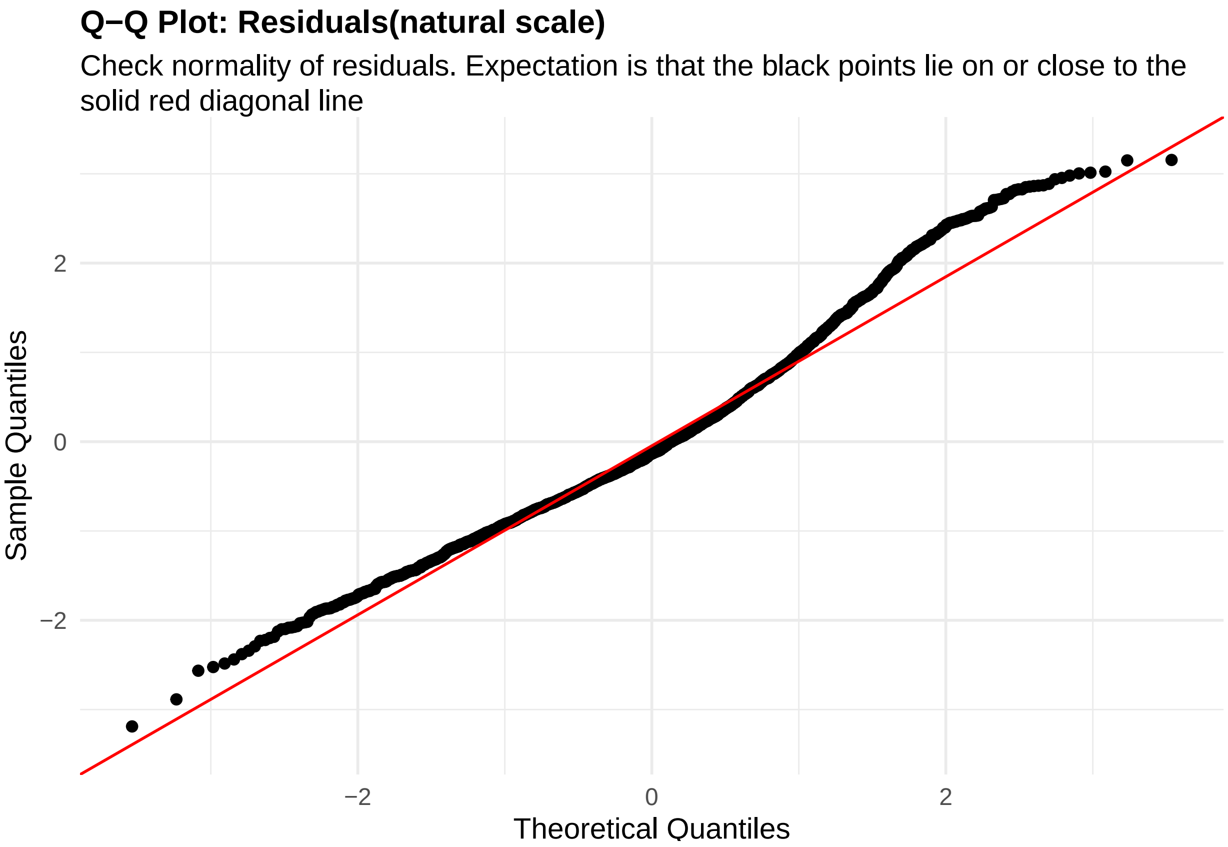


**Figure 1:** residuals_QQ. Check normality of residuals (natural scale). Expectation is that the black points lie on or close to the solid red diagonal line. Check the Normal Q-Q plots for good and bad examples here: https://library.virginia.edu/data/articles/diagnostic-plots

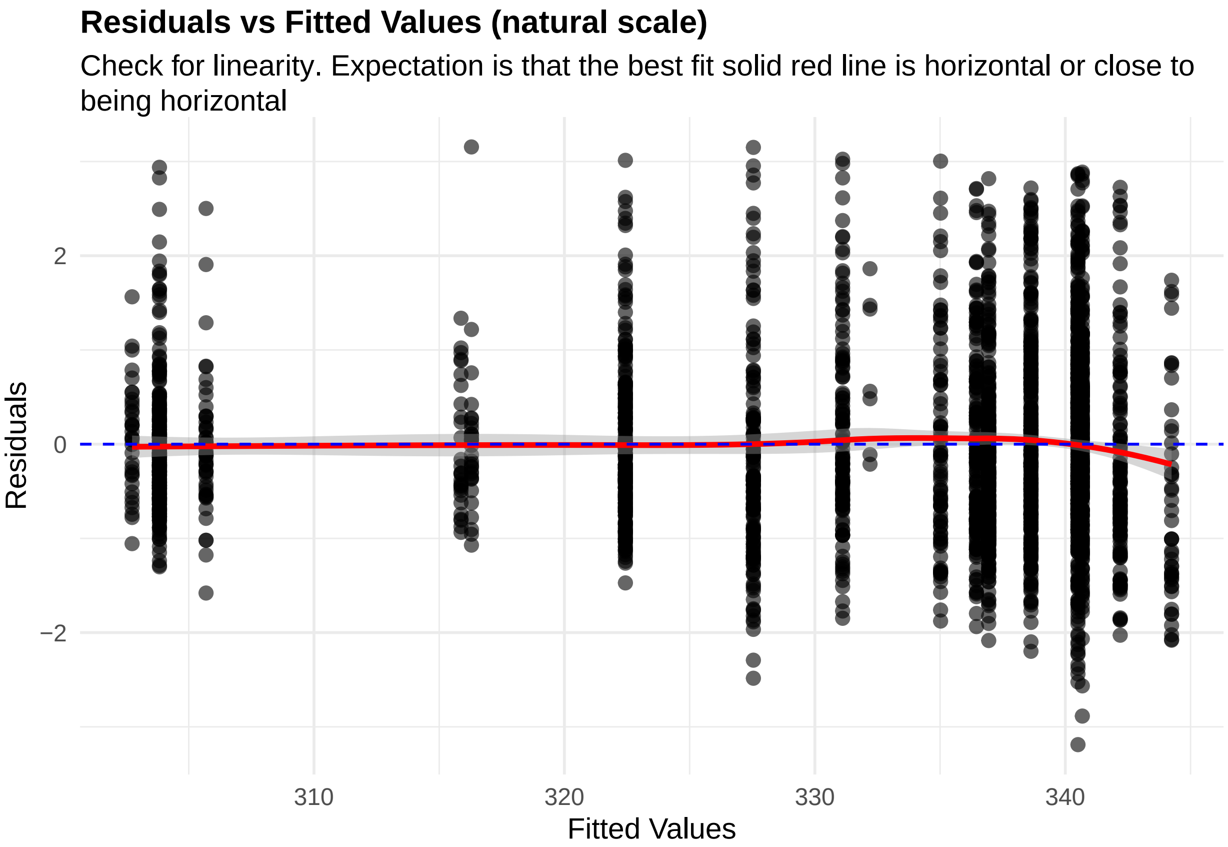


**Figure 2:** residuals_vs_fitted. Check for linearity (natural scale). Expectation is that the best fit solid red line is horizontal or close to being horizontal. Check the Residuals vs Fitted plots for good and bad examples here: https://library.virginia.edu/data/articles/diagnostic-plots

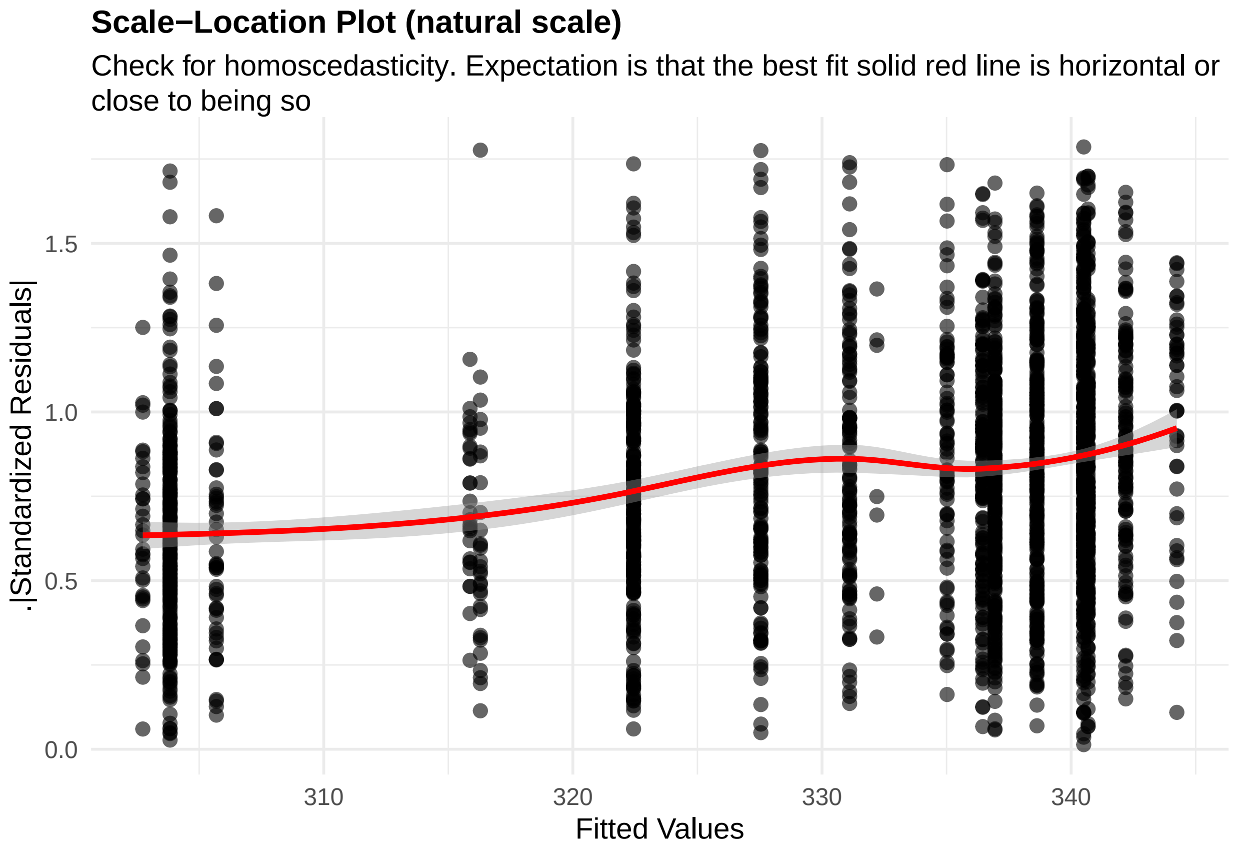


**Figure 3:** residuals_homoscedasticity. Check for homoscedasticity (natural scale). Expectation is that the best fit solid red line is horizontal or close to being so. Check the Scale-Location plots for good and bad examples here: https://library.virginia.edu/data/articles/diagnostic-plots

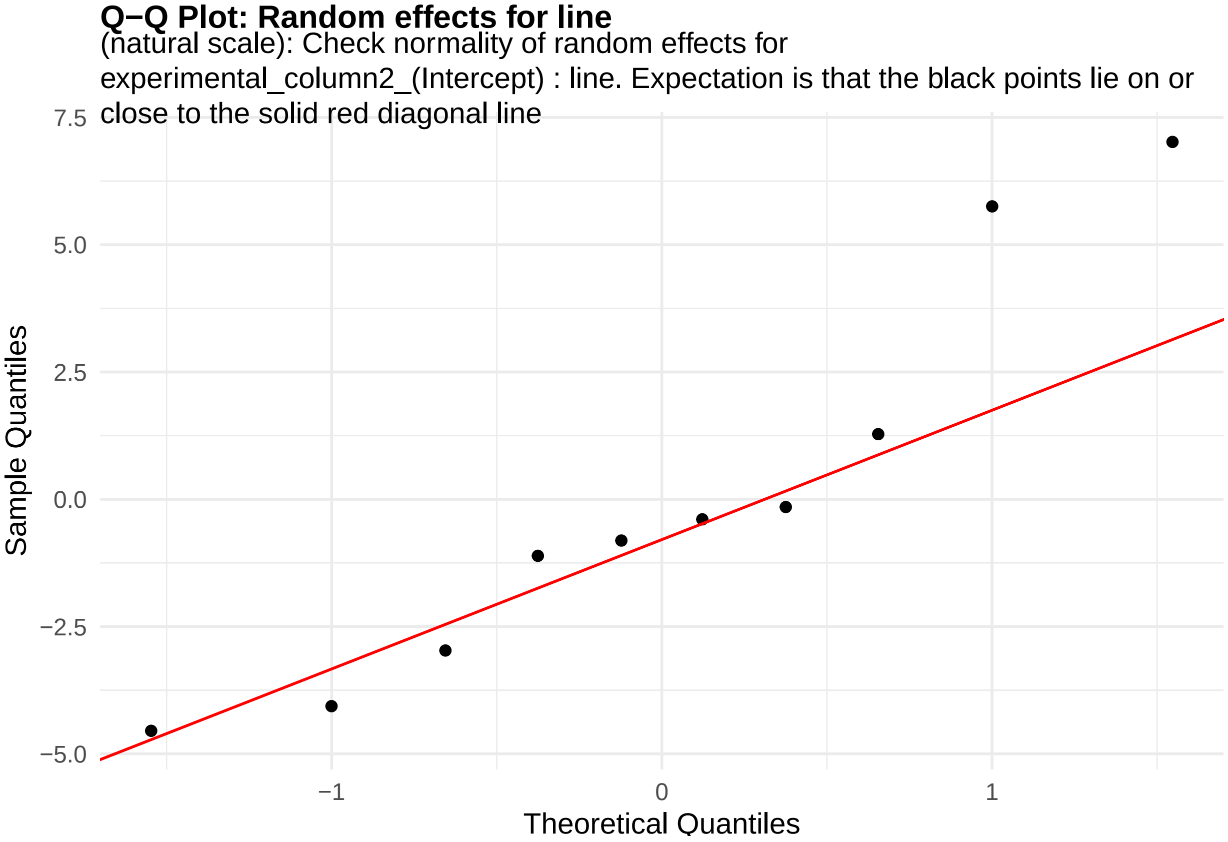


**Figure 4:** random_effects_QQ_1. Check normality of random effects for experimental_column2_(Intercept) (natural scale): line. Expectation is that the black points lie on or close to the solid red diagonal line. Check the Normal Q-Q plots for good and bad examples here: https://library.virginia.edu/data/articles/diagnostic-plots

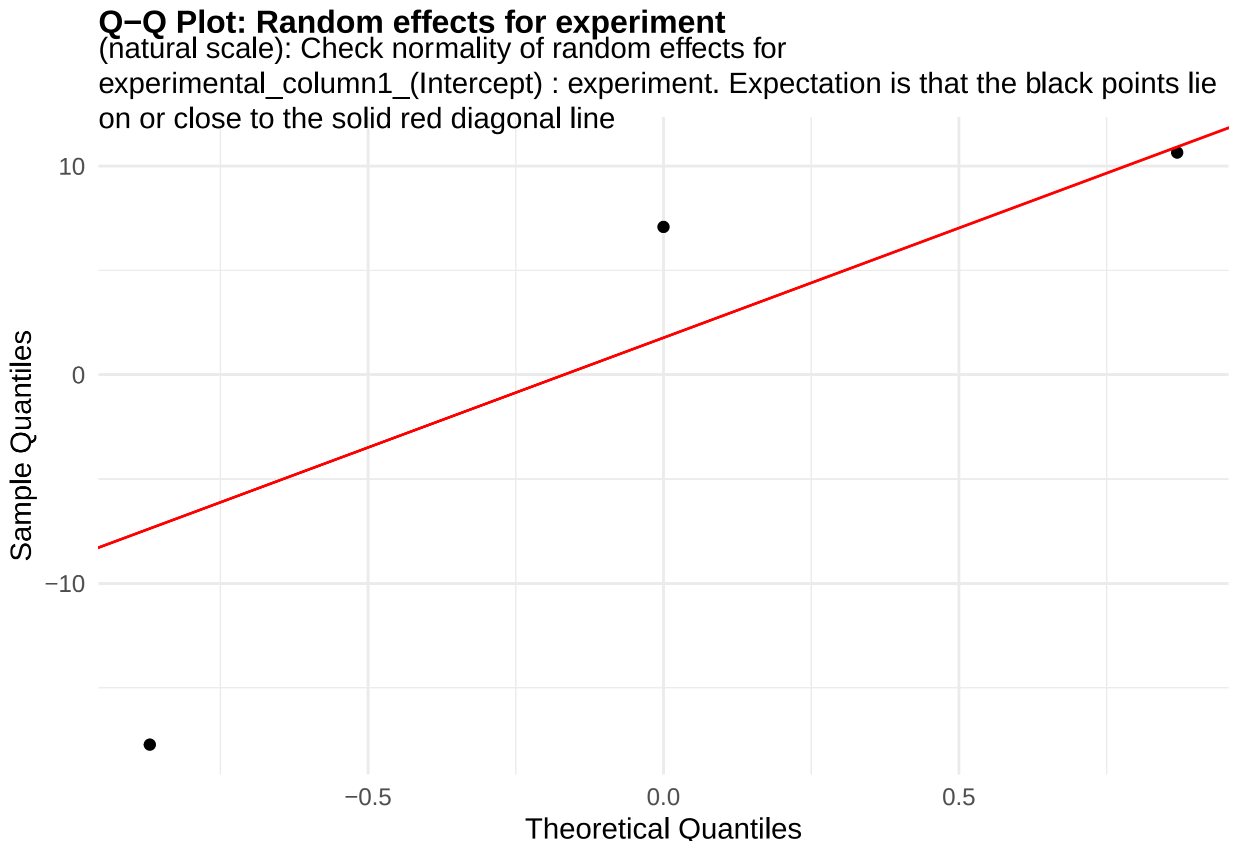


**Figure 5:** random_effects_QQ_2. Check normality of random effects for experimental_column1_(Intercept) (natural scale): experiment. Expectation is that the black points lie on or close to the solid red diagonal line. Check the Normal Q-Q plots for good and bad examples here: https://library.virginia.edu/data/articles/diagnostic-plots

```
cat("\n\n\n")
```

```
cooks_plots <- diagnose_res$cooks_plots$natural_scale
for(i in 1:length(cooks_plots)) {
  print(cooks_plots[[i]])
  cat(sprintf("\n\n\n**Figure %d:**\n\n", plot_no))
  plot_no <- plot_no + 1
}
```

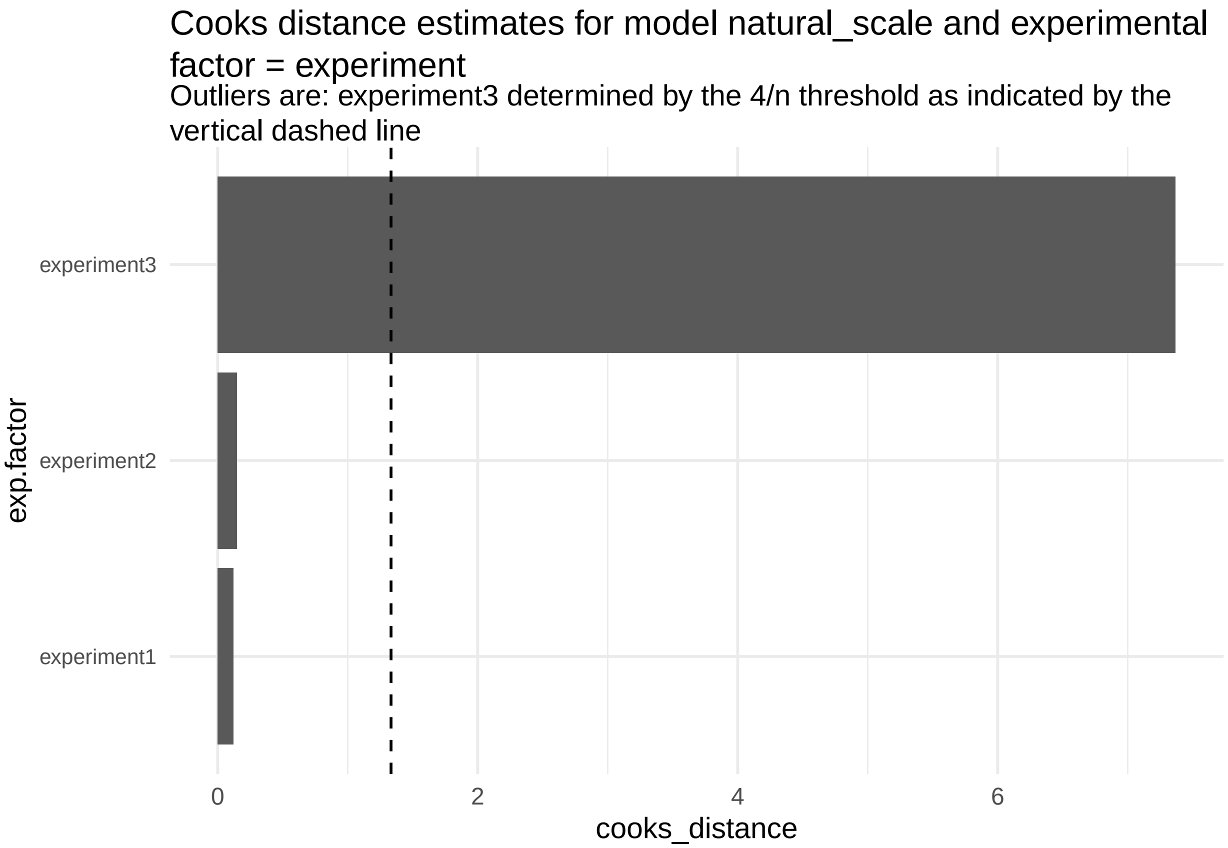
Cooks distance estimates for model natural_scale and experimental factor = experiment
Outliers are: experiment3 determined by the 4/n threshold as indicated by the vertical dashed line
experiment3
experiment2
experiment1
exp.factor
0
2
4
6
cooks_distance

**Figure 6:**

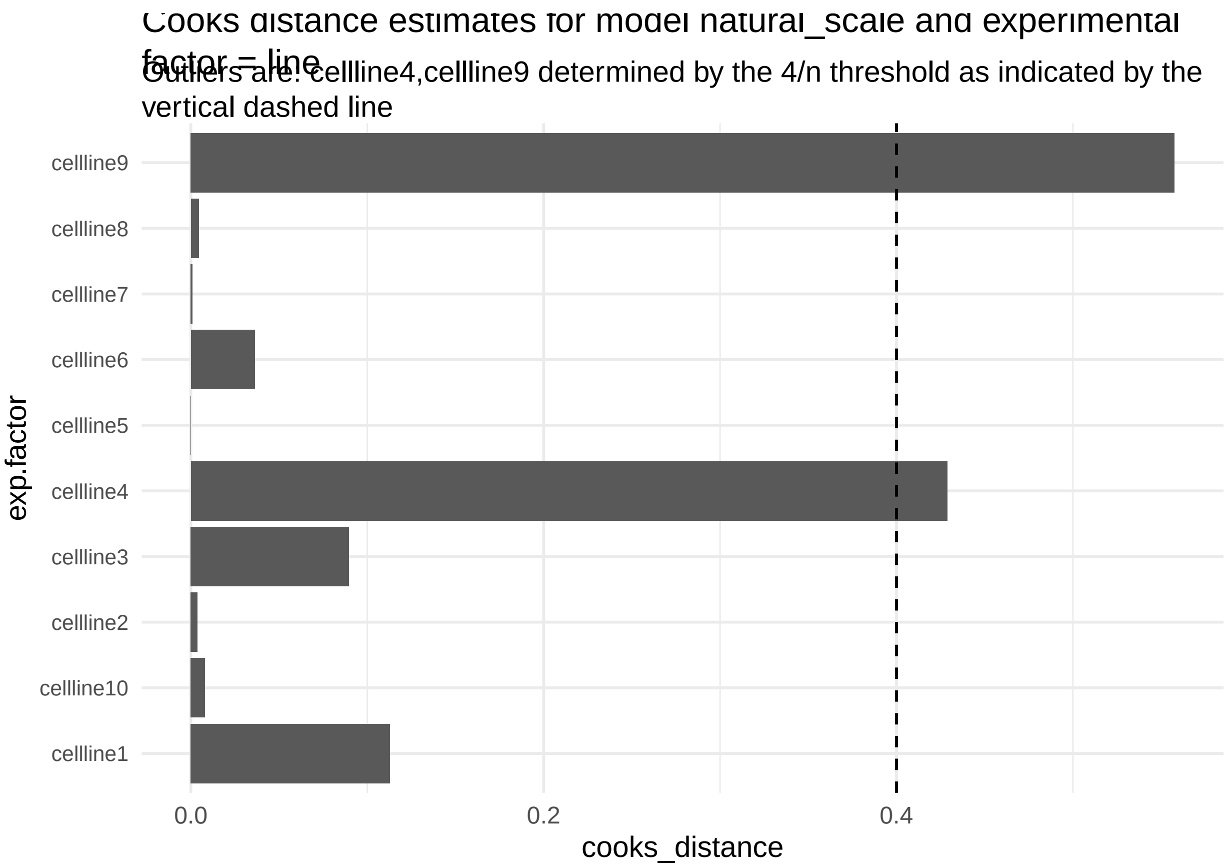


**Figure 7:**

```r
print("Inferred outlier groups are")
```

[1] "Inferred outlier groups are"

```r
print(diagnose_res$inferred_outlier_groups$natural_scale)
```

$experiment [1] "experiment3"

$line [1] "cellline4" "cellline9"

Now, let us look at the same set of QC plots for models on the log-transformed responses.

```r
plots <- diagnose_res$diagnostic_plots$log_scale$plots
captions <- diagnose_res$diagnostic_plots$log_scale$captions
for (i in seq_along(plots)) {
  print(plots[[i]])
  cat(sprintf("\n\n\n**Figure %d:** %s. %s\n\n", plot_no,names(plots)[i], captions[i]))
  plot_no <- plot_no + 1
}
```

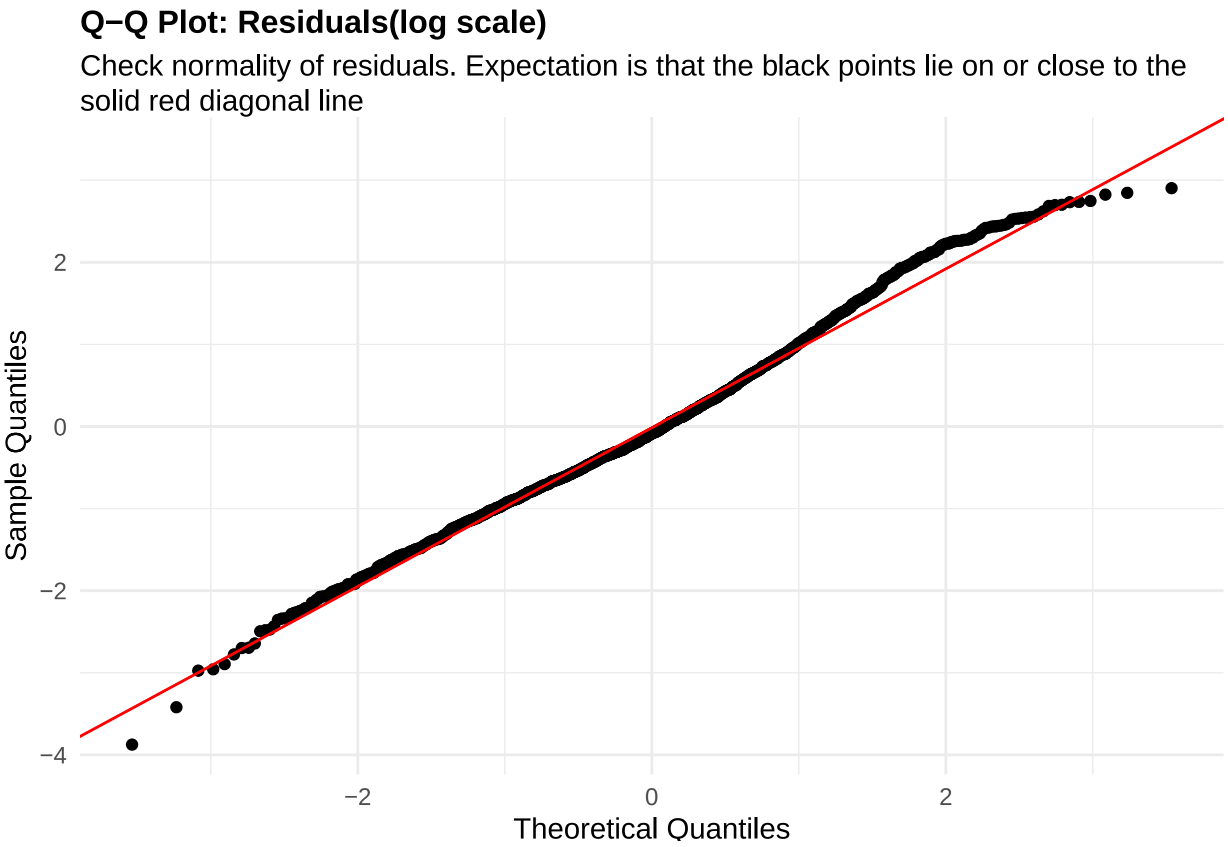


**Figure 8:** residuals_QQ. Check normality of residuals (log scale). Expectation is that the black points lie on or close to the solid red diagonal line. Check the Normal Q-Q plots for good and bad examples here: https://library.virginia.edu/data/articles/diagnostic-plots

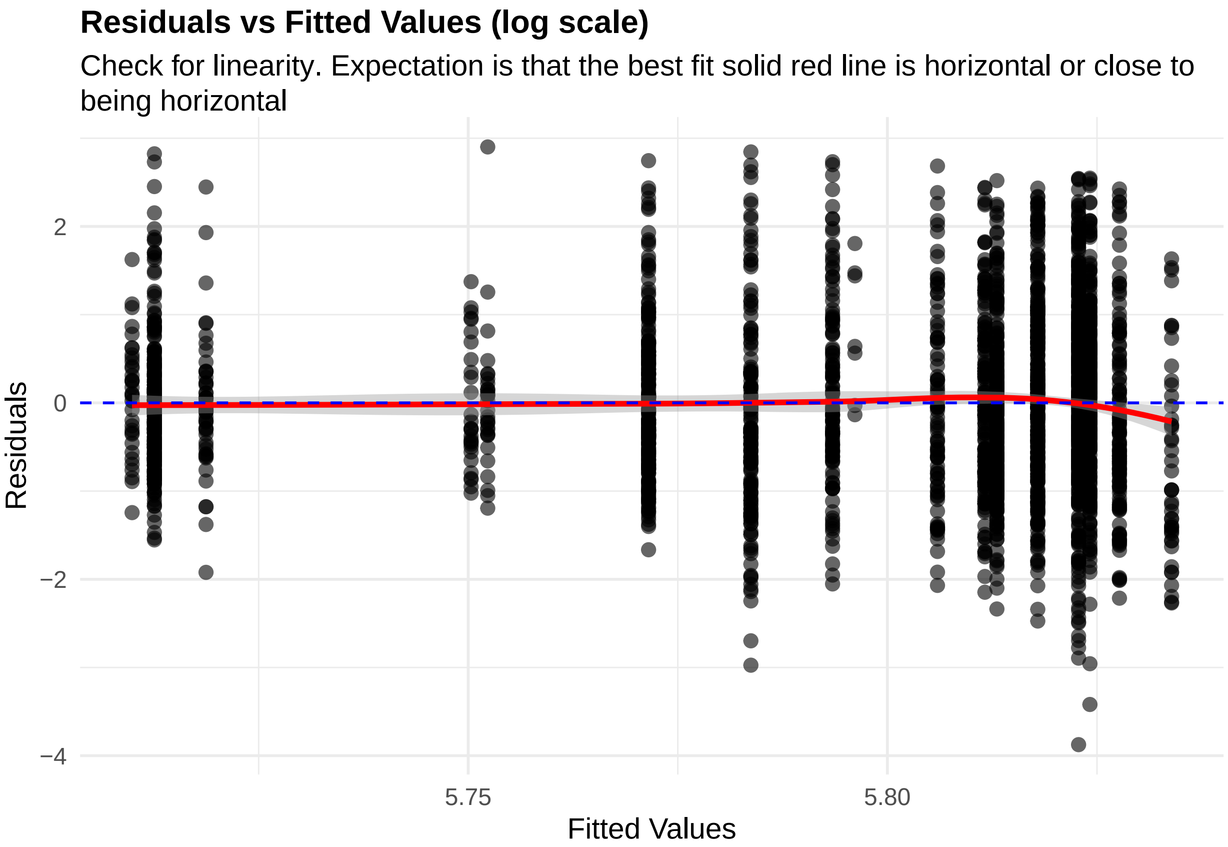


**Figure 9:** residuals_vs_fitted. Check for linearity (log scale). Expectation is that the best fit solid red line is horizontal or close to being horizontal. Check the Residuals vs Fitted plots for good and bad examples here: https://library.virginia.edu/data/articles/diagnostic-plots

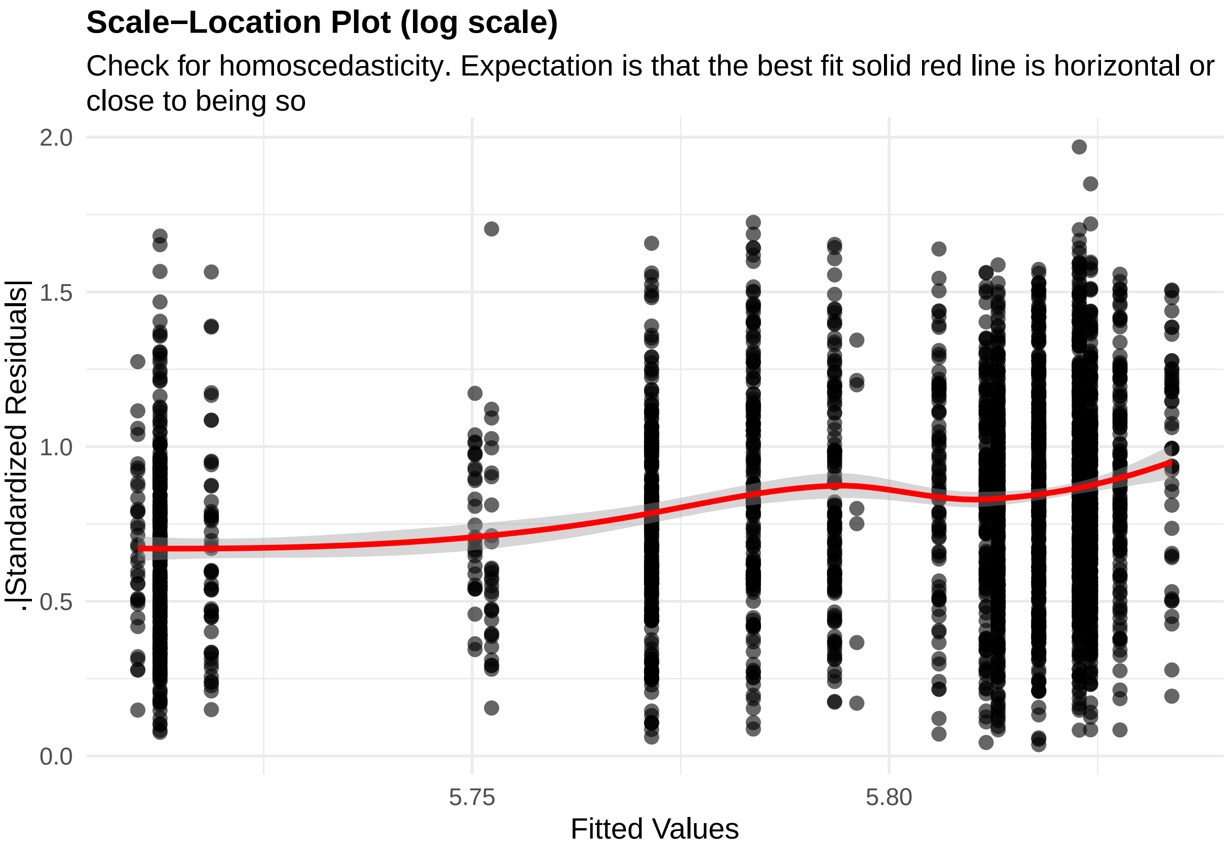


**Figure 10:** residuals_homoscedasticity. Check for homoscedasticity (log scale). Expectation is that the best fit solid red line is horizontal or close to being so. Check the Scale-Location plots for good and bad examples here: https://library.virginia.edu/data/articles/diagnostic-plots

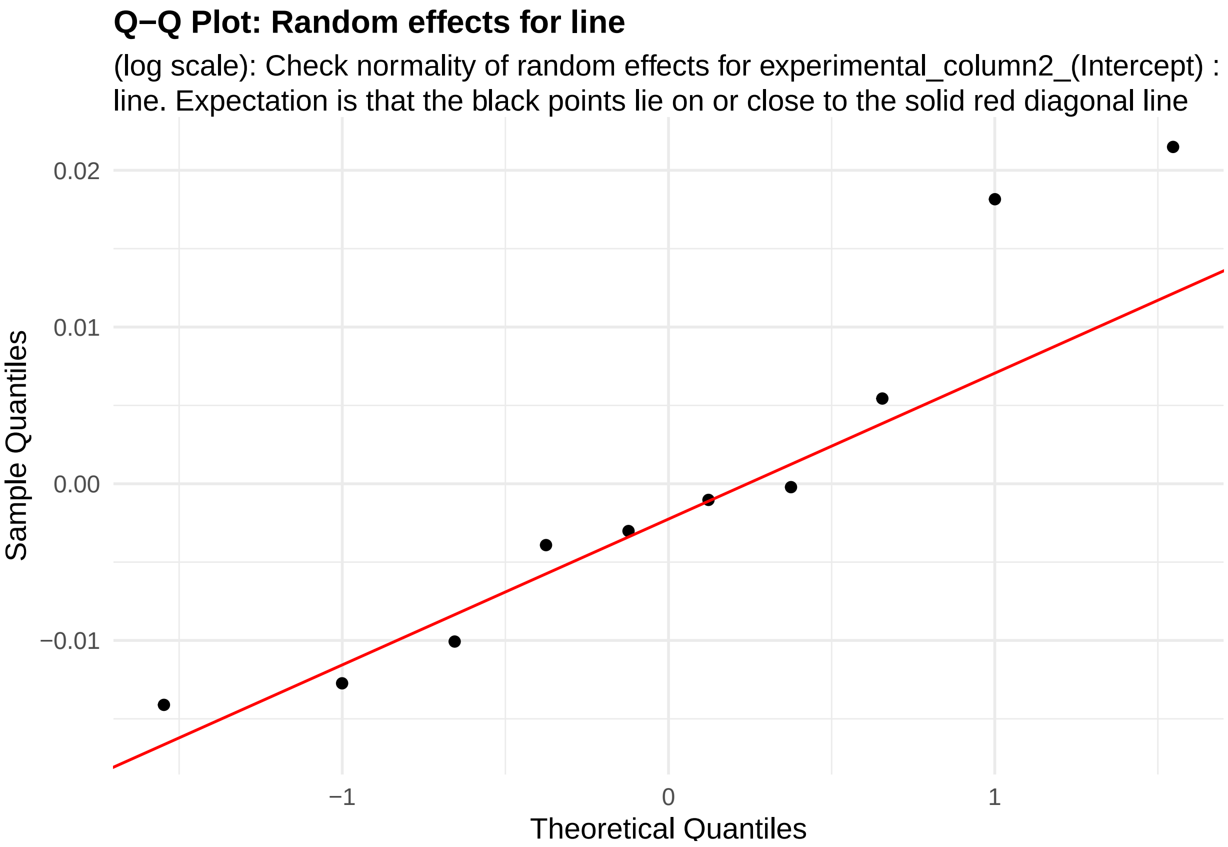


**Figure 11:** random_effects_QQ_1. Check normality of random effects for experimental_column2_(Intercept) (log scale): line. Expectation is that the black points lie on or close to the solid red diagonal line. Check the Normal Q-Q plots for good and bad examples here: https://library.virginia.edu/data/articles/diagnostic-plots

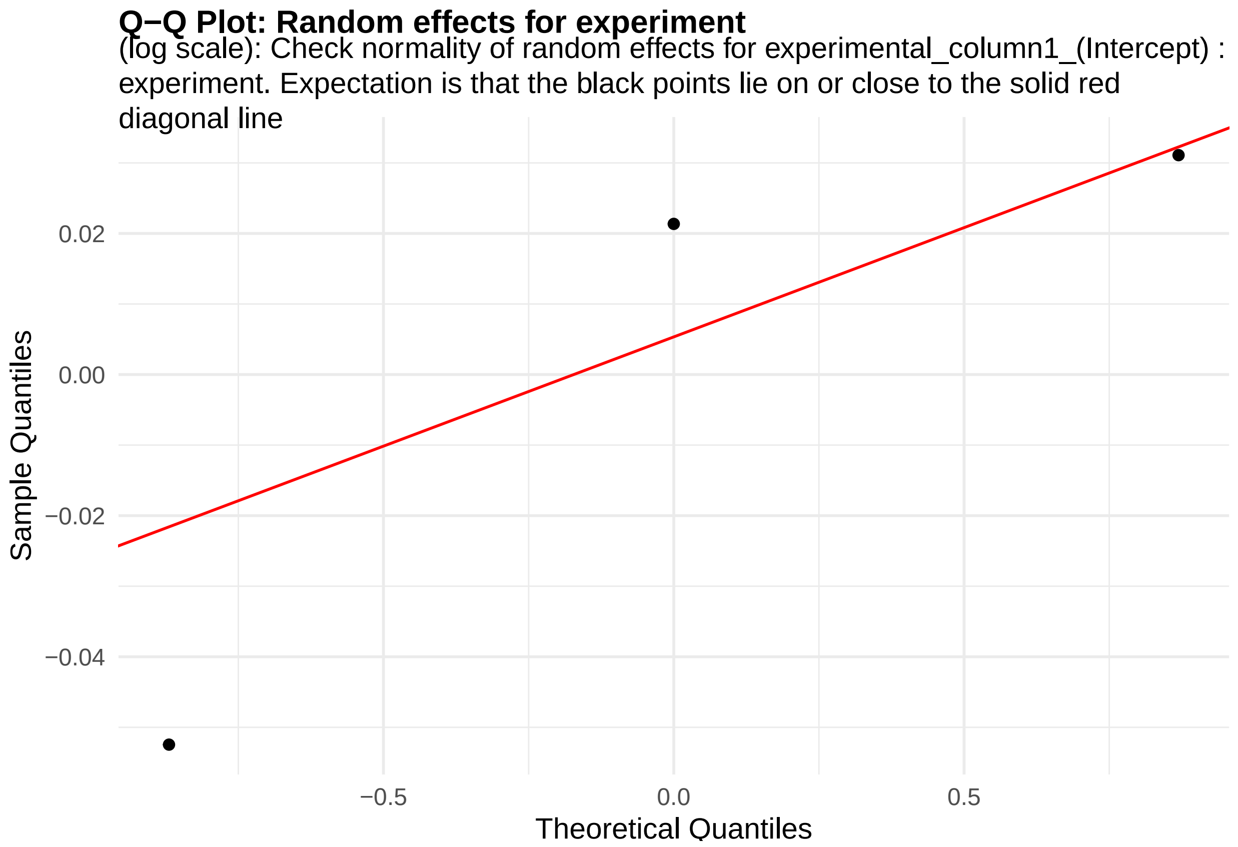


**Figure 12:** random_effects_QQ_2. Check normality of random effects for experimental_column1_(Intercept) (log scale): experiment. Expectation is that the black points lie on or close to the solid red diagonal line. Check the Normal Q-Q plots for good and bad examples here: https://library.virginia.edu/data/articles/diagnostic-plots

```
cooks_plots <- diagnose_res$cooks_plots$log_scale

cat("\n\n\n")
```

```
for(i in 1:length(cooks_plots)) {
  print(cooks_plots[[i]])
  cat(sprintf("\n\n\n**Figure %d:**\n\n", plot_no))
  plot_no <- plot_no + 1
}
```

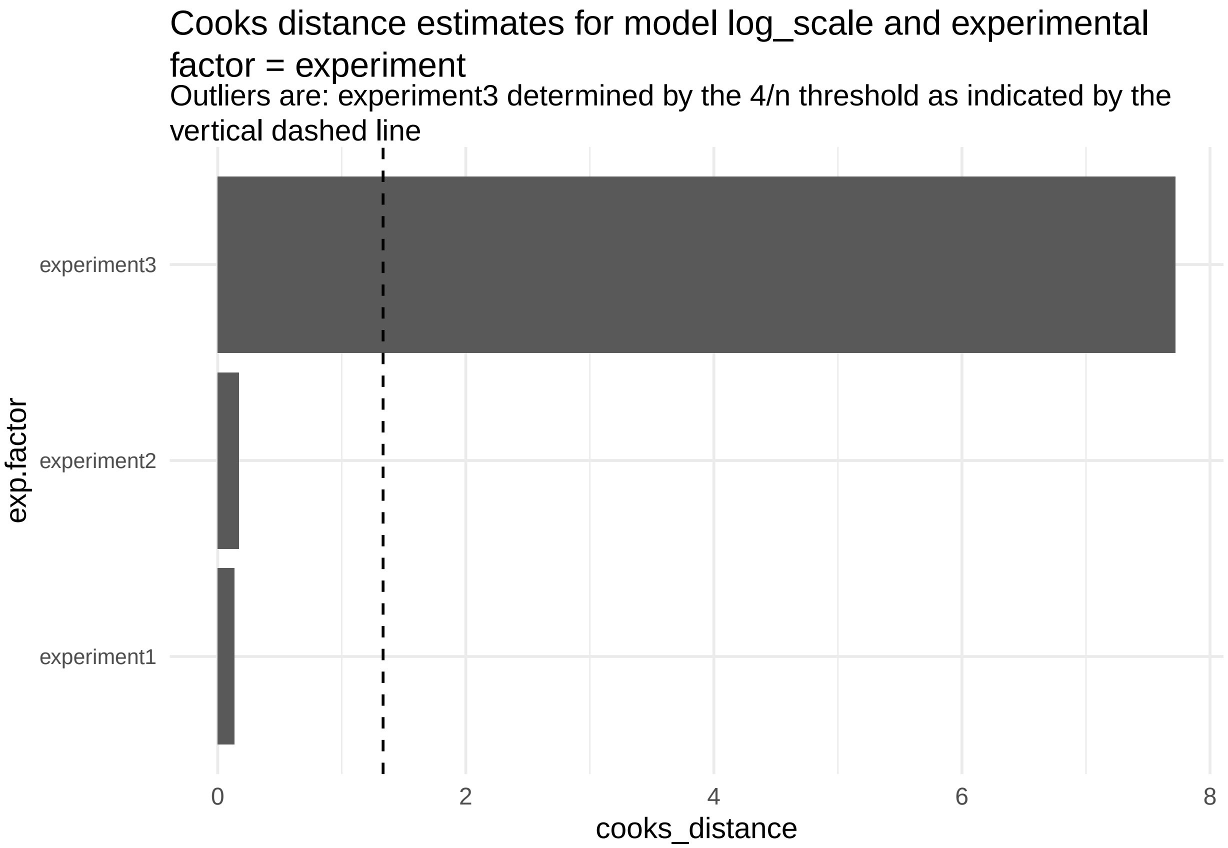
Cooks distance estimates for model log_scale and experimental factor = experiment
Outliers are: experiment3 determined by the 4/n threshold as indicated by the vertical dashed line
experiment3
experiment2
experiment1
exp.factor
0
2
4
6
8
cooks_distance

**Figure 13:**

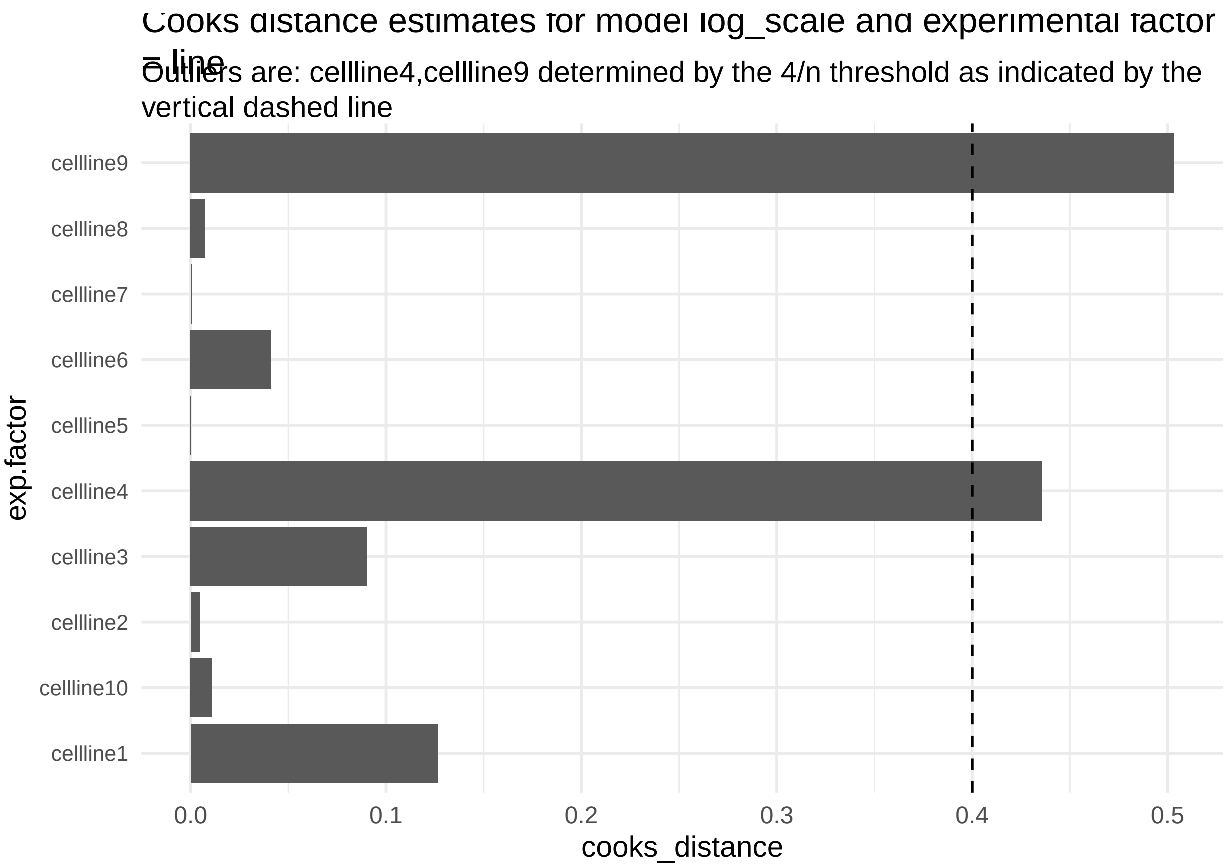


**Figure 14:**

```
print("Inferred outlier groups are")
```

[1] "Inferred outlier groups are"

```
print(diagnose_res$inferred_outlier_groups$natural_scale)
```

$experiment [1] "experiment3"

$line [1] "cellline4" "cellline9"

We see normality assumption is better met with the log-transformed model (Figure 6) compared to the model in the natural scale (Figure 1).

The code also outputs Cook's distance measures for each experiment and cell-line that is intended to capture how influential each of these were in determining the final parameter estimates in the LMM model.

The observations in experimental batch 3 and those for cell lines 4 and 9 were identified as outliers using a 4/n (where n is the number of replicates, n=3 for experimental batches and n=10 for cell-lines) threshold applied to the estimates of Cook's distance for the estimated model parameters. This warranted further investigation on whether or not there was something unusual with the observations made in experimental batch 3 (example: the images from the microscope displayed low signal to noise and may have skewed the measurements) and also for cell line 4# and 9# (example: differentiation runs, genetic background was different from the other lines) that would suggest observations linked to this experiment and cell lines may need to be removed from the analysis. Our investigation did not indicate anything untoward and therefore we ran a sensitivity analysis where we dropped all observations linked to either experiment 3, cell line 4# or cell line 9#, refit the model and visualized the resulting change in ALS status associated mean soma perimeter value. There were only relatively slight changes in the estimated changes in the mean, so that the conclusions we would draw of an increase in the soma cell perimeter in ALS lines remains irrespective of whether or not we drop observations from the experimental batch or the two cell lines.

We will use the log transformed cell_size2 data for parameter estimation using the following commands.

```
design_update <- design
data_update <- diagnose_res$Data_updated
print(colnames(data_update))
```

```
##  [1] "cell_size2_logTransformed" "experiment"
##  [3] "line"                      "classification"
##  [5] "cell_size1"                "cell_size2"
##  [7] "cell_size3"                "cell_size4"
##  [9] "cell_size5"                "cell_size6"
```

```
design_update@response_column <-  "cell_size2_logTransformed"
estimate_res <- getEstimatesOfInterest(data_update, design_update, model,print_plots = FALSE)
```

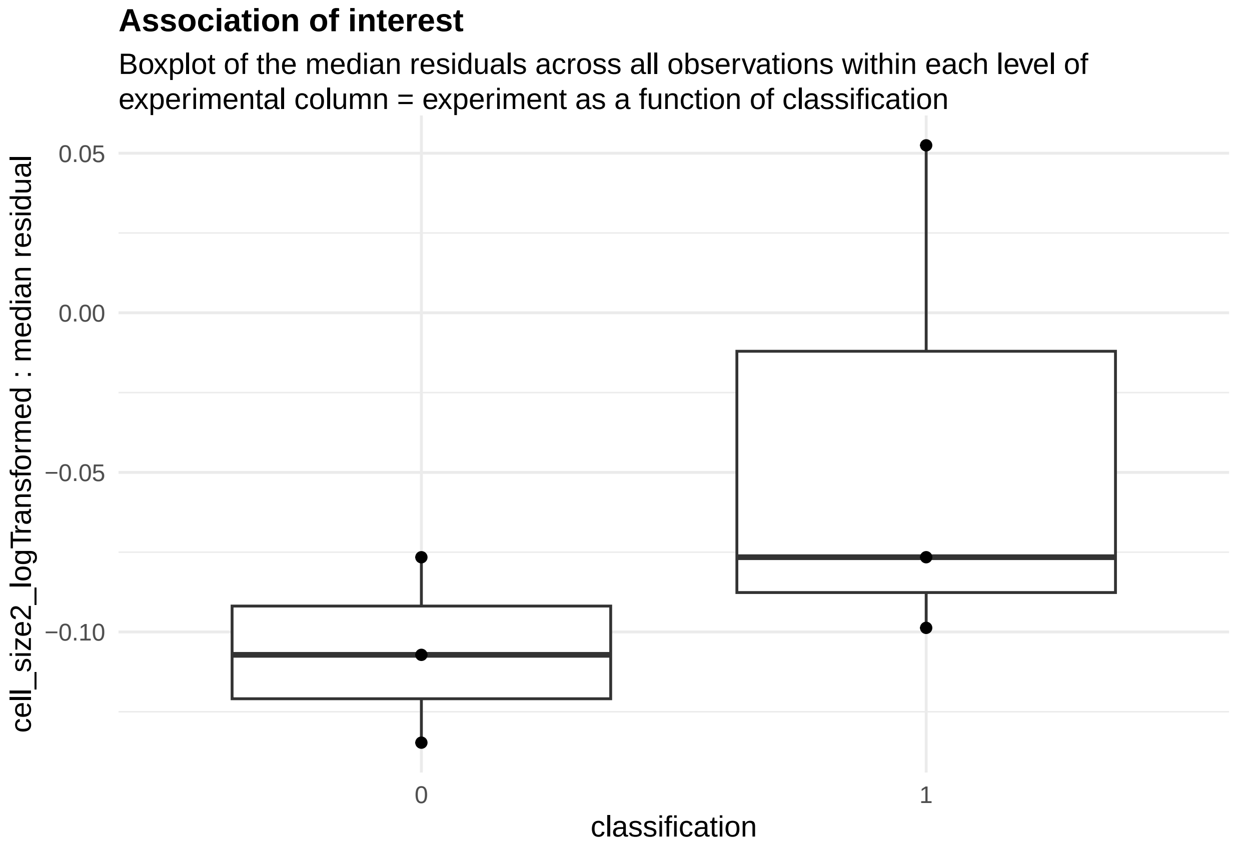


**Figure 15:**

The median residual boxplot shows the differences in the distribution of the response variable for the two conditions.

```
##print model summary output
print(estimate_res[[1]])
```

```
## Linear mixed model fit by REML. t-tests use Satterthwaite's method [
## lmerModLmerTest]
## Formula: response_column ~ condition_column + (1 | experimental_column1) +
##     (1 | experimental_column2)
##    Data: fixed_global_variable_data
##
```

```
## REML criterion at convergence: -3678.5
##
## Scaled residuals:
##     Min      1Q  Median      3Q     Max
## -3.8747 -0.6665 -0.0869  0.6377  2.9014
##
## Random effects:
##  Groups               Name        Variance  Std.Dev.
##  experimental_column2 (Intercept) 0.0002438 0.01561
##  experimental_column1 (Intercept) 0.0021256 0.04610
##  Residual                         0.0129115 0.11363
## Number of obs: 2458, groups:  experimental_column2, 10; experimental_column1, 3
##
## Fixed effects:
##                   Estimate Std. Error      df t value Pr(>|t|)
## (Intercept)        5.77509    0.02764 2.19350 208.924  9.3e-06 ***
## condition_column1  0.03072    0.01182 8.47900   2.599   0.0302 *
## ---
## Signif. codes:  0 '***' 0.001 '**' 0.01 '*' 0.05 '.' 0.1 ' ' 1
##
## Correlation of Fixed Effects:
##             (Intr)
## cndtn_clmn1 -0.171
```

As a result of linear mixed model-based association analysis, the response variable was found to be significantly associated with the condition variable with an estimate of 0.03.

We can also estimate the statistical needed to estimate the observed differences in mean perimeter across 15 experimental batches

```
power_param@max_size <- 15
power_res <- calculatePower(data_update, design_update, model, power_param)
```

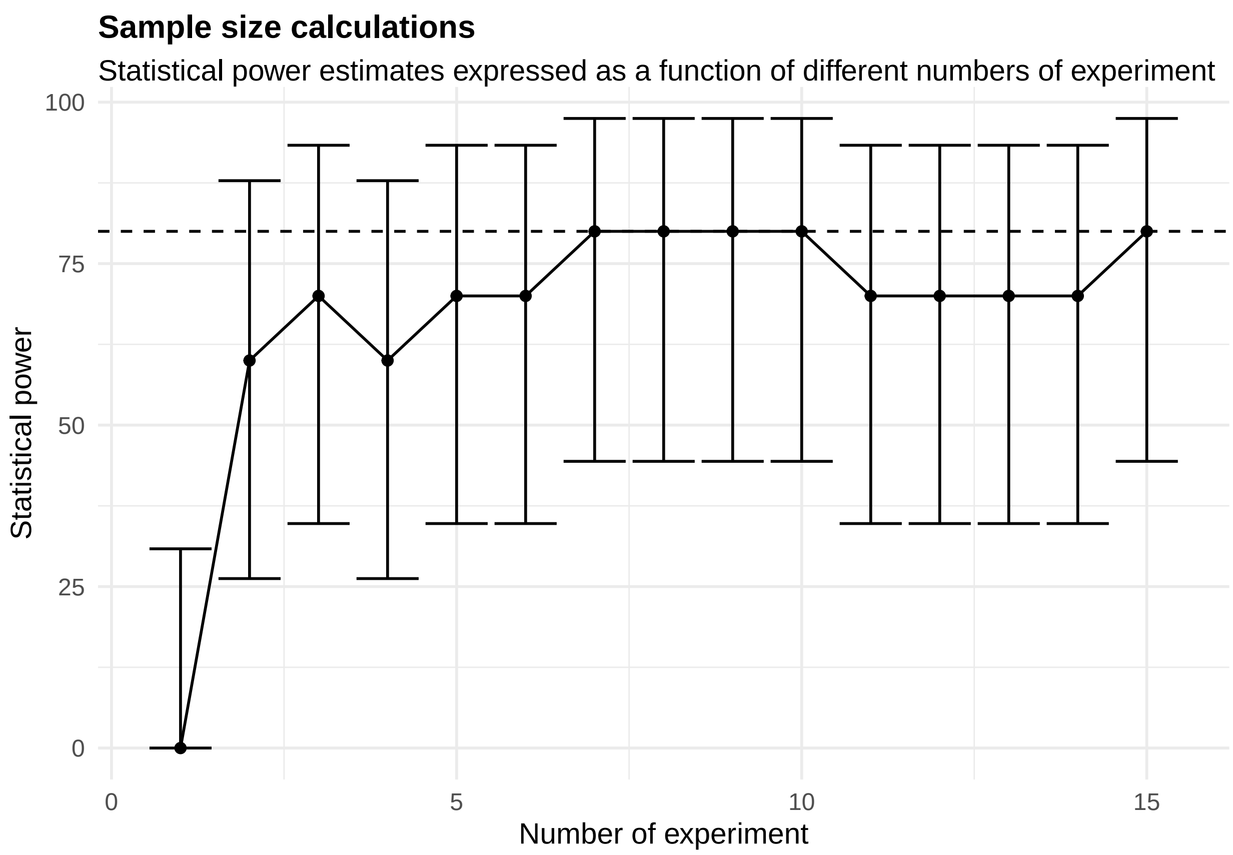


[[1]]

**Figure 16:**

The power curve shows that at least eight or more experimental batches are required to achieve a power of 80% or greater. The points in the plot represent the mean estimate of power over the simulations while the error bars represent the 95% confidence intervals.

Accordingly, we will now load data from 8 experimental batches and estimate our parameter of interest.

```
full_data <- plate_assay_full_data

##filter the top 5% of the observations
full_data %<>% filter(perim_2th_effect < quantile(perim_2th_effect, 0.95))

design@response_column <- "perim_2th_effect"
design@condition_column <- "classif"
full_diag_res <- diagnoseDataModel(full_data, design, model)
full_data_update <- full_diag_res$Data_updated
full_design <- design
full_design@response_column <- "perim_2th_effect_logTransformed"
```

Note now, none of the experimental batches or the 13 cell-lines are considered outliers or unduly influencing the final parameter estimates.

```
cooks_plots <- full_diag_res$cooks_plots$log_scale
cat("\n\n\n")
```

```
for(i in 1:length(cooks_plots)) {
  print(cooks_plots[[i]])
  cat(sprintf("\n\n\n**Figure %d:**\n\n", plot_no))
  plot_no <- plot_no + 1
```

```
}
```

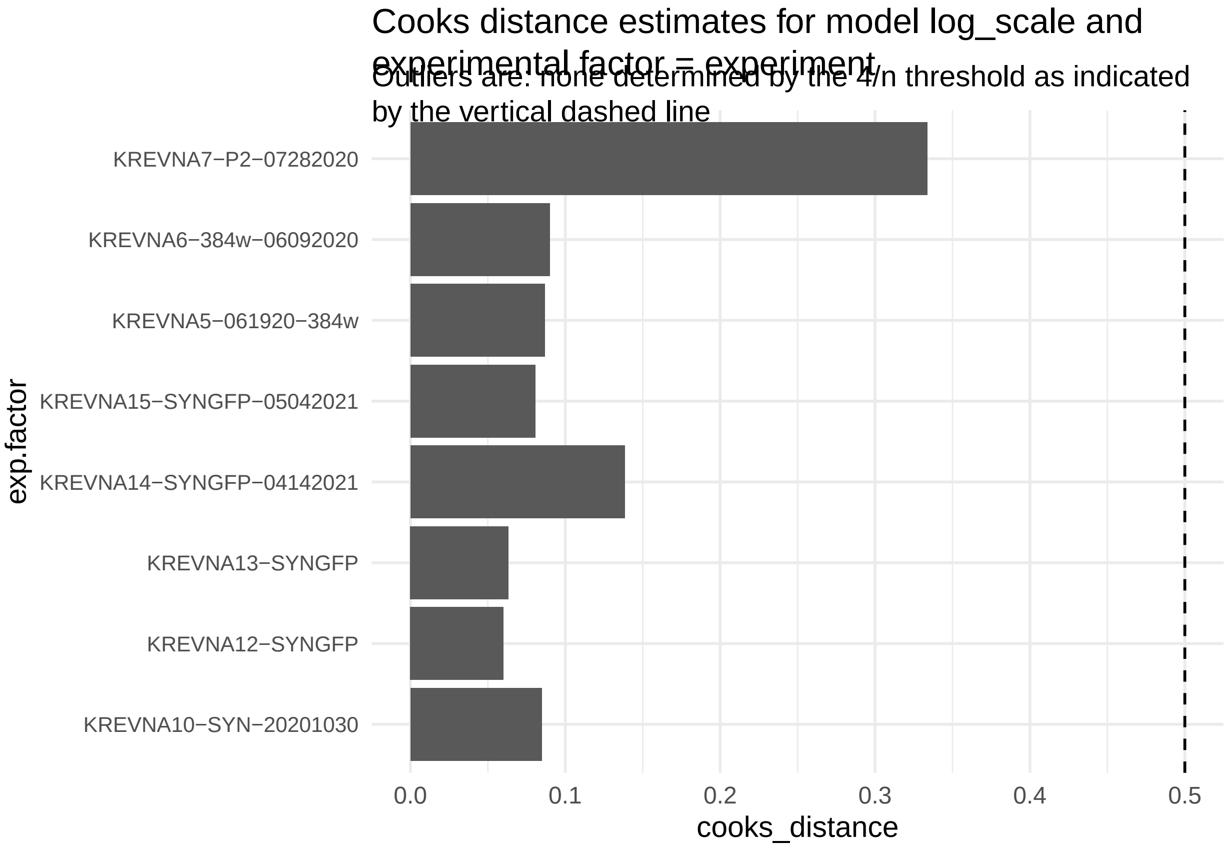

Cooks distance estimates for model log_scale and experimental factor = experiment
Outliers are: none determined by the 4/n threshold as indicated by the vertical dashed line
KREVNA7−P2−07282020
KREVNA6−384w−06092020
KREVNA5−061920−384w
KREVNA15−SYNGFP−05042021
KREVNA14−SYNGFP−04142021
KREVNA13−SYNGFP
KREVNA12−SYNGFP
KREVNA10−SYN−20201030
exp.factor
0.0
0.1
0.2
0.3
0.4
0.5
cooks_distance


**Figure 17:**

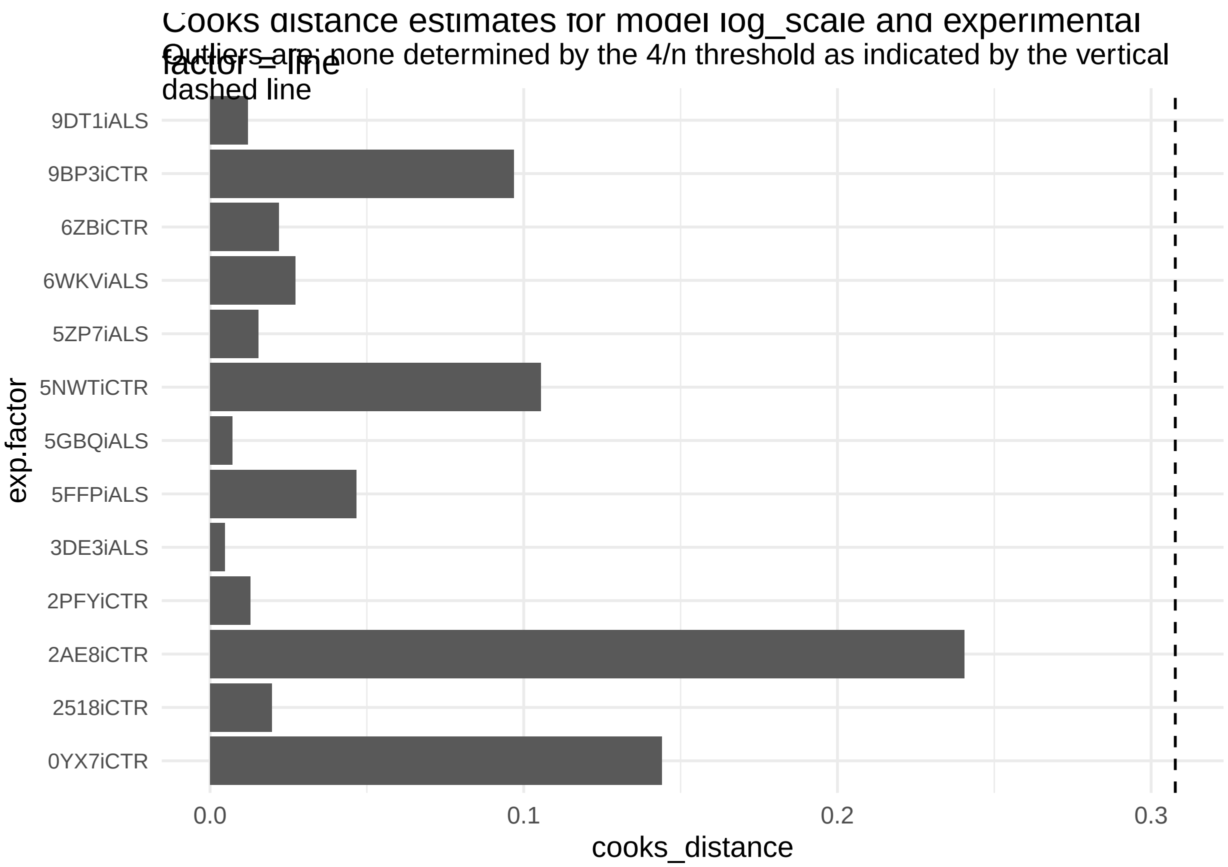


**Figure 18:**

```r
print("Inferred outlier groups are")
```

[1] "Inferred outlier groups are"

```r
print(diagnose_res$inferred_outlier_groups$natural_scale)
```

$experiment [1] "experiment3"

$line [1] "cellline4" "cellline9"

We derive an estimate of 0.027 as the mean difference in perimeter with an associated significance of 0.006 using the data from 8 experimental batches.

```r
full_res <- getEstimatesOfInterest(full_data_update, full_design, model, print_plots = FALSE)
print(full_res[[1]])
```

```
## Linear mixed model fit by REML. t-tests use Satterthwaite's method [
## lmerModLmerTest]
## Formula: response_column ~ condition_column + (1 | experimental_column1) +
##     (1 | experimental_column2)
##    Data: fixed_global_variable_data
##
## REML criterion at convergence: -10215.6
##
## Scaled residuals:
##     Min      1Q  Median      3Q     Max
## -4.1653 -0.6837 -0.0505  0.6413  3.0906
##
```

```
## Random effects:
##  Groups               Name        Variance Std.Dev.
##  experimental_column2 (Intercept) 0.000168 0.01296 
##  experimental_column1 (Intercept) 0.000883 0.02971 
##  Residual                         0.010516 0.10255 
## Number of obs: 5989, groups:  experimental_column2, 13; experimental_column1, 8
## 
## Fixed effects:
##                   Estimate Std. Error        df t value Pr(>|t|)    
## (Intercept)       5.753966   0.011811 10.213599 487.158   <2e-16 ***
## condition_column1  0.027159   0.007924 10.925415   3.427   0.0057 ** 
## ---
## Signif. codes:  0 '***' 0.001 '**' 0.01 '*' 0.05 '.' 0.1 ' ' 1
## 
## Correlation of Fixed Effects:
##             (Intr)
## cndtn_clmn1 -0.309
```

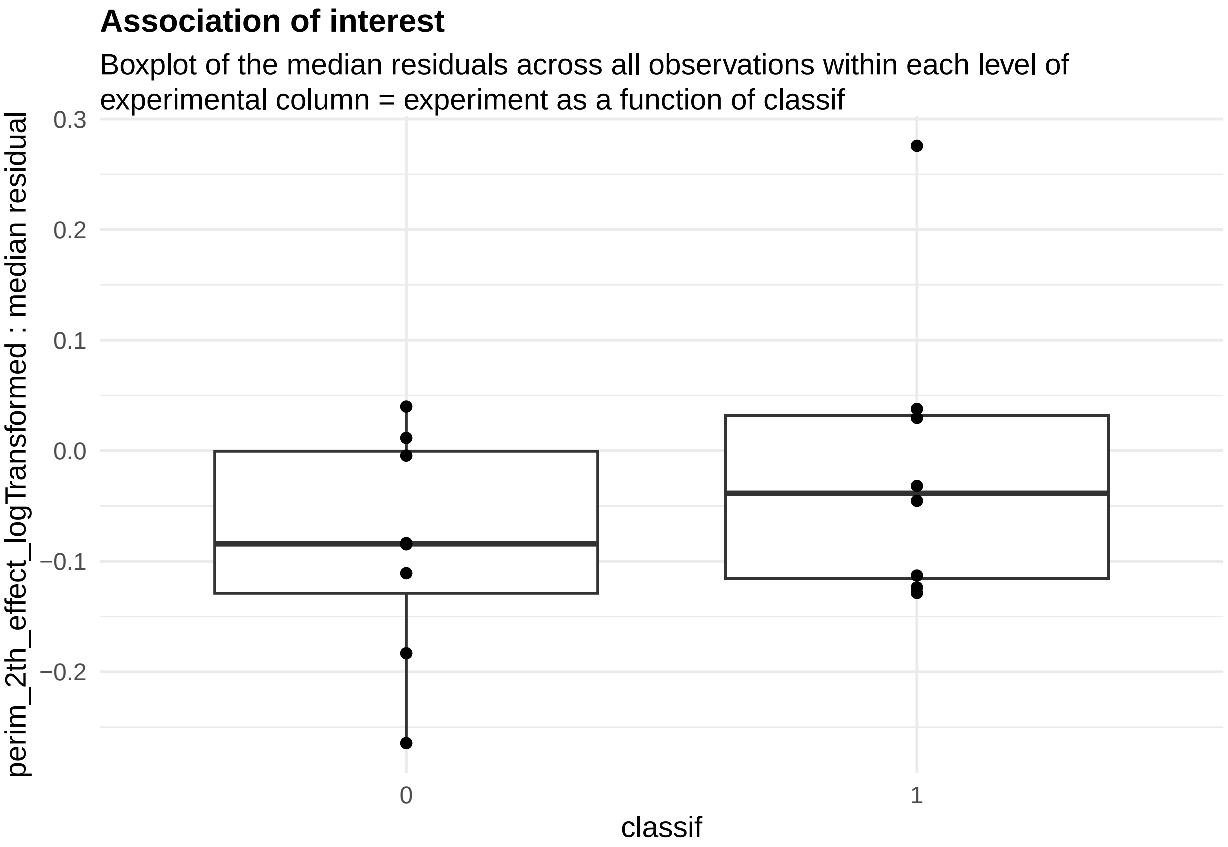


**Figure 19:**

##### 2.2 Single-cell RNA-seq assays

Here we will use data derived from the single-nuclei RNA-seq experiments performed in Koutsodendris et al[]. paper One of the goals in these experiments was to assess the effect of the APOE gene isoform on brain pathology in a mouse model of Alzheimer's Disease (AD). Specifically, the *APOE4* haplotype is a known risk factor for AD compared to those with the *APOE3* gene isoform. Koutsodendris et a., investigates the role of expression of neuronal *APOE4* expression in the development of disease pathology in the *APOE4* knock-in mouse model. The scRNA seq dataset consists of over 1000 cells derived from the hippocampus brain region across 4 mice harboring the human *APOE4* gene knocked-in the mouse *Apoe* locus and 4 other mice (called E4-KI-Syn1-Cre) with the human *APOE4* gene also knocked-in in the same mouse. However, the *APOE4* expression in these mice is knocked out specifically in the neurons. The single cell data allowed the identification of multiple clusters or cell-types including cluster 7 representing a group of excitatory neurons.

##### 2.2.1 Cell-type/cluster associations

One typical use of scRNA seq data is clustering analysis. First, we can ask whether the proportion of nuclei derived from APOE4-KI mice differs from the proportion of nuclei derived from APOE4-KI-Syn1-Cre mice in cluster 7. We want to examine the proportion of cells in each mouse assigned cluster 7 by testing whether there were differences in the log odds of cluster membership in cluster 7 between APOE4-KI mice and APOE4-KI-Syn1-Cre. The data consists of the number of nuclei per animal present in the cluster and also the total number of nuclei that were isolated from this animal.

**2.2.1.1 Associations with genotype** The following commands load data and set parameters to specify the experimental design, distribution information, and power simulation conditions.

The first few lines of data display cell meta information.

```
template_dir <- system.file("input_templates/single_cell_data/celltype_proportions_with_genotype", packa
data <- snRNAseq_cluster_count_data
head(data)
```

```
##   sample_id      animal_model total_numbers_of_cells_per_sample cluster_id
## 1       S10          PS19-fE4                              6552          7
## 2       S11          PS19-fE4                              6214          7
## 3       S12 PS19-fE4 Syn1-Cre                              7873          7
## 4       S13 PS19-fE4 Syn1-Cre                              6577          7
## 5        S4          PS19-fE4                             16223          7
## 6        S5          PS19-fE4                             12357          7
##   number_of_cells_per_sample_in_cluster          Genotype Mouse_. Sex PS19 Cre
## 1                                   263          PS19-fE4     154   F    +   -
## 2                                     8          PS19-fE4     413   M    +   -
## 3                                    58 PS19-fE4_Syn1-Cre     305   M    + +_-
## 4                                     1 PS19-fE4_Syn1-Cre     504   F    + +_-
## 5                                  2139          PS19-fE4     129   F    +   -
## 6                                  1811          PS19-fE4     156   M    +   -
##   ApoE      DOB Age_Perfused Date_Perfused Date_of_Nuc_Isolation
## 1 E4_4 10_30_19         10.3        9/7/20                9/9/21
## 2 E4_4  10_2_20         10.1        8/4/21                9/9/21
## 3 E4_4  3_19_20         10.5        1/8/21                9/9/21
## 4 E4_4   9_8_20         10.8        8/4/21                9/9/21
## 5 E4_4 12_18_19          9.9      10/16/20                9/7/21
## 6 E4_4 12_18_19          9.9      10/16/20                9/7/21
##   Hippocampus_Vol_mm X._AT8_Coverage_Area X._GFAP_Coverage_Area
## 1               6.75                31.52                 31.75
## 2               8.88                 4.43                  4.98
## 3               9.26                15.43                  5.60
## 4               7.58                 6.92                 11.90
## 5               3.73                69.89                 38.28
## 6               3.62                88.85                 39.84
##   X._S100B_Coverage_Area X._IBA1_Coverage_Area X._CD68_Coverage_Area
## 1                   6.94                 20.20                 10.45
## 2                   1.02                 10.09                  1.58
```

```
## 3                   0.38                   1.17                   0.42
## 4                   5.05                  13.09                   3.49
## 5                   9.71                  30.51                  25.49
## 6                  18.11                  31.49                  19.85
##   X._MBP_Coverage_Area X._OPC_Coverage_Area
## 1                 9.43                10.30
## 2                 4.32                 6.04
## 3                14.08                24.32
## 4                10.21                21.60
## 5                13.28                11.68
## 6                 7.72                10.95
```

The following command reads parameters from json files and print corresponding objects, allowing us to check which parameters have been assigned.

```r
##load the design
design <- readDesign(file.path(template_dir, "design_celltype.json"))
print(design)
```

```
## An object of class "RMeDesign"
## Slot "response_column":
## [1] "number_of_cells_per_sample_in_cluster"
##
## Slot "covariate":
## [1] "Sex"
##
## Slot "condition_column":
## [1] "Genotype"
##
## Slot "condition_is_categorical":
## [1] TRUE
##
## Slot "experimental_columns":
## [1] "sample_id"
##
## Slot "crossed_columns":
## NULL
##
## Slot "total_column":
## [1] "total_numbers_of_cells_per_sample"
##
## Slot "outlier_alpha":
## [1] 0.05
##
## Slot "na_action":
## [1] "complete"
##
## Slot "include_interaction":
## [1] FALSE
##
## Slot "random_slope_variable":
## NULL
##
## Slot "covariate_is_categorical":
## [1] TRUE
```

```r
##load the probability model
model <- readProbabilityModel(file.path(template_dir, "prob_model.json"))
print(model)
```

```
## An object of class "ProbabilityModel"
## Slot "error_is_non_normal":
## [1] TRUE
##
## Slot "family_p":
## [1] "binomial"
```

```r
##load the relevant power parameters
power_param <- readPowerParams(file.path(template_dir, "power_param.json"))
print(power_param)
```

```
## An object of class "PowerParams"
## Slot "target_columns":
## [1] "sample_id"
##
## Slot "power_curve":
## [1] 1
##
## Slot "nsimn":
## [1] 10
##
## Slot "levels":
## [1] 1
##
## Slot "max_size":
## [1] 1
##
## Slot "alpha":
## [1] 0.05
##
## Slot "breaks":
## NULL
##
## Slot "effect_size":
## NULL
##
## Slot "icc":
## NULL
```

Now let's use the 'diagnoseDataModel' function to diagnose the data quality and distributional assumptions of the model.

```r
diagnose_res <- diagnoseDataModel(data, design, model)
```

```
## [1] "No outlier detected from the raw Data"
## [1] "No outlier detected from the raw Data"
```

To identify all the models fit to the data. There were no outliers identified so only one model fit to natural scale is fit.

```r
print(diagnose_res$models)
```

```
## [1] "Natural scale"
```

```r
plots <- diagnose_res$diagnostic_plots$natural_scale$plots
captions <- diagnose_res$diagnostic_plots$natural_scale$captions
for (i in seq_along(plots)) {
  print(plots[[i]])
    cat(sprintf("\n\n\n**Figure %d:** %s. %s\n\n", plot_no,names(plots)[i], captions[i]))
  plot_no <- plot_no + 1
}
```

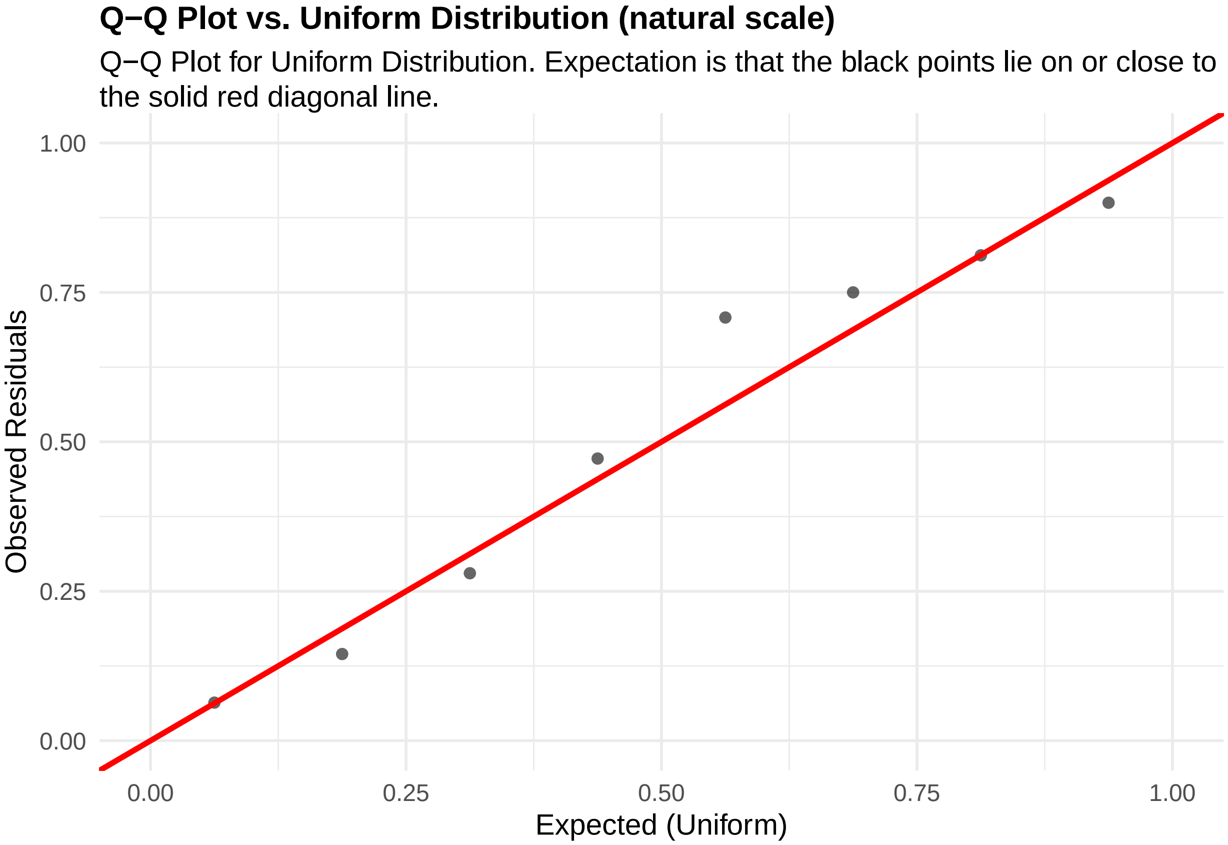


**Figure 20:** residuals_QQ. Q-Q Plot for Uniform Distribution. Expectation is that the black points lie on or close to the solid red diagonal line. Check the Q-Q plots for good and bad examples here: https://library.virginia.edu/data/articles/diagnostic-plots

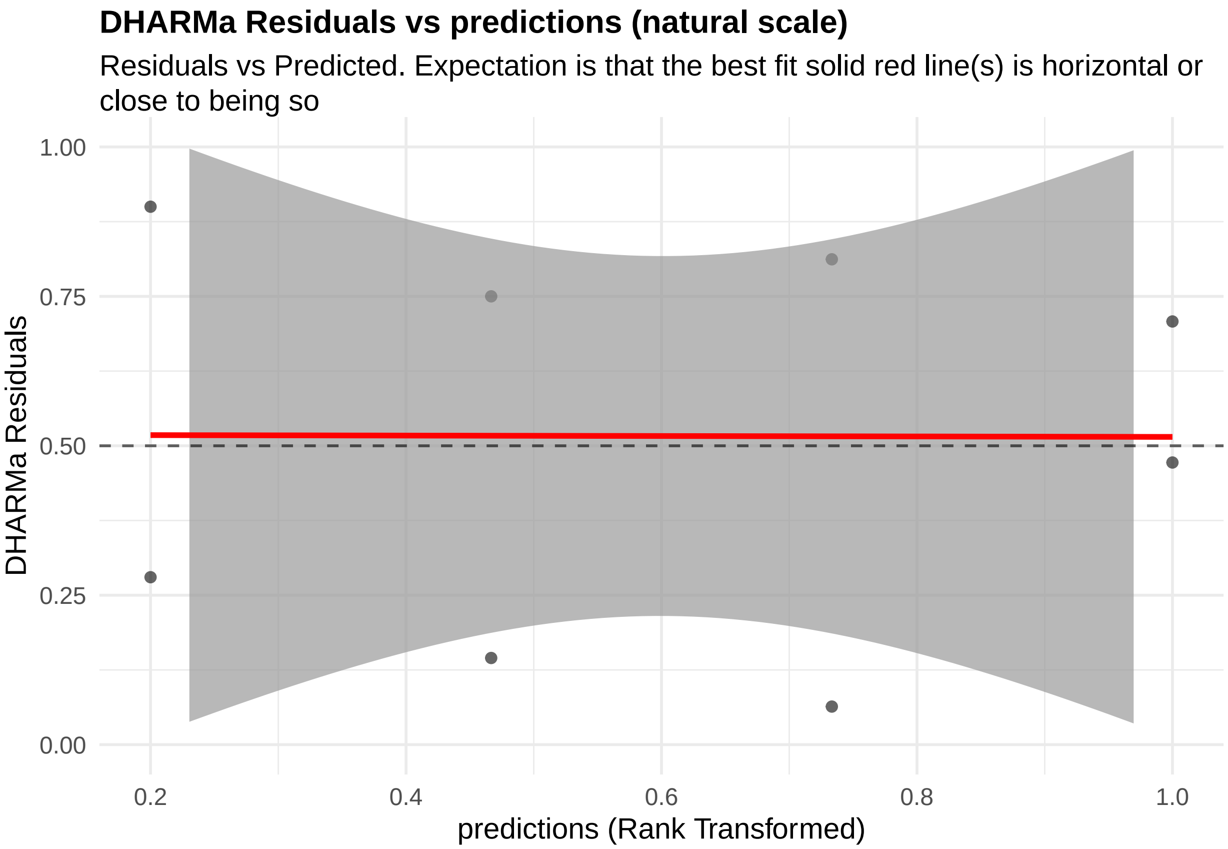


**Figure 21:** residuals_vs_predicted. Residuals vs Predicted Check the Residuals vs Fitted plots for good and bad examples here: https://library.virginia.edu/data/articles/diagnostic-plots

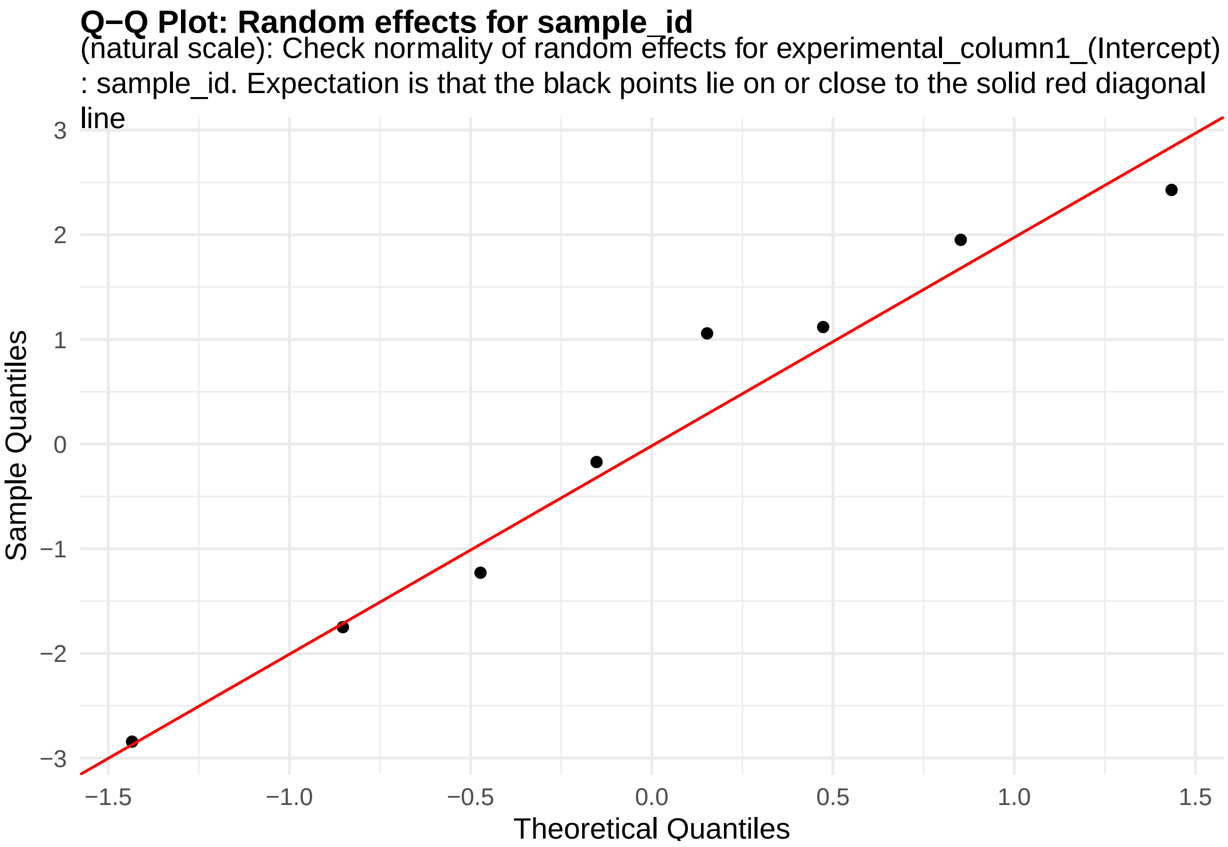


**Figure 22:** random_effects_QQ_1. Check normality of random effects for experimental_column1_(Intercept) (natural scale): sample_id. Expectation is that the black points lie on or close to the solid red diagonal line. Check the Normal Q-Q plots for good and bad examples here: https://library.virginia.edu/data/articles/diagnostic-plots

```
cooks_plots <- diagnose_res$cooks_plots$natural_scale
cat("\n\n\n")
```

```
for(i in 1:length(cooks_plots)) {
  print(cooks_plots[[i]])
  cat(sprintf("\n\n\n**Figure %d:**\n\n", plot_no))
  plot_no <- plot_no + 1
}
```

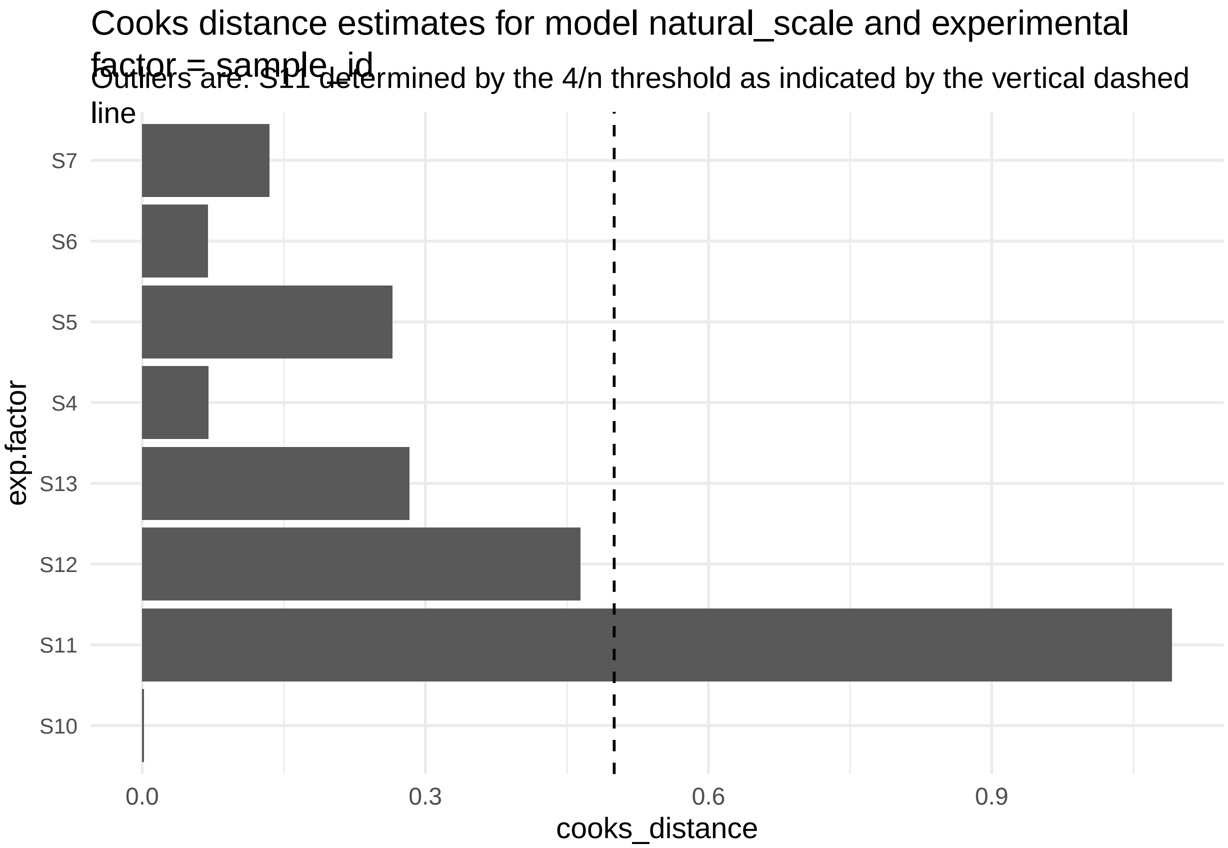


**Figure 23:**

```
print("Inferred outlier groups are")
```

[1] "Inferred outlier groups are"

```
print(diagnose_res$inferred_outlier_groups$natural_scale)
```

$sample_id [1] "S11"

We can now estimate differences in the log odds of cluster membership in cluster 7 between APOE4-KI mice and APOE4-KI-Syn1-Cre using the function 'getEstimatesOfInterest'.

```
estimate_res <- getEstimatesOfInterest(data, design, model, print_plots = F)
print(estimate_res[[1]])
```

```
## Generalized linear mixed model fit by maximum likelihood (Laplace
##   Approximation) [glmerMod]
##  Family: binomial  ( logit )
## Formula:
## cbind(response_column, (total_column - response_column)) ~ condition_column +
##     covariate + (1 | experimental_column1)
##    Data: fixed_global_variable_data
##
##       AIC       BIC    logLik -2*log(L)  df.resid
##     100.9     101.2     -46.4      92.9         4
##
## Scaled residuals:
##      Min       1Q   Median       3Q      Max
## -0.41163 -0.28356  0.00219  0.02755  0.09292
```

```
##
## Random effects:
##  Groups               Name        Variance Std.Dev.
##  experimental_column1 (Intercept) 3.464    1.861
## Number of obs: 8, groups:  experimental_column1, 8
##
## Fixed effects:
##                                  Estimate Std. Error z value Pr(>|z|)
## (Intercept)                       -3.0030     1.1493  -2.613  0.00898 **
## condition_columnPS19-fE4_Syn1-Cre -3.6302     1.3514  -2.686  0.00723 **
## covariateM                        -0.7104     1.3482  -0.527  0.59823
## ---
## Signif. codes:  0 '***' 0.001 '**' 0.01 '*' 0.05 '.' 0.1 ' ' 1
##
## Correlation of Fixed Effects:
##             (Intr) c_PS19
## c_PS19-E4_S -0.560
## covariateM  -0.582 -0.003
```

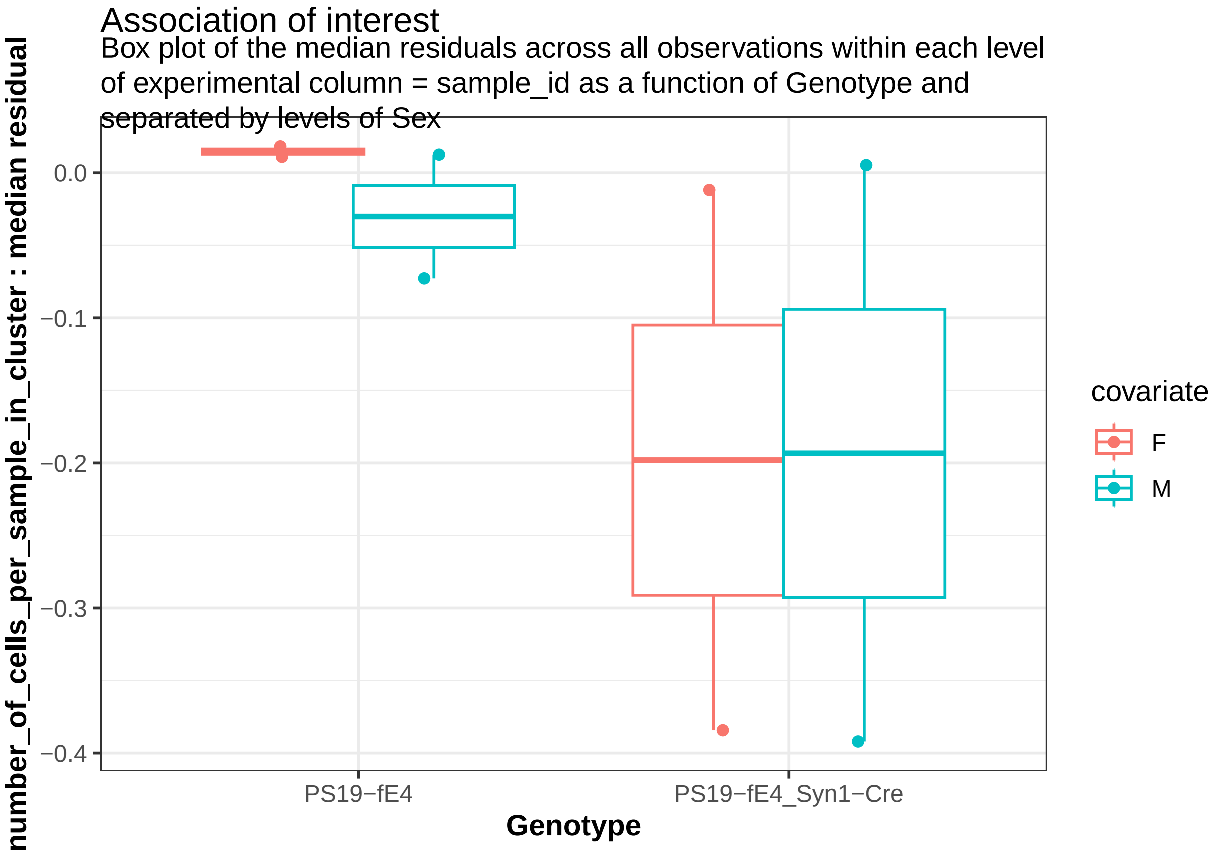


**Figure 24:**

And estimate whether or not statistical power we have to estimate the given observed association with varying number of mice per group.

```
print(power_param@target_columns)
```

```
## [1] "sample_id"
```

```r
power_res <- calculatePower(data, design, model, power_param)
```

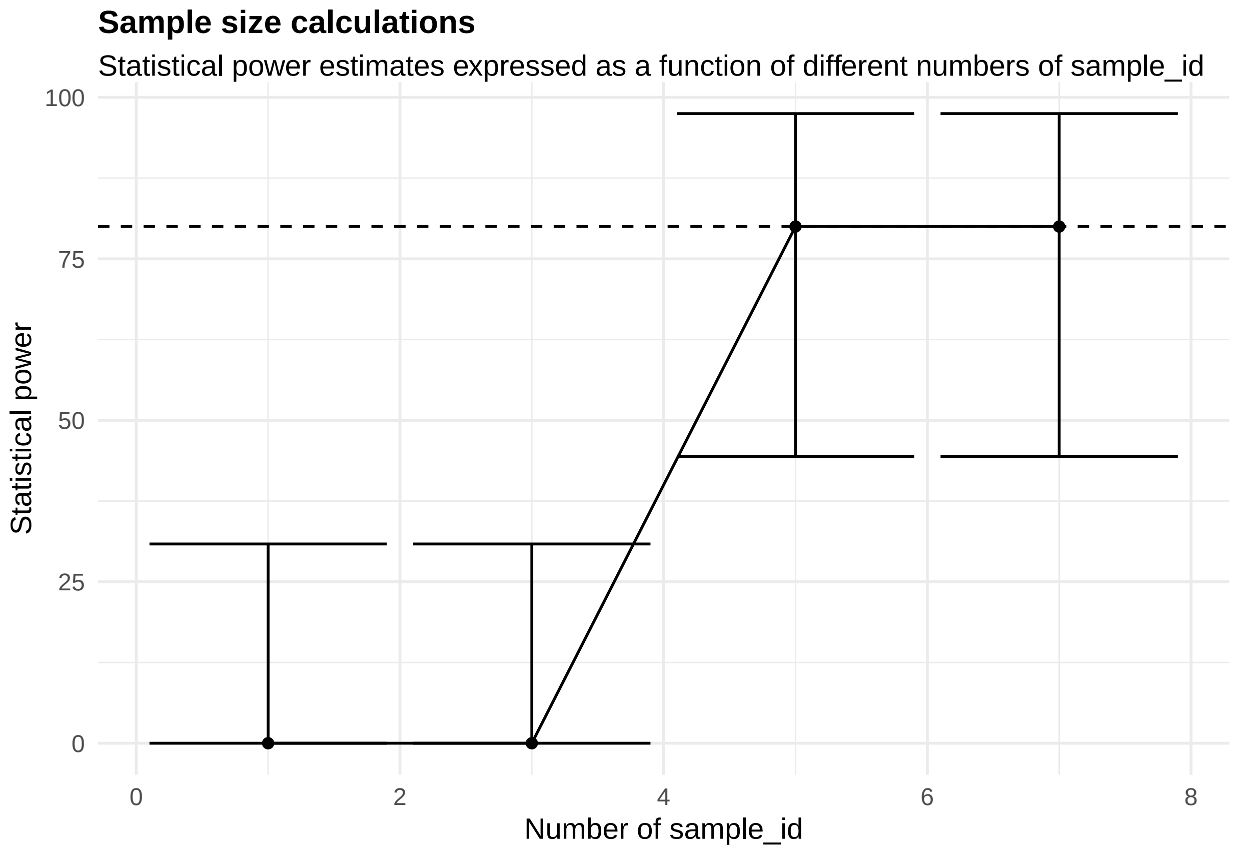


[[1]]

**Figure 25:**

Alternatively, we may believe that effect of genotype on the cluster membership may be cluster specific. Therefore we would include an interaction term in the model

```r
design@include_interaction <- TRUE
```

The estimates after including the interaction term could be recovered by,

```r
estimate_res_w_interaction <- getEstimatesOfInterest(data, design, model,print_plots = F)
print(estimate_res_w_interaction[[1]])
```

```
## Generalized linear mixed model fit by maximum likelihood (Laplace
##   Approximation) [glmerMod]
##  Family: binomial  ( logit )
## Formula:
## cbind(response_column, (total_column - response_column)) ~ condition_column *
##     covariate + (1 | experimental_column1)
##    Data: fixed_global_variable_data
##
##      AIC       BIC    logLik -2*log(L)  df.resid
##    102.3     102.7     -46.2      92.3         3
##
## Scaled residuals:
##      Min       1Q   Median       3Q      Max
## -0.42015 -0.27409 -0.00398  0.03400  0.10782
##
```

```
## Random effects:
##  Groups               Name        Variance Std.Dev.
##  experimental_column1 (Intercept) 3.202    1.789
## Number of obs: 8, groups:  experimental_column1, 8
##
## Fixed effects:
##                                              Estimate Std. Error z value
## (Intercept)                                    -2.530      1.266  -1.999
## condition_columnPS19-fE4_Syn1-Cre              -4.615      1.838  -2.511
## covariateM                                     -1.660      1.798  -0.923
## condition_columnPS19-fE4_Syn1-Cre:covariateM    2.003      2.598   0.771
##                                              Pr(>|z|)
## (Intercept)                                    0.0456 *
## condition_columnPS19-fE4_Syn1-Cre              0.0120 *
## covariateM                                     0.3558
## condition_columnPS19-fE4_Syn1-Cre:covariateM   0.4406
## ---
## Signif. codes:  0 '***' 0.001 '**' 0.01 '*' 0.05 '.' 0.1 ' ' 1
##
## Correlation of Fixed Effects:
##                  (Intr) cn_PS19-E4_S1-C covrtM
## cn_PS19-E4_S1-C -0.689
## covariateM      -0.704  0.486
## c_PS19-E4_S1-C:  0.487 -0.706          -0.692
```

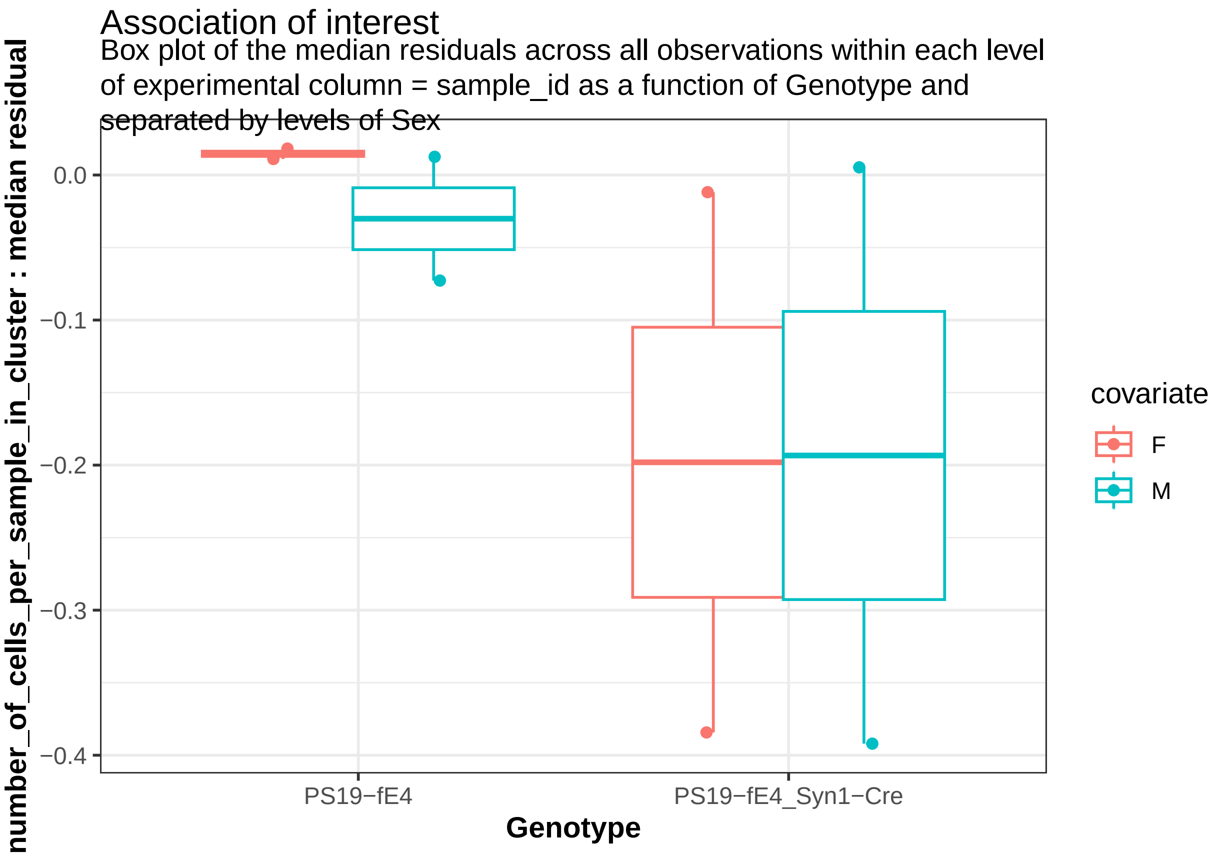


**Figure 26:** We don't see a significant interaction of genotype with sex given that the interaction term is not significantly different from zero (p-value = 0.446)

###### 2.2.1.2 Associations with hippocampus volume

Next, we can first ask the question whether or not the proportion of nuclei derived from each of the mice are associated with their hippocampus volumes. Note, more neuro-degeneration is linked to smaller hippocampus volumes. More exactly, we want to test whether or not there is a change in the log odds of cluster membership in cluster 7 corresponding to a unit change in the hippocampus volume. The above data provided includes the hippocampus volume for each mouse.

```
##Associations of cell-type proportions with hippocampus volume
template_dir <- system.file("input_templates/single_cell_data/celltype_proportions_with_hippocampus_vol

design <- readDesign(file.path(template_dir, "design_celltype.json"))

diagnose_res <- diagnoseDataModel(data, design, model)
```

```
## [1] "Not enough levels to perform cooks test for experimental_column1 with the assumed binomial dist
```

```
## [1] "No outlier detected from the raw Data"
## [1] "Not enough levels to perform cooks test for experimental_column1 with the assumed binomial dist
```

To identify all the models fit to the data. There were outliers identified so two models were fit with and without outliers.

```
print(diagnose_res$models)
```

```
## [1] "Natural scale"              "Natural scale wo outliers"
```

Let us look the basic QC plots for the models fit in the natural scale of the responses. There are only two levels for the animal model. So cooks distance estimates not generated for them.

```
plots <- diagnose_res$diagnostic_plots$natural_scale$plots
captions <- diagnose_res$diagnostic_plots$natural_scale$captions
for (i in seq_along(plots)) {
  print(plots[[i]])
    cat(sprintf("\n\n\n**Figure %d:** %s. %s\n\n", plot_no,names(plots)[i], captions[i]))
  plot_no <- plot_no + 1
}
```

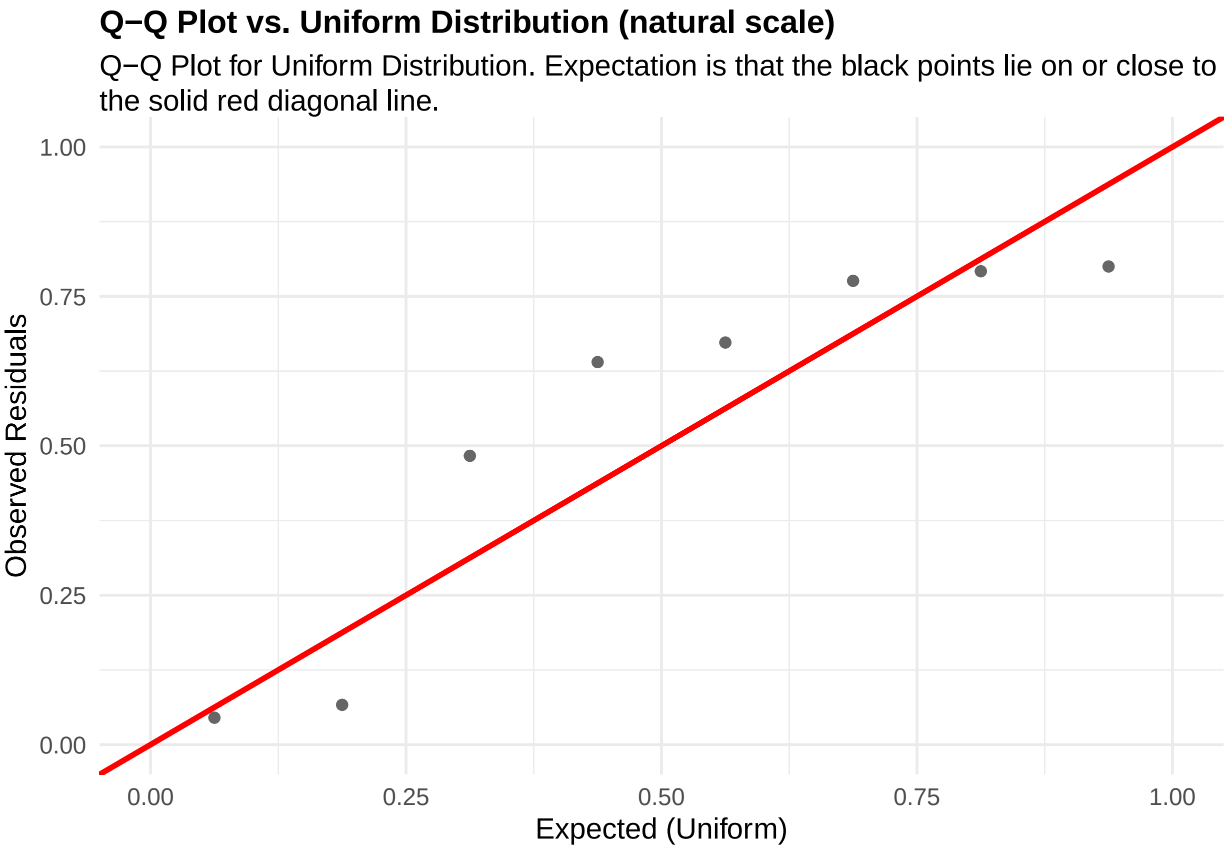


**Figure 27:** residuals_QQ. Q-Q Plot for Uniform Distribution. Expectation is that the black points lie on or close to the solid red diagonal line. Check the Q-Q plots for good and bad examples here: https://library.virginia.edu/data/articles/diagnostic-plots

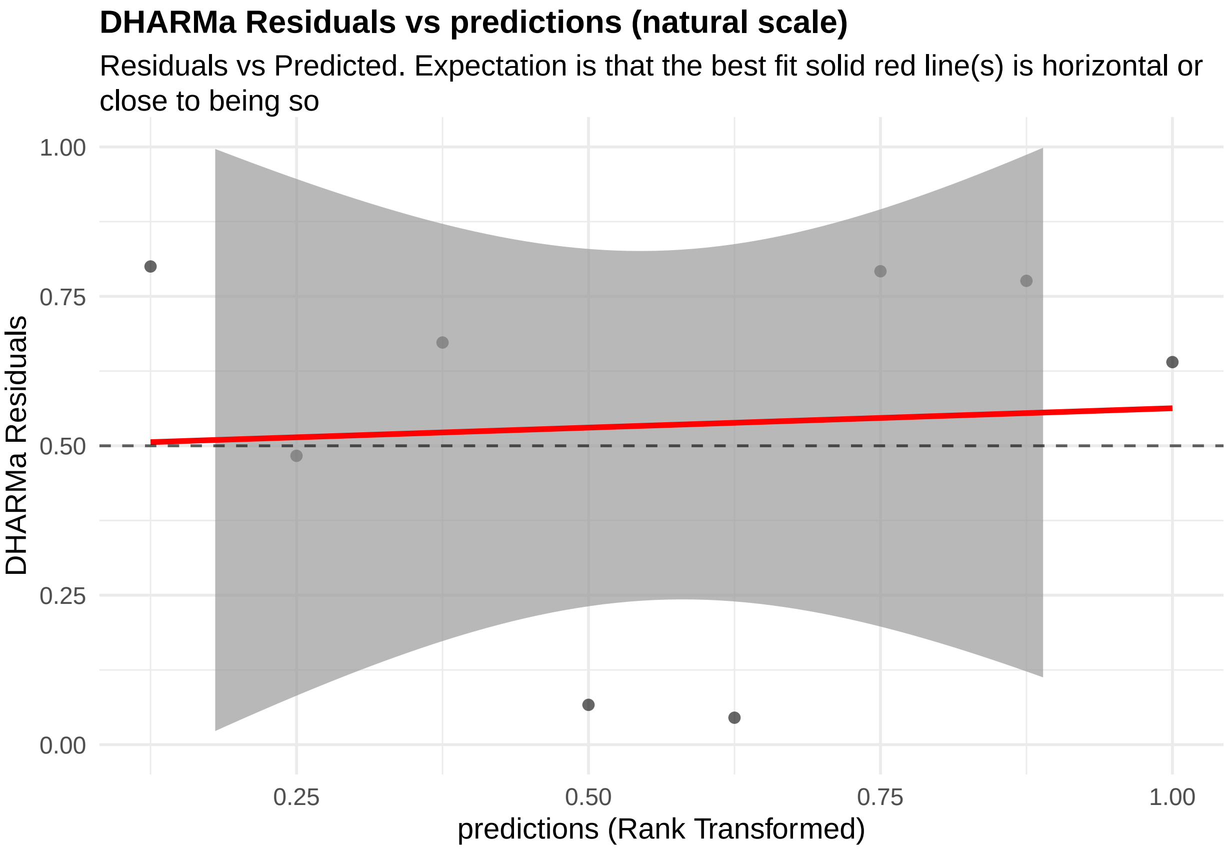


**Figure 28:** residuals_vs_predicted. Residuals vs Predicted Check the Residuals vs Fitted plots for good and bad examples here: https://library.virginia.edu/data/articles/diagnostic-plots

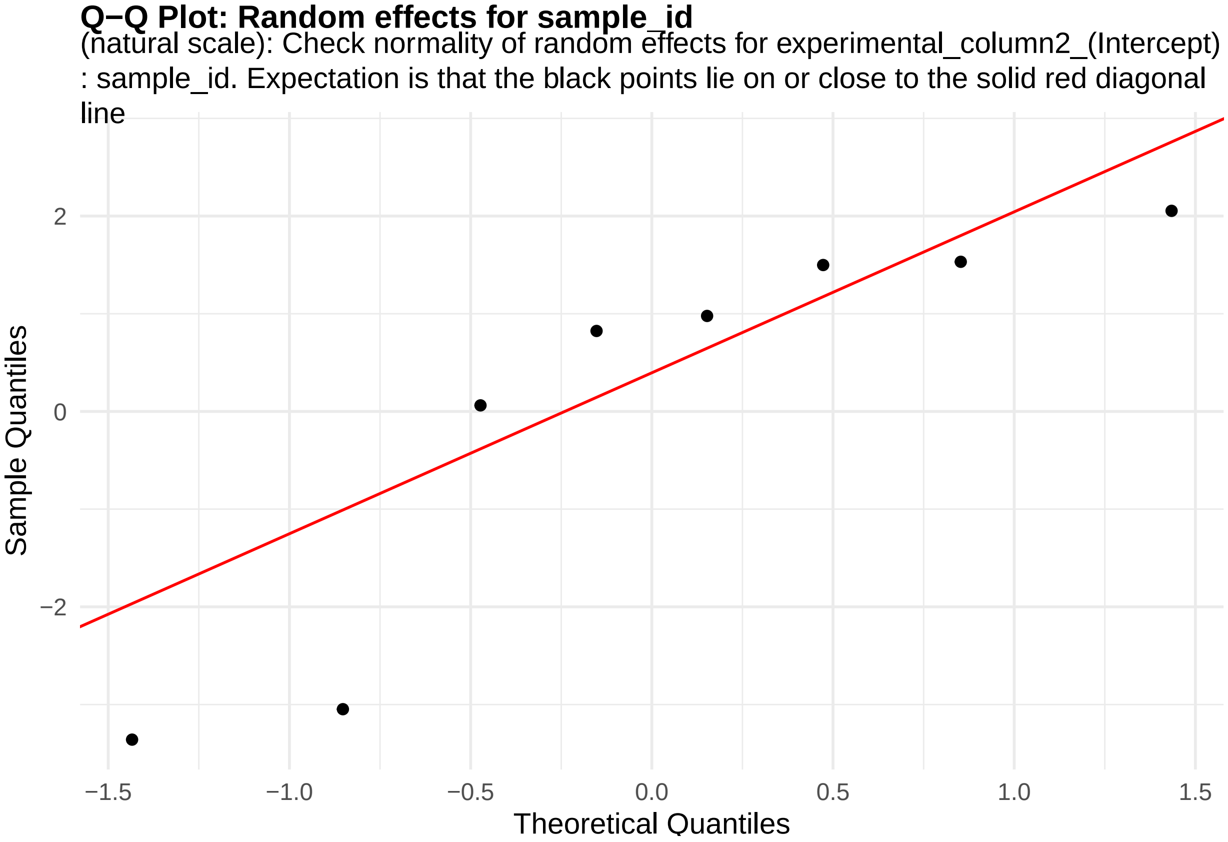


**Figure 29:** random_effects_QQ_1. Check normality of random effects for experimental_column2_(Intercept) (natural scale): sample_id. Expectation is that the black points lie on or close to the solid red diagonal line. Check the Normal Q-Q plots for good and bad examples here: https://library.virginia.edu/data/articles/diagnostic-plots

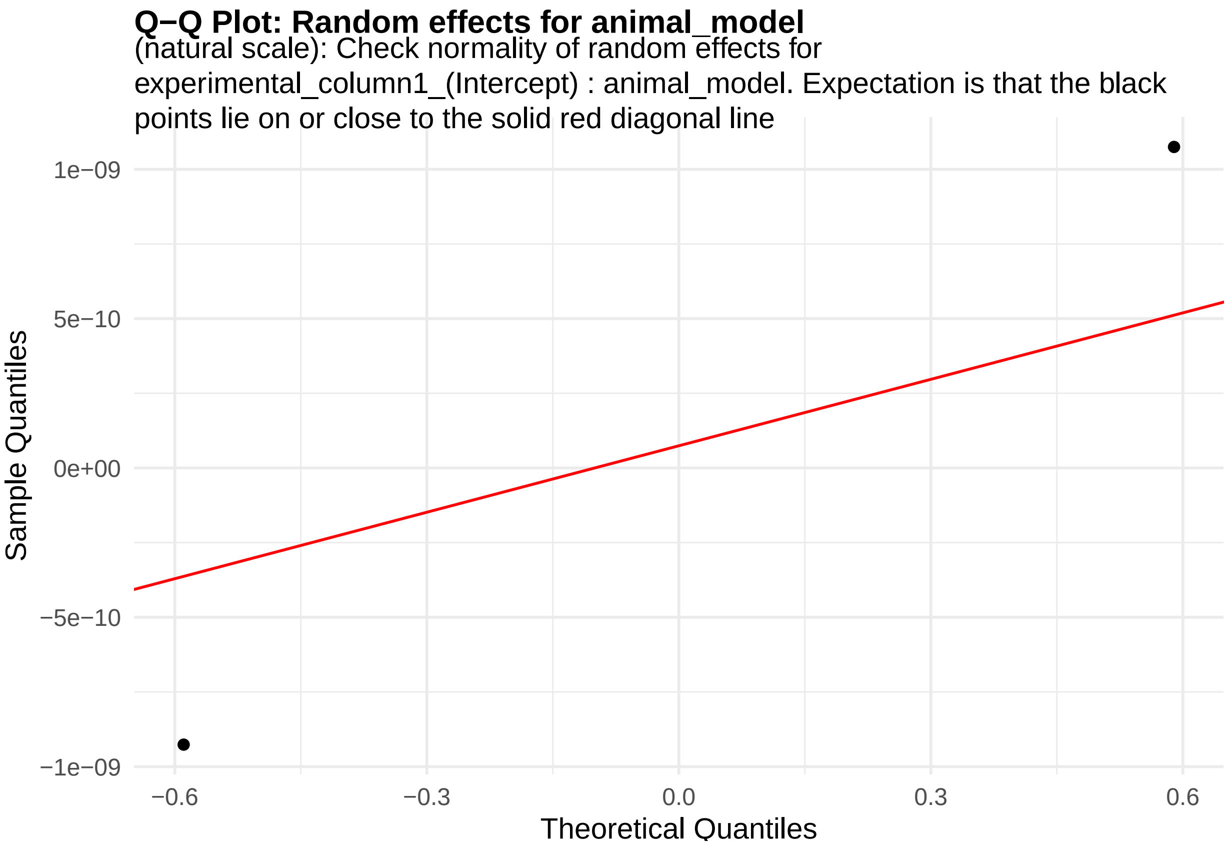


**Figure 30:** random_effects_QQ_2. Check normality of random effects for experimental_column1_(Intercept) (natural scale): animal_model. Expectation is that the black points lie on or close to the solid red diagonal line. Check the Normal Q-Q plots for good and bad examples here: https://library.virginia.edu/data/articles/diagnostic-plots

#### Histogram of Residuals Values

Histogram of residuals values. Red vertical lines (if present) are the location of the cutoffs to identify the outliers. No red vertical lines implies that the residual values that were estimated to be infinite were identified as outliers

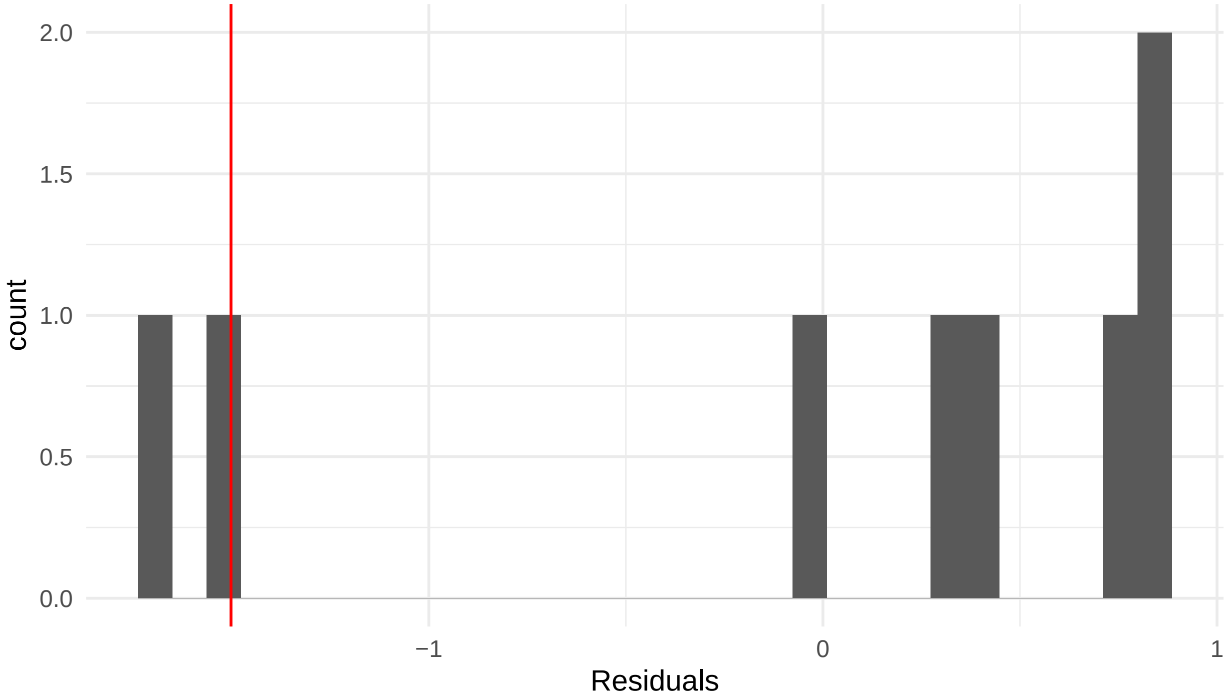


**Figure 31:** residuals_histogram. Histogram of residuals values. Red vertical lines (if present) are the location of the cutoffs to identify the outliers. No red vertical lines implies that the residual values that were estimated to be infinite were identified as outliers

```
cat("\n\n\n")
```

Let us also now look at the QC plots after removing the identified outliers

```
plots <- diagnose_res$diagnostic_plots$natural_scale_wo_outliers$plots
captions <- diagnose_res$diagnostic_plots$natural_scale_wo_outliers$captions

for (i in seq_along(plots)) {
  print(plots[[i]])
    cat(sprintf("\n\n\n**Figure %d:** %s. %s\n\n", plot_no,names(plots)[i], captions[i]))
  plot_no <- plot_no + 1
}
```

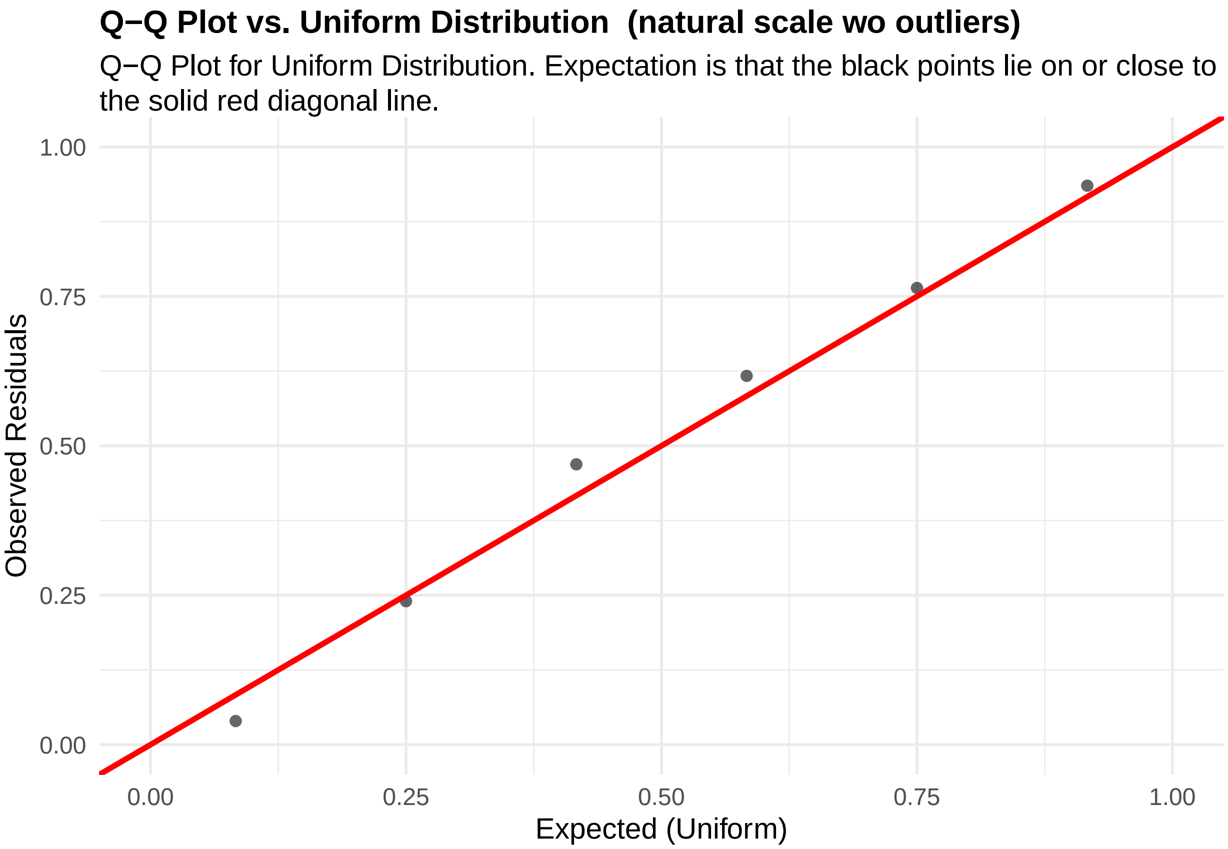


**Figure 32:** residuals_QQ. Q-Q Plot for Uniform Distribution. Expectation is that the black points lie on or close to the solid red diagonal line. Check the Q-Q plots for good and bad examples here: https://library.virginia.edu/data/articles/diagnostic-plots

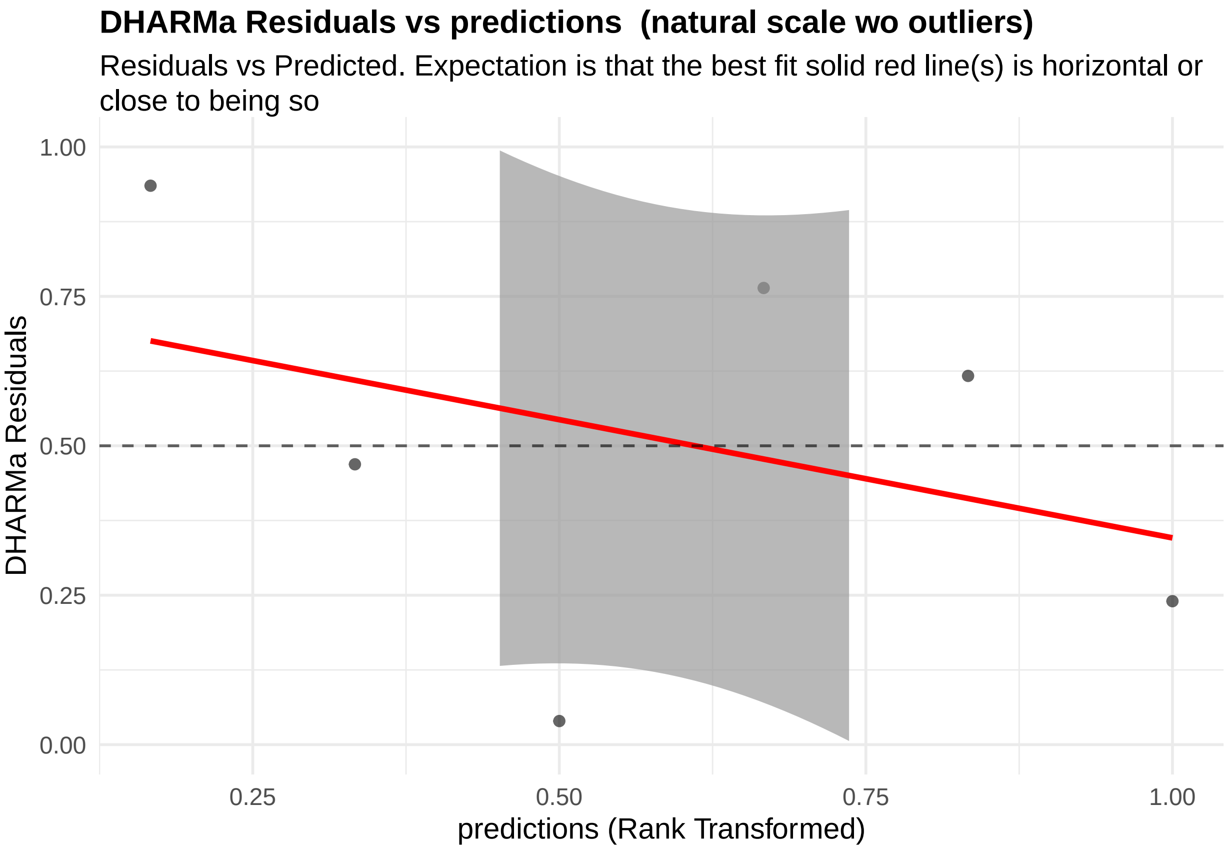


**Figure 33:** residuals_vs_predicted. Residuals vs Predicted Check the Residuals vs Fitted plots for good and bad examples here: https://library.virginia.edu/data/articles/diagnostic-plots

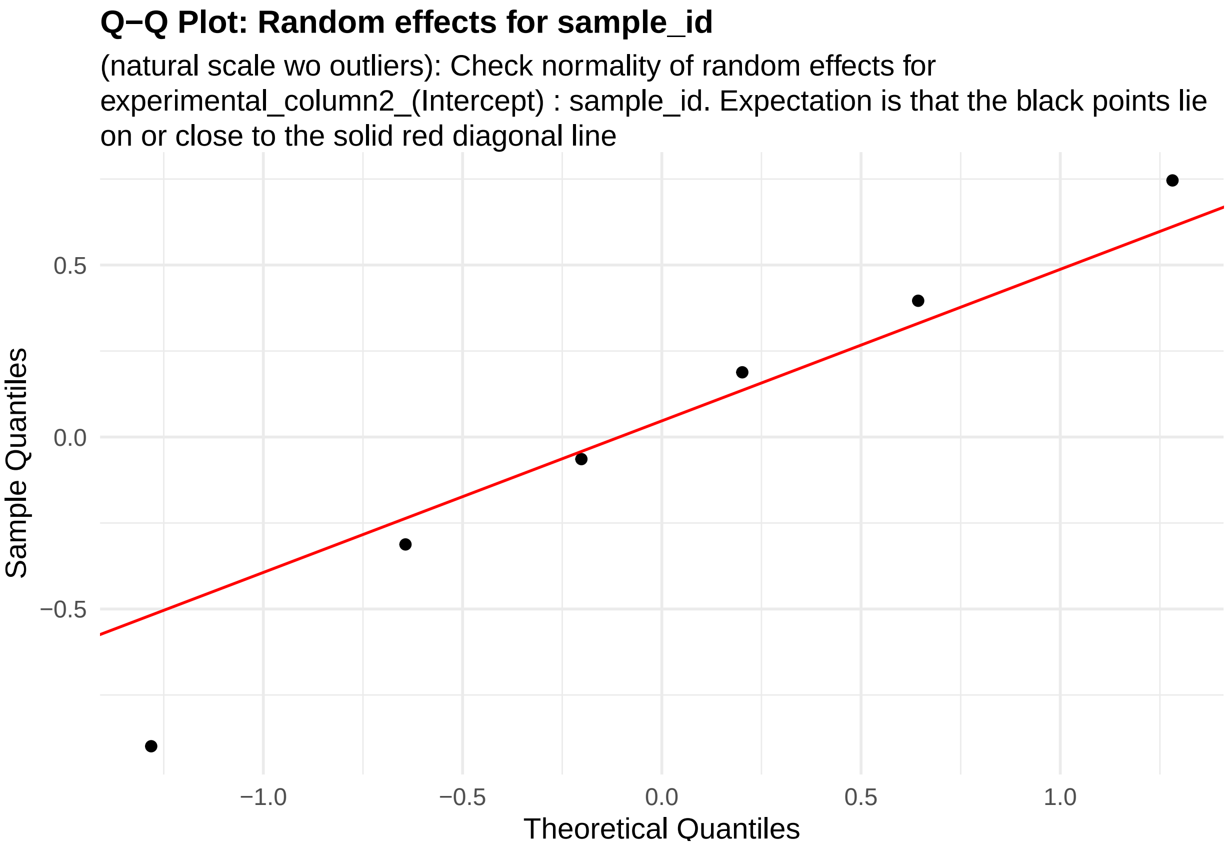


**Figure 34:** random_effects_QQ_1. Check normality of random effects for experimental_column2_(Intercept) (natural scale wo outliers): sample_id. Expectation is that the black points lie on or close to the solid red diagonal line. Check the Normal Q-Q plots for good and bad examples here: https://library.virginia.edu/data/articles/diagnostic-plots

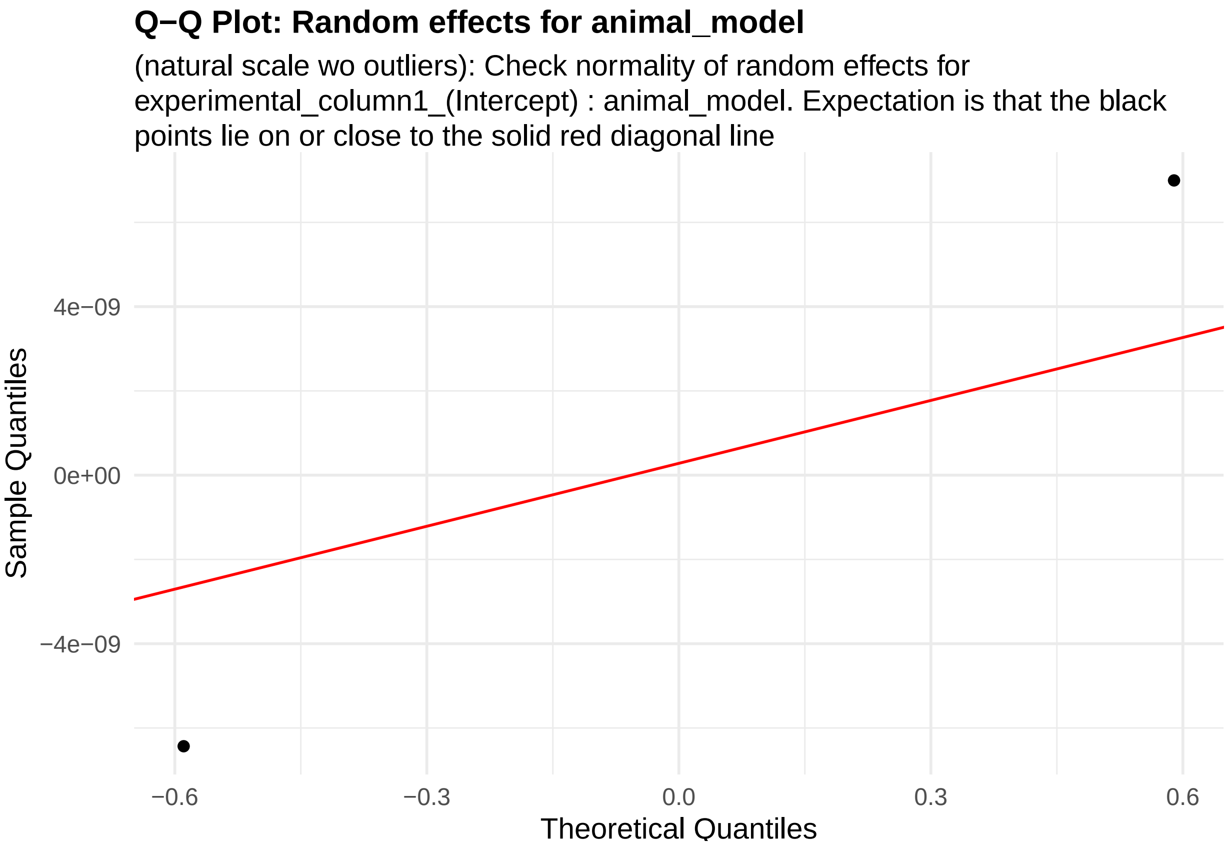


**Figure 35:** random_effects_QQ_2. Check normality of random effects for experimental_column1_(Intercept) (natural scale wo outliers): animal_model. Expectation is that the black points lie on or close to the solid red diagonal line. Check the Normal Q-Q plots for good and bad examples here: https://library.virginia.edu/data/articles/diagnostic-plots

```
cat("\n\n\n")
```

We can also estimate the parameter of interest - log odds ratio of cluster 7 membership per unit increase in hippocampus volume of associated mouse.

```
estimate_res <- getEstimatesOfInterest(data, design, model,print_plots = F)
print(estimate_res[[1]])
```

```
## Generalized linear mixed model fit by maximum likelihood (Laplace
##   Approximation) [glmerMod]
##  Family: binomial  ( logit )
## Formula:
## cbind(response_column, (total_column - response_column)) ~ condition_column +
##     covariate + (1 | experimental_column1) + (1 | experimental_column2)
##    Data: fixed_global_variable_data
##
##      AIC      BIC   logLik -2*log(L)  df.resid
##    105.2    105.6    -47.6     95.2         3
##
## Scaled residuals:
##      Min       1Q   Median       3Q      Max
## -0.57752 -0.13043  0.00696  0.02846  0.06210
##
## Random effects:
```

```
##  Groups                Name        Variance  Std.Dev.
##  experimental_column2 (Intercept) 4.375e+00 2.092e+00
##  experimental_column1 (Intercept) 1.201e-09 3.465e-05
## Number of obs: 8, groups:  experimental_column2, 8; experimental_column1, 2
##
## Fixed effects:
##                  Estimate Std. Error z value Pr(>|z|)
## (Intercept)       -0.2808     2.5591  -0.110   0.9126
## condition_column  -0.6509     0.3375  -1.929   0.0537 .
## covariateM        -0.6567     1.5052  -0.436   0.6626
## ---
## Signif. codes:  0 '***' 0.001 '**' 0.01 '*' 0.05 '.' 0.1 ' ' 1
##
## Correlation of Fixed Effects:
##             (Intr) cndtn_
## condtn_clmn -0.909
## covariateM  -0.274 -0.021
## optimizer (Nelder_Mead) convergence code: 0 (OK)
## boundary (singular) fit: see help('isSingular')
```

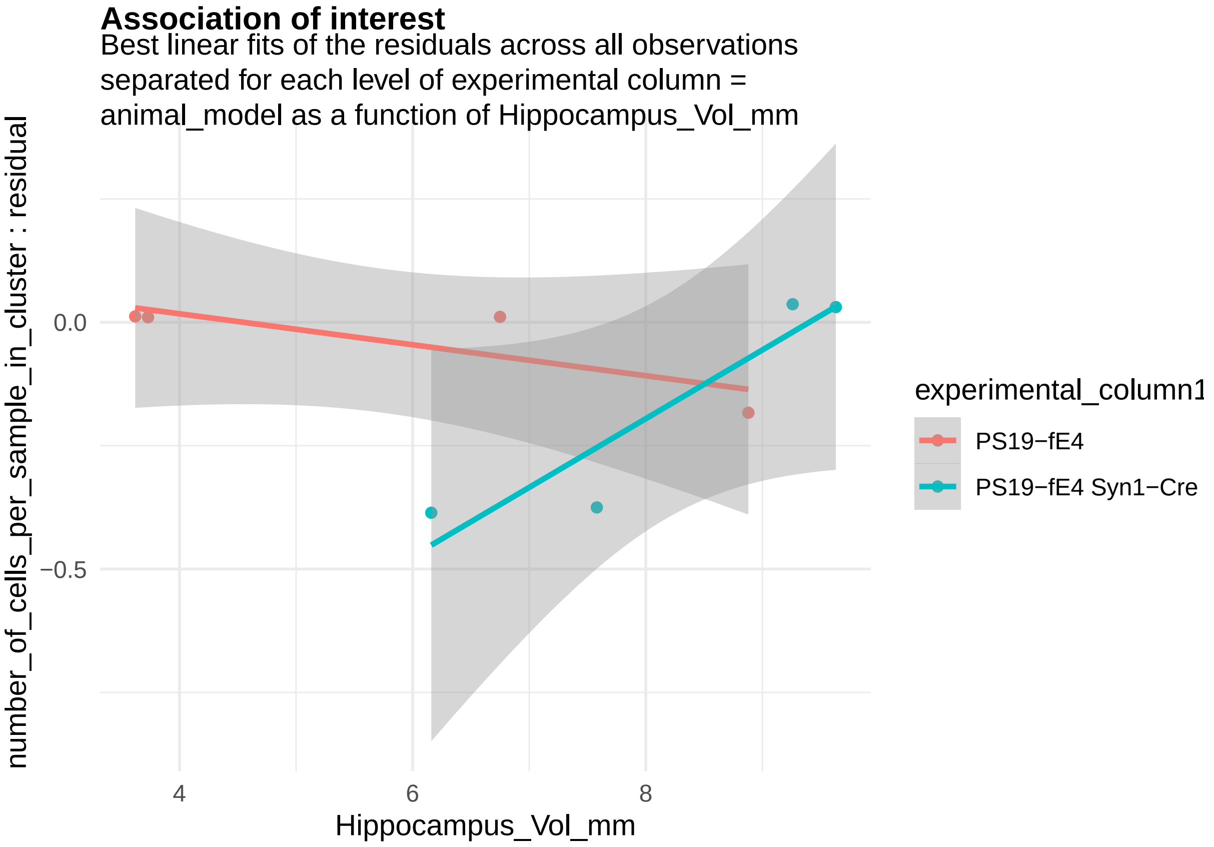


**Figure 36:**

And calculate the statistical power to estimate this association for different numbers of animals per group.

```
power_plots <- calculatePower(data, design, model, power_param)
```

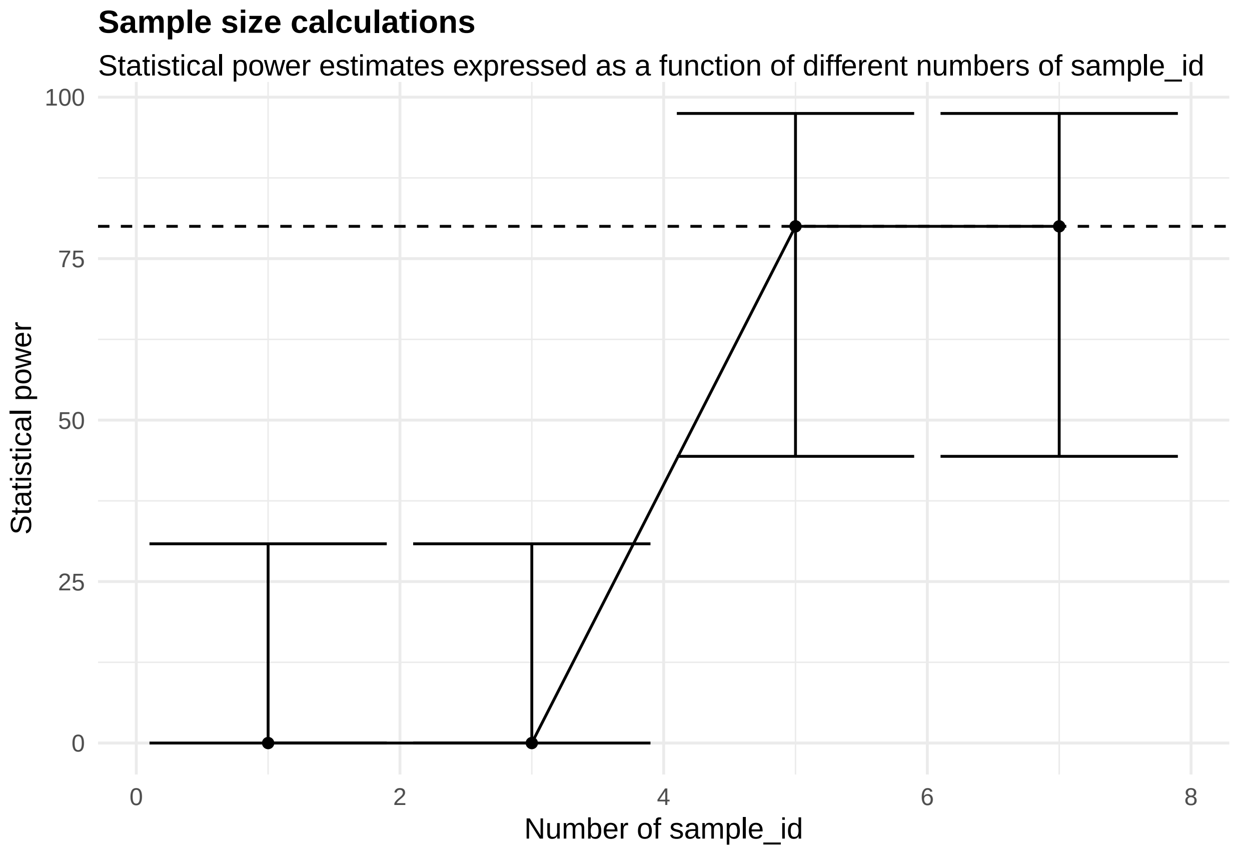


[[1]]

**Figure 37:**

##### 2.2.2 Gene expression associations with genotype

We will now estimate the association of the expression levels of the Snhg11 gene with genotype among cells in cluster 7 in the same data set. We will now load the raw counts for this gene and create the relevant RMeDesign, ProbabilityModel and PowerParams objects.

```
template_dir <- system.file("input_templates/single_cell_data/gene_expression_with_genotype", package =

data <- snRNAseq_gene_count_data
head(data)
```

```
##    y    orig.ident nCount_RNA nFeature_RNA percent.mt sample_number
## 1 20 SeuratProject       2878         1716 0.03474635           S10
## 2  6 SeuratProject       9350         3682 0.05347594           S10
## 3 22 SeuratProject       2949         1735 0.03390980           S10
## 4 12 SeuratProject       4529         2494 0.04415986           S10
## 5 14 SeuratProject       3044         1866 0.03285151           S10
## 6  9 SeuratProject      10636         3957 0.03760812           S10
##   mouse_number genotype sex Cre ApoE      dob age_perfused date_perfused
## 1          154 PS19-fE4   F   - E4/4 10/30/19         10.3        9/7/20
## 2          154 PS19-fE4   F   - E4/4 10/30/19         10.3        9/7/20
## 3          154 PS19-fE4   F   - E4/4 10/30/19         10.3        9/7/20
## 4          154 PS19-fE4   F   - E4/4 10/30/19         10.3        9/7/20
## 5          154 PS19-fE4   F   - E4/4 10/30/19         10.3        9/7/20
## 6          154 PS19-fE4   F   - E4/4 10/30/19         10.3        9/7/20
##   date_of_nuclear_isolation nCount_SCT nFeature_SCT SCT_snn_res.0.7
```

```
## 1                    9/9/21       2762        1716              6
## 2                    9/9/21       2910        2040              6
## 3                    9/9/21       2813        1735              6
## 4                    9/9/21       3253        2414              6
## 5                    9/9/21       2877        1866              6
## 6                    9/9/21       2675        1843              6
##   seurat_clusters orig_seurat_clusters y.samples.lib.size library_size
## 1               7                    6               2762         2762
## 2               7                    6               2910         2910
## 3               7                    6               2813         2813
## 4               7                    6               3253         3253
## 5               7                    6               2877         2877
## 6               7                    6               2675         2675
##   constant_library_size
## 1                     1
## 2                     1
## 3                     1
## 4                     1
## 5                     1
## 6                     1
```

```
design <- readDesign(file.path(template_dir, "design_gene_expression.json"))
print(design)
```

```
## An object of class "RMeDesign"
## Slot "response_column":
## [1] "y"
##
## Slot "covariate":
## [1] "sex"
##
## Slot "condition_column":
## [1] "genotype"
##
## Slot "condition_is_categorical":
## [1] TRUE
##
## Slot "experimental_columns":
## [1] "sample_number"
##
## Slot "crossed_columns":
## NULL
##
## Slot "total_column":
## [1] "library_size"
##
## Slot "outlier_alpha":
## [1] 0.05
##
## Slot "na_action":
## [1] "complete"
##
## Slot "include_interaction":
## [1] FALSE
##
```

```
## Slot "random_slope_variable":
## NULL
##
## Slot "covariate_is_categorical":
## [1] TRUE
```

```
model <- readProbabilityModel(file.path(template_dir, "prob_model.json"))
print(model)
```

```
## An object of class "ProbabilityModel"
## Slot "error_is_non_normal":
## [1] TRUE
##
## Slot "family_p":
## [1] "negative_binomial"
```

```
power_param <- readPowerParams(file.path(template_dir, "power_param.json"))
print(power_param)
```

```
## An object of class "PowerParams"
## Slot "target_columns":
## [1] "sample_number"
##
## Slot "power_curve":
## [1] 1
##
## Slot "nsimn":
## [1] 10
##
## Slot "levels":
## [1] 1
##
## Slot "max_size":
## [1] 1
##
## Slot "alpha":
## [1] 0.05
##
## Slot "breaks":
## NULL
##
## Slot "effect_size":
## NULL
##
## Slot "icc":
## NULL
```

Note, we will assume the underlying probability distribution for the counts is negative binomial and will control for differences in the sequencing depths or the total reads each cell receives using `library_size`.

Diagnosis of the data and the model

```
diagnose_res <- diagnoseDataModel(data, design, model)
```

To identify all the models fit to the data. There were outliers identified so two models were fit in the natural scale.

```r
print(diagnose_res$models)
```

```
## [1] "Natural scale"              "Natural scale wo outliers"
```

Let us look the basic QC plots for the models fit in the natural scale of the responses.

```r
plots <- diagnose_res$diagnostic_plots$natural_scale$plots
captions <- diagnose_res$diagnostic_plots$natural_scale$captions

for (i in seq_along(plots)) {
  print(plots[[i]])
    cat(sprintf("\n\n\n**Figure %d:** %s. %s\n\n", plot_no,names(plots)[i], captions[i]))
  plot_no <- plot_no + 1
}
```

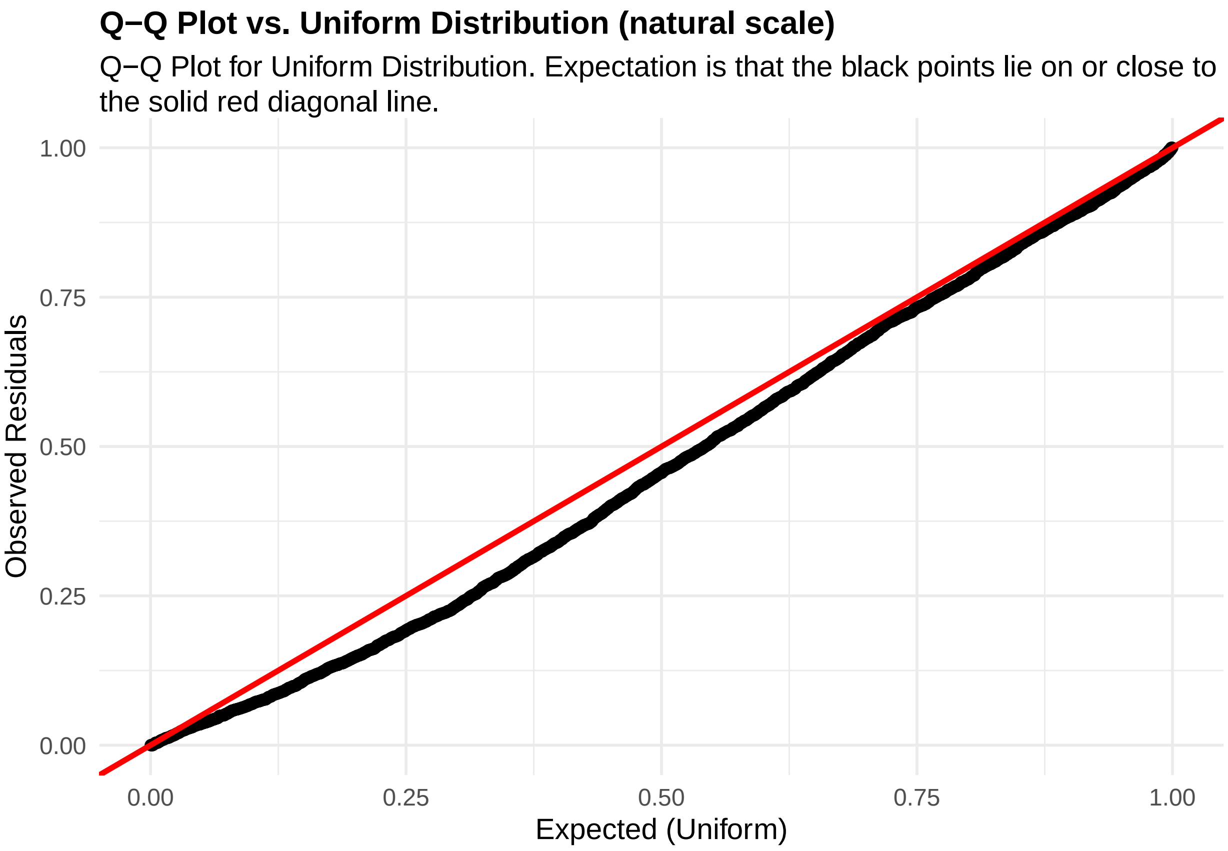


**Figure 38:** residuals_QQ. Q-Q Plot for Uniform Distribution. Expectation is that the black points lie on or close to the solid red diagonal line. Check the Q-Q plots for good and bad examples here: https://library.virginia.edu/data/articles/diagnostic-plots

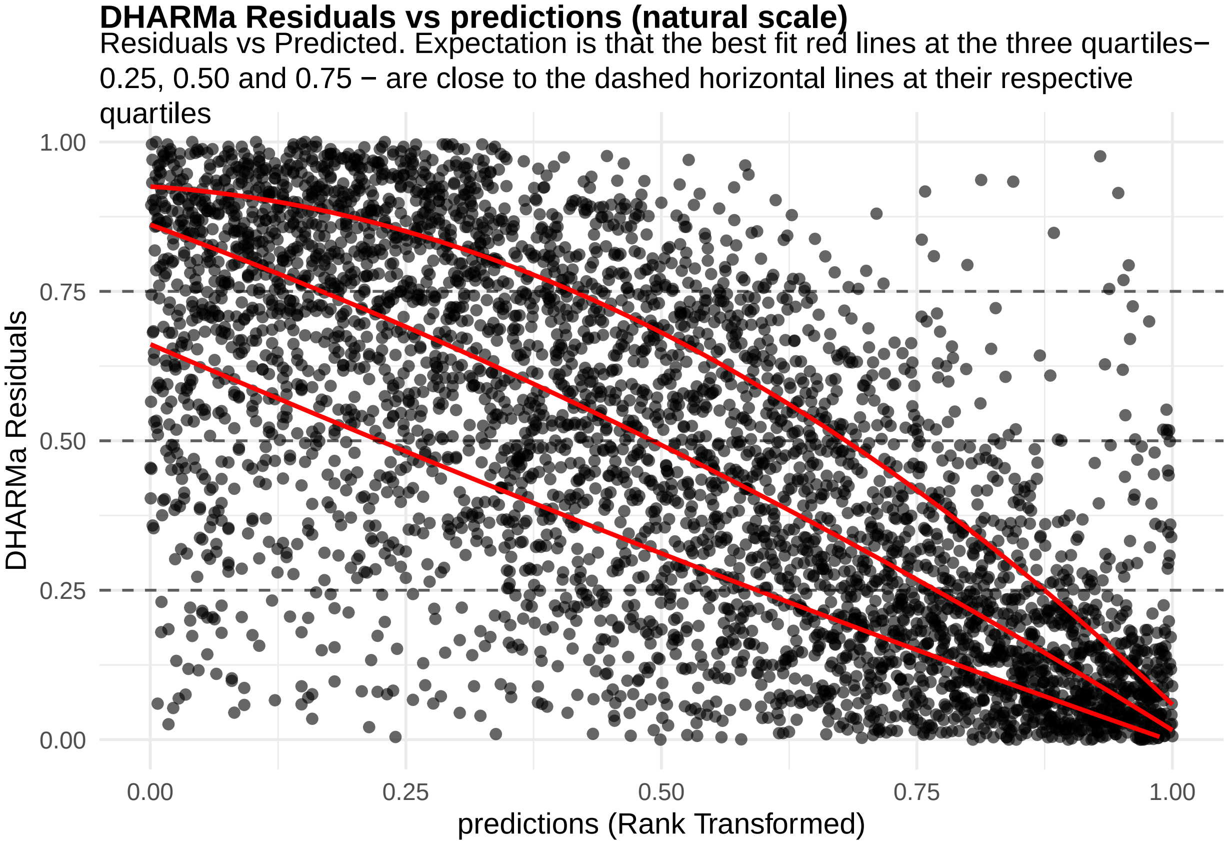


**Figure 39:** residuals_vs_predicted. Residuals vs Predicted Check the Residuals vs Fitted plots for good and bad examples here: https://library.virginia.edu/data/articles/diagnostic-plots

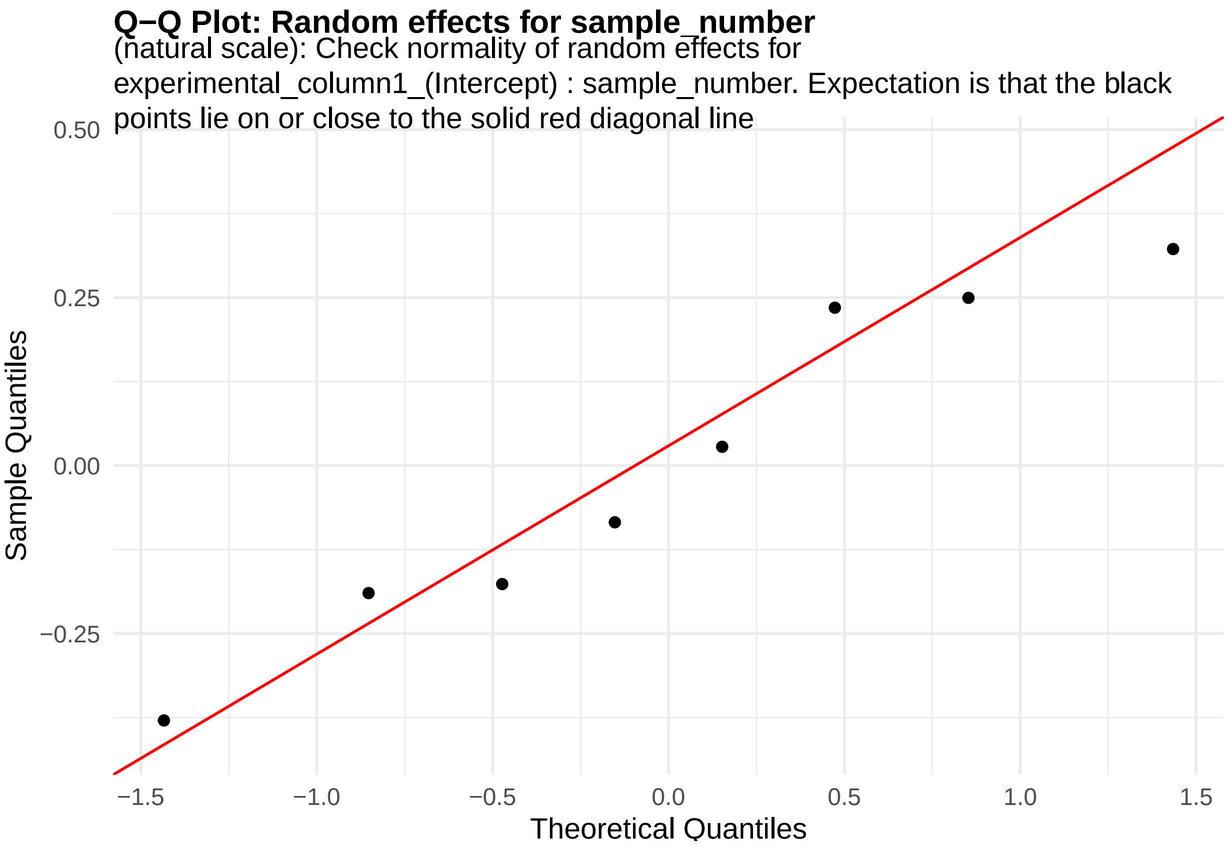


**Figure 40:** random_effects_QQ_1. Check normality of random effects for experimental_column1_(Intercept) (natural scale): sample_number. Expectation is that the black points lie on or close to the solid red diagonal line. Check the Normal Q-Q plots for good and bad examples here: https://library.virginia.edu/data/articles/diagnostic-plots

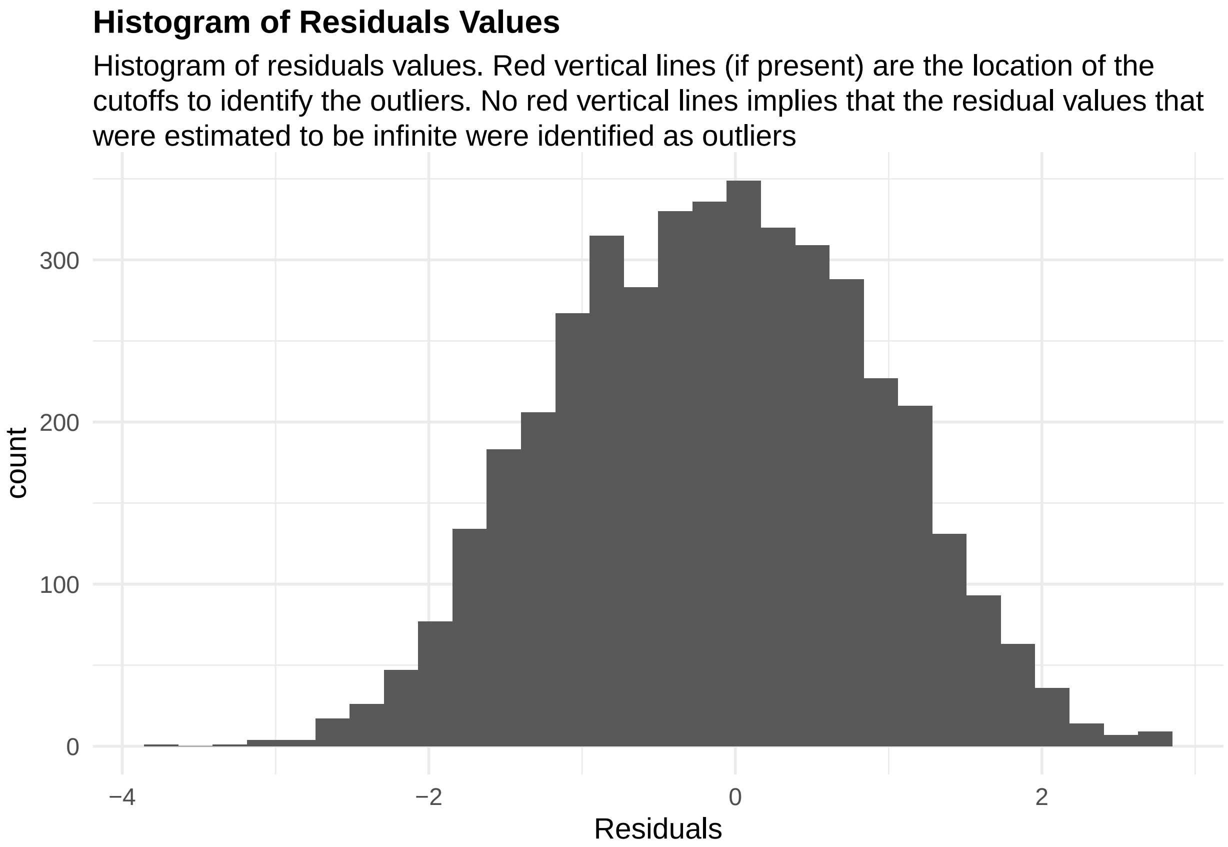


**Figure 41:** residuals_histogram. Histogram of residuals values. Red vertical lines (if present) are the location of the cutoffs to identify the outliers. No red vertical lines implies that the residual values that were estimated to be infinite were identified as outliers

```
cooks_plots <- diagnose_res$cooks_plots$natural_scale
cat("\n\n\n")
```

```
for(i in 1:length(cooks_plots)) {
  print(cooks_plots[[i]])
  cat(sprintf("\n\n\n**Figure %d:**\n\n", plot_no))
  plot_no <- plot_no + 1
}
```

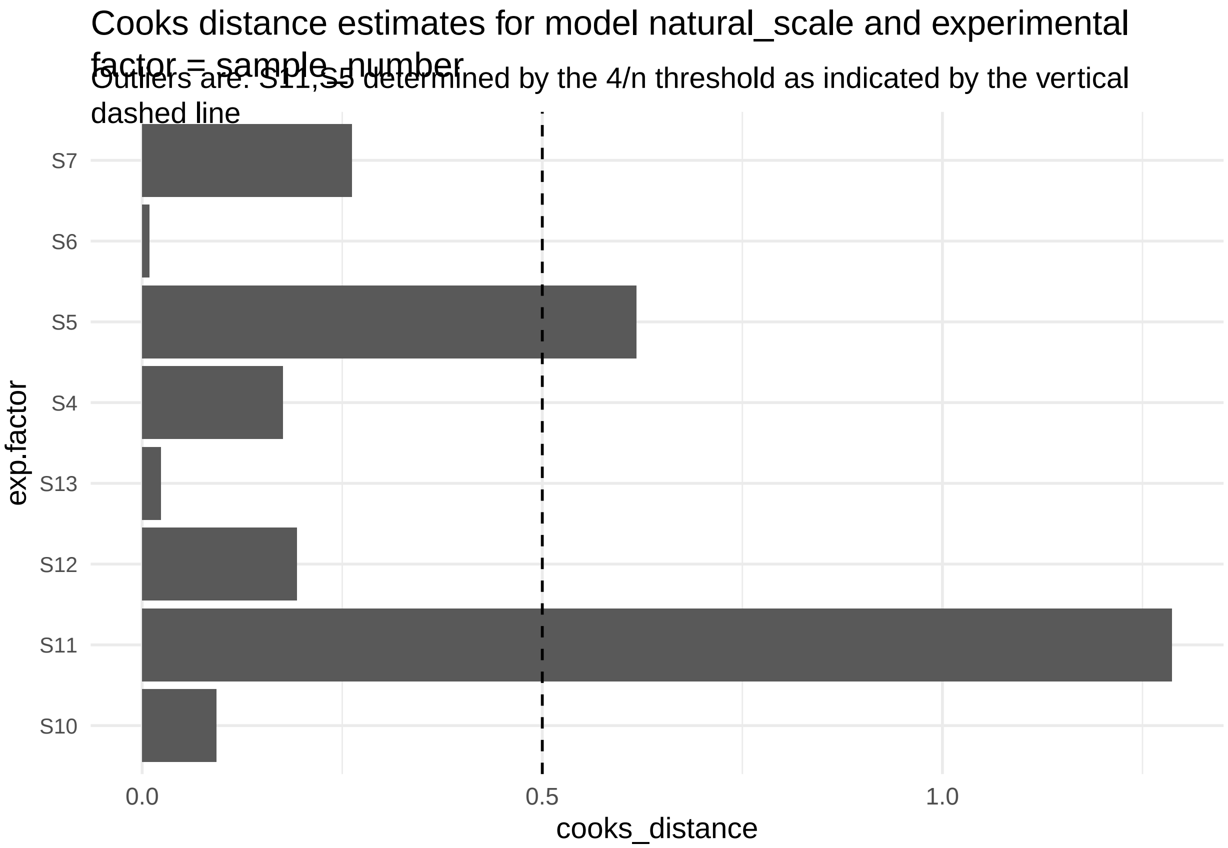


**Figure 42:**

```r
print("Inferred outlier groups are")
```

[1] "Inferred outlier groups are"

```r
print(diagnose_res$inferred_outlier_groups$natural_scale)
```

$sample_number [1] "S11" "S5"

Let us also now look at the QC plots after removing the identified outliers

```r
plots <- diagnose_res$diagnostic_plots$natural_scale_wo_outliers$plots
captions <- diagnose_res$diagnostic_plots$natural_scale_wo_outliers$captions

for (i in seq_along(plots)) {
  print(plots[[i]])
    cat(sprintf("\n\n\n**Figure %d:** %s. %s\n\n", plot_no,names(plots)[i], captions[i]))
  plot_no <- plot_no + 1
}
```

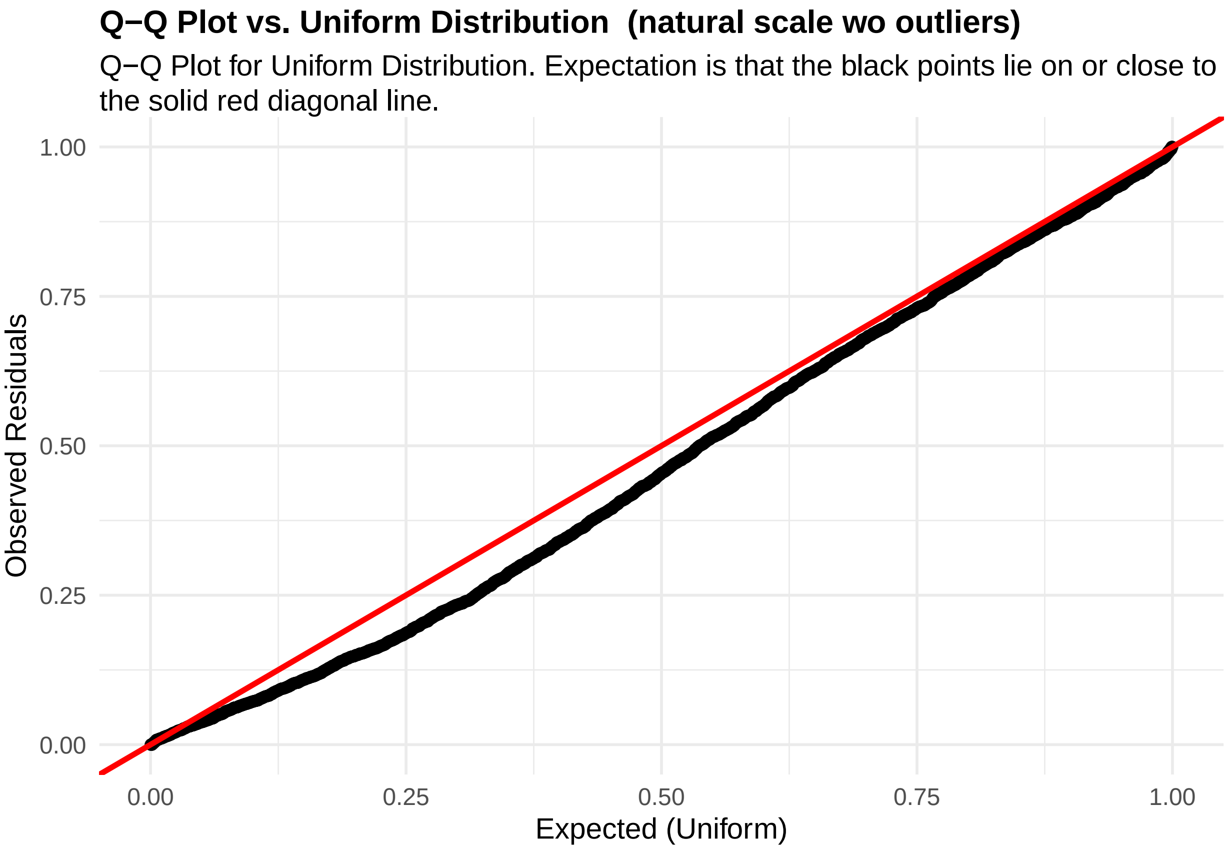


**Figure 43:** residuals_QQ. Q-Q Plot for Uniform Distribution. Expectation is that the black points lie on or close to the solid red diagonal line. Check the Q-Q plots for good and bad examples here: `https://library.virginia.edu/data/articles/diagnostic-plots`

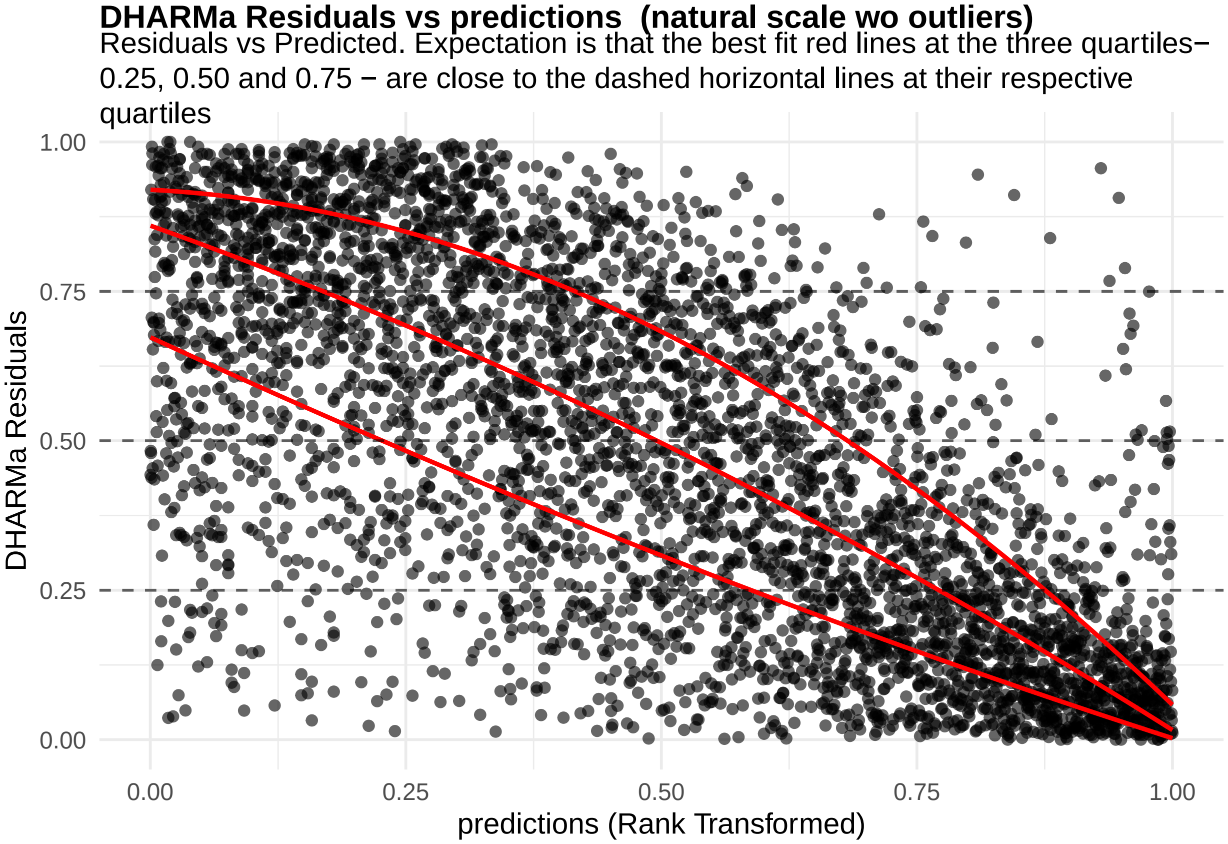


**Figure 44:** residuals_vs_predicted. Residuals vs Predicted Check the Residuals vs Fitted plots for good and bad examples here: https://library.virginia.edu/data/articles/diagnostic-plots

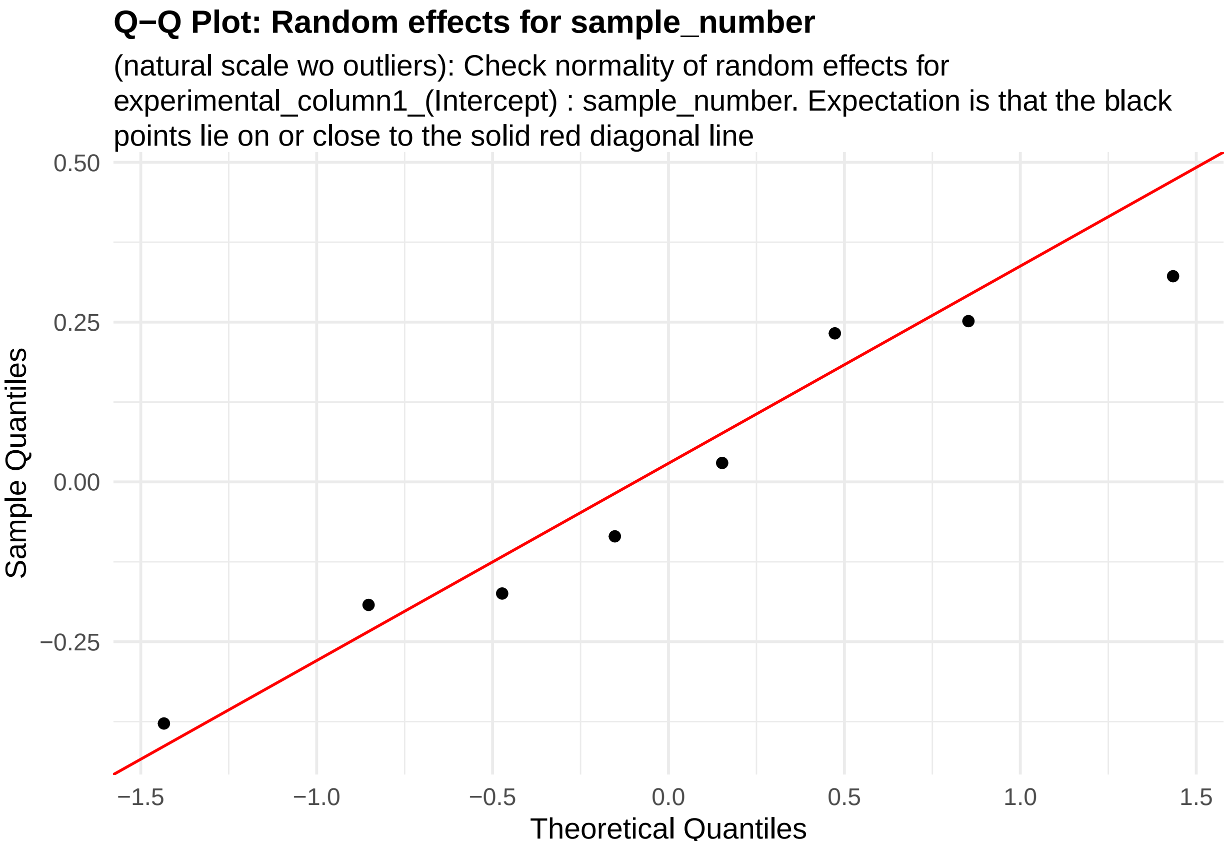


**Figure 45:** random_effects_QQ_1. Check normality of random effects for experimental_column1_(Intercept) (natural scale wo outliers): sample_number. Expectation is that the black points lie on or close to the solid red diagonal line. Check the Normal Q-Q plots for good and bad examples here: https://library.virginia.edu/data/articles/diagnostic-plots

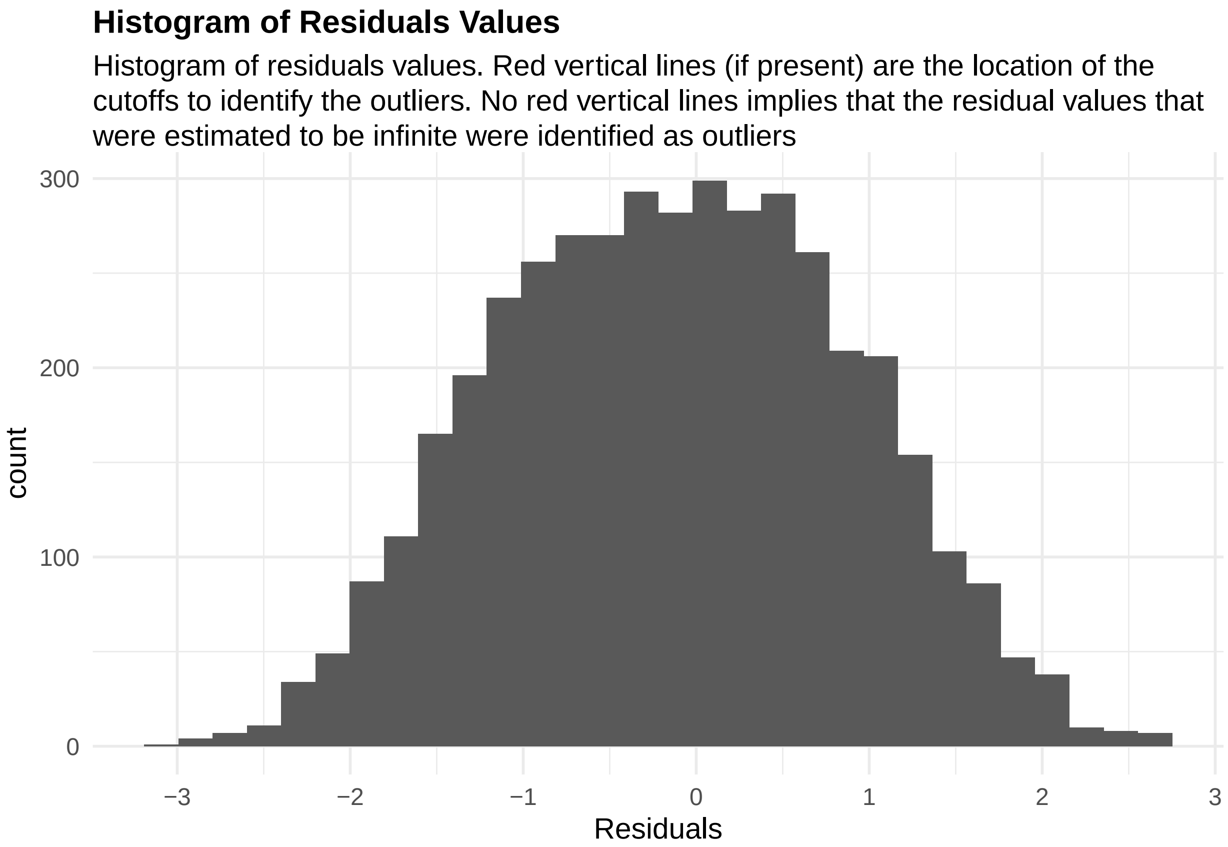


**Figure 46:** residuals_histogram. Histogram of residuals values. Red vertical lines (if present) are the location of the cutoffs to identify the outliers. No red vertical lines implies that the residual values that were estimated to be infinite were identified as outliers

```
cooks_plots <- diagnose_res$cooks_plots$natural_scale_wo_outliers
cat("\n\n\n")
```

```
for(i in 1:length(cooks_plots)) {
  print(cooks_plots[[i]])
  cat(sprintf("\n\n\n**Figure %d:**\n\n", plot_no))
  plot_no <- plot_no + 1
}
```

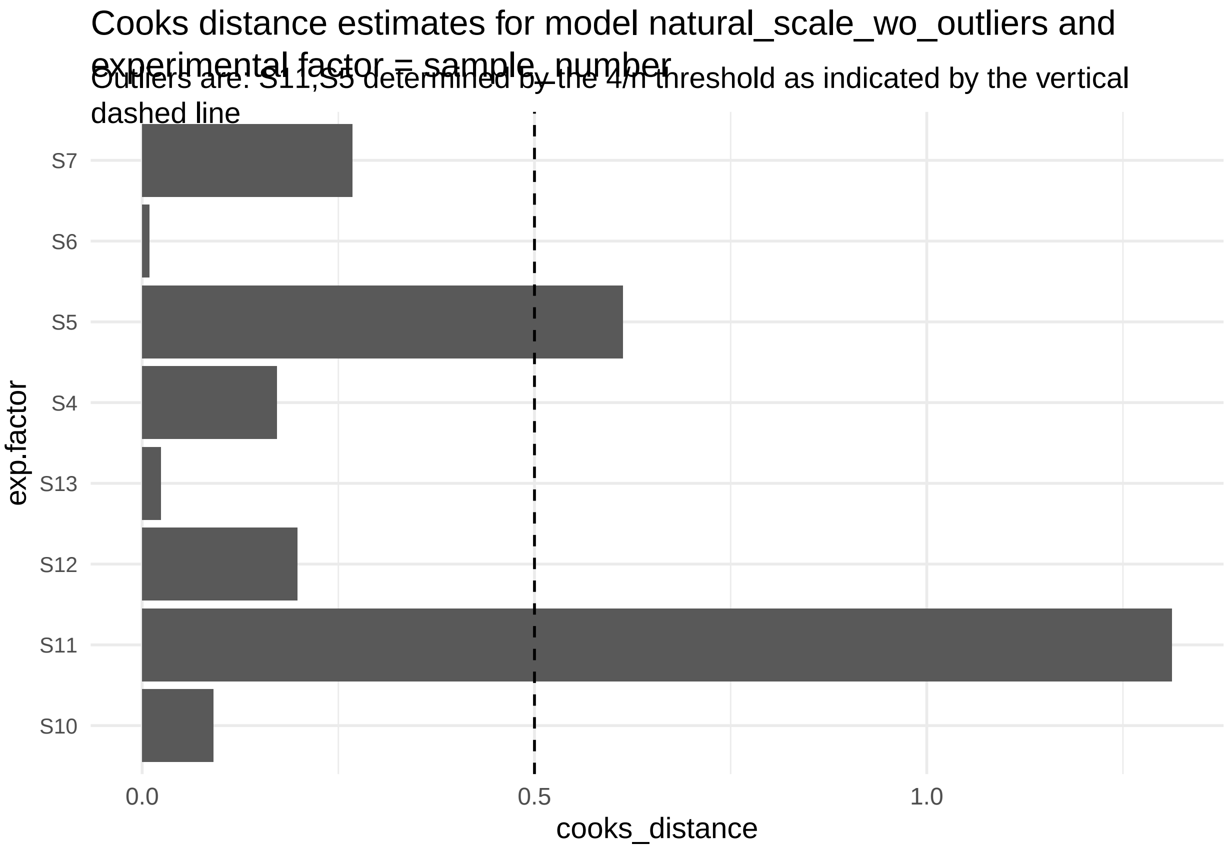


**Figure 47:**

```
print("Inferred outlier groups are")
```

[1] "Inferred outlier groups are"

```
print(diagnose_res$inferred_outlier_groups$natural_scale)
```

$sample_number [1] "S11" "S5"

We don't see a good fit to the assumption of negative binomial distribution based on the residual vs predicted plots suggesting that GLMM models for single cell RNA-seq data, even though they have been proposed as alternatives to dealing with the issue of pseudo-replication, are not good models to assess for differential expression.

Estimate log-fold change of the Snhg11 gene with genotype.

```
estimate_res <- getEstimatesOfInterest(data, design, model, print_plots = F)
print(estimate_res[[1]])
```

```
## Generalized linear mixed model fit by maximum likelihood (Laplace
##   Approximation) [glmerMod]
##  Family: Negative Binomial(4.914)  ( log )
## Formula:
## response_column ~ condition_column + covariate + (1 | experimental_column1) +
##     offset(log(total_column))
##    Data: fixed_global_variable_data
##
##       AIC       BIC    logLik -2*log(L)  df.resid
##   28790.0   28821.9  -14390.0   28780.0      4297
```

```
##
## Scaled residuals:
##     Min      1Q  Median      3Q     Max
## -1.8485 -0.7485 -0.1820  0.5882  5.4549
##
## Random effects:
##  Groups               Name        Variance Std.Dev.
##  experimental_column1 (Intercept) 0.07378  0.2716
## Number of obs: 4302, groups:  experimental_column1, 8
##
## Fixed effects:
##                                  Estimate Std. Error z value Pr(>|z|)
## (Intercept)                       -5.1736     0.1756 -29.460   <2e-16 ***
## condition_columnPS19-fE4-Syn1-Cre   0.4622     0.2280   2.027   0.0426 *
## covariateM                         0.1356     0.2335   0.581   0.5616
## ---
## Signif. codes:  0 '***' 0.001 '**' 0.01 '*' 0.05 '.' 0.1 ' ' 1
##
## Correlation of Fixed Effects:
##             (Intr) c_PS19
## c_PS19-E4-S -0.450
## covariateM  -0.588 -0.076
```

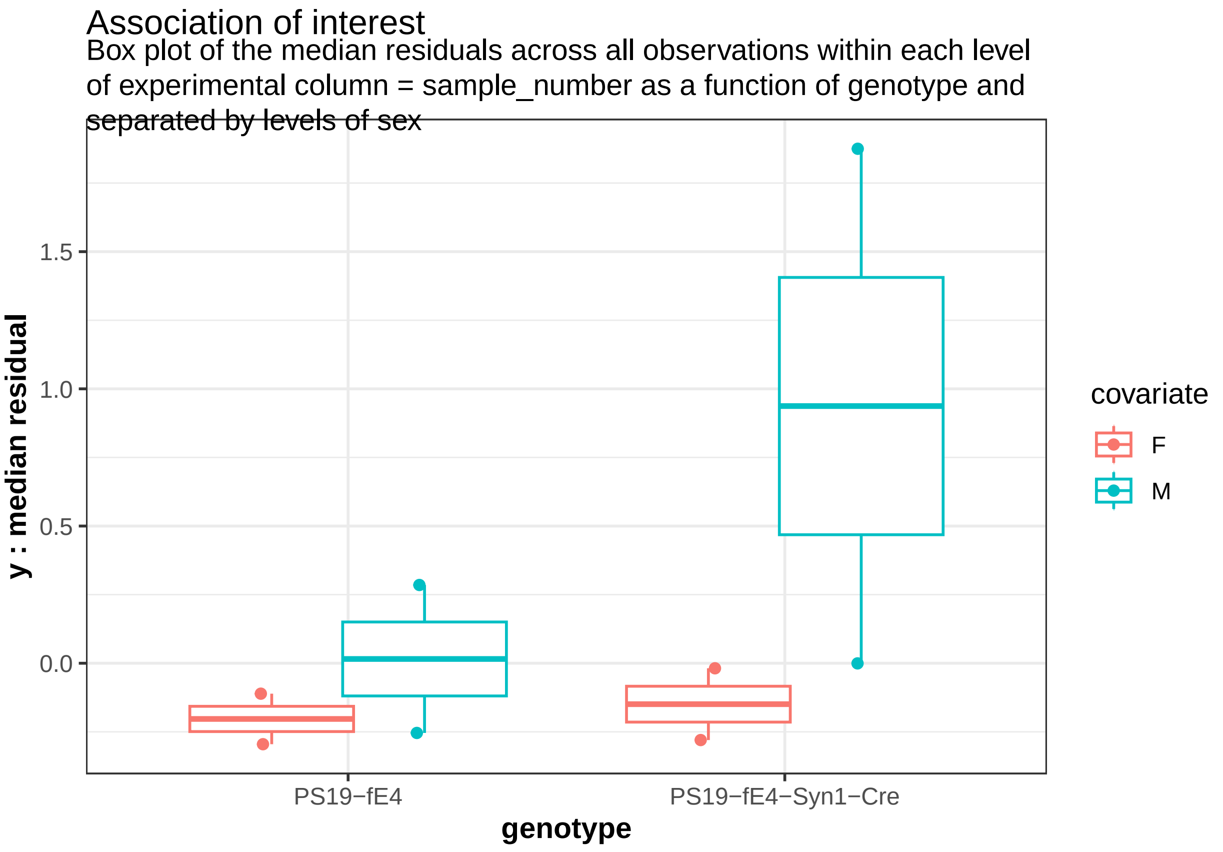


**Figure 48:** We can also perform a power analyses to detect a log(2) fold-change in this gene.

```
power_param@effect_size <- log(2)
power_res <- calculatePower(data, design, model, power_param)
```

```
##
## [1] "_________________________________Effect size of the covariate is now NA"
```

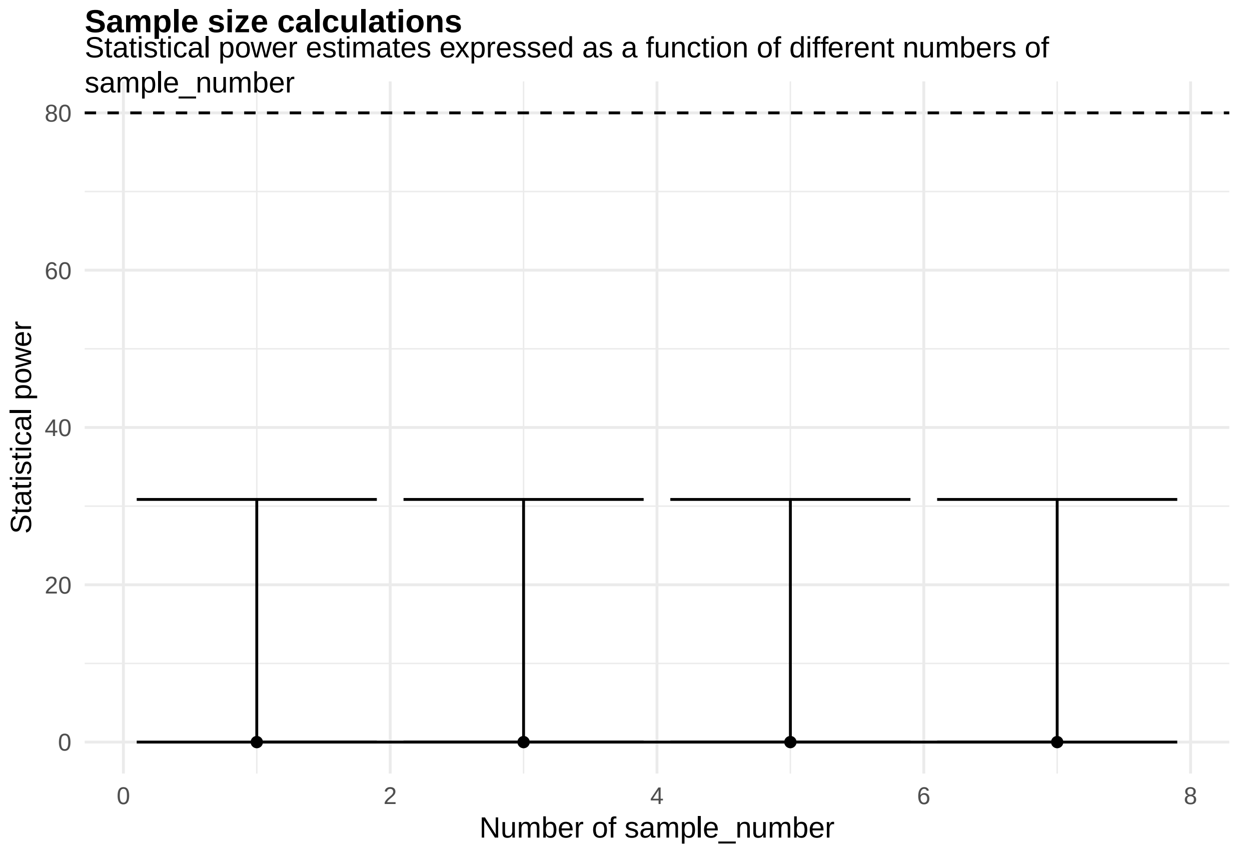


[[1]]

**Figure 49:**

#### 2.3 Behavior assays

Morris Water Maze (MWM) is an assay used to test spatial learning in animal models of diseases like Alzheimer's disease. The time recorded by each mouse to reach a target region (in the MWM) over multiple trials denoted as latency is the response of interest. Mouse models deficient in spatial learning would demonstrate an increased latency to reach a target region (invisible platform in Figure @ref(fig:MWM)) across the learning trials.

```
template_dir <- system.file("input_templates/behavior_data", package = "RMeDPower2")
data <- mouse_behavior_MWM_assay_data
design <- readDesign(file.path(template_dir, "design_behavior.json"))
model <- readProbabilityModel(file.path(template_dir, "prob_model.json"))
power_param <- readPowerParams(file.path(template_dir, "power_param.json"))
```

```
diagnose_res <- diagnoseDataModel(data, design, model)
```

```
## [1] "No outlier detected from the raw Data"
## [1] "No outlier detected from the raw Data"
```

To identify all the models fit to the data

```
print(diagnose_res$models)
```

```
## [1] "Natural scale"         "Log scale"             "Log scale wo outliers"
```

We see that three models were fit to this data - one in the natural scale of cell size, other in the logarithmic scale and the last one in log scale without outliers.

Let us look the basic QC plots for the models fit in the natural scale of the responses.

```
plots <- diagnose_res$diagnostic_plots$natural_scale$plots
captions <- diagnose_res$diagnostic_plots$natural_scale$captions

for (i in seq_along(plots)) {
  print(plots[[i]])
    cat(sprintf("\n\n\n**Figure %d:** %s. %s\n\n", plot_no,names(plots)[i], captions[i]))
  plot_no <- plot_no + 1
}
```

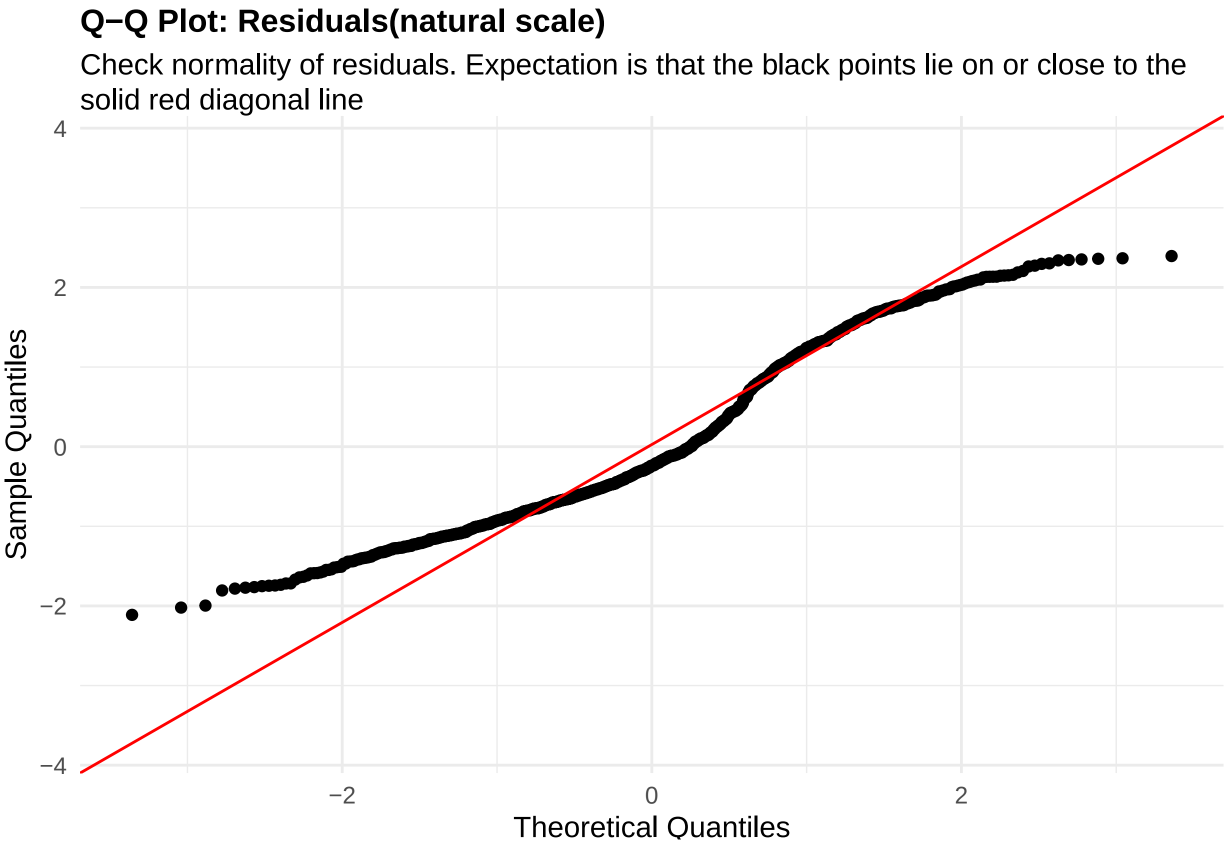


**Figure 50:** residuals_QQ. Check normality of residuals (natural scale). Expectation is that the black points lie on or close to the solid red diagonal line. Check the Normal Q-Q plots for good and bad examples here: https://library.virginia.edu/data/articles/diagnostic-plots

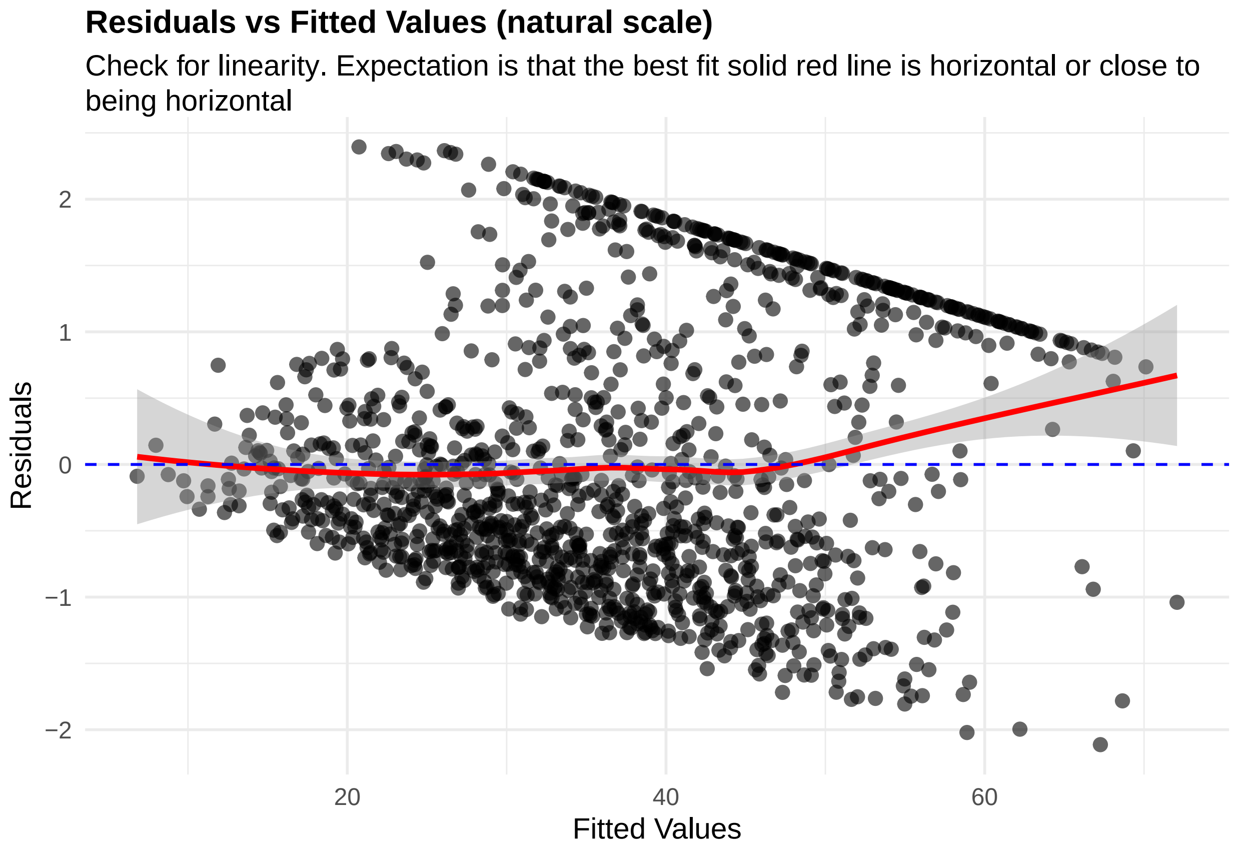


**Figure 51:** residuals_vs_fitted. Check for linearity (natural scale). Expectation is that the best fit solid red line is horizontal or close to being horizontal. Check the Residuals vs Fitted plots for good and bad examples here: https://library.virginia.edu/data/articles/diagnostic-plots

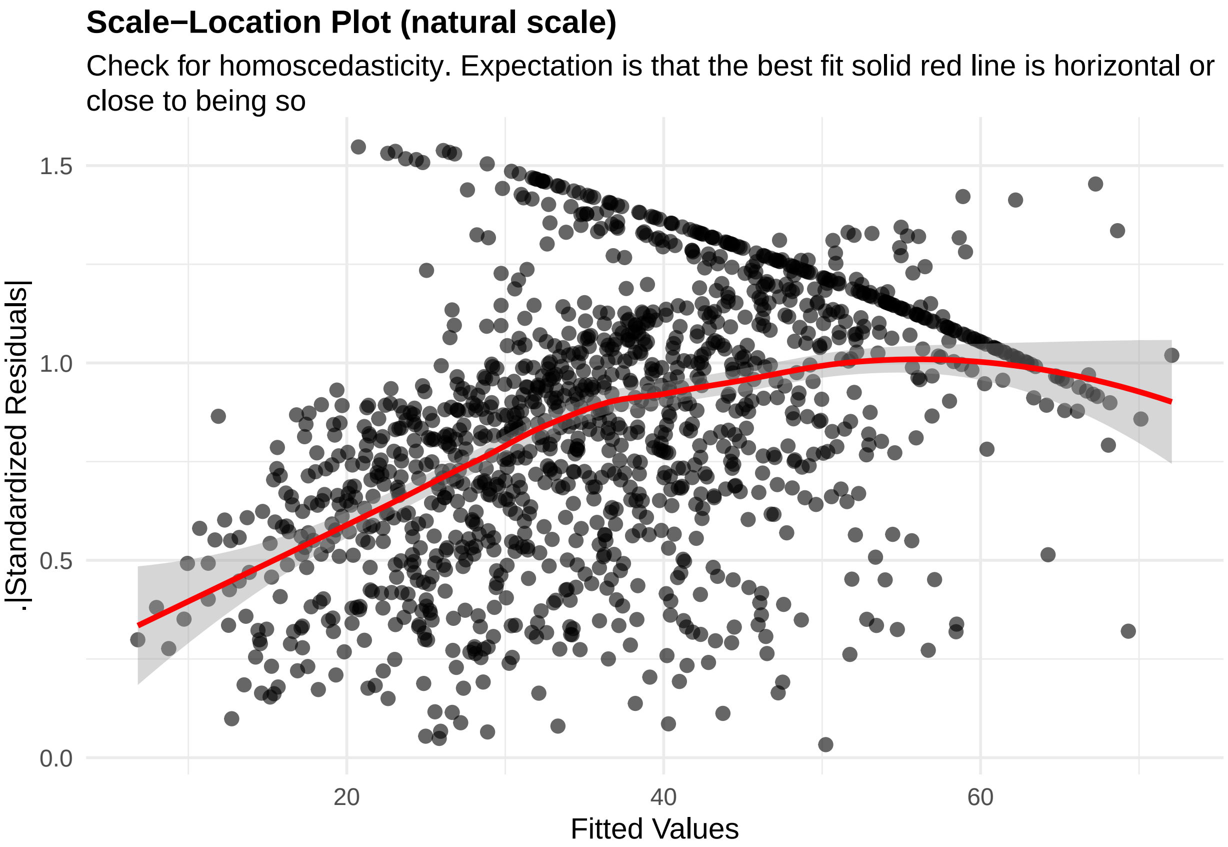


**Figure 52:** residuals_homoscedasticity. Check for homoscedasticity (natural scale). Expectation is that the best fit solid red line is horizontal or close to being so. Check the Scale-Location plots for good and bad examples here: https://library.virginia.edu/data/articles/diagnostic-plots

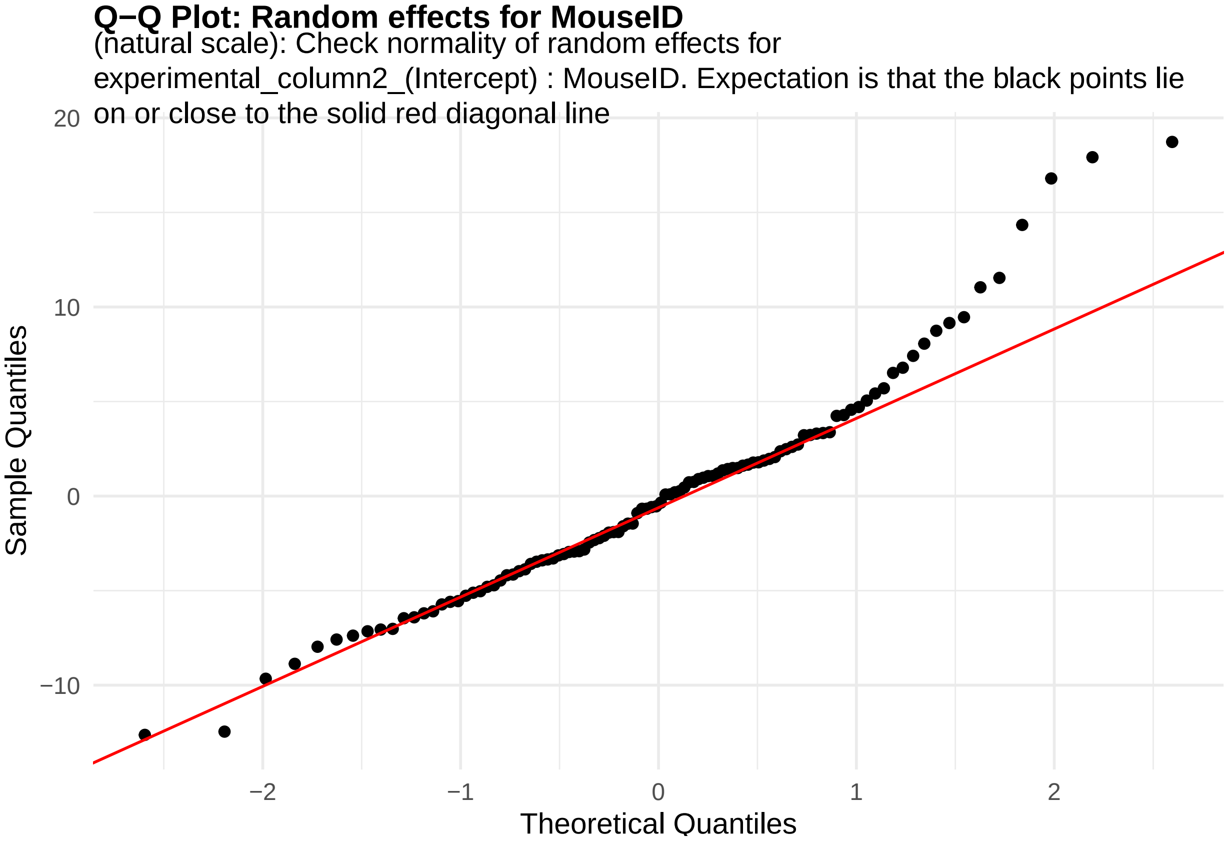


**Figure 53:** random_effects_QQ_1. Check normality of random effects for experimental_column2_(Intercept) (natural scale): MouseID. Expectation is that the black points lie on or close to the solid red diagonal line. Check the Normal Q-Q plots for good and bad examples here: https://library.virginia.edu/data/articles/diagnostic-plots

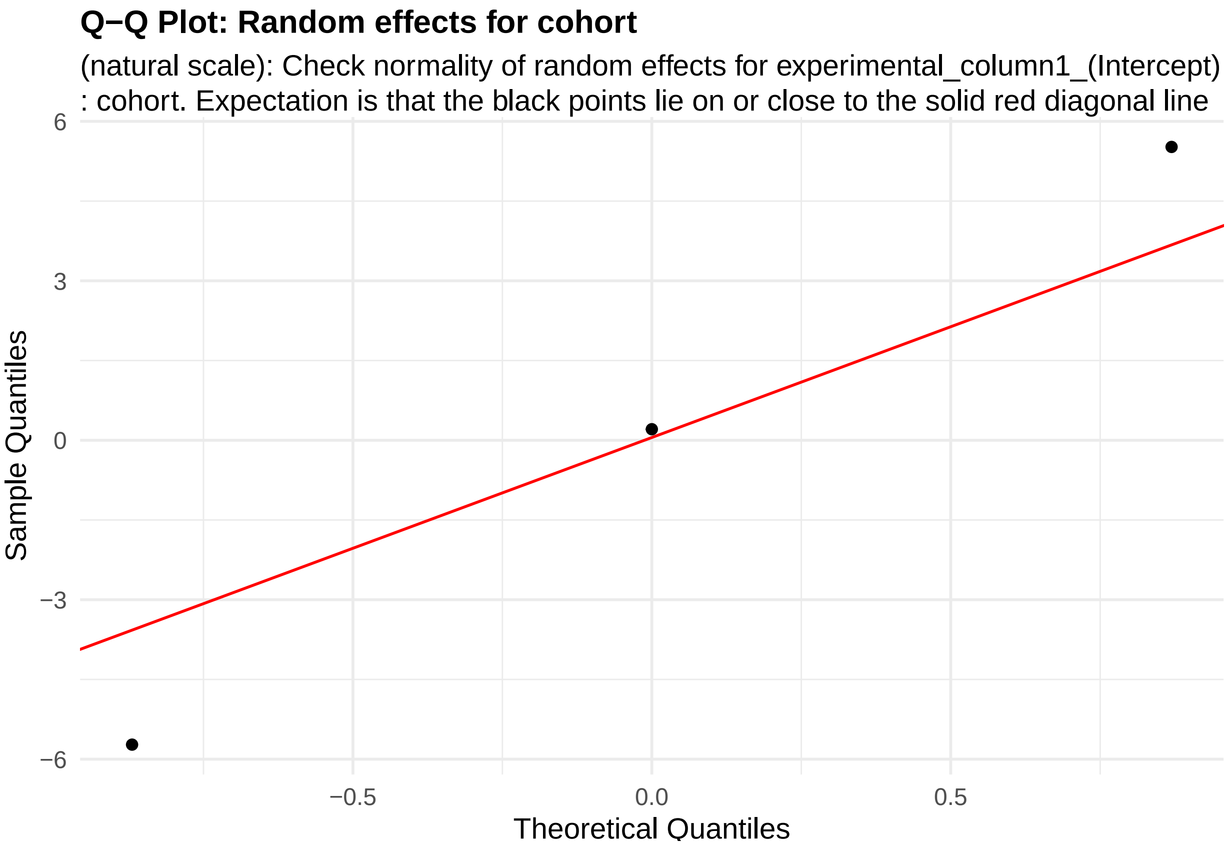


**Figure 54:** random_effects_QQ_2. Check normality of random effects for experimental_column1_(Intercept) (natural scale): cohort. Expectation is that the black points lie on or close to the solid red diagonal line. Check the Normal Q-Q plots for good and bad examples here: https://library.virginia.edu/data/articles/diagnostic-plots

```
cooks_plots <- diagnose_res$cooks_plots$natural_scale
cat("\n\n\n")
```

```
for(i in 1:length(cooks_plots)) {
  print(cooks_plots[[i]])
  cat(sprintf("\n\n\n**Figure %d:**\n\n", plot_no))
  plot_no <- plot_no + 1
}
```

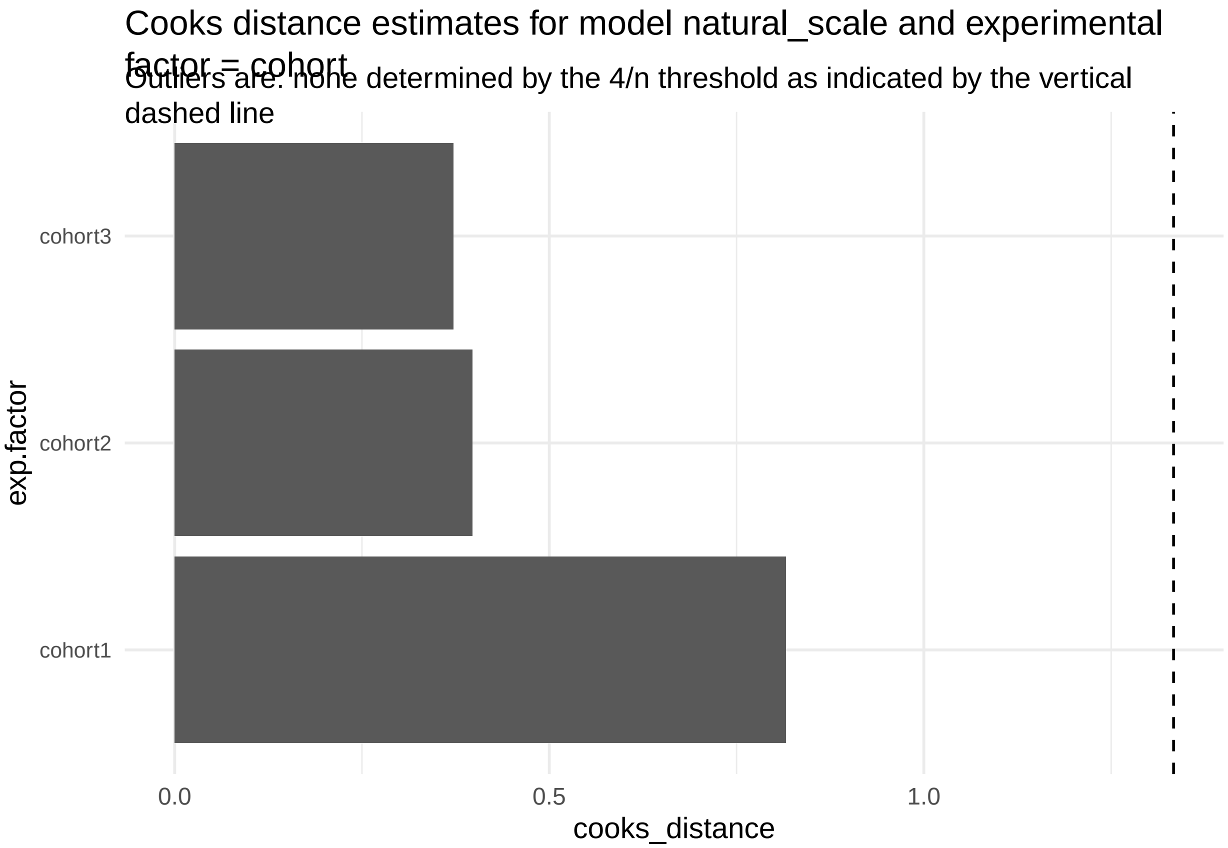

Cooks distance estimates for model natural_scale and experimental factor = cohort
Outliers are: none determined by the 4/n threshold as indicated by the vertical dashed line
cohort3
cohort2
cohort1
exp.factor
0.0
0.5
1.0
cooks_distance


**Figure 55:**

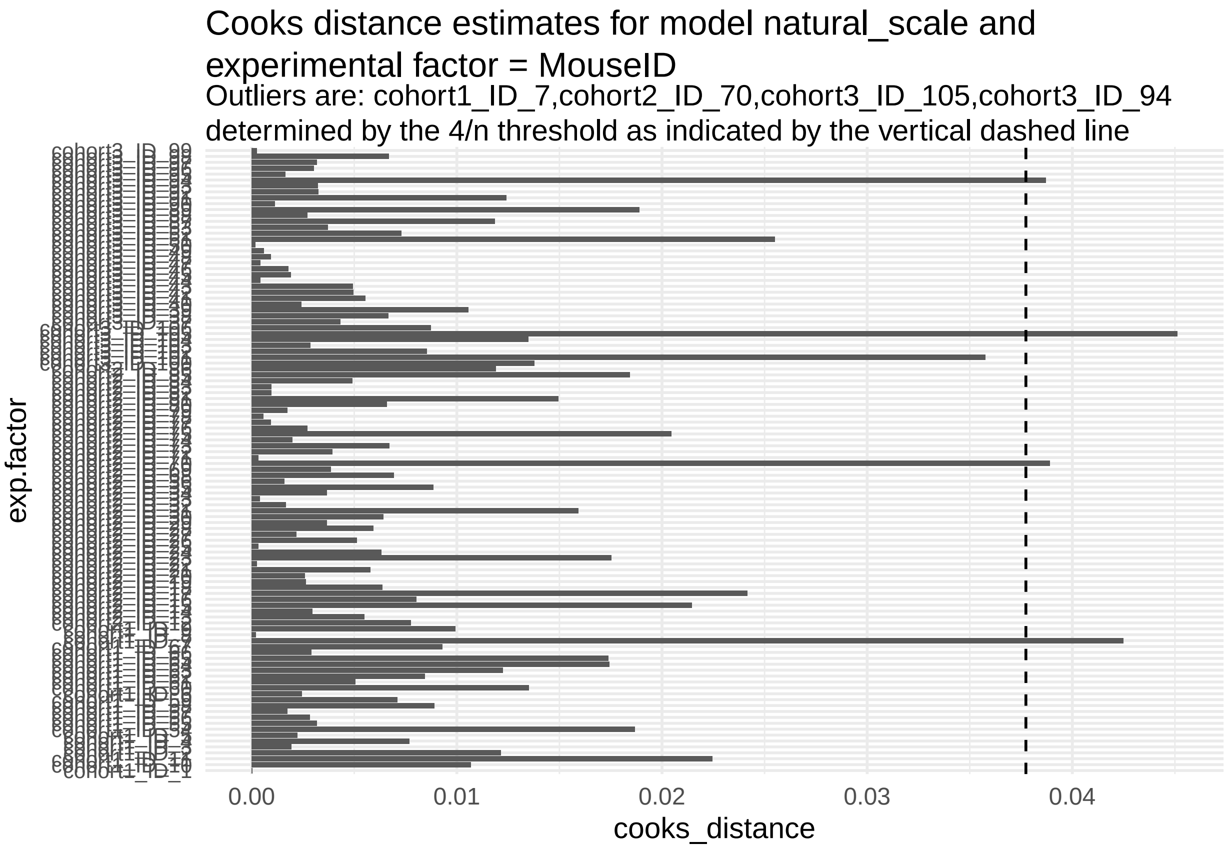


**Figure 56:**

```r
print("Inferred outlier groups are")
```

[1] "Inferred outlier groups are"

```r
print(diagnose_res$inferred_outlier_groups$natural_scale)
```

$cohort [1] "none"

$MouseID [1] "cohort1_ID_7" "cohort2_ID_70" "cohort3_ID_105" "cohort3_ID_94"

Now, let us look at the same set of QC plots for models on the log-transformed responses.

```r
plots <- diagnose_res$diagnostic_plots$log_scale$plots
captions <- diagnose_res$diagnostic_plots$log_scale$captions

for (i in seq_along(plots)) {
  print(plots[[i]])
    cat(sprintf("\n\n\n**Figure %d:** %s. %s\n\n", plot_no,names(plots)[i], captions[i]))
  plot_no <- plot_no + 1
}
```

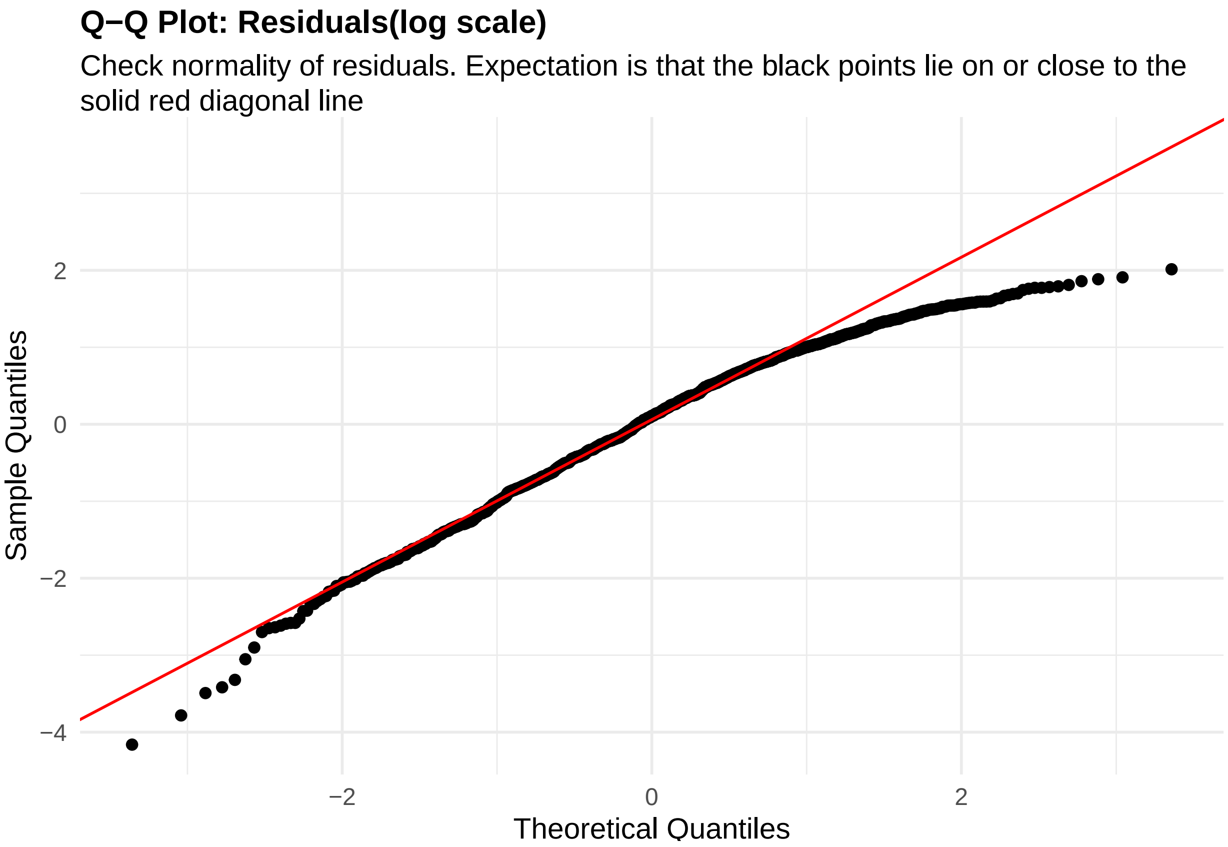


**Figure 57:** residuals_QQ. Check normality of residuals (log scale). Expectation is that the black points lie on or close to the solid red diagonal line. Check the Normal Q-Q plots for good and bad examples here: https://library.virginia.edu/data/articles/diagnostic-plots

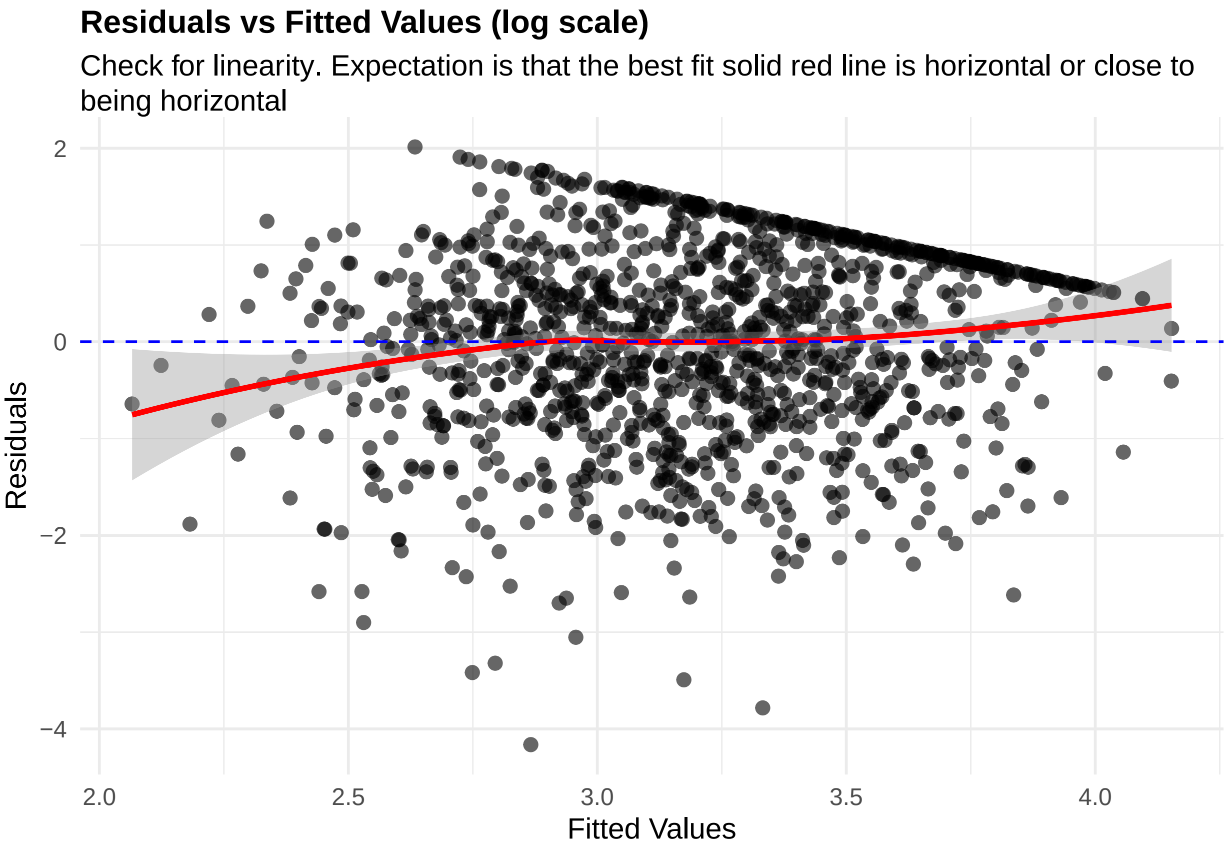


**Figure 58:** residuals_vs_fitted. Check for linearity (log scale). Expectation is that the best fit solid red line is horizontal or close to being horizontal. Check the Residuals vs Fitted plots for good and bad examples here: https://library.virginia.edu/data/articles/diagnostic-plots

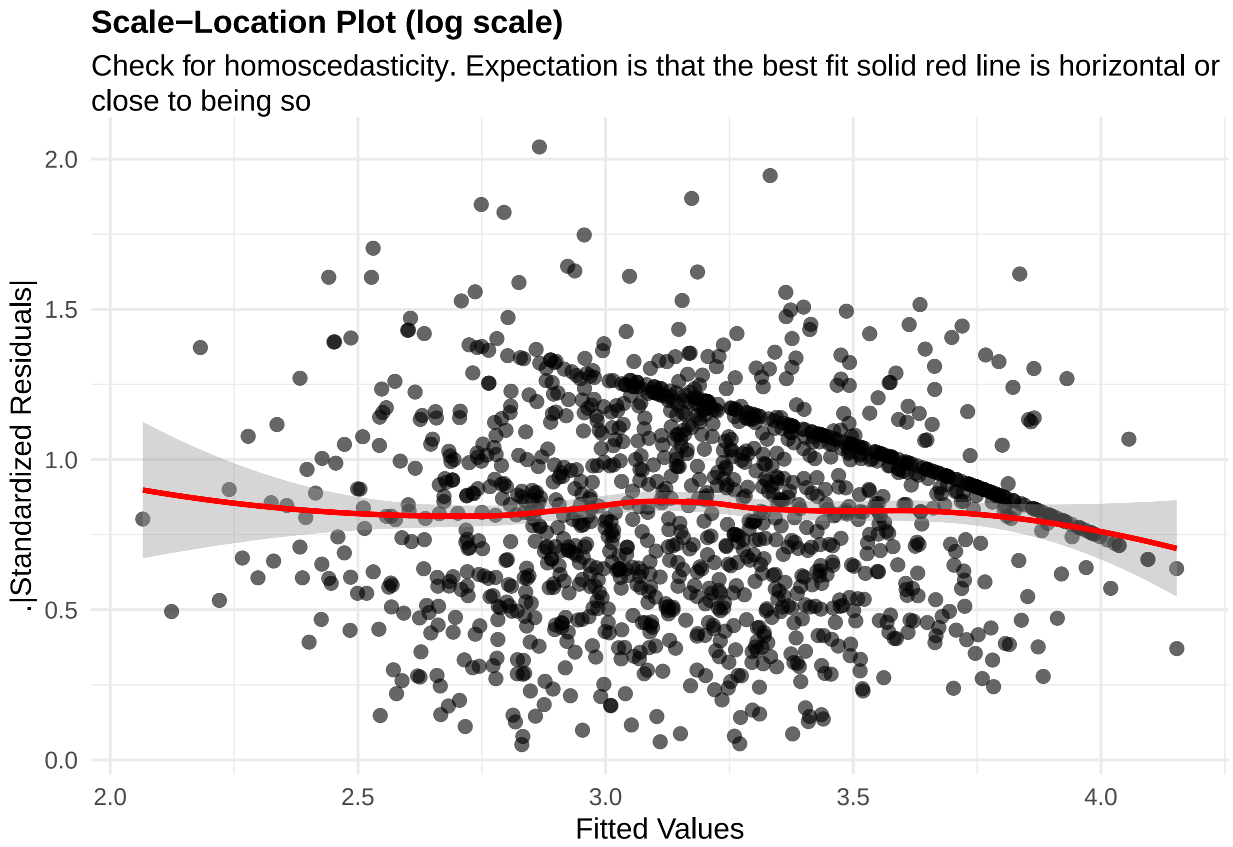


**Figure 59:** residuals_homoscedasticity. Check for homoscedasticity (log scale). Expectation is that the best fit solid red line is horizontal or close to being so. Check the Scale-Location plots for good and bad examples here: https://library.virginia.edu/data/articles/diagnostic-plots

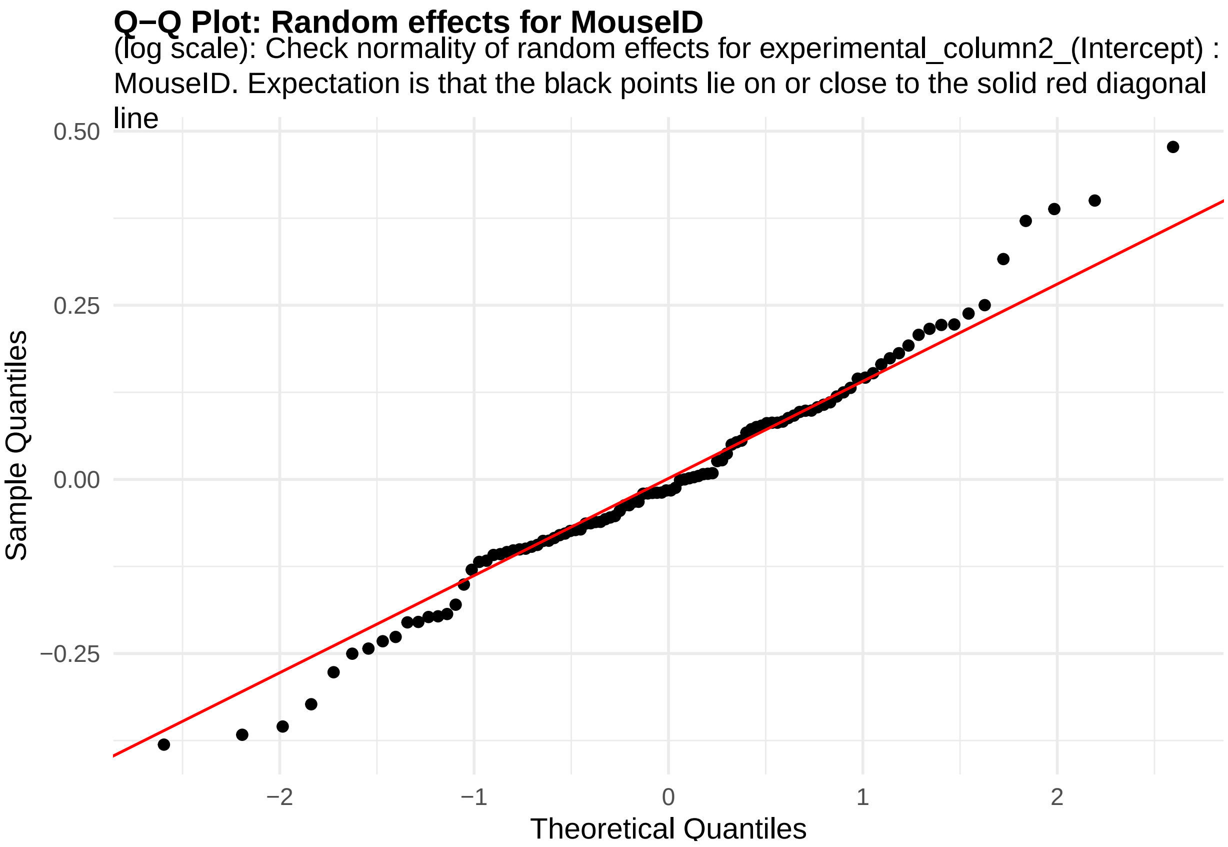


**Figure 60:** random_effects_QQ_1. Check normality of random effects for experimental_column2_(Intercept) (log scale): MouseID. Expectation is that the black points lie on or close to the solid red diagonal line. Check the Normal Q-Q plots for good and bad examples here: https://library.virginia.edu/data/articles/diagnostic-plots

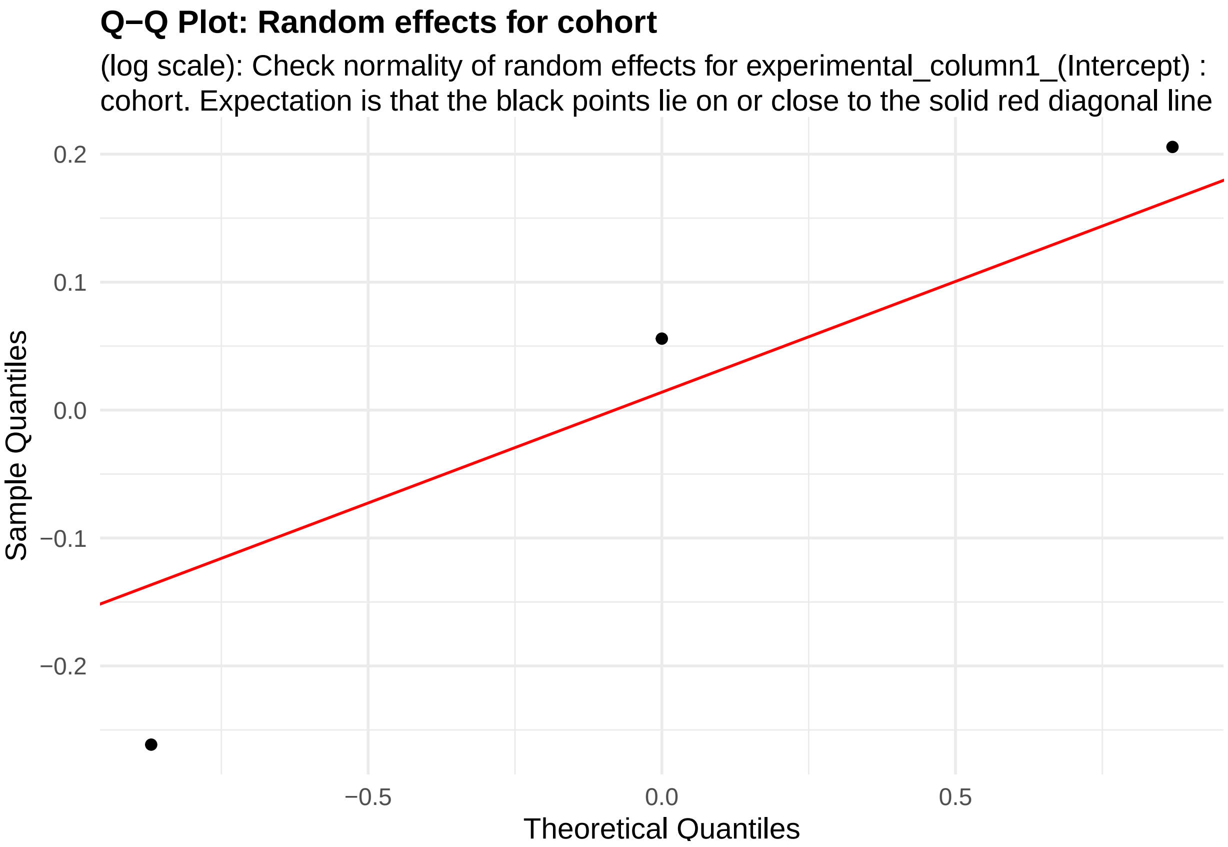


**Figure 61:** random_effects_QQ_2. Check normality of random effects for experimental_column1_(Intercept) (log scale): cohort. Expectation is that the black points lie on or close to the solid red diagonal line. Check the Normal Q-Q plots for good and bad examples here: https://library.virginia.edu/data/articles/diagnostic-plots

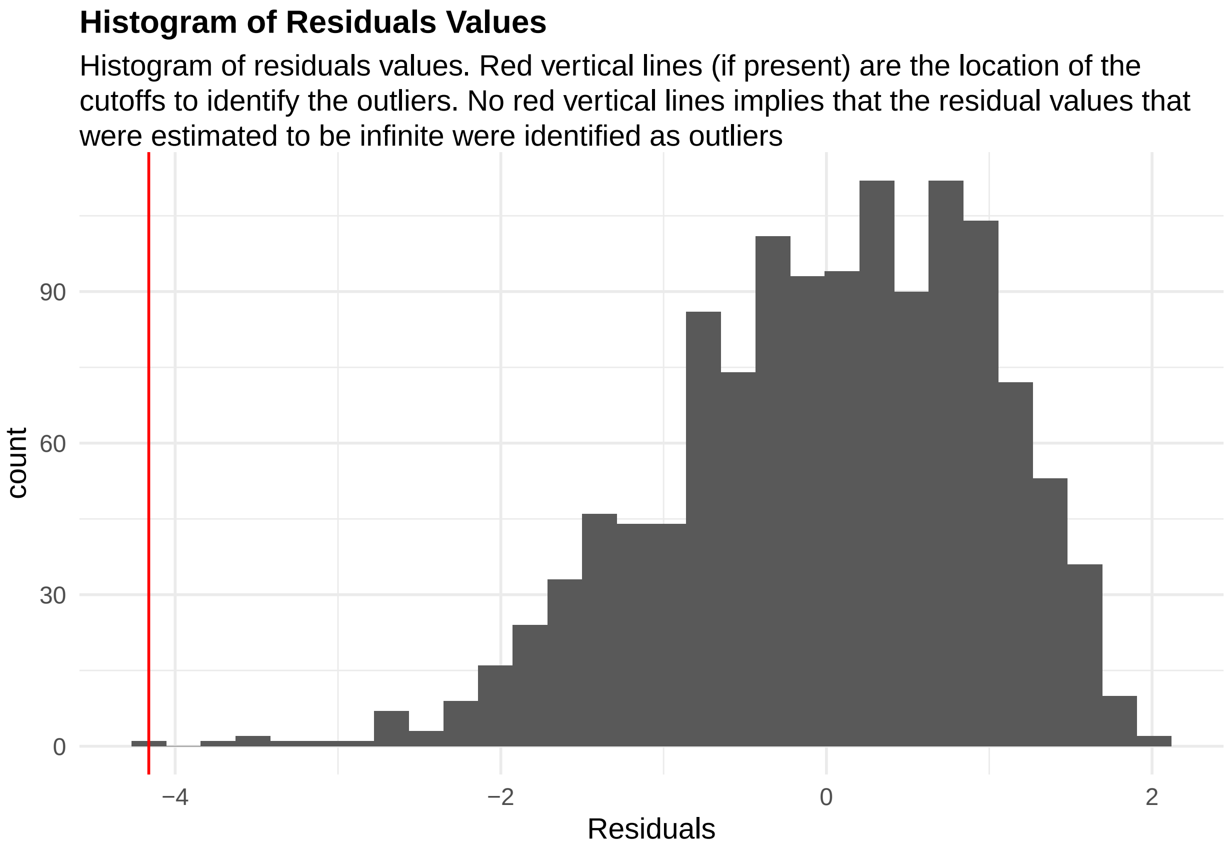


**Figure 62:** residuals_histogram. Histogram of residuals values. Red vertical lines (if present) are the location of the cutoffs to identify the outliers. No red vertical lines implies that the residual values that were estimated to be infinite were identified as outliers

```
cooks_plots <- diagnose_res$cooks_plots$log_scale
cat("\n\n\n")
```

```
for(i in 1:length(cooks_plots)) {
  print(cooks_plots[[i]])
  cat(sprintf("\n\n\n**Figure %d:**\n\n", plot_no))
  plot_no <- plot_no + 1
}
```

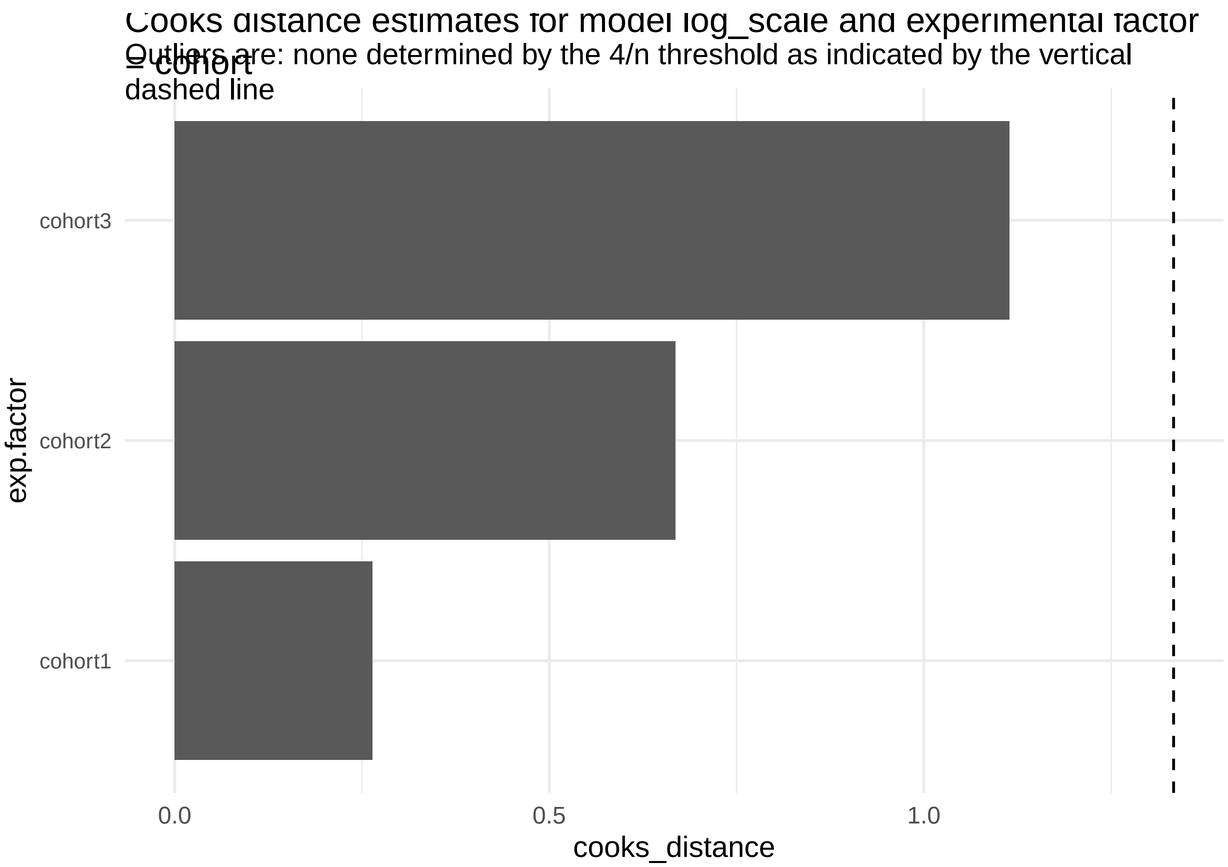
Cooks distance estimates for model log_scale and experimental factor = cohort
Outliers are: none determined by the 4/n threshold as indicated by the vertical dashed line
cohort3
cohort2
cohort1
exp.factor
0.0
0.5
1.0
cooks_distance

**Figure 63:**

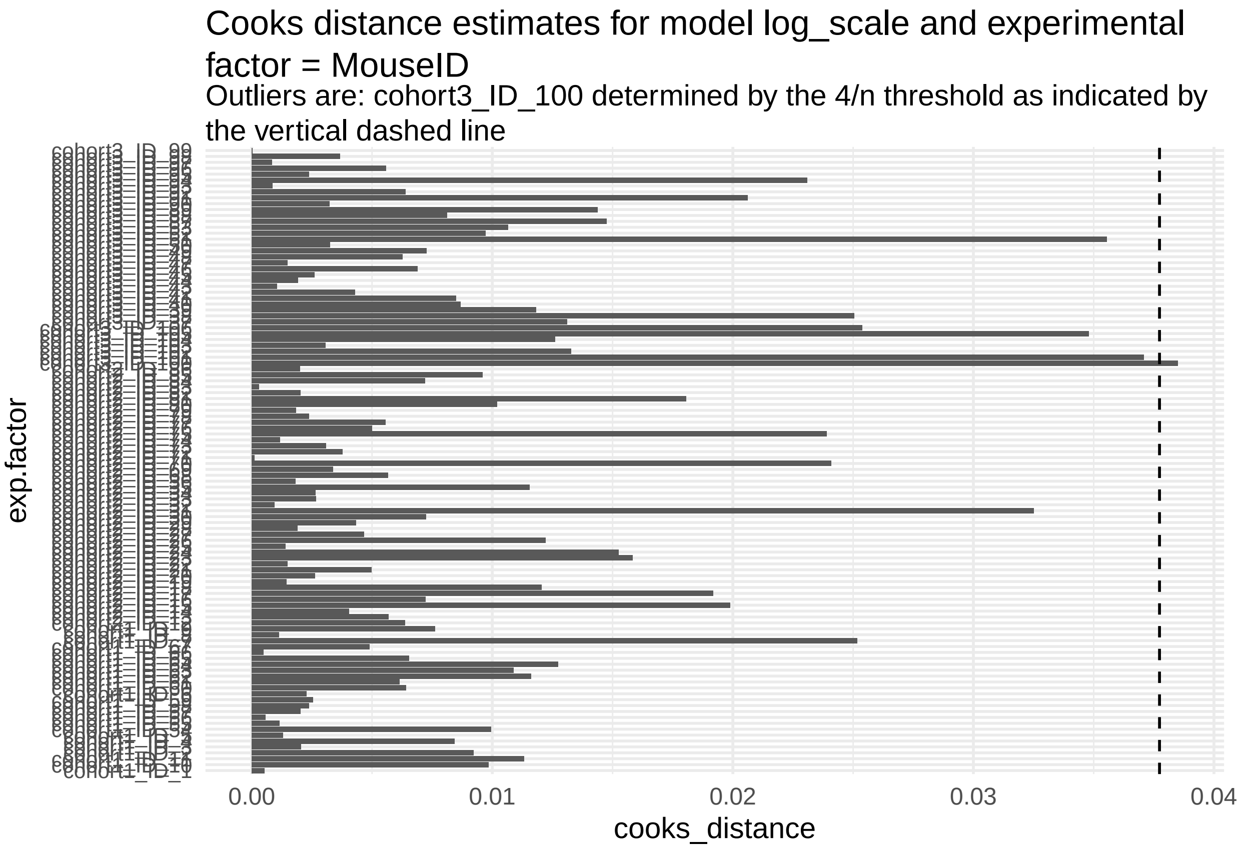


**Figure 64:**

```r
print("Inferred outlier groups are")
```

[1] "Inferred outlier groups are"

```r
print(diagnose_res$inferred_outlier_groups$natural_scale)
```

$cohort [1] "none"

$MouseID [1] "cohort1_ID_7" "cohort2_ID_70" "cohort3_ID_105" "cohort3_ID_94"

We see normality assumption is better met with the log-transformed model (Figure 6) compared to the model in the natural scale (Figure 1).

The code also outputs Cook's distance measures for each cohort and mouse that is intended to capture how influential each of these were in determining the final parameter estimates in the LMM model.

There aren't any obvious outliers in any of these models for cohort or mice.

Based on the above plots, it seems like the log transformed versions of the latency responses better fit the model assumptions. Therefore using these transformed responses we will estimate and visualize our parameter of interest.

```r
data_update <- diagnose_res$Data_updated
design_update <- design
design_update@response_column %<>% paste0(., "_logTransformed")
estimate_res <- getEstimatesOfInterest(data_update, design_update, model, print_plots = F)
print(estimate_res[[1]])
```

```
## Linear mixed model fit by REML. t-tests use Satterthwaite's method [
## lmerModLmerTest]
```

```
## Formula:
## response_column ~ condition_column + covariate + (1 | experimental_column1) +
##     (1 | experimental_column2)
##    Data: fixed_global_variable_data
##
## REML criterion at convergence: 3430.9
##
## Scaled residuals:
##     Min      1Q  Median      3Q     Max
## -3.8168 -0.6592  0.1087  0.7697  1.9260
##
## Random effects:
##  Groups               Name        Variance Std.Dev.
##  experimental_column2 (Intercept) 0.06004  0.2450
##  experimental_column1 (Intercept) 0.05800  0.2408
##  Residual                         0.81359  0.9020
## Number of obs: 1271, groups:
## experimental_column2, 106; experimental_column1, 3
##
## Fixed effects:
##                     Estimate Std. Error         df t value Pr(>|t|)
## (Intercept)        3.416e+00  1.554e-01  2.722e+00  21.983 0.000381 ***
## condition_column1  3.716e-01  6.992e-02  1.022e+02   5.315 6.29e-07 ***
## covariate         -5.878e-02  7.328e-03  1.164e+03  -8.022 2.52e-15 ***
## ---
## Signif. codes:  0 '***' 0.001 '**' 0.01 '*' 0.05 '.' 0.1 ' ' 1
##
## Correlation of Fixed Effects:
##             (Intr) cndt_1
## cndtn_clmn1 -0.230
## covariate   -0.307  0.000
```

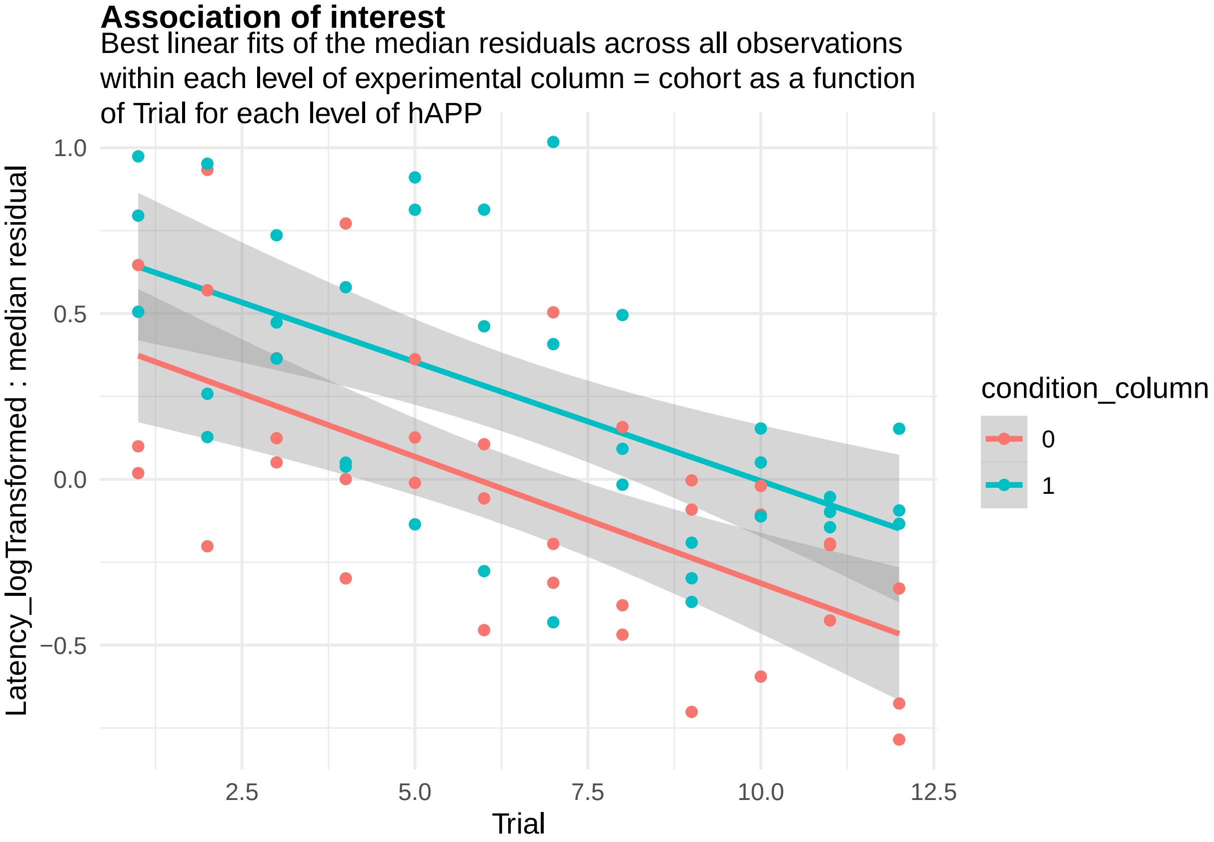


**Figure 65:** We will also perform sample-size calculations for the number of mice needed per cohort.

```
power_param@target_columns <- "MouseID"
power_res <- calculatePower(data_update, design_update, model, power_param)
```

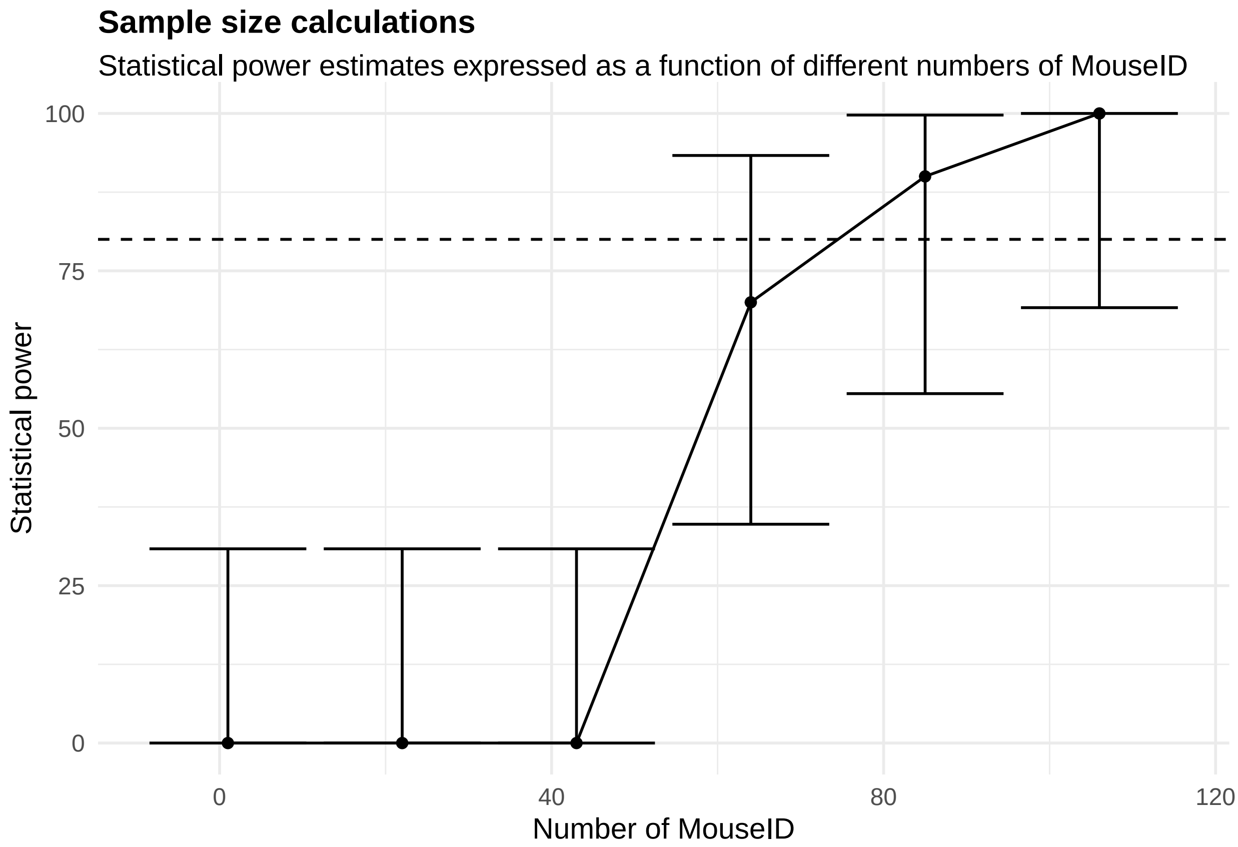


[[1]]

**Figure 66:**

#### 2.4 Brain electrical signal assays

Patch clamp is a technique that can be used to measure membrane action potential in excitable cells like neurons. We will work with (simulated) data from two sets of mice - one representing a Alzheimer's Disease model (hAPP) and the other representing control or wild-type mice. In each of these mice, multiple brain tissue slices are isolated and the membrane action potential for at least one neuronal cell in each slice is recorded.

```
template_dir <- system.file("input_templates/electrical_patch_clamp_data",package = "RMeDPower2")
data <- mouse_brain_electro_physiology_data
head(data)
```

```
##   negative_action_potential_mV    Mice Genotype    Sex slice_number
## 1                     56.23547 Mouse_1     hAPP   Male            1
## 2                     47.89597 Mouse_3     hAPP   Male            1
## 3                     56.20910 Mouse_3     hAPP   Male            2
## 4                     53.14394 Mouse_9     hAPP Female            1
## 5                     53.67793 Mouse_9     hAPP Female            1
## 6                     61.21780 Mouse_9     hAPP Female            1
```

```
design <- readDesign(file.path(template_dir, "design_patch_clamp.json"))
print(design)
```

```
## An object of class "RMeDesign"
## Slot "response_column":
## [1] "negative_action_potential_mV"
##
```

```
## Slot "covariate":
## NULL
##
## Slot "condition_column":
## [1] "Genotype"
##
## Slot "condition_is_categorical":
## [1] TRUE
##
## Slot "experimental_columns":
## [1] "Mice"         "slice_number"
##
## Slot "crossed_columns":
## NULL
##
## Slot "total_column":
## NULL
##
## Slot "outlier_alpha":
## [1] 0.05
##
## Slot "na_action":
## [1] "complete"
##
## Slot "include_interaction":
## [1] NA
##
## Slot "random_slope_variable":
## NULL
##
## Slot "covariate_is_categorical":
## [1] NA
```

```r
model <- readProbabilityModel(file.path(template_dir, "prob_model.json"))
print(model)
```

```
## An object of class "ProbabilityModel"
## Slot "error_is_non_normal":
## [1] FALSE
##
## Slot "family_p":
## NULL
```

```r
power_param <- readPowerParams(file.path(template_dir, "power_param.json"))
print(power_param)
```

```
## An object of class "PowerParams"
## Slot "target_columns":
## [1] "Mice"
##
## Slot "power_curve":
## [1] 1
##
## Slot "nsimn":
## [1] 10
```

```
##
## Slot "levels":
## [1] 1
##
## Slot "max_size":
## [1] 1
##
## Slot "alpha":
## [1] 0.05
##
## Slot "breaks":
## NULL
##
## Slot "effect_size":
## NULL
##
## Slot "icc":
## NULL
```

We will now evaluate the model.

```
diagnose_res <-diagnoseDataModel(data, design, model)
```

```
## [1] "No outlier detected from the raw Data"
```

```
## [1] "No outlier detected from the raw Data"
```

To identify all the models fit to the data

```
print(diagnose_res$models)
```

```
## [1] "Natural scale"              "Natural scale wo outliers"
## [3] "Log scale"
```

We see that three models were fit to this data - one in the natural scale of negative_action_potential_mV, other in the natural scale without outliers and the last one in log scale.

Let us look the basic QC plots for the models fit in the natural scale of the responses.

```
plots <- diagnose_res$diagnostic_plots$natural_scale$plots
captions <- diagnose_res$diagnostic_plots$natural_scale$captions
for (i in seq_along(plots)) {
  print(plots[[i]])
    cat(sprintf("\n\n\n**Figure %d:** %s. %s\n\n", plot_no,names(plots)[i], captions[i]))
  plot_no <- plot_no + 1
}
```

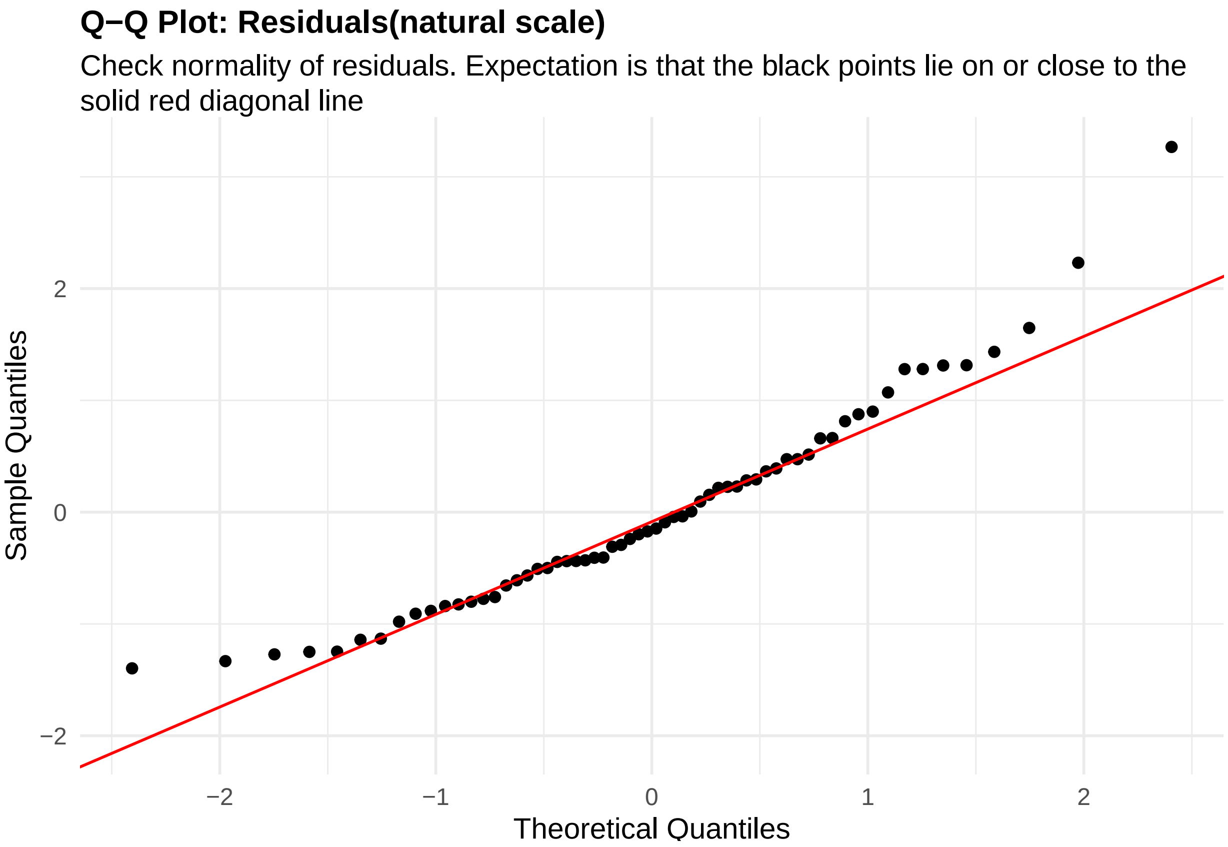


**Figure 67:** residuals_QQ. Check normality of residuals (natural scale). Expectation is that the black points lie on or close to the solid red diagonal line. Check the Normal Q-Q plots for good and bad examples here: https://library.virginia.edu/data/articles/diagnostic-plots

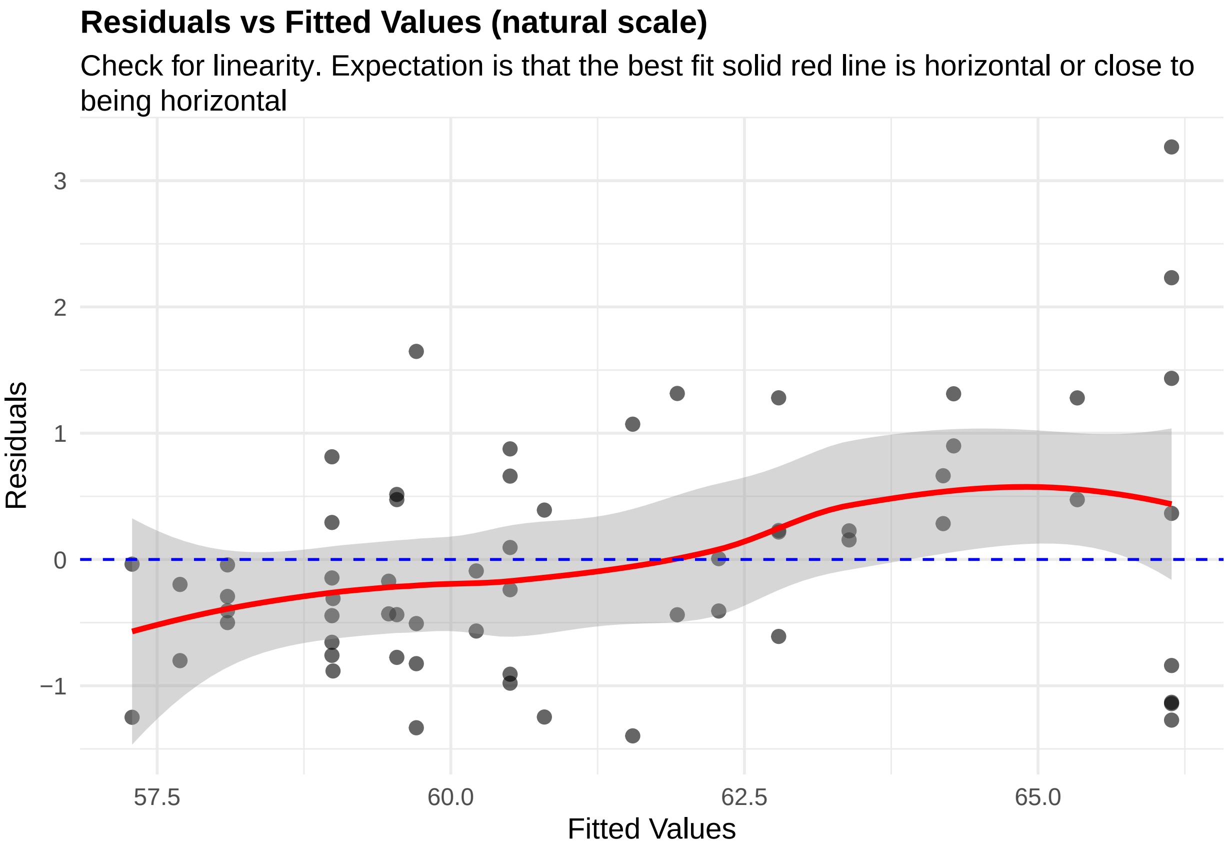


**Figure 68:** residuals_vs_fitted. Check for linearity (natural scale). Expectation is that the best fit solid red line is horizontal or close to being horizontal. Check the Residuals vs Fitted plots for good and bad examples here: https://library.virginia.edu/data/articles/diagnostic-plots

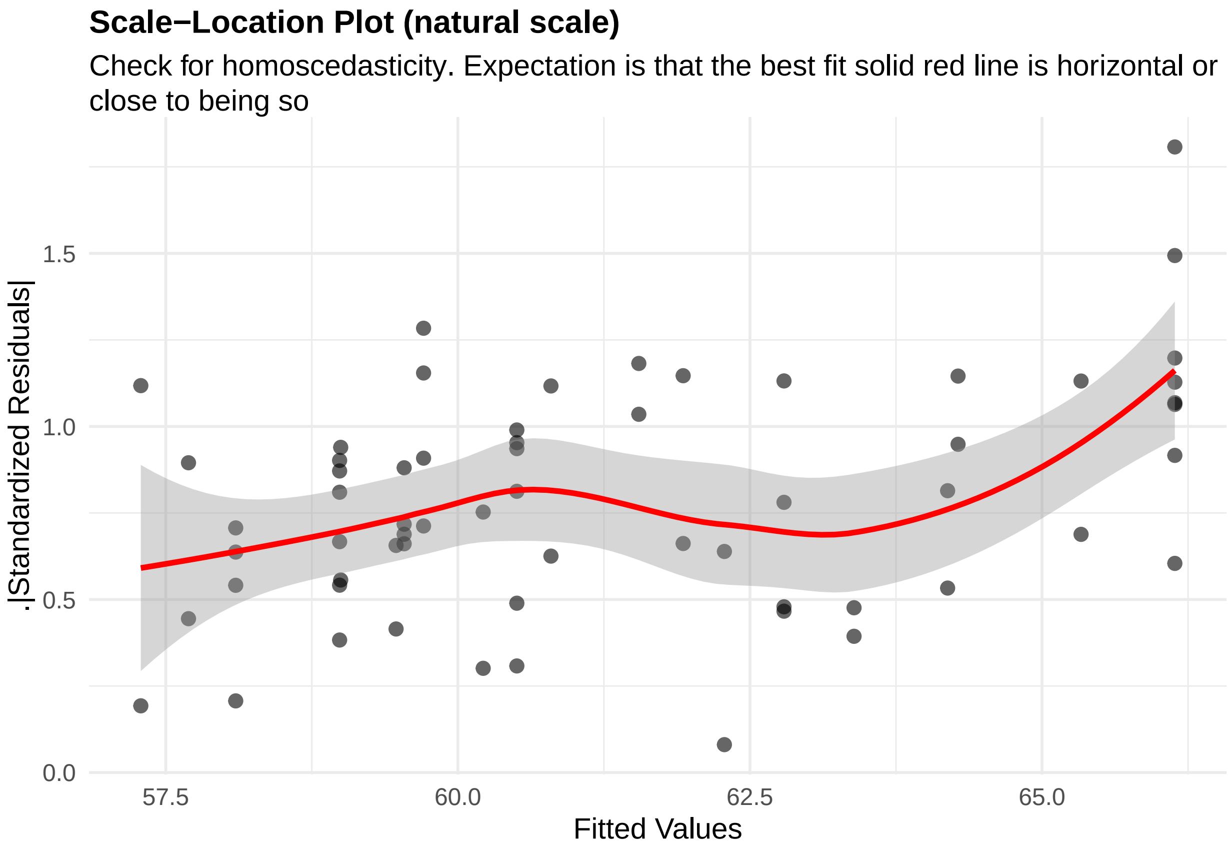


**Figure 69:** residuals_homoscedasticity. Check for homoscedasticity (natural scale). Expectation is that the best fit solid red line is horizontal or close to being so. Check the Scale-Location plots for good and bad examples here: https://library.virginia.edu/data/articles/diagnostic-plots

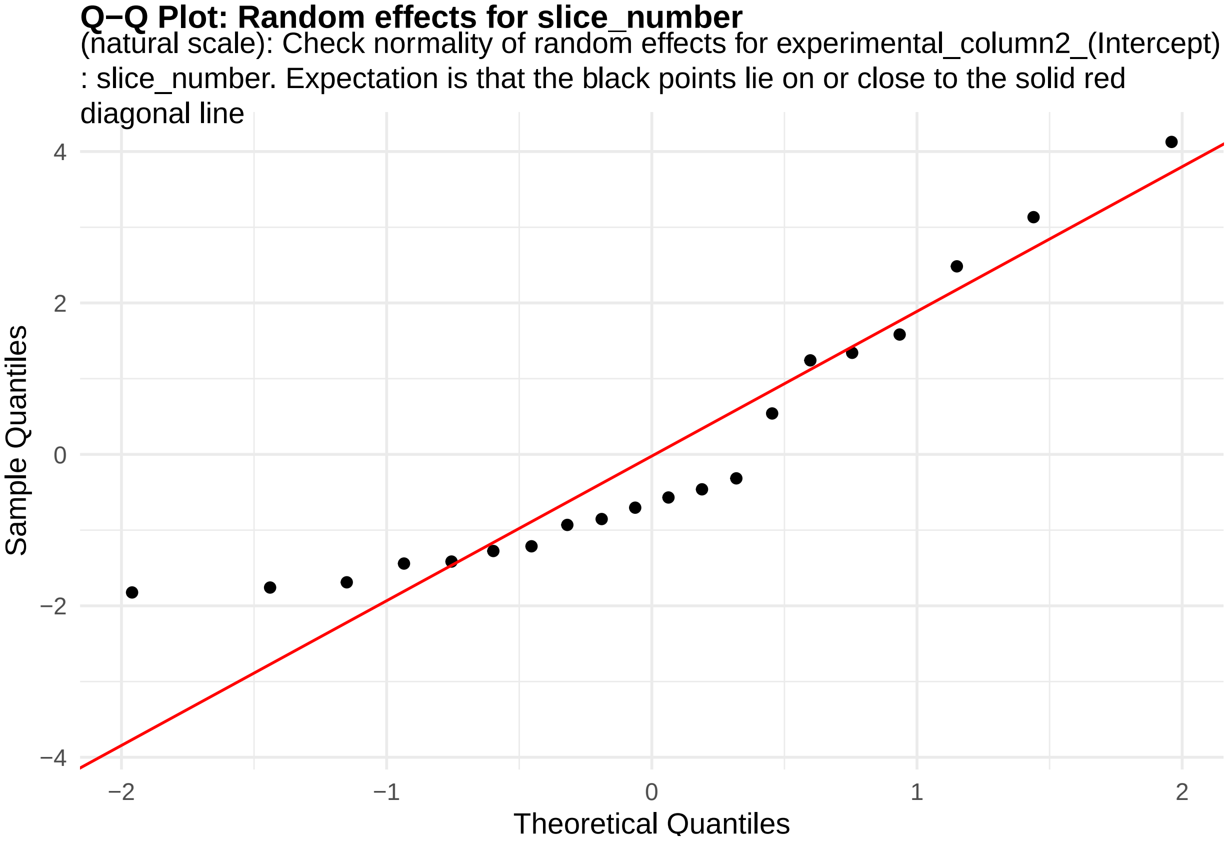


**Figure 70:** random_effects_QQ_1. Check normality of random effects for experimental_column2_(Intercept) (natural scale): slice_number. Expectation is that the black points lie on or close to the solid red diagonal line. Check the Normal Q-Q plots for good and bad examples here: https://library.virginia.edu/data/articles/diagnostic-plots

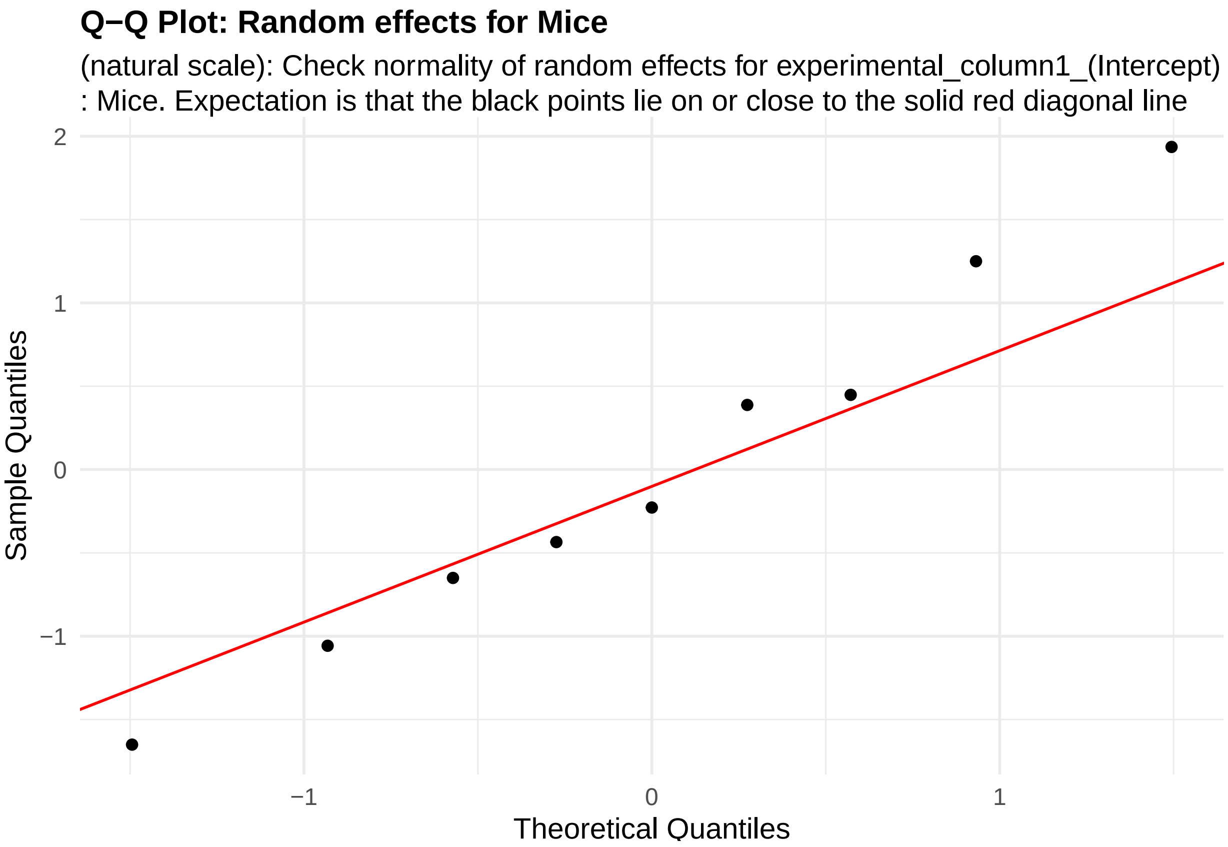


**Figure 71:** random_effects_QQ_2. Check normality of random effects for experimental_column1_(Intercept) (natural scale): Mice. Expectation is that the black points lie on or close to the solid red diagonal line. Check the Normal Q-Q plots for good and bad examples here: https://library.virginia.edu/data/articles/diagnostic-plots

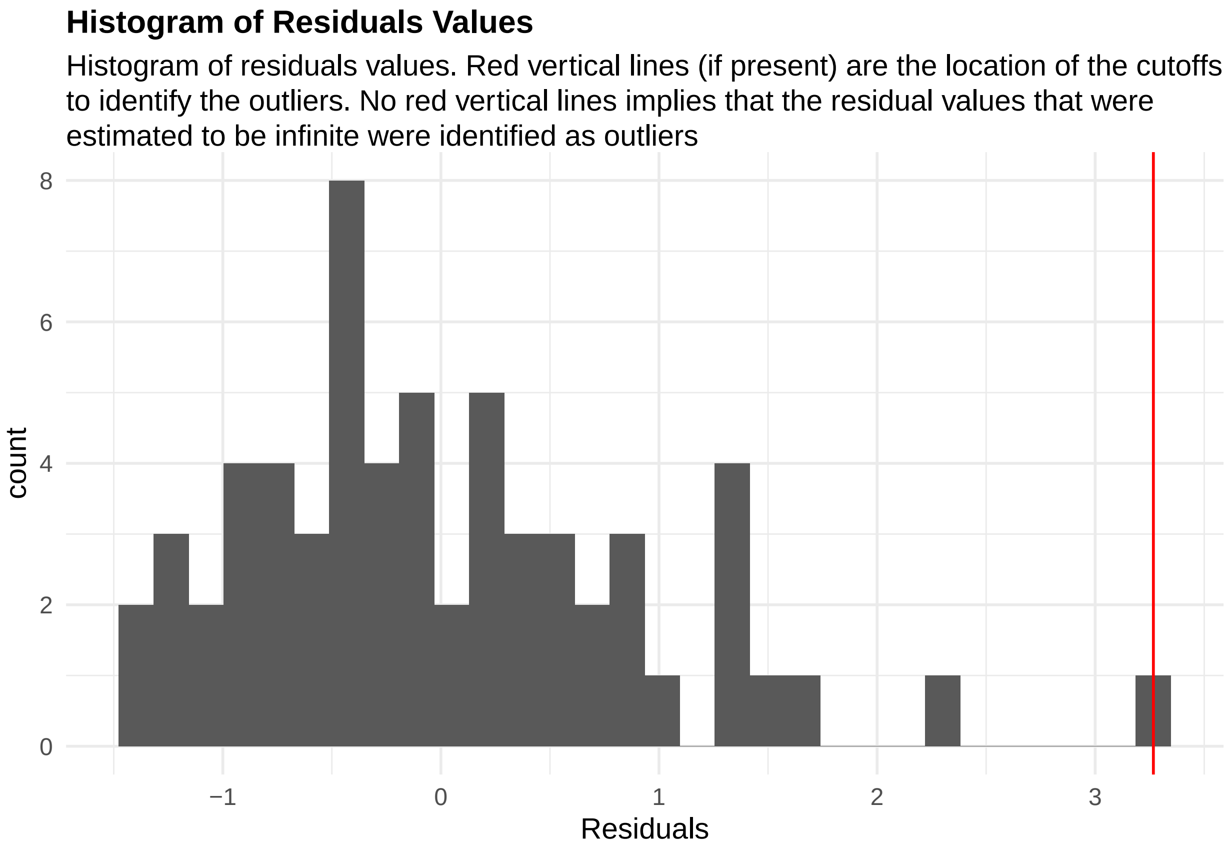


**Figure 72:** residuals_histogram. Histogram of residuals values. Red vertical lines (if present) are the location of the cutoffs to identify the outliers. No red vertical lines implies that the residual values that were estimated to be infinite were identified as outliers

```r
cooks_plots <- diagnose_res$cooks_plots$natural_scale
cat("\n\n\n")
```

```r
for(i in 1:length(cooks_plots)) {
  print(cooks_plots[[i]])
  cat(sprintf("\n\n\n**Figure %d:**\n\n", plot_no))
  plot_no <- plot_no + 1
}
```

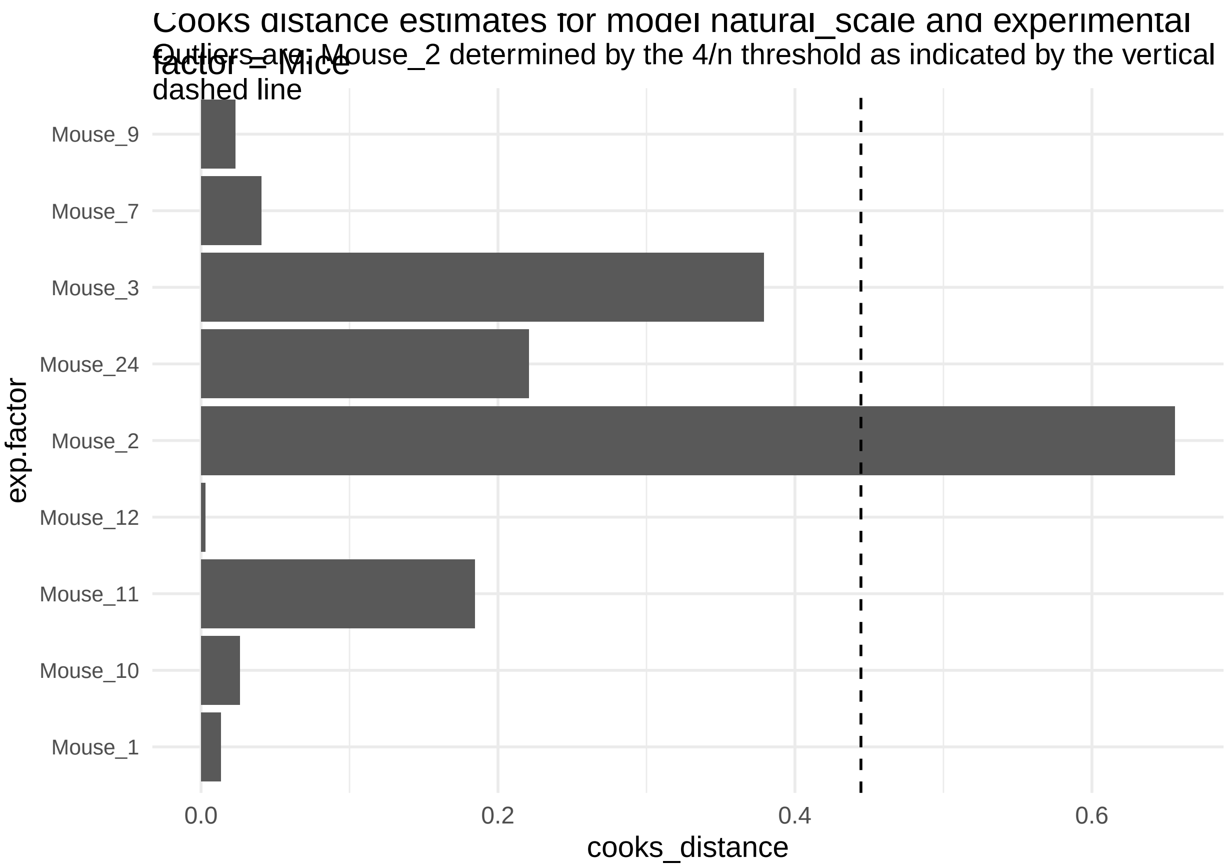
Cooks distance estimates for model natural_scale and experimental factor = Mice
Outliers are: Mouse_2 determined by the 4/n threshold as indicated by the vertical dashed line
Mouse_9
Mouse_7
Mouse_3
Mouse_24
Mouse_2
Mouse_12
Mouse_11
Mouse_10
Mouse_1
exp.factor
0.0
0.2
0.4
0.6
cooks_distance

**Figure 73:**

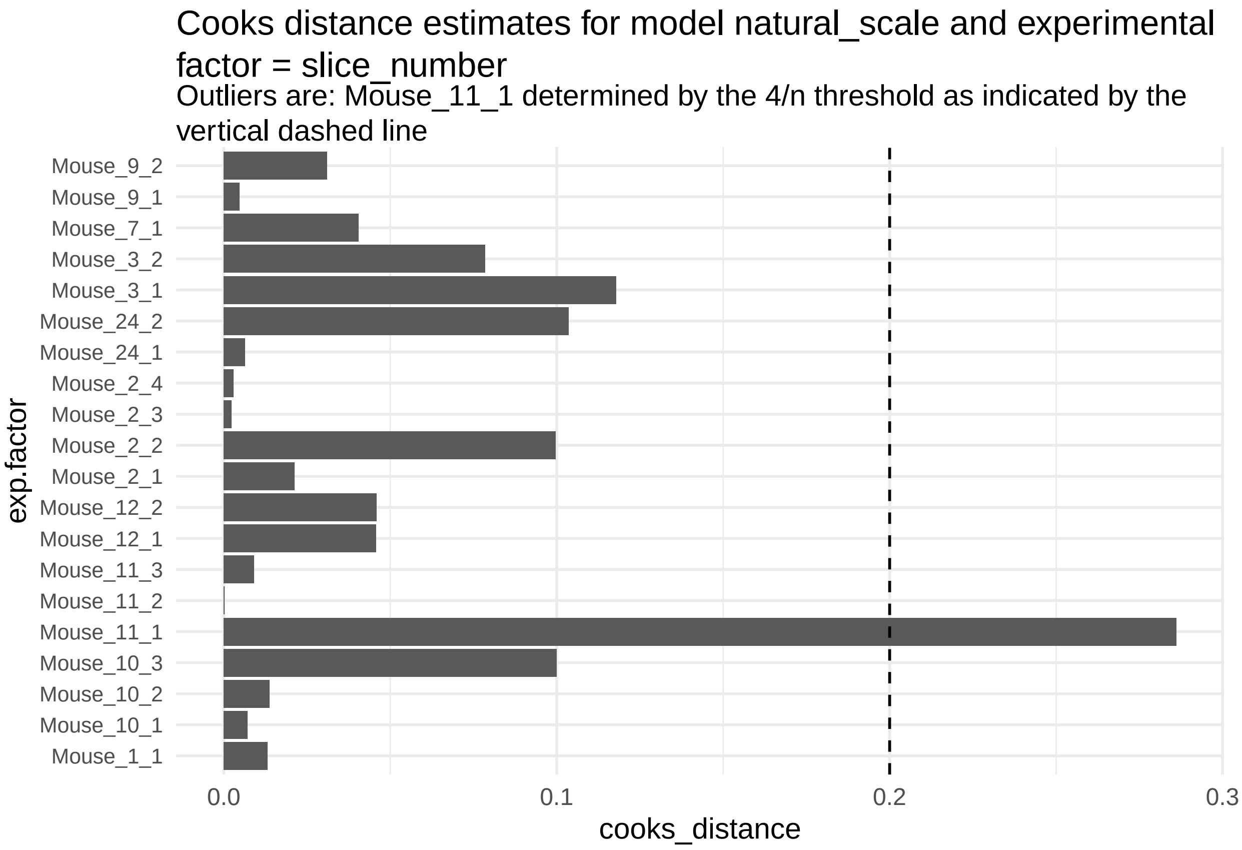


**Figure 74:**

```r
print("Inferred outlier groups are")
```

[1] "Inferred outlier groups are"

```r
print(diagnose_res$inferred_outlier_groups$natural_scale)
```

$Mice [1] "Mouse_2"

$slice_number [1] "Mouse_11_1"

Let us look the basic QC plots for the models fit in the natural scale of the responses without the outliers.

```r
plots <- diagnose_res$diagnostic_plots$natural_scale_wo_outliers$plots
captions <- diagnose_res$diagnostic_plots$natural_scale_wo_outliers$captions

for (i in seq_along(plots)) {
  print(plots[[i]])
    cat(sprintf("\n\n\n**Figure %d:** %s. %s\n\n", plot_no,names(plots)[i], captions[i]))
  plot_no <- plot_no + 1
}
```

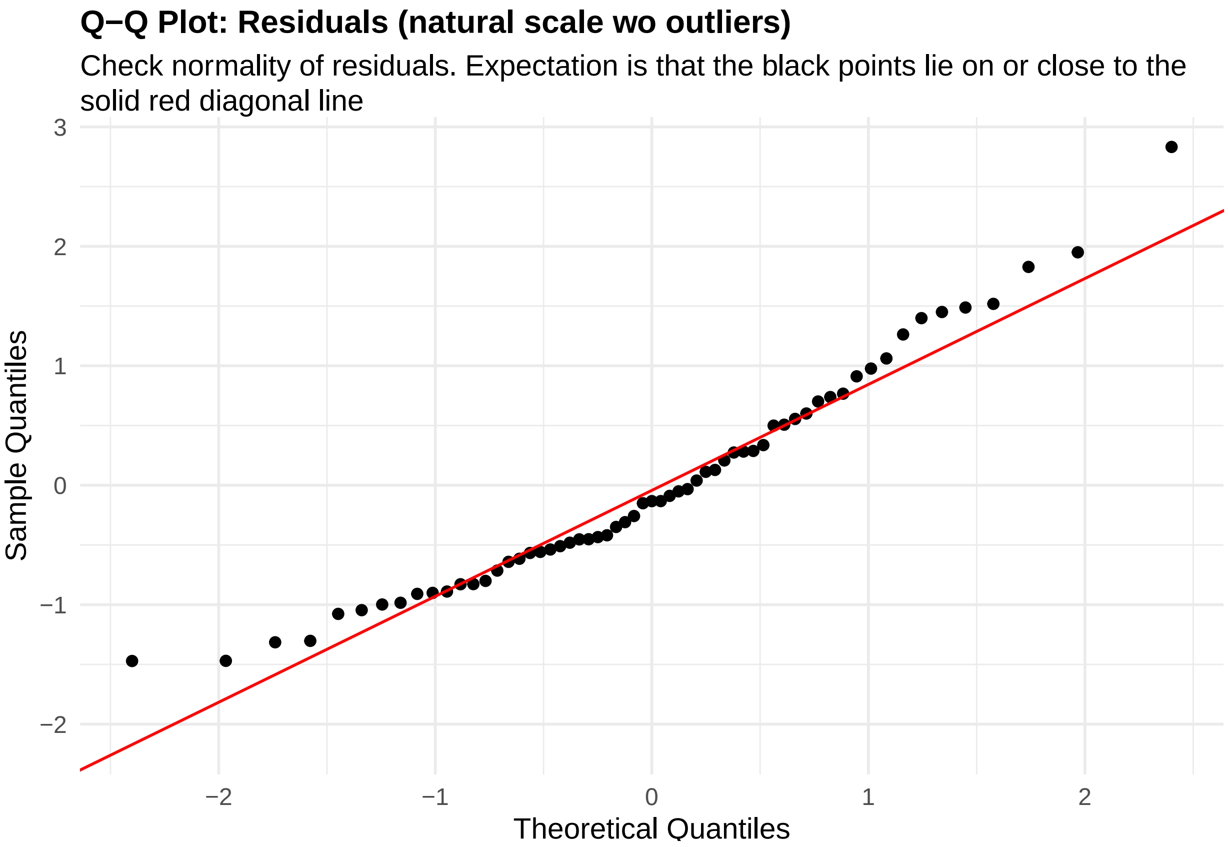


**Figure 75:** residuals_QQ. Check normality of residuals (natural scale wo outliers). Expectation is that the black points lie on or close to the solid red diagonal line. Check the Normal Q-Q plots for good and bad examples here: https://library.virginia.edu/data/articles/diagnostic-plots

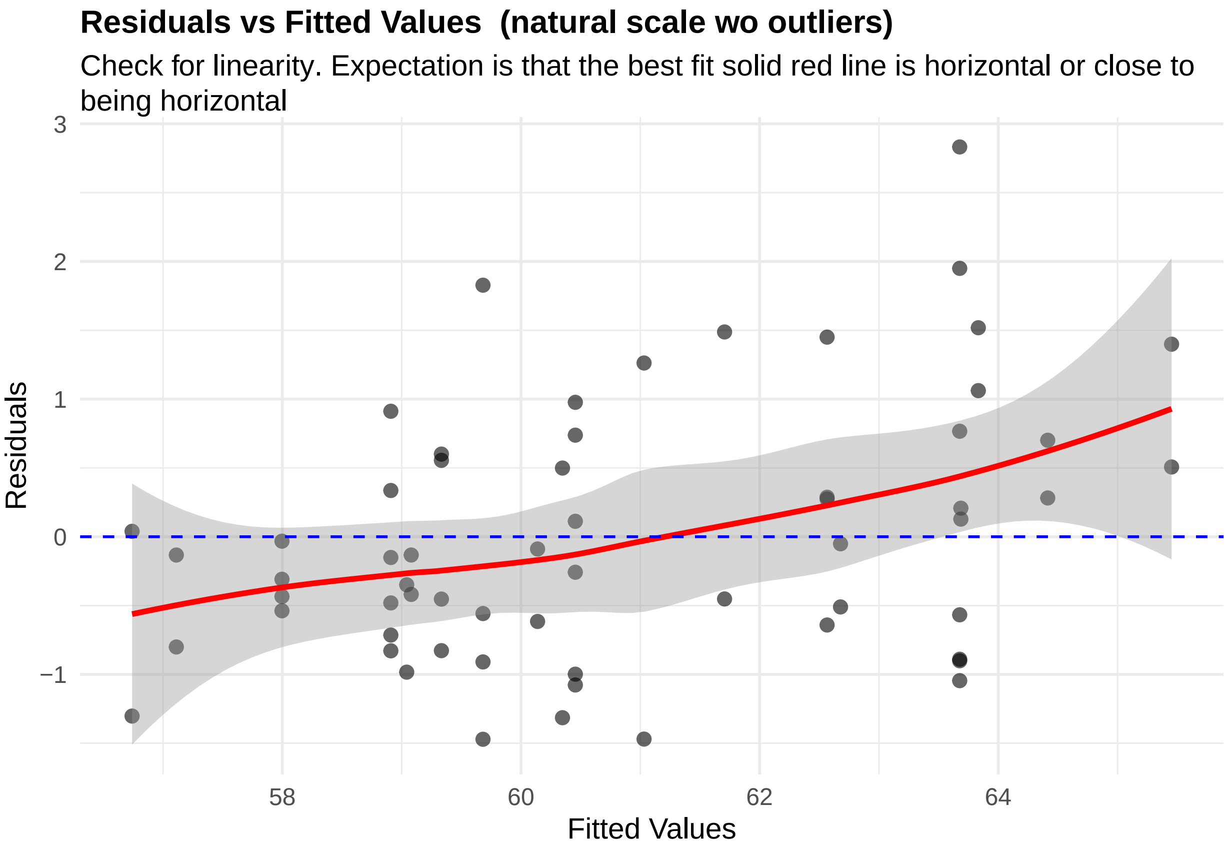


**Figure 76:** residuals_vs_fitted. Check for linearity (natural scale wo outliers). Expectation is that the best fit solid red line is horizontal or close to being horizontal. Check the Residuals vs Fitted plots for good and bad examples here: https://library.virginia.edu/data/articles/diagnostic-plots

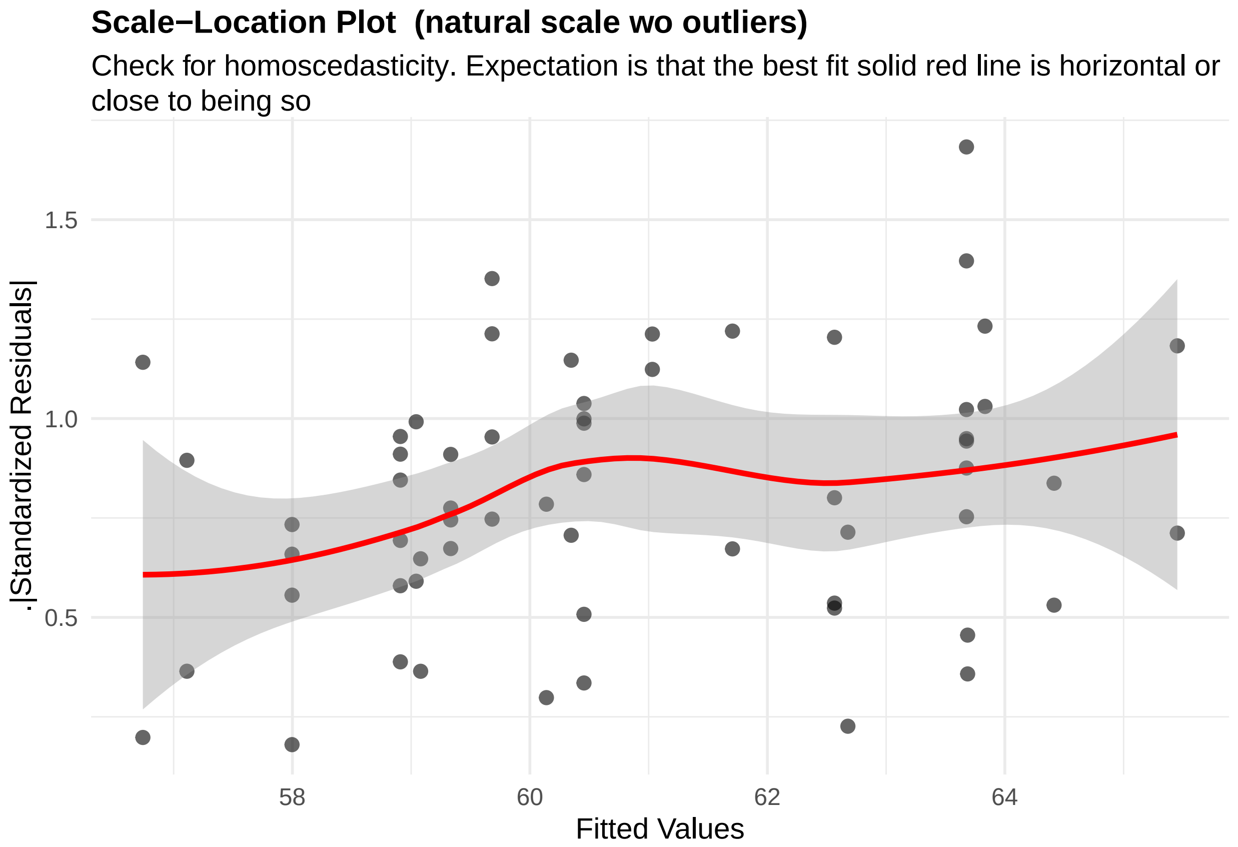


**Figure 77:** residuals_homoscedasticity. Check for homoscedasticity (natural scale wo outliers). Expectation is that the best fit solid red line is horizontal or close to being so. Check the Scale-Location plots for good and bad examples here: https://library.virginia.edu/data/articles/diagnostic-plots

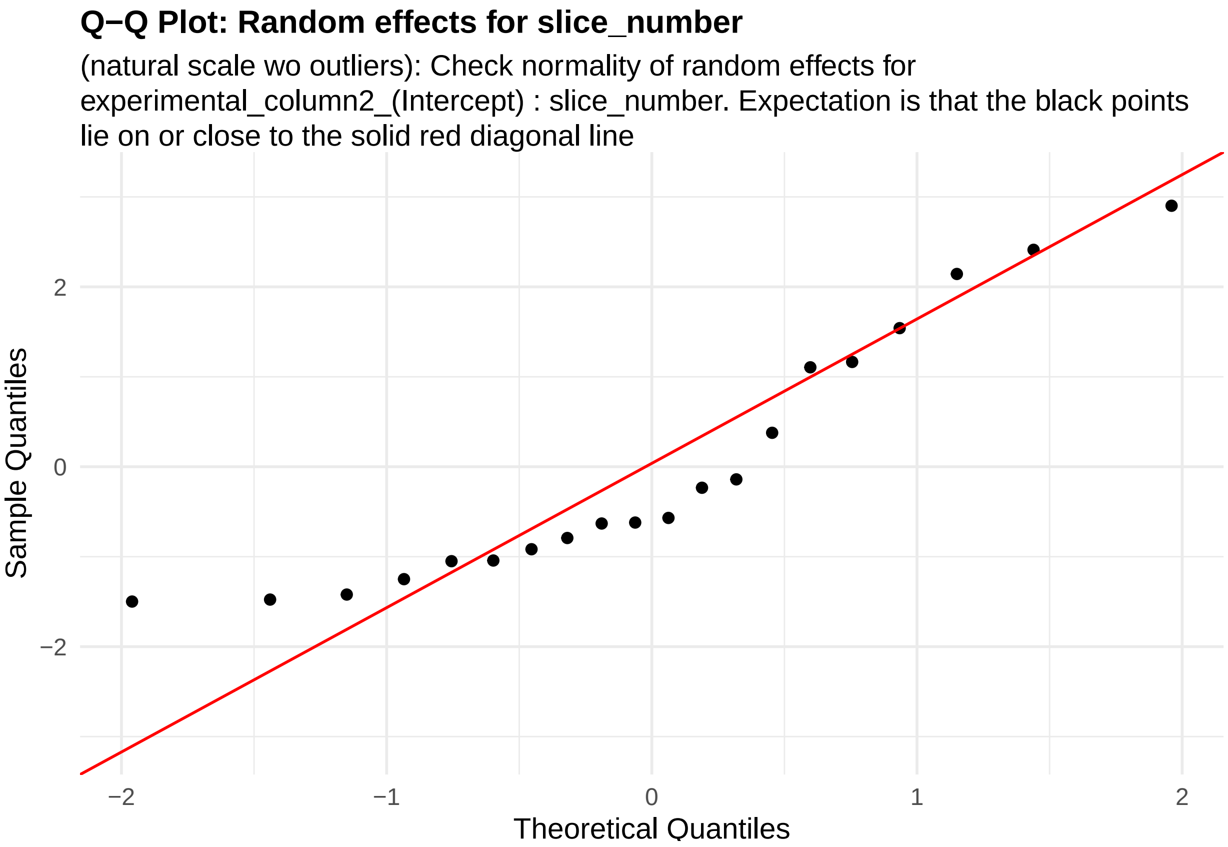


**Figure 78:** random_effects_QQ_1. Check normality of random effects for experimental_column2_(Intercept) (natural scale wo outliers): slice_number. Expectation is that the black points lie on or close to the solid red diagonal line. Check the Normal Q-Q plots for good and bad examples here: https://library.virginia.edu/data/articles/diagnostic-plots

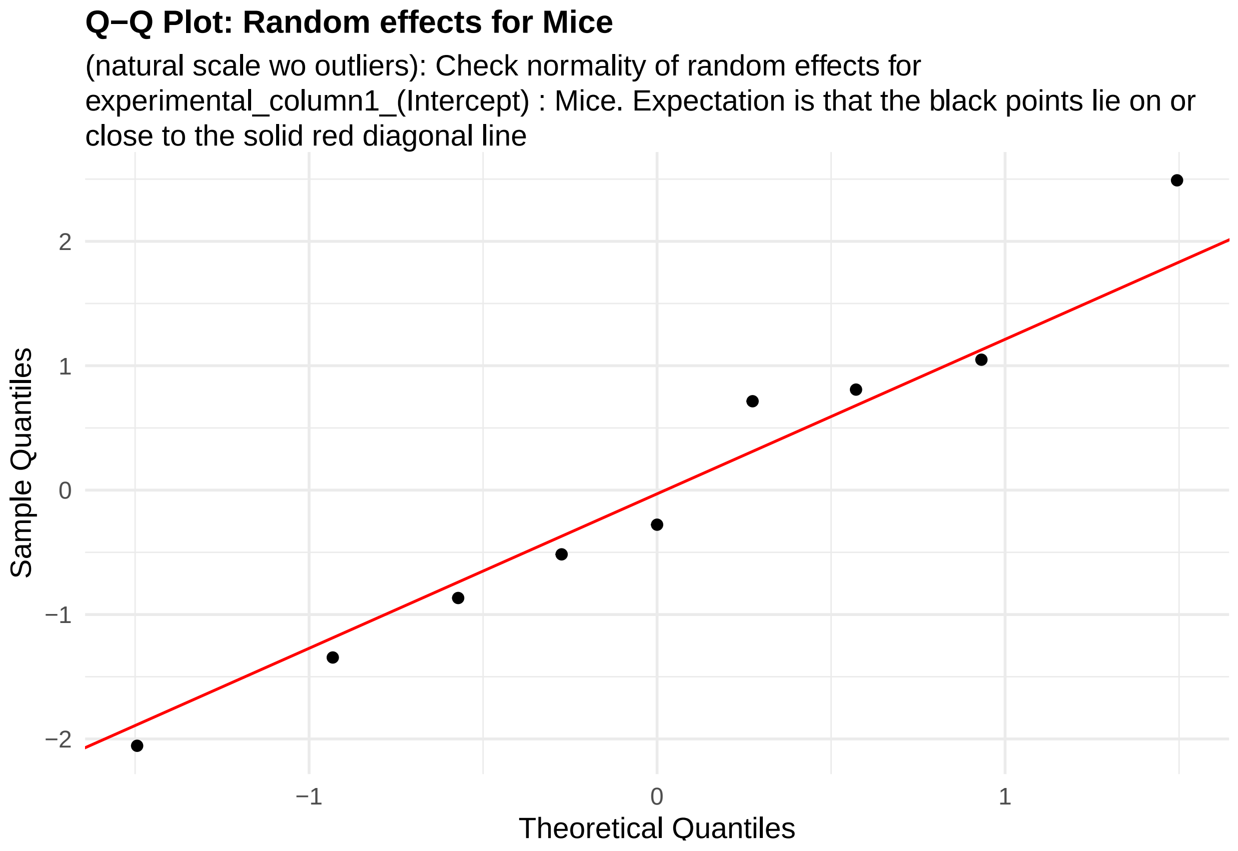


**Figure 79:** random_effects_QQ_2. Check normality of random effects for experimental_column1_(Intercept) (natural scale wo outliers): Mice. Expectation is that the black points lie on or close to the solid red diagonal line. Check the Normal Q-Q plots for good and bad examples here: https://library.virginia.edu/data/articles/diagnostic-plots

```
cooks_plots <- diagnose_res$cooks_plots$natural_scale_wo_outliers
cat("\n\n\n")
```

```
for(i in 1:length(cooks_plots)) {
  print(cooks_plots[[i]])
  cat(sprintf("\n\n\n**Figure %d:**\n\n", plot_no))
  plot_no <- plot_no + 1
}
```

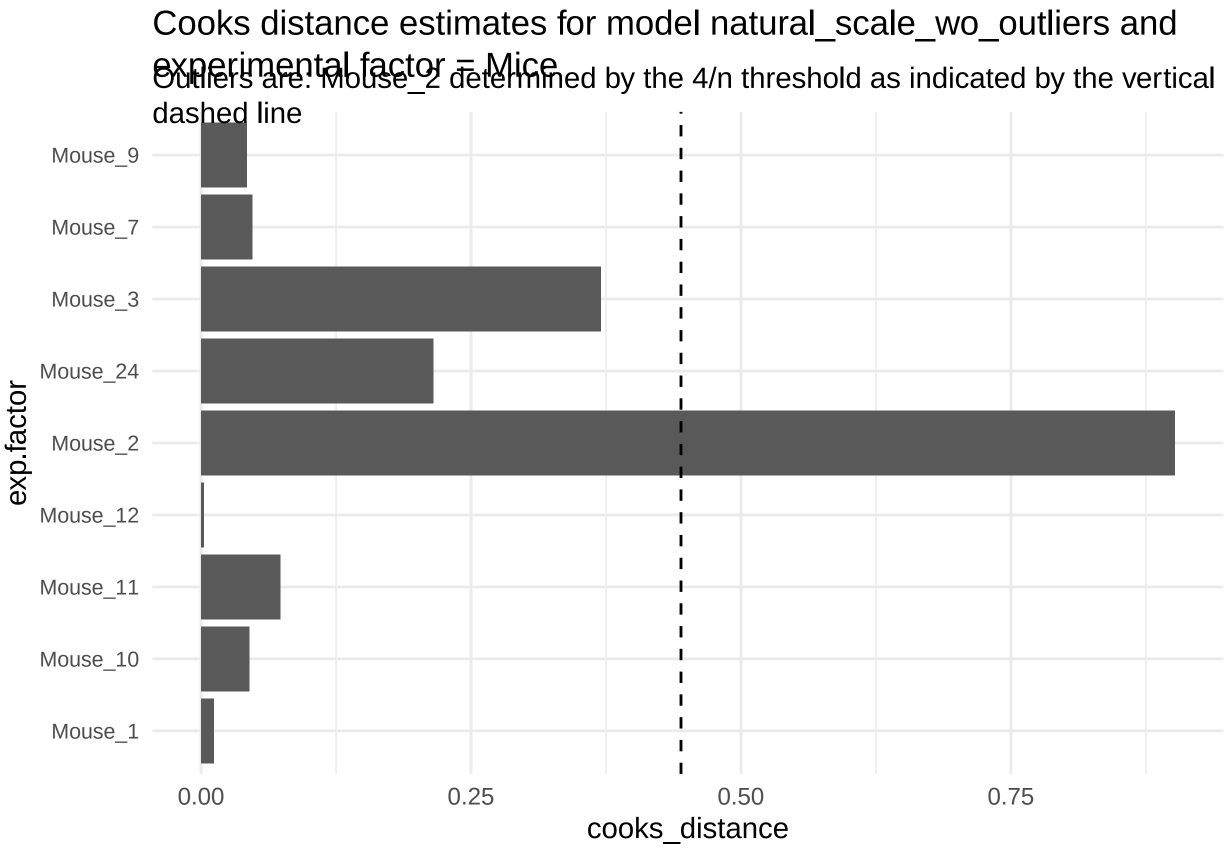
Cooks distance estimates for model natural_scale_wo_outliers and experimental factor = Mice
Outliers are: Mouse_2 determined by the 4/n threshold as indicated by the vertical dashed line
Mouse_9
Mouse_7
Mouse_3
Mouse_24
Mouse_2
Mouse_12
Mouse_11
Mouse_10
Mouse_1
exp.factor
0.00
0.25
0.50
0.75
cooks_distance

**Figure 80:**

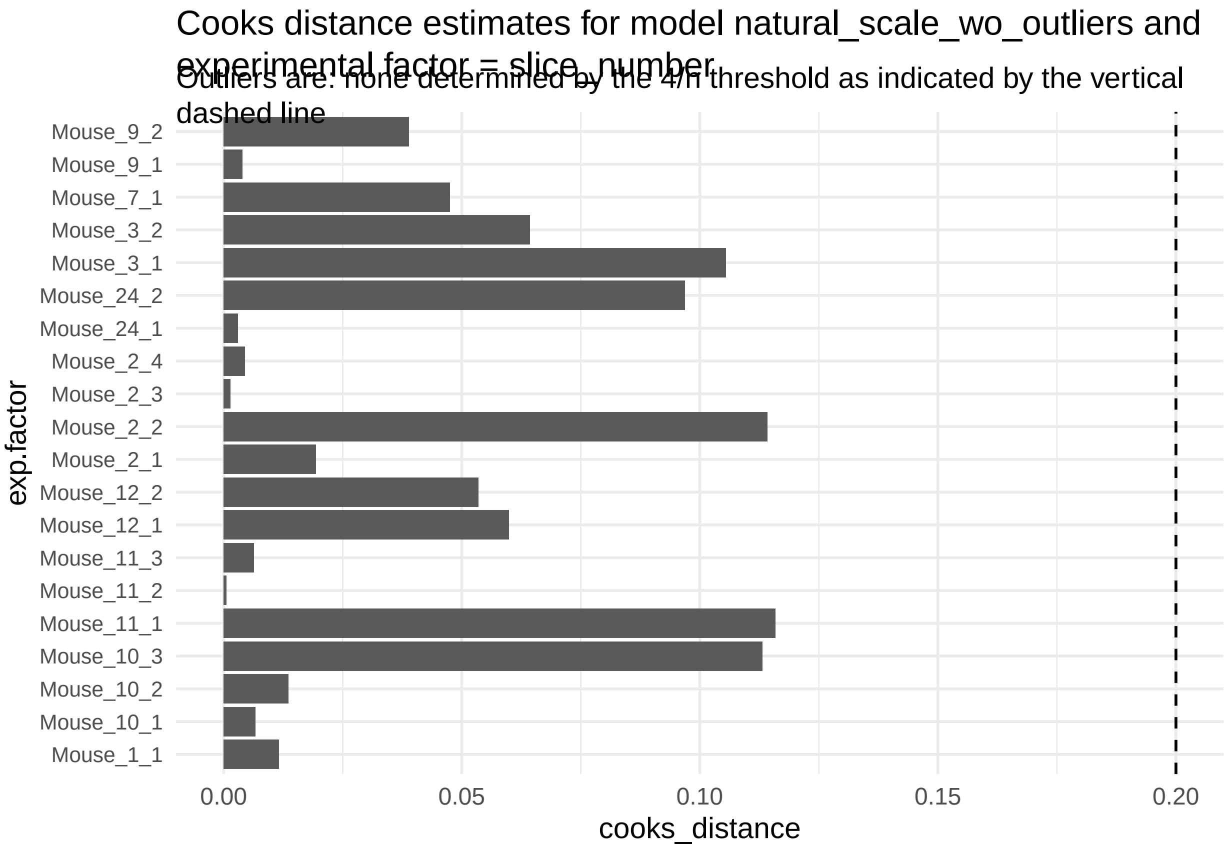


**Figure 81:**

```r
print("Inferred outlier groups are")
```

[1] "Inferred outlier groups are"

```r
print(diagnose_res$inferred_outlier_groups$natural_scale)
```

$Mice [1] "Mouse_2"

$slice_number [1] "Mouse_11_1"

Now, let us look at the same set of QC plots for models on the log-transformed responses.

```r
plots <- diagnose_res$diagnostic_plots$log_scale$plots
captions <- diagnose_res$diagnostic_plots$log_scale$captions
for (i in seq_along(plots)) {
  print(plots[[i]])
    cat(sprintf("\n\n\n**Figure %d:** %s. %s\n\n", plot_no,names(plots)[i], captions[i]))
  plot_no <- plot_no + 1
}
```

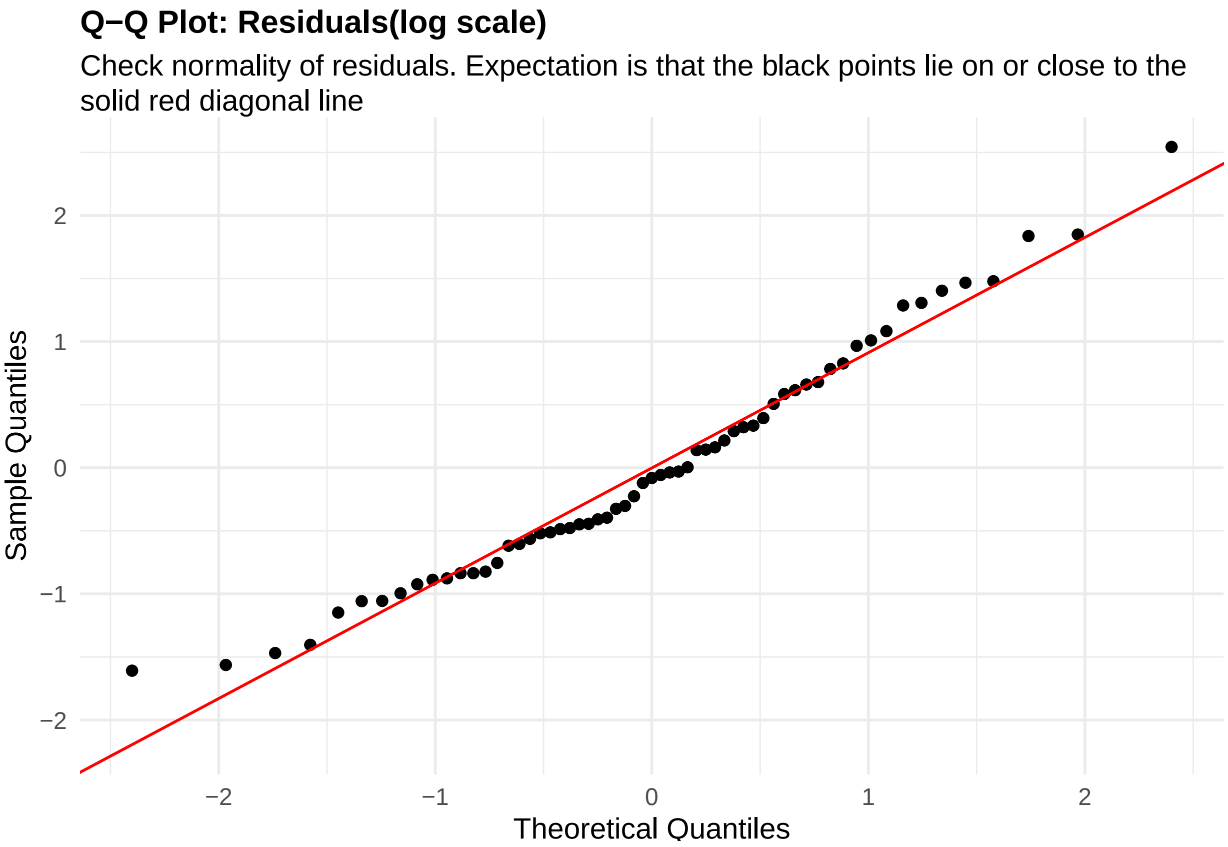


**Figure 82:** residuals_QQ. Check normality of residuals (log scale). Expectation is that the black points lie on or close to the solid red diagonal line. Check the Normal Q-Q plots for good and bad examples here: https://library.virginia.edu/data/articles/diagnostic-plots

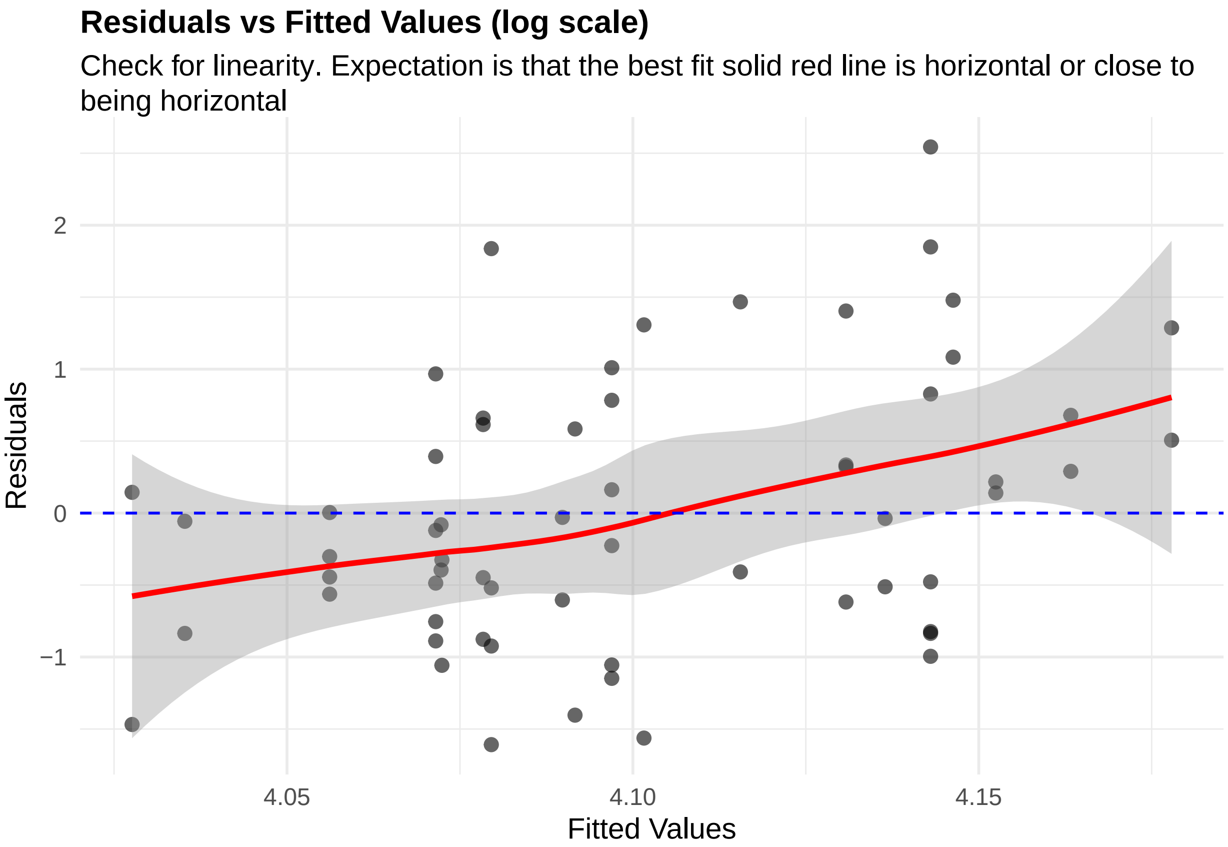


**Figure 83:** residuals_vs_fitted. Check for linearity (log scale). Expectation is that the best fit solid red line is horizontal or close to being horizontal. Check the Residuals vs Fitted plots for good and bad examples here: https://library.virginia.edu/data/articles/diagnostic-plots

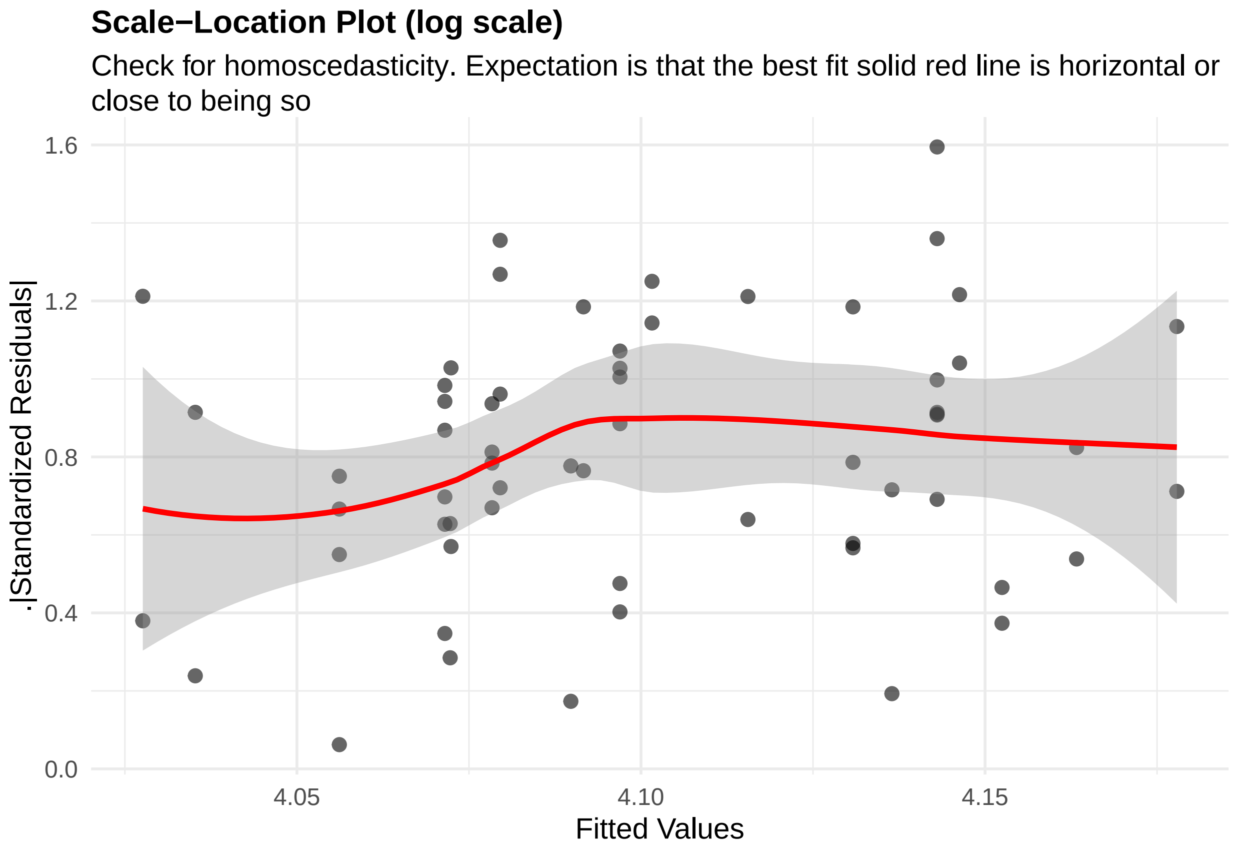


**Figure 84:** residuals_homoscedasticity. Check for homoscedasticity (log scale). Expectation is that the best fit solid red line is horizontal or close to being so. Check the Scale-Location plots for good and bad examples here: https://library.virginia.edu/data/articles/diagnostic-plots

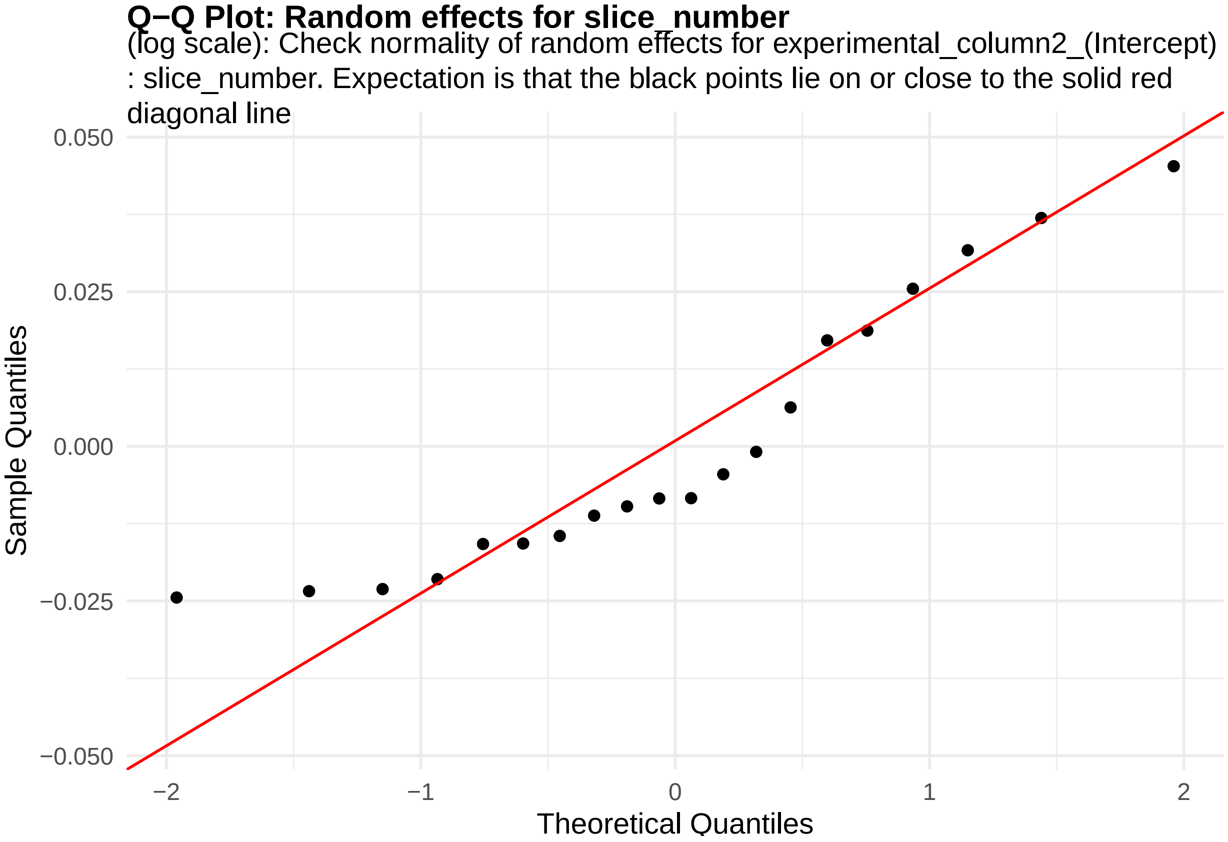


**Figure 85:** random_effects_QQ_1. Check normality of random effects for experimental_column2_(Intercept) (log scale): slice_number. Expectation is that the black points lie on or close to the solid red diagonal line. Check the Normal Q-Q plots for good and bad examples here: https://library.virginia.edu/data/articles/diagnostic-plots

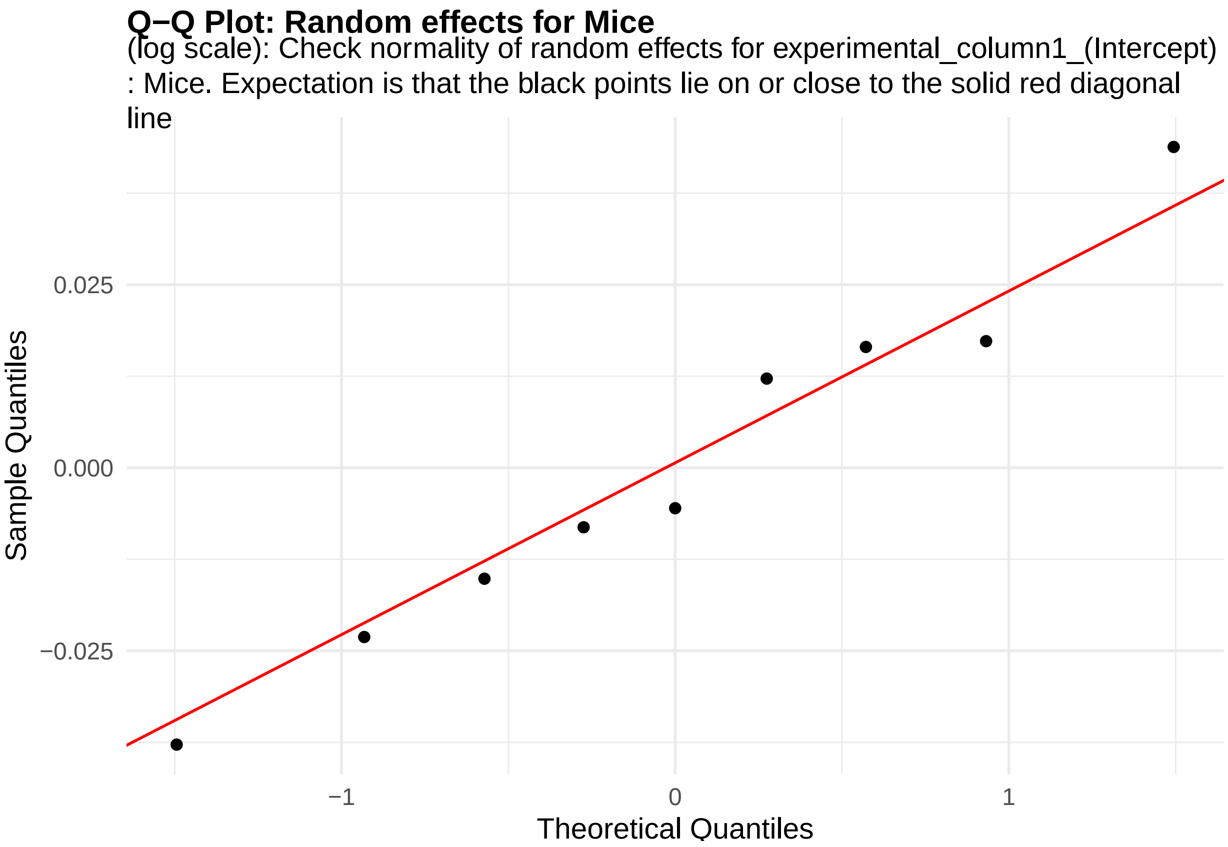


**Figure 86:** random_effects_QQ_2. Check normality of random effects for experimental_column1_(Intercept) (log scale): Mice. Expectation is that the black points lie on or close to the solid red diagonal line. Check the Normal Q-Q plots for good and bad examples here: https://library.virginia.edu/data/articles/diagnostic-plots

```
cooks_plots <- diagnose_res$cooks_plots$log_scale
cat("\n\n\n")
```

```
for(i in 1:length(cooks_plots)) {
  print(cooks_plots[[i]])
  cat(sprintf("\n\n\n**Figure %d:**\n\n", plot_no))
  plot_no <- plot_no + 1
}
```

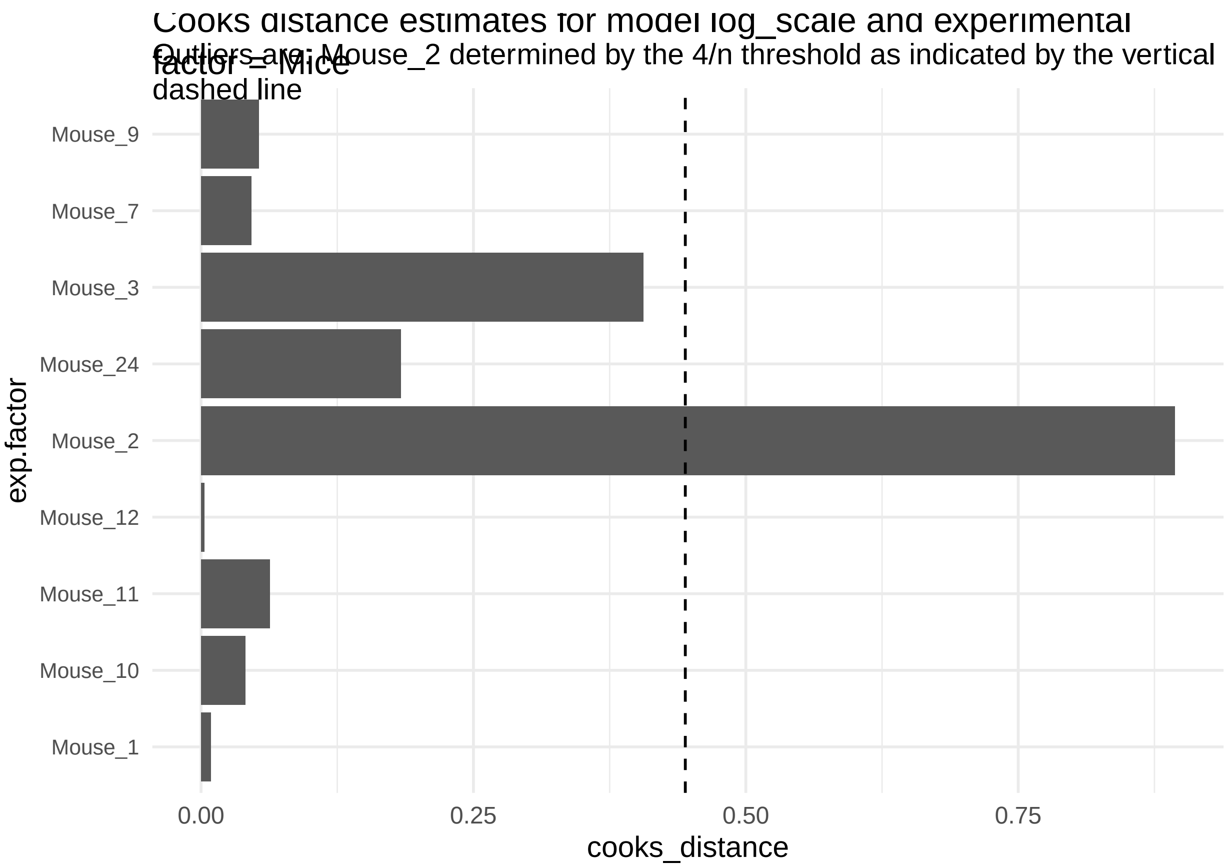
Cooks distance estimates for model log_scale and experimental factor = Mice
Outliers are: Mouse_2 determined by the 4/n threshold as indicated by the vertical dashed line
exp.factor
Mouse_9
Mouse_7
Mouse_3
Mouse_24
Mouse_2
Mouse_12
Mouse_11
Mouse_10
Mouse_1
0.00
0.25
0.50
0.75
cooks_distance

**Figure 87:**

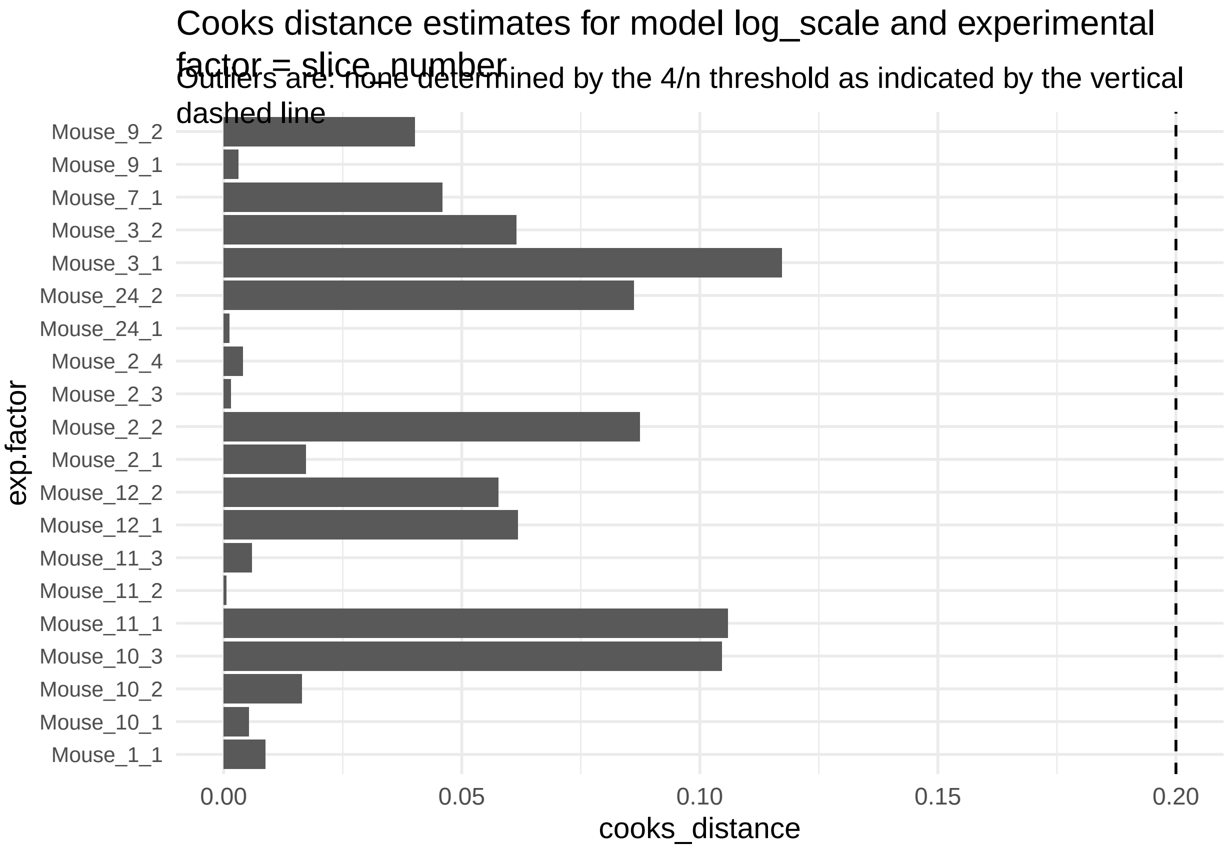


**Figure 88:**

```
print("Inferred outlier groups are")
```

[1] "Inferred outlier groups are"

```
print(diagnose_res$inferred_outlier_groups$natural_scale)
```

$Mice [1] "Mouse_2"

$slice_number [1] "Mouse_11_1"

We will work with the model of the electrical responses in the natural scale without outliers

The code also outputs Cook's distance measures for each cohort and mouse that is intended to capture how influential each of these were in determining the final parameter estimates in the LMM model.

Mouse_2 appears to be an outlier based on its cook's distance estimate (of 0.90) from the natural scale without outliers model and a 4/n=4/9=0.44 threshold.

We will therefore drop all observations from Mouse_2.

```
data_update <- diagnose_res$Data_updated
data_update %<>% filter(Mice != "Mouse_2")
design_update <- design
design_update@response_column %<>% paste0(., "_noOutlier")
diagnose_res_update <- diagnoseDataModel(data_update, design_update, model)
```

```
## [1] "No outlier detected from the raw Data"
```

```
## [1] "No outlier detected from the raw Data"
```

To identify all the models fit to the data

```r
print(diagnose_res_update$models)
```

```
## [1] "Natural scale" "Log scale"
```

We see that two models were fit to this data - one in the natural scale of negative_action_potential_mV, other in the log scale.

Let us look the basic QC plots for the models fit in the natural scale of the responses.

```r
plots <- diagnose_res_update$diagnostic_plots$natural_scale$plots
captions <- diagnose_res_update$diagnostic_plots$natural_scale$captions
for (i in seq_along(plots)) {
  print(plots[[i]])
    cat(sprintf("\n\n\n**Figure %d:** %s. %s\n\n", plot_no,names(plots)[i], captions[i]))
  plot_no <- plot_no + 1
}
```

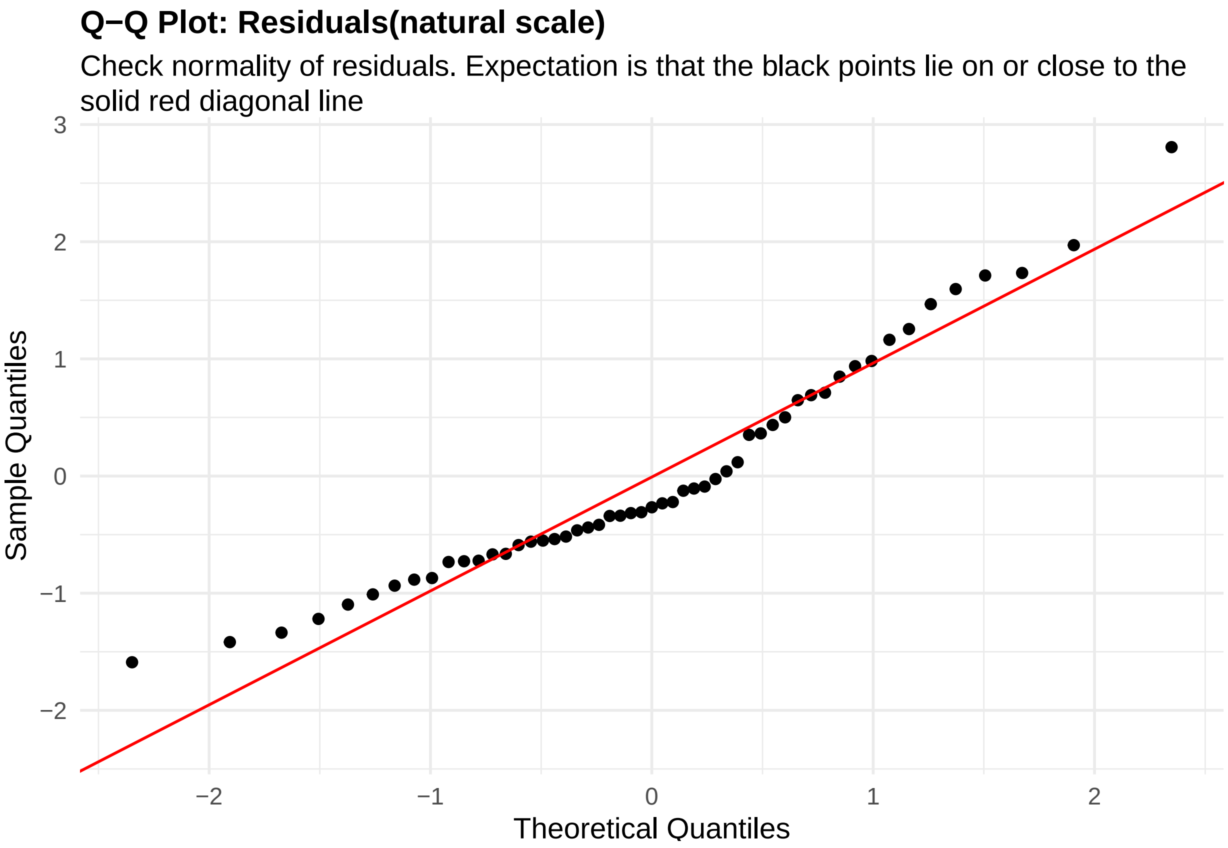


**Figure 89:** residuals_QQ. Check normality of residuals (natural scale). Expectation is that the black points lie on or close to the solid red diagonal line. Check the Normal Q-Q plots for good and bad examples here: https://library.virginia.edu/data/articles/diagnostic-plots

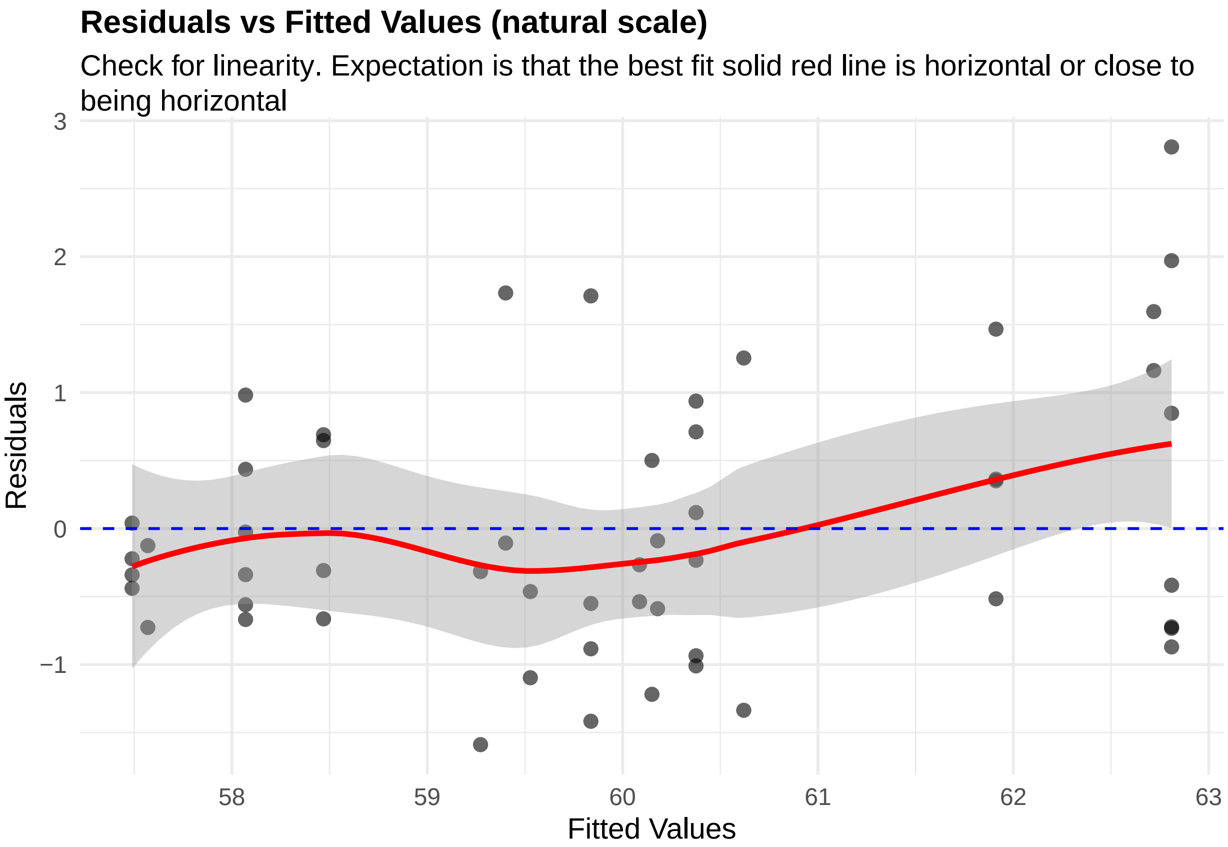


**Figure 90:** residuals_vs_fitted. Check for linearity (natural scale). Expectation is that the best fit solid red line is horizontal or close to being horizontal. Check the Residuals vs Fitted plots for good and bad examples here: https://library.virginia.edu/data/articles/diagnostic-plots

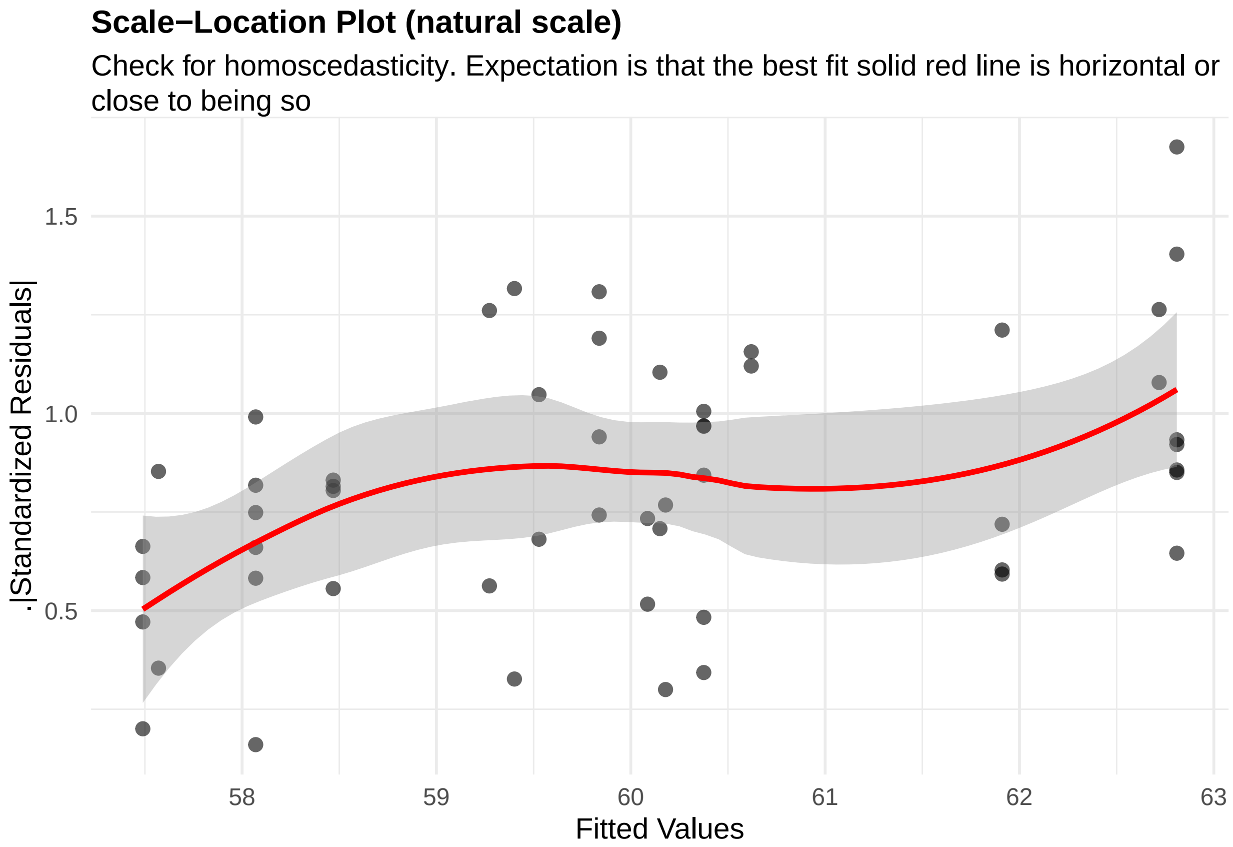


**Figure 91:** residuals_homoscedasticity. Check for homoscedasticity (natural scale). Expectation is that the best fit solid red line is horizontal or close to being so. Check the Scale-Location plots for good and bad examples here: https://library.virginia.edu/data/articles/diagnostic-plots

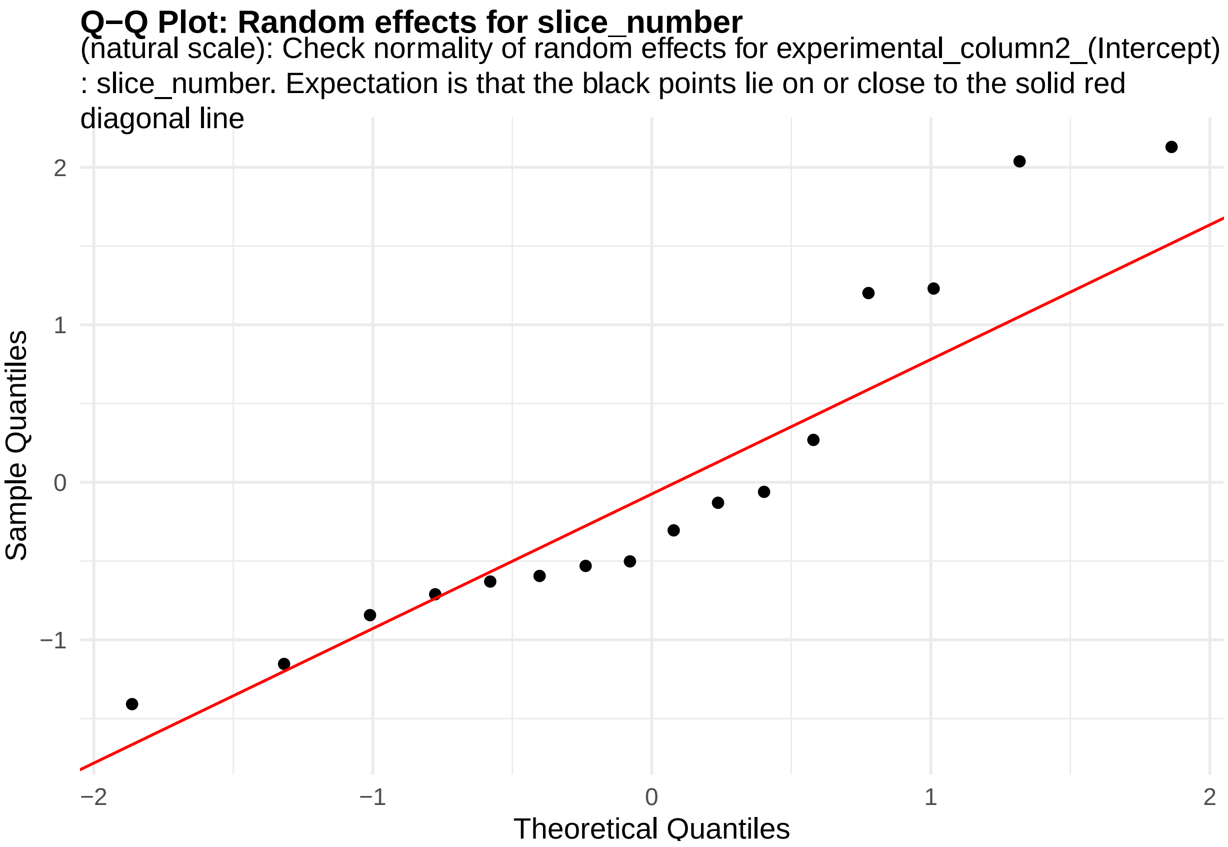


**Figure 92:** random_effects_QQ_1. Check normality of random effects for experimental_column2_(Intercept) (natural scale): slice_number. Expectation is that the black points lie on or close to the solid red diagonal line. Check the Normal Q-Q plots for good and bad examples here: https://library.virginia.edu/data/articl es/diagnostic-plots

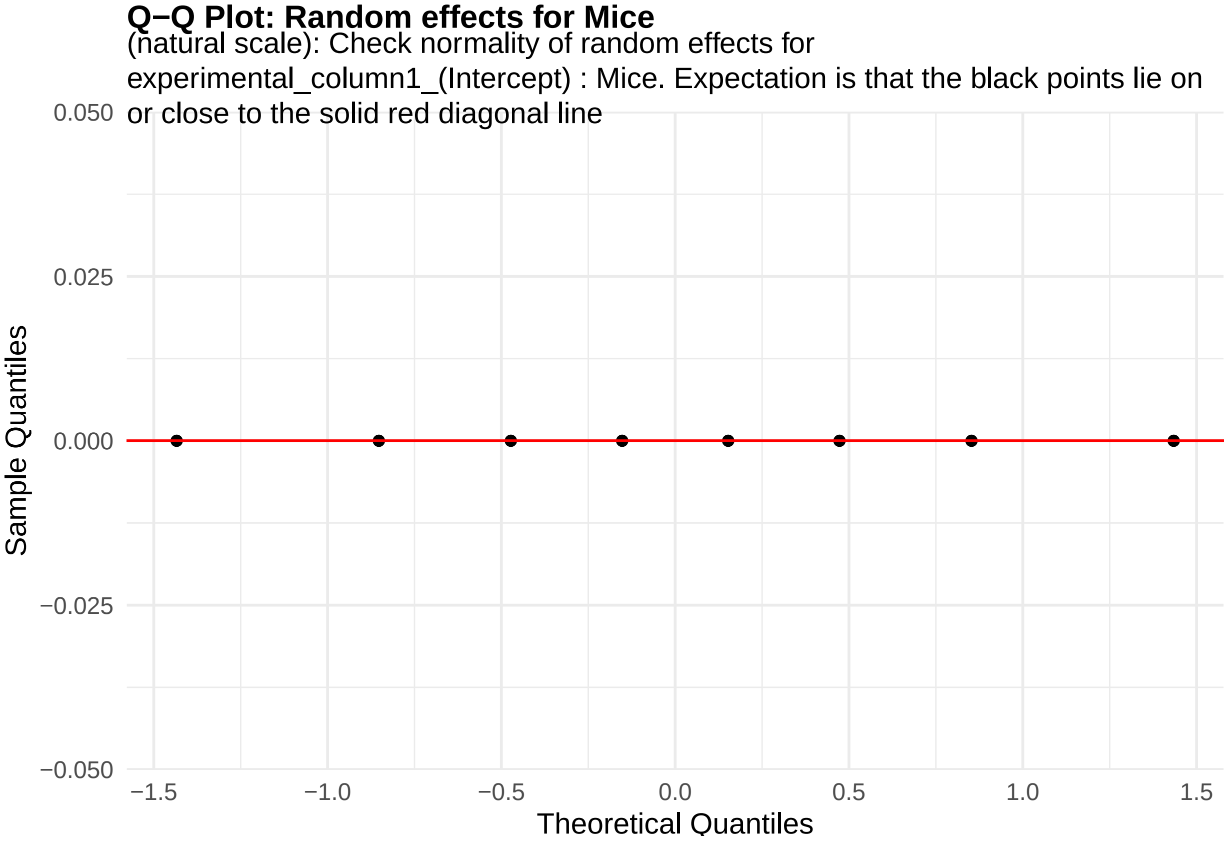


**Figure 93:** random_effects_QQ_2. Check normality of random effects for experimental_column1_(Intercept) (natural scale): Mice. Expectation is that the black points lie on or close to the solid red diagonal line. Check the Normal Q-Q plots for good and bad examples here: https://library.virginia.edu/data/articles/diagnostic-plots

```
cooks_plots <- diagnose_res_update$cooks_plots$natural_scale
cat("\n\n\n")
```

```
for(i in 1:length(cooks_plots)) {
  print(cooks_plots[[i]])
  cat(sprintf("\n\n\n**Figure %d:**\n\n", plot_no))
  plot_no <- plot_no + 1
}
```

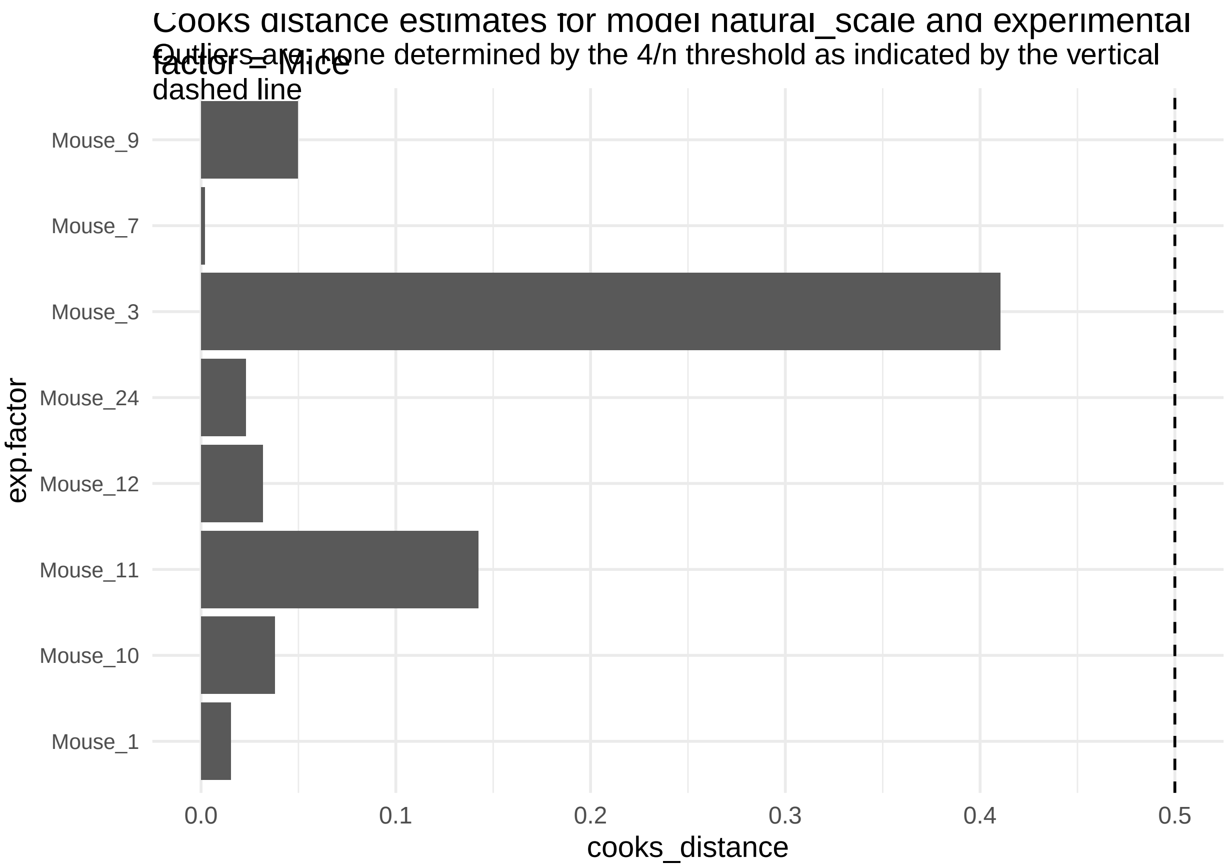
Cooks distance estimates for model natural_scale and experimental factor = Mice
Outliers are: none determined by the 4/n threshold as indicated by the vertical dashed line
Mouse_9
Mouse_7
Mouse_3
Mouse_24
Mouse_12
Mouse_11
Mouse_10
Mouse_1
exp.factor
0.0
0.1
0.2
0.3
0.4
0.5
cooks_distance

**Figure 94:**

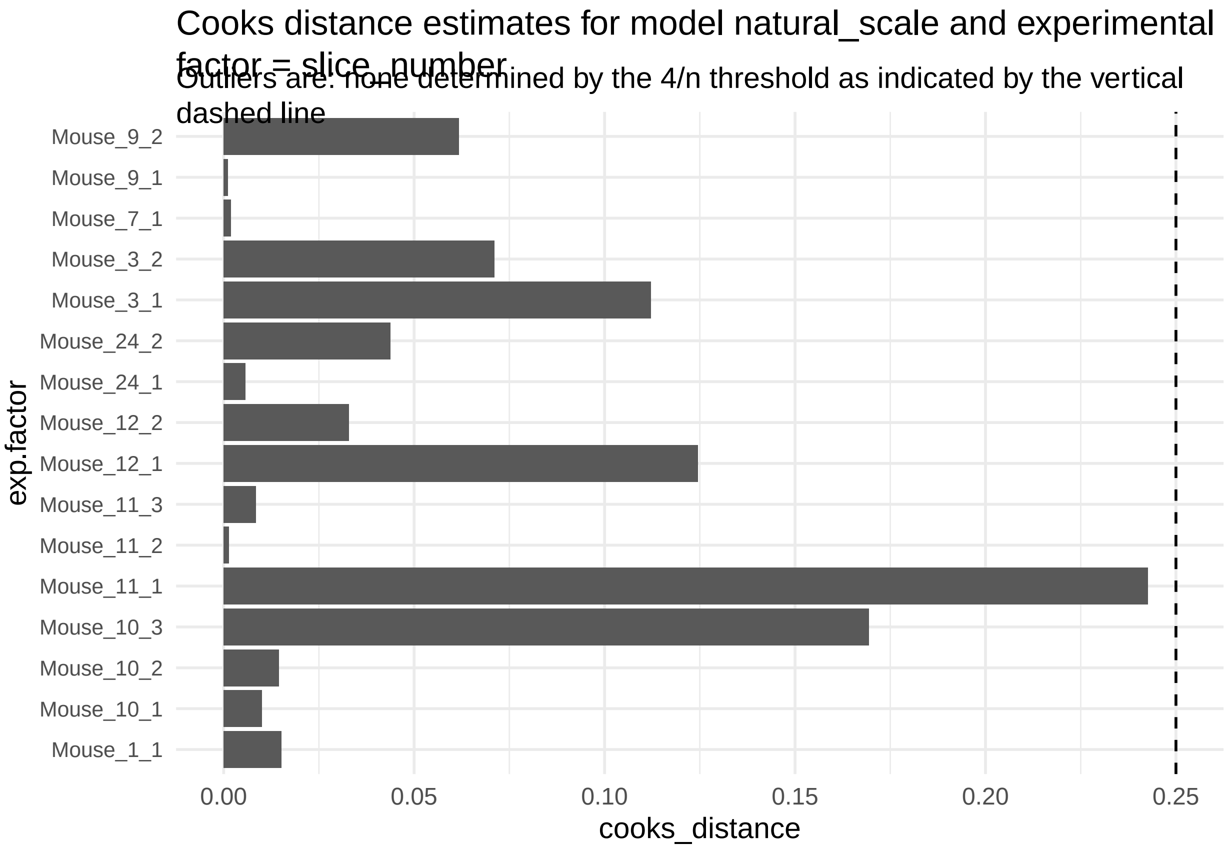


**Figure 95:**

```
print("Inferred outlier groups are")
```

[1] "Inferred outlier groups are"

```
print(diagnose_res$inferred_outlier_groups$natural_scale)
```

$Mice [1] "Mouse_2"

$slice_number [1] "Mouse_11_1"

Let us look the basic QC plots for the models fit in the log scale of the responses.

```
plots <- diagnose_res_update$diagnostic_plots$log_scale$plots
captions <- diagnose_res_update$diagnostic_plots$log_scale$captions
for (i in seq_along(plots)) {
  print(plots[[i]])
    cat(sprintf("\n\n\n**Figure %d:** %s. %s\n\n", plot_no,names(plots)[i], captions[i]))
  plot_no <- plot_no + 1
}
```

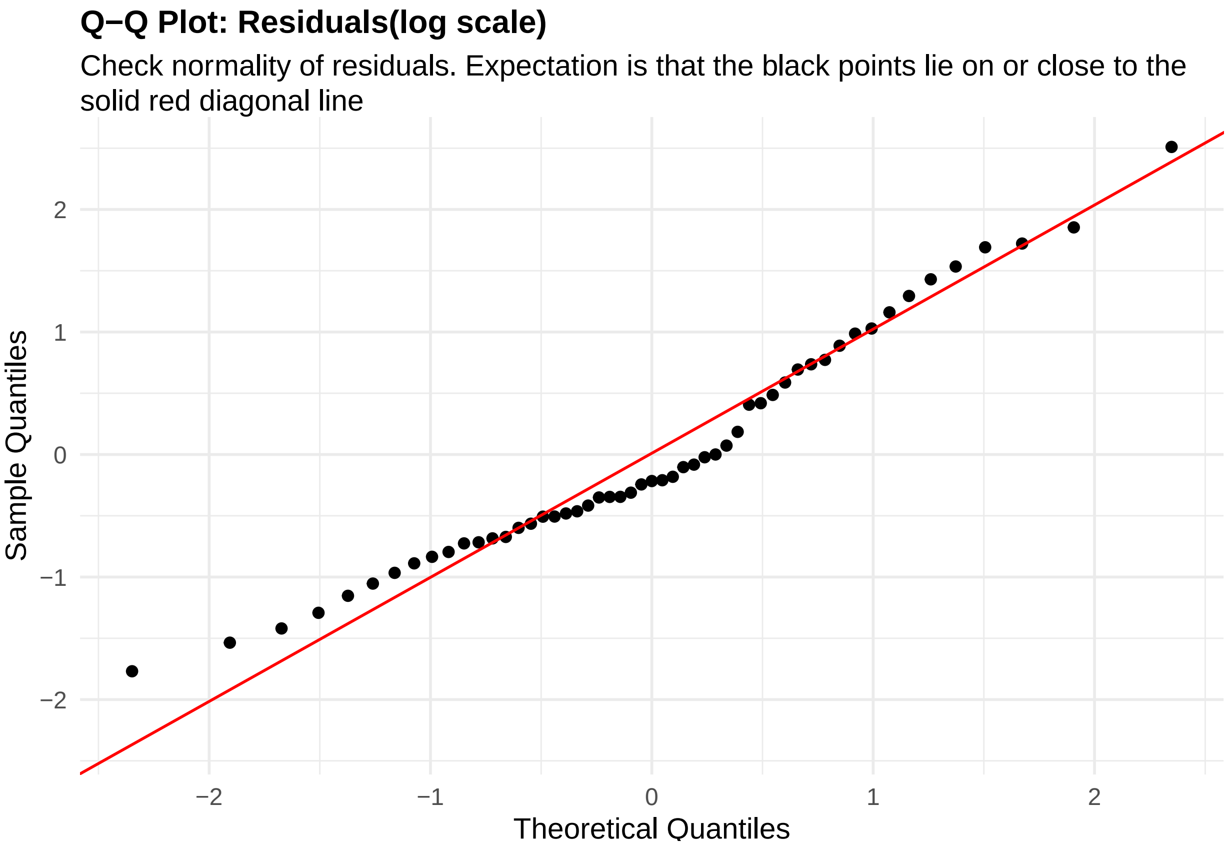


**Figure 96:** residuals_QQ. Check normality of residuals (log scale). Expectation is that the black points lie on or close to the solid red diagonal line. Check the Normal Q-Q plots for good and bad examples here: https://library.virginia.edu/data/articles/diagnostic-plots

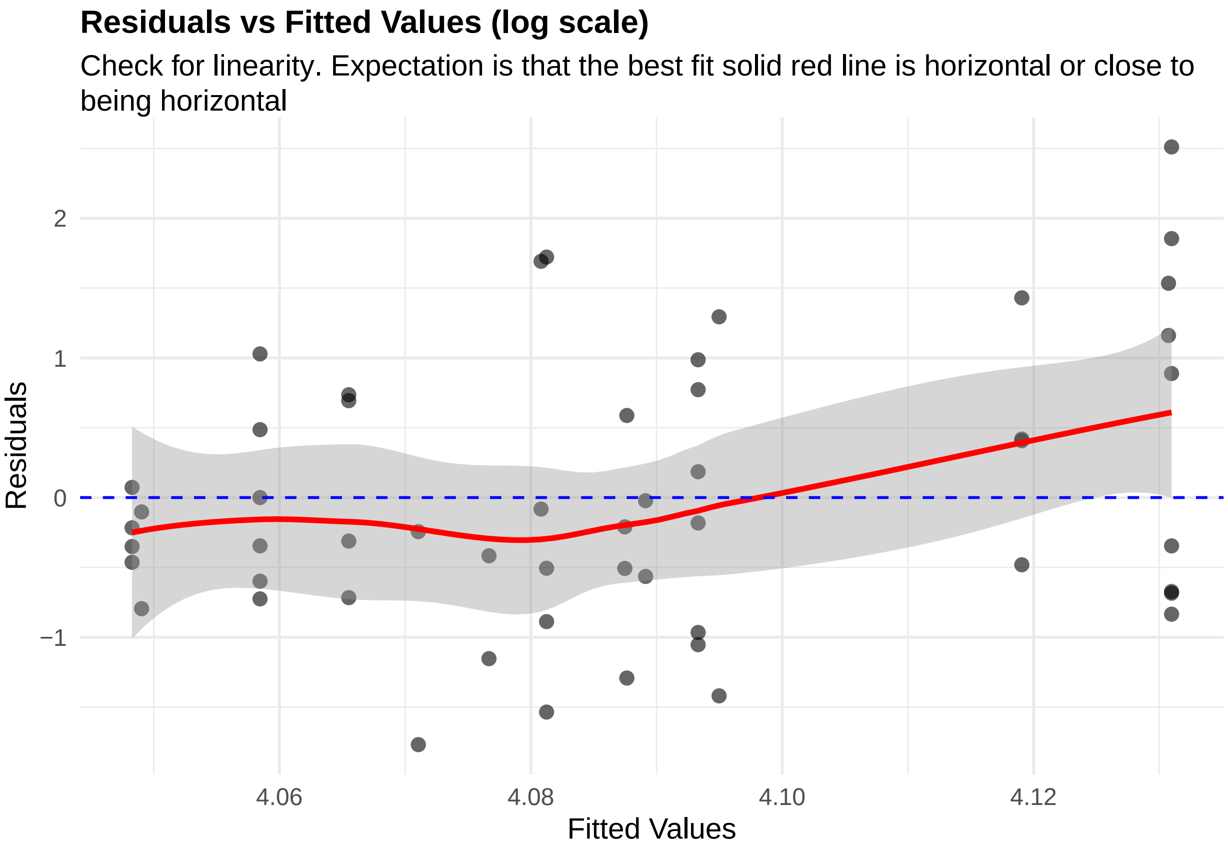


**Figure 97:** residuals_vs_fitted. Check for linearity (log scale). Expectation is that the best fit solid red line is horizontal or close to being horizontal. Check the Residuals vs Fitted plots for good and bad examples here: https://library.virginia.edu/data/articles/diagnostic-plots

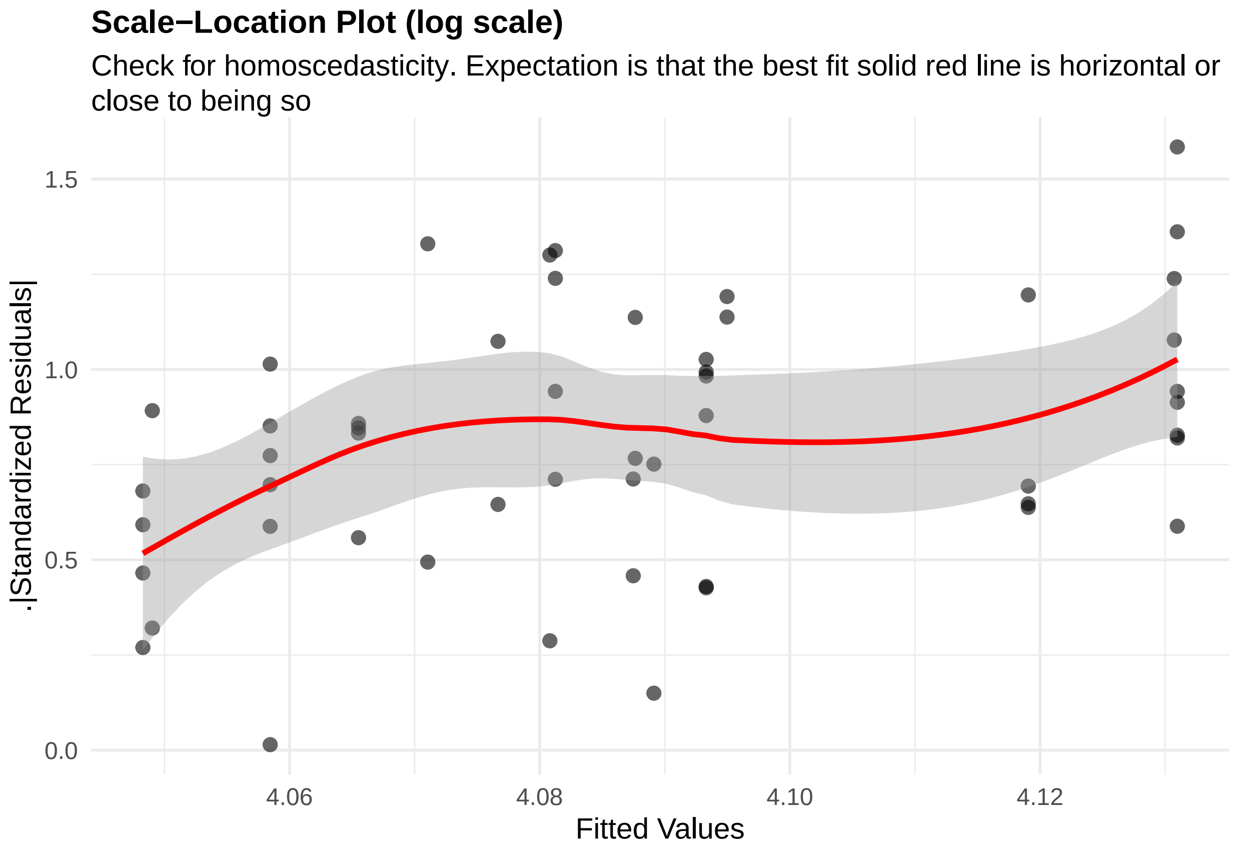


**Figure 98:** residuals_homoscedasticity. Check for homoscedasticity (log scale). Expectation is that the best fit solid red line is horizontal or close to being so. Check the Scale-Location plots for good and bad examples here: https://library.virginia.edu/data/articles/diagnostic-plots

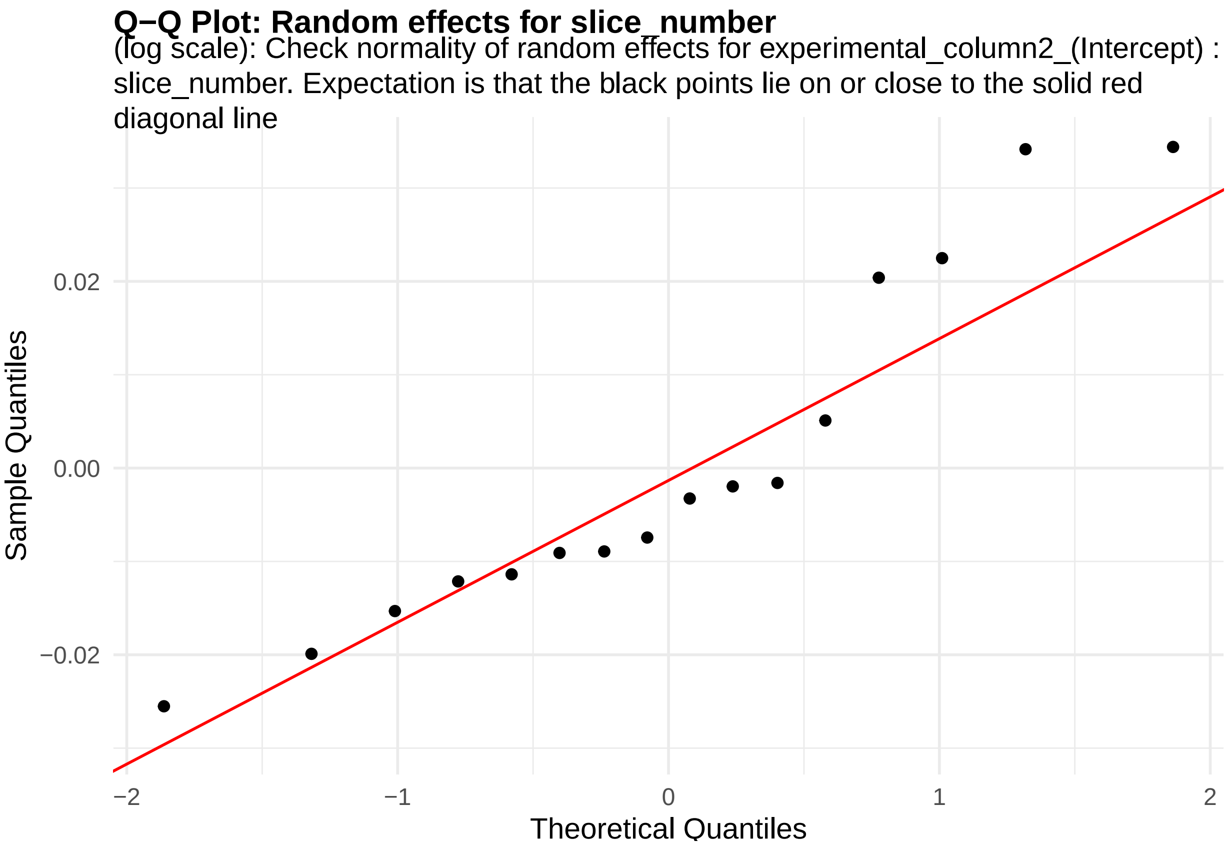


**Figure 99:** random_effects_QQ_1. Check normality of random effects for experimental_column2_(Intercept) (log scale): slice_number. Expectation is that the black points lie on or close to the solid red diagonal line. Check the Normal Q-Q plots for good and bad examples here: https://library.virginia.edu/data/articles/diagnostic-plots

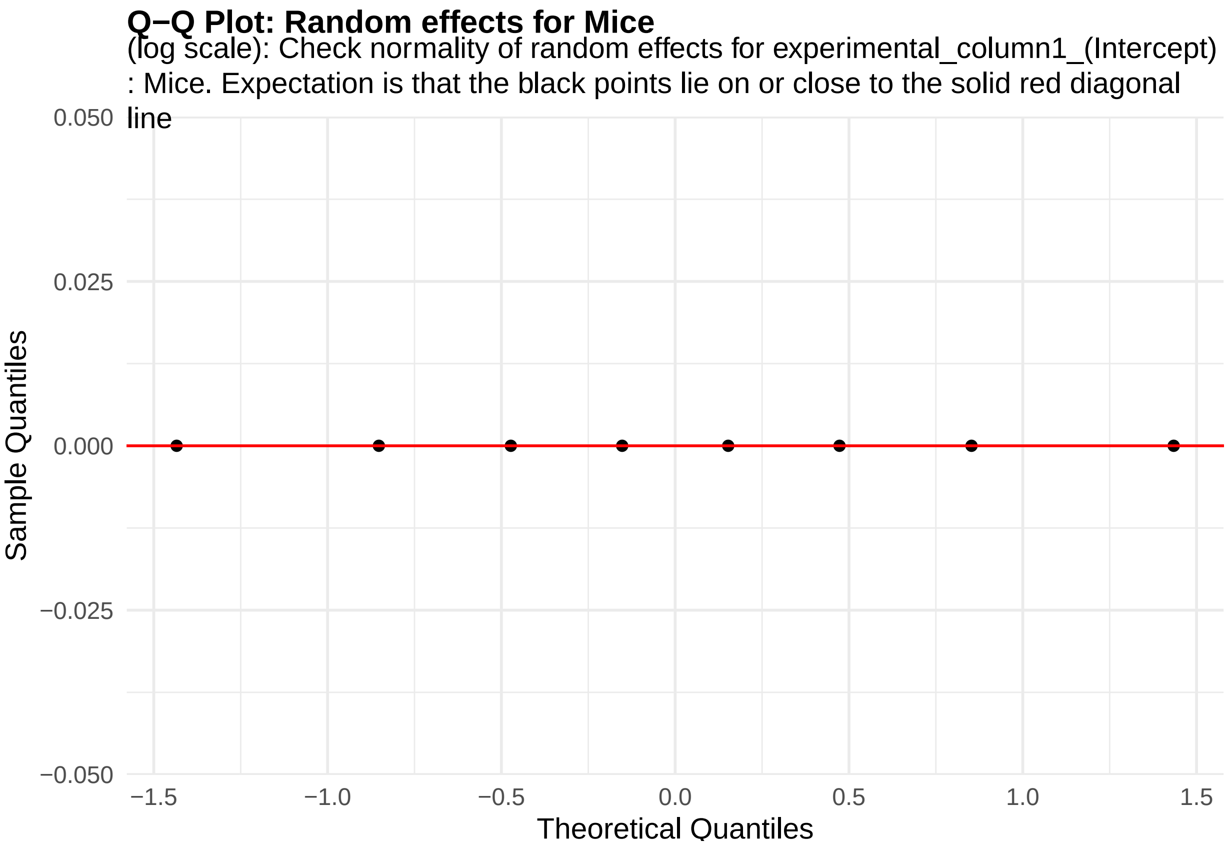


**Figure 100:** random_effects_QQ_2. Check normality of random effects for experimental_column1_(Intercept) (log scale): Mice. Expectation is that the black points lie on or close to the solid red diagonal line. Check the Normal Q-Q plots for good and bad examples here: https://library.virginia.edu/data/articles/diagnostic-plots

```
cooks_plots <- diagnose_res_update$cooks_plots$log_scale
cat("\n\n\n")
```

```
for(i in 1:length(cooks_plots)) {
  print(cooks_plots[[i]])
  cat(sprintf("\n\n\n**Figure %d:**\n\n", plot_no))
  plot_no <- plot_no + 1
}
```

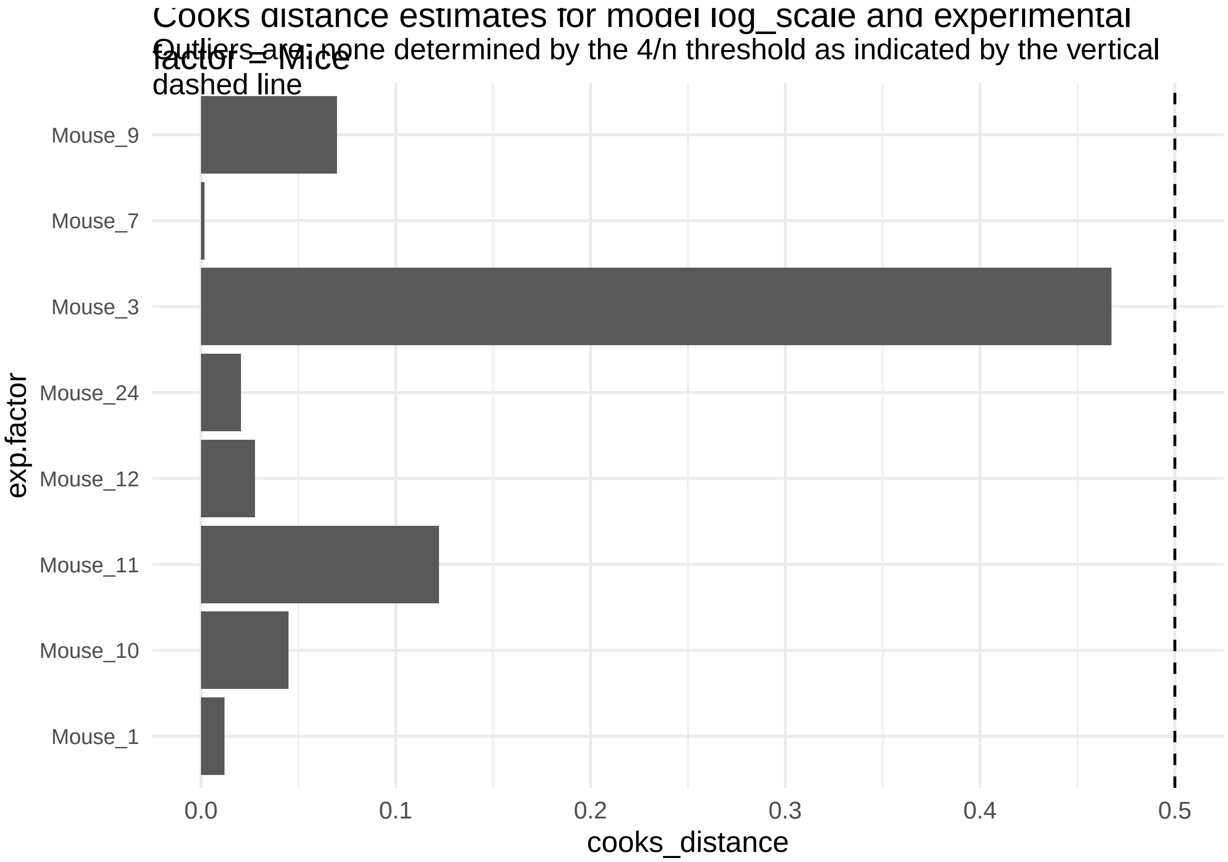
Cooks distance estimates for model log_scale and experimental factor = Mice
Outliers are: none determined by the 4/n threshold as indicated by the vertical dashed line
Mouse_9
Mouse_7
Mouse_3
Mouse_24
Mouse_12
Mouse_11
Mouse_10
Mouse_1
exp.factor
0.0
0.1
0.2
0.3
0.4
0.5
cooks_distance

**Figure 101:**

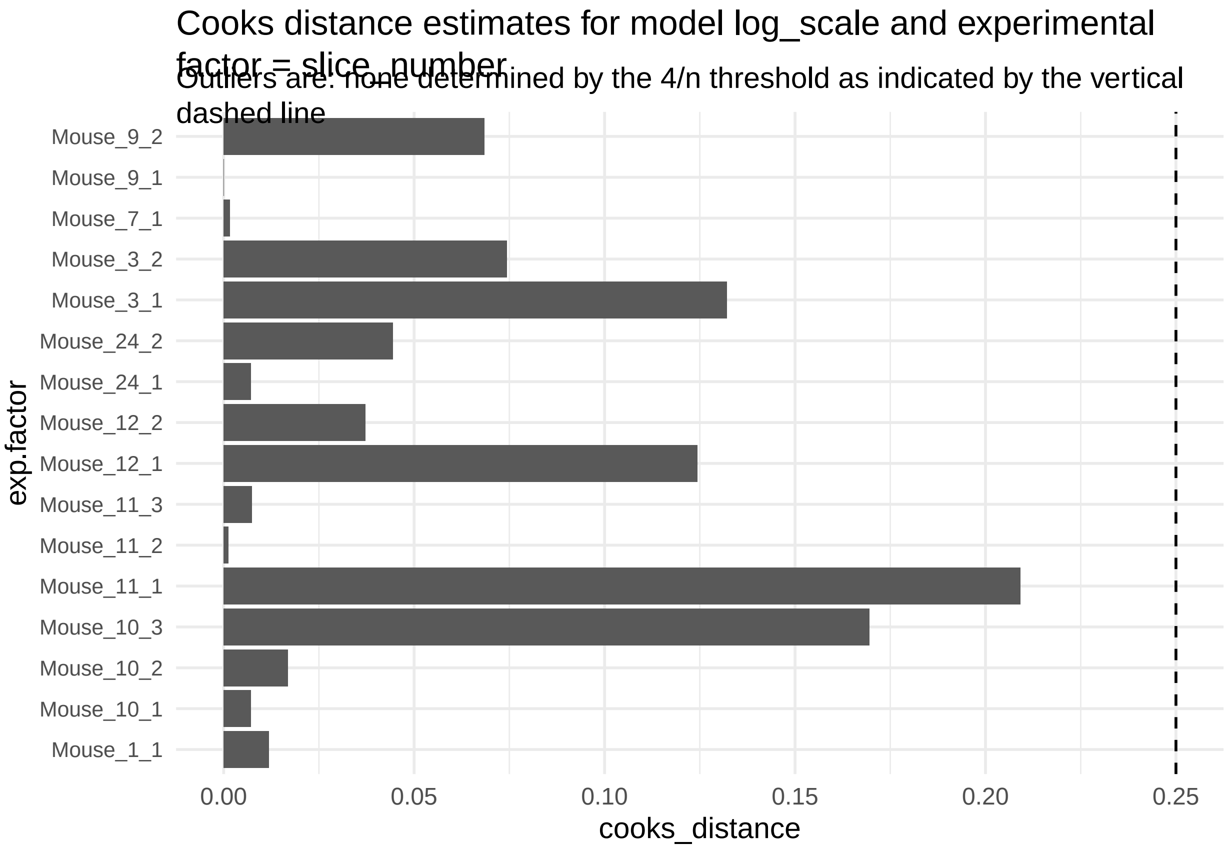


**Figure 102:**

```
print("Inferred outlier groups are")
```

[1] "Inferred outlier groups are"

```
print(diagnose_res$inferred_outlier_groups$natural_scale)
```

$Mice [1] "Mouse_2"

$slice_number [1] "Mouse_11_1"

It does not look like the log transformation resulted significantly changed how well the model assumptions were being met. We will estimate the association of genotype using the responses in their natural scale.

```
estimate_res <- getEstimatesOfInterest(data_update, design, model, print_plots = F)
print(estimate_res[[1]])
```

```
## Linear mixed model fit by REML. t-tests use Satterthwaite's method [
## lmerModLmerTest]
## Formula: response_column ~ condition_column + (1 | experimental_column1) +
##     (1 | experimental_column2)
##    Data: fixed_global_variable_data
##
## REML criterion at convergence: 355.8
##
## Scaled residuals:
##     Min      1Q  Median      3Q     Max
## -1.5894 -0.6642 -0.2666  0.6472  2.8072
##
```

```
## Random effects:
##  Groups               Name        Variance Std.Dev.
##  experimental_column2 (Intercept)  5.288   2.300
##  experimental_column1 (Intercept)  0.000   0.000
##  Residual                         51.232   7.158
## Number of obs: 53, groups:  experimental_column2, 16; experimental_column1, 8
##
## Fixed effects:
##                     Estimate Std. Error     df t value Pr(>|t|)
## (Intercept)           60.681      1.431 10.601  42.419 3.53e-13 ***
## condition_columnWT    -2.481      2.467  9.736  -1.006    0.339
## ---
## Signif. codes:  0 '***' 0.001 '**' 0.01 '*' 0.05 '.' 0.1 ' ' 1
##
## Correlation of Fixed Effects:
##             (Intr)
## cndtn_clmWT -0.580
## optimizer (nloptwrap) convergence code: 0 (OK)
## boundary (singular) fit: see help('isSingular')
```

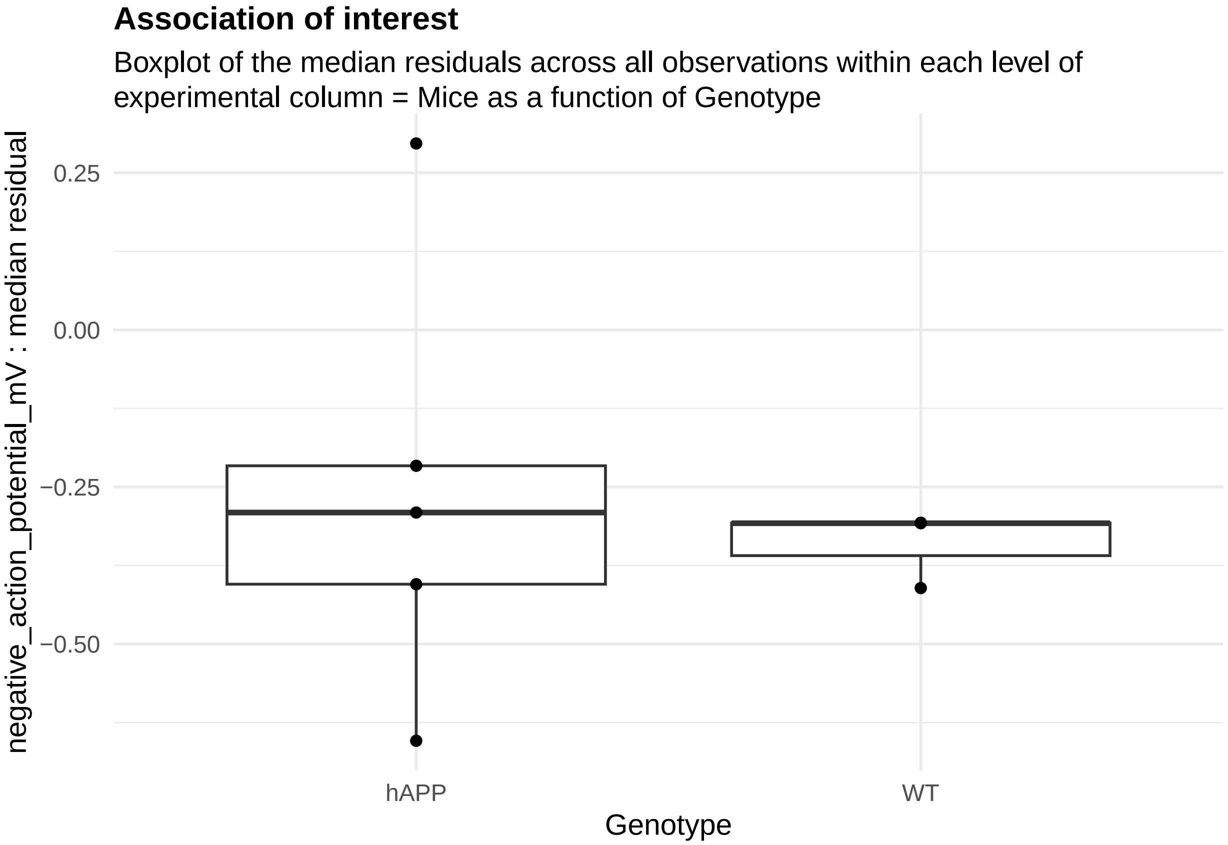


**Figure 103:**

This is not significant at all. How many mice do we need to have 80% power to estimate the observed difference?

```
power_param@target_columns <- "Mice"
power_param@effect_size <- c(7, NA, NA)
power_res <- calculatePower(data, design, model, power_param)
```

```
## [1] "effect_size must be a 3 element numeric vector"
```

NULL

**Figure 104:**

```
power_param@target_columns <- "slice_number"
power_res_slice <- calculatePower(data, design, model, power_param)
```

```
## [1] "effect_size must be a 3 element numeric vector"
```

NULL

**Figure 105:**

### 3 Understanding simulations to estimate the power

`RMeDPower2` is designed to simulate the effect of variability of the responses which could come from differences in the processing batch, plates or cell lines on the responses of interest the cell assay data (described above for exampel). There are several ways to assess the impact of different choices for the experimental variables in this package as outlined below.

First, a user can assess how increasing the number of independent experiments affects power. For example, if a user has a pilot dataset that consists of 3 experimental batches that each contain 2 plates, expanding this dataset two-fold would have the effect of simulating 6 experiments with a total of 12 plates (Figure @ref(fig:exp) ).

In the code that tests power using the example data 'plate_assay_pilot_data', this scenario can be simulated by setting 'target_columns' to 'line' and 'levels' to 1 where 'line' corresponds to the cell line. The 'condition_column' of the data corresponds to 'classification' indicating the cell state, and there are two experimental variables: 'experiment' and 'line'. Since the same cell line may appear in multiple experiments, "line" is treated as "crossed_column". "cell_size2" is used as 'response_column' for the simulation (Figure @ref(fig:Sim1)).

```
template_dir <- system.file("input_templates/understanding_simulations",
                            package = "RMeDPower2")

##load the design
design <- readDesign(file.path(template_dir, "design_cell_assay_sim1.json"))
##load the probability model
model <- readProbabilityModel(file.path(template_dir, "prob_model_sim1.json"))
##load the relevant power parameters
power_param <- readPowerParams(file.path(template_dir, "power_param_sim1.json"))

##filter out the top 5% of observations
plate_assay_pilot_data %<>% filter(cell_size2 < quantile(cell_size2, 0.95))

print(design)
```

```
## An object of class "RMeDesign"
## Slot "response_column":
## [1] "cell_size2"
##
## Slot "covariate":
## NULL
##
## Slot "condition_column":
## [1] "classification"
```

```
##
## Slot "condition_is_categorical":
## [1] TRUE
##
## Slot "experimental_columns":
## [1] "experiment" "line"
##
## Slot "crossed_columns":
## [1] "line"
##
## Slot "total_column":
## NULL
##
## Slot "outlier_alpha":
## [1] 0.05
##
## Slot "na_action":
## [1] "complete"
##
## Slot "include_interaction":
## [1] NA
##
## Slot "random_slope_variable":
## NULL
##
## Slot "covariate_is_categorical":
## [1] NA
```

```
print(model)
```

```
## An object of class "ProbabilityModel"
## Slot "error_is_non_normal":
## [1] FALSE
##
## Slot "family_p":
## NULL
```

```
print(power_param)
```

```
## An object of class "PowerParams"
## Slot "target_columns":
## [1] "experiment"
##
## Slot "power_curve":
## [1] 1
##
## Slot "nsimn":
## [1] 10
##
## Slot "levels":
## [1] 1
##
## Slot "max_size":
## NULL
##
```

```
## Slot "alpha":
## [1] 0.05
##
## Slot "breaks":
## NULL
##
## Slot "effect_size":
## NULL
##
## Slot "icc":
## NULL
```

The first step involves running a diagnosis of the data and the model.

```
diagnose_res <- diagnoseDataModel(plate_assay_pilot_data, design, model)
design_update <- design
data_update <- diagnose_res$Data_updated
print(colnames(data_update))
```

```
##  [1] "cell_size2_logTransformed" "experiment"
##  [3] "line"                      "classification"
##  [5] "cell_size1"                "cell_size2"
##  [7] "cell_size3"                "cell_size4"
##  [9] "cell_size5"                "cell_size6"
```

```
design_update@response_column <-  "cell_size2_logTransformed"
```

We can now perform sample-size estimations for the effects of increasing the number of experimental batches.

```
##calculate power
power_res <- calculatePower(data_update, design_update, model, power_param)
```

In a similar context, users can evaluate power for a given number of experiments. To do this, we assign a value corresponding to the number of experiments to max_size and set 'power_curve' to 0. We can set the 'output' variable to specify the output file name.

```
template_dir <- system.file("input_templates/understanding_simulations",
                            package = "RMeDPower2")

##load the relevant power parameters
power_param <- readPowerParams(file.path(template_dir, "power_param_sim3.json"))

print(power_param)
```

```
## An object of class "PowerParams"
## Slot "target_columns":
## [1] "experiment"
##
## Slot "power_curve":
## [1] 0
##
## Slot "nsimn":
## [1] 10
##
## Slot "levels":
## [1] 1
##
```

```
## Slot "max_size":
## [1] 15
##
## Slot "alpha":
## [1] 0.05
##
## Slot "breaks":
## NULL
##
## Slot "effect_size":
## NULL
##
## Slot "icc":
## NULL
```

```
##calculate power
power_res <- calculatePower(data_update, design_update, model, power_param)
```

```
## Power for predictor 'condition_column', (95% confidence interval):
##       80.00% (44.39, 97.48)
##
## Test: Likelihood ratio
##
## Based on 10 simulations, (0 warnings, 0 errors)
## alpha = 0.05, nrow = 12290
##
## Time elapsed: 0 h 0 m 3 s
##
## nb: result might be an observed power calculation
```

Consider the situation where a user has a specific difference of mean perimeters in mind based on prior information from related previous experiments, but the pilot data itself does not reflect the a priori assumed difference. For example, previously an experimenter found a significant difference comparing a given response between control cell lines "A", "B", "C", and cell lines from patients with a disease, "D", "E" and "F", and there was a sufficient sample size to estimate the effect size reliably. The experimenter then performs another set of experiments comparing the same disease lines ("D", "E" and "F") but this time uses different control cell lines ("G" and "H"), and the data did not have a sufficient sample size in terms of the number of experimental batches. It seems reasonable to expect that the magnitude of the associations of the response variable between the new control lines and the original disease lines would be similar to the first set of experiments. In this case, the user can then run power simulations using the known difference instead of estimating it from the pilot data. For our illustration, in the pilot data set the observed difference in mean log-transformed perimeter was 0.031 units. The power analysis shows that the power to detect this specific association between ALS status and the cell perimeter using 8 experimental batches is 72.1% with a 95% confidence interval (69.2, 74.9) at type I error rate of 0.05 reflecting what we observed in Figure 10 while if we expect the difference of means to be equal to 0.02 then the corresponding power estimate is 38.7% with a 95% confidence interval (35.7, 41.8).

```
set.seed(12345)
##observed differences of means = 0.03072
power_param@max_size <- 8
power_param@target_columns <- "experiment"
power_param@levels <- 1
power_param@power_curve <- 0
power_param@effect_size <- c(0.03072, NA, NA)
calculatePower(data_update, design_update, model, power_param)
```

```
## [1] "effect_size must be a 3 element numeric vector"
```

```
## NULL
```

```
set.seed(12345)
##asssumed difference of means = 0.02
power_param@max_size <- 8
power_param@target_columns <- "experiment"
power_param@levels <- 1
power_param@power_curve <- 0
power_param@effect_size <- c(0.02, NA, NA)
calculatePower(data_update, design_update, model, power_param)
```

```
## [1] "effect_size must be a 3 element numeric vector"
```

```
## NULL
```

We can also test the power of two experimental variables by assigning a vector of variable names to 'target_columns'.

```
power_param <- readPowerParams(file.path(template_dir,"power_param_sim5.json"))

print(power_param)
```

```
## An object of class "PowerParams"
## Slot "target_columns":
## [1] "experiment" "line"
##
## Slot "power_curve":
## [1] 1
##
## Slot "nsimn":
## [1] 10
##
## Slot "levels":
## [1] 1 0
##
## Slot "max_size":
## [1]   9 142
##
## Slot "alpha":
## [1] 0.05
##
## Slot "breaks":
## NULL
##
## Slot "effect_size":
## NULL
##
## Slot "icc":
## NULL
```

```
##calculate power
power_res <- calculatePower(data_update, design_update, model, power_param)
```

Our power simulations depend on the variance components estimated from the input dataset. However, in some cases there may not be enough data to estimate the variance component. For example, the pilot data might have only a single category for 'experimental batch', 'plate' or 'line'. When this occurs, the user needs to provide ICC values. These values can be estimated from another dataset for which the variance components are assumed to be similar to those inherent for the new response being considered in the input dataset. ICC can be estimated by taking the ratio between the variance estimates using the following formula:

$$ICC_i = \frac{V_i}{\sum_{j \in S} V_i + e}$$

where V_i represents the variance of the random effect linked to experimental variable (e.g., experimental batch, plate or cell-line) i, and epsilon represents the variance of the residual error. S represents all the modeled sources of variability of the response under consideration. We will test this scenario using the example RMeDPower_data2 with only single experiment and cell line. To test power in scenarios with varying number of experiments, we set the 'icc' parameter to 0.8, 0.05, 0.05.

```r
template_dir <- system.file("input_templates/understanding_simulations",
                            package = "RMeDPower2")

##load the design
design <- readDesign(file.path(template_dir, "design_cell_assay_sim6.json"))
##load the probability model
model <- readProbabilityModel(file.path(template_dir, "prob_model_sim6.json"))
##load the relevant power parameters
power_param <- readPowerParams(file.path(template_dir, "power_param_sim6.json"))

#print(design)
#print(model)
print(power_param)
```

```
## An object of class "PowerParams"
## Slot "target_columns":
## [1] "experiment"
##
## Slot "power_curve":
## [1] 1
##
## Slot "nsimn":
## [1] 10
##
## Slot "levels":
## [1] 1
##
## Slot "max_size":
## NULL
##
## Slot "alpha":
## [1] 0.05
##
## Slot "breaks":
## NULL
##
## Slot "effect_size":
## NULL
##
## Slot "icc":
## [1] 0.80 0.05 0.05
```

```
##calculate power
power_res <- calculatePower(plate_assay_pilot_data_wo_repeats, design, model, power_param)
```

Alternatively, a user can assess how increasing the number of plates per experiment affects power. In the case where there are two experiments that each contain two plates, the user can double the number of plates used per experiment. In this way, the user can simulate how the statistical power changes as a result of increasing the number of plates used per experiment rather than increasing the number of experimental batches (Figure @ref(fig:plate)).

In a third example, a user can examine the effect of expanding the number of cell lines within each experiment affects power (Figure @ref(fig:line)). This would capture the effect of increasing the number of cells assayed as a consequence of having more cell lines.

Alternatively, one might want to assess the effect of increasing the total number of cells by increasing the number of cells per cell line (Figure @ref(fig:lineCell)).

Tables 1-3, in addition to the figures, show the change in cell number after variable expansion in experiment, plate or cell line, especially when the cell distribution is asymmetric. A user may want to examine the power of increasing the total number of cells measured from each experimental variable per experiment. For example, if there are 12 cells per cell line on plate 1, doubling the number of cells from each plate will result in assessing the effect in twice the number of cells per cell line, even if the number of experiments and cell lines remain the same (Figure @ref(fig:plateCell)).

Table 2: Example of changes in cell counts in a simulation of experimental batch variability. (A) Original data structure and (B) changed data structure after simulation. Simulation based on level=1 keeps the same structure and increases the number of experiments. This describes the situation where the pilot study involved 2 experiments, in the first experiment two cell lines are used with 3 and 6 cells assayed, respectively, and in the second experiment only 3 cells from the first experiment are assayed.

| (A) | Cell line 1 | Cell line 2 |
|---|---|---|
| Exp 1 | 3 | 6 |
| Exp 2 | 3 | 0 |
| (B) | Cell line 1 | Cell line2 |
| Exp 1 | 3 | 6 |
| Exp 2 | 3 | 0 |
| Exp 3 | 3 | 6 |
| Exp 4 | 3 | 0 |

Table 3: Example of changes in cell counts in a simulation of plate variability. (A) Original data structure and (B) changed data structure after simulation. Simulation based on level=1 keeps the same structure and increases the number of plates.

| (A) | Cell line 1 | Cell line 2 |
|---|---|---|
| Plate 1 | 4 | 5 |
| Plate 2 | 2 | 0 |
| (B) | Cell line 1 | Cell line 2 |
| Plate 1 | 4 | 5 |
| Plate 2 | 2 | 0 |
| Plate 1’ | 4 | 5 |
| Plate 2’ | 2 | 0 |

Table 4: Example of changes in cell counts in a simulation of cell line variability. (A) Original data structure and (B) changed data structure after simulation. If level=1 is set, RMeDPower simulates new cell lines by inheriting the experimental design structure from the existing data. An example of a simulation in tabular format shows the change in the number of cells per cell line and experiment after simulation. Simulation based on level=1 keeps the same structure and increases the number of cell lines.

| | | | | | | |
|---|---|---|---|---|---|---|
| (A) | Cell line 1 | Cell line 2 | | | | |
| Exp 1 | 3 | 6 | | | | |
| Exp 2 | 3 | 0 | | | | |
| (B) | Cell line 1a | Cell line 2a | Cell line 1b | Cell line 2b | Cell line 1c | Cell line 2c |
| Exp 1 | 3 | 6 | 3 | 6 | 3 | 6 |
| Exp 2 | 3 | 0 | 3 | 0 | 3 | 0 |

Expansion results for plates and cell lines may differ if cells are asymmetrically distributed in different settings. Tables 4 and 5 show the different results users can get in an asymmetric distributed cell scenario. This type of variable expansion can be accomplished by setting ‘level=0’ in the calculate_power function.

Table 5: Example of plate-based cell count expansion simulation. (A) Original data structure and (B) changed data structure after simulation. The simulation is based on level=0 which results in increased number of cells per plate.

| | | |
|---|---|---|
| (A) | Cell line 1 | Cell line2 |
| Plate 1 | 3 | 6 |
| Plate 2 | 3 | 0 |
| (B) | Cell line 1 | Cell line 2 |
| Plate 1 | 6 | 12 |
| Plate 2 | 6 | 0 |

Table 6: Example of cell line-based cell count expansion simulation. (A) Original data structure and (B) changed data structure after simulation. An example of a simulation in tabular format shows the change in the number of cells per cell line and experiment after simulation. The simulation is based on level=0 which results in an increased number of cells per cell line.

| | | |
|---|---|---|
| (A) | Cell line 1 | Cell line 2 |
| Exp 1 | 3 | 6 |
| Exp 2 | 3 | 0 |
| (B) | Cell line 1 | Cell line 2 |
| Exp 1 | 9 | 18 |
| Exp 2 | 9 | 0 |

The type of variable expansion shown in Table 3 and 5 can be accomplished by setting ‘target_columns’ to ‘line’ and ‘levels’ to 0. In the code example, we modified the maximum number of cell lines by setting max_size to 700 (Figure @ref(fig:Sim2)).

### References

### 4 Session info

```
## R version 4.5.1 (2025-06-13)
## Platform: aarch64-apple-darwin20
## Running under: macOS Sonoma 14.7.6
##
## Matrix products: default
## BLAS:   /Library/Frameworks/R.framework/Versions/4.5-arm64/Resources/lib/libRblas.0.dylib
## LAPACK: /Library/Frameworks/R.framework/Versions/4.5-arm64/Resources/lib/libRlapack.dylib;  LAPACK ve
##
## locale:
## [1] en_US.UTF-8/en_US.UTF-8/en_US.UTF-8/C/en_US.UTF-8/en_US.UTF-8
##
## time zone: America/Los_Angeles
## tzcode source: internal
##
## attached base packages:
## [1] stats     graphics  grDevices utils     datasets  methods   base
##
## other attached packages:
##  [1] RMeDPower2_0.1.0 quantreg_6.1     SparseM_1.84-2   ggtext_0.1.2
##  [5] ggplot2_4.0.0    magrittr_2.0.4   simr_1.0.8       dplyr_1.1.4
##  [9] lme4_1.1-37      Matrix_1.7-3
##
## loaded via a namespace (and not attached):
##  [1] influence.ME_0.9-9 gtable_0.3.6       xfun_0.53
##  [4] lattice_0.22-7     numDeriv_2016.8-1.1 vctrs_0.6.5
##  [7] tools_4.5.1        Rdpack_2.6.4       generics_0.1.4
## [10] parallel_4.5.1     pbkrtest_0.5.5     tibble_3.3.0
## [13] pkgconfig_2.0.3    DHARMa_0.4.7       RColorBrewer_1.1-3
## [16] S7_0.2.0           lifecycle_1.0.4    binom_1.1-1.1
## [19] compiler_4.5.1     farver_2.1.2       stringr_1.6.0
## [22] MatrixModels_0.5-4 tinytex_0.57       lmerTest_3.1-3
## [25] carData_3.0-5      litedown_0.7       htmltools_0.5.8.1
## [28] yaml_2.3.10        Formula_1.2-5      pillar_1.11.1
## [31] car_3.1-3          nloptr_2.2.1       tidyr_1.3.1
## [34] MASS_7.3-65        reformulas_0.4.1   iterators_1.0.14
## [37] boot_1.3-31        abind_1.4-8        nlme_3.1-168
## [40] commonmark_2.0.0   tidyselect_1.2.1   digest_0.6.37
## [43] stringi_1.8.7      purrr_1.2.0        labeling_0.4.3
## [46] splines_4.5.1      fastmap_1.2.0      grid_4.5.1
## [49] RLRsim_3.1-8       cli_3.6.5          survival_3.8-3
## [52] broom_1.0.10       withr_3.0.2        scales_1.4.0
## [55] backports_1.5.0    plotrix_3.8-4      rmarkdown_2.29
## [58] png_0.1-8          evaluate_1.0.5     knitr_1.50
## [61] EnvStats_3.1.0     rbibutils_2.3      markdown_2.0
## [64] mgcv_1.9-3         rlang_1.1.6        gridtext_0.1.5
## [67] Rcpp_1.1.0         glue_1.8.0         xml2_1.4.0
## [70] jsonlite_2.0.0     rstudioapi_0.17.1  minqa_1.2.8
## [73] R6_2.6.1           plyr_1.8.9
```
